# Clonal breeding strategies to harness heterosis: insights from stochastic simulation

**DOI:** 10.1101/2022.07.01.497810

**Authors:** Marlee R. Labroo, Jeffrey B. Endelman, Dorcus C. Gemenet, Christian R. Werner, R. Chris Gaynor, Giovanny E. Covarrubias-Pazaran

## Abstract

To produce genetic gain, hybrid crop breeding can change the additive as well as dominance genetic value of populations, which can lead to utilization of heterosis. A common hybrid breeding strategy is reciprocal recurrent selection (RRS), in which parents of hybrids are typically recycled within pools based on general combining ability (GCA). However, the relative performance of RRS and other possible breeding strategies have not been thoroughly compared. RRS can have relatively increased costs and longer cycle lengths which reduce genetic gain, but these are sometimes outweighed by its ability to harness heterosis due to dominance and increase genetic gain. Here, we used stochastic simulation to compare gain per unit cost of various clonal breeding strategies with different amounts of population inbreeding depression and heterosis due to dominance, relative cycle lengths, time horizons, estimation methods, selection intensities, and ploidy levels. In diploids with phenotypic selection at high intensity, whether RRS was the optimal breeding strategy depended on the initial population heterosis. However, in diploids with rapid cycling genomic selection at high intensity, RRS was the optimal breeding strategy after 50 years over almost all amounts of initial population heterosis under the study assumptions. RRS required more population heterosis to outperform other strategies as its relative cycle length increased and as selection intensity decreased. Use of diploid fully inbred parents vs. outbred parents with RRS typically did not affect genetic gain. In autopolyploids, RRS typically was not beneficial regardless of the amount of population inbreeding depression.

**Key Message:** Reciprocal recurrent selection sometimes increases genetic gain per unit cost in clonal diploids with heterosis due to dominance, but it typically does not benefit autopolyploids.

## Introduction

Hybrid breeding may achieve genetic gain by changing the additive as well as dominance genetic value of populations over breeding cycles. Hybrid breeding strategies are widely used in diploid, inbred-hybrid crops such as maize (*Zea mays* L.) and sorghum (*Sorghum bicolor* L.), but an assessment of these strategies’ genetic gain per unit cost over a wide range of dominance genetic architectures has not yet been conducted (Duvick, 2005; Longin et al., 2012). Additionally, breeding strategies to cost-effectively utilize dominance in clonal breeding programs, particularly autopolyploids, have not been fully explored or widely initiated (Diaz et al., 2021; Ceballos et al., 2020; Darkwa et al., 2020; Batte et al, 2020; Lindhout et al., 2018). Dominance has been observed via inbreeding depression and heterosis in economically important traits of various clonal species, such as fresh yield in diploid cassava (*Manihot esculenta*) and autohexaploid sweetpotato (*Ipomoea batatas*; Ceballos et al., 2015; Diaz et al., 2021). However, clonal crops have differences from inbred-hybrid crops which could affect the optimal breeding strategy to achieve genetic gain when heterosis due to dominance is present. Therefore, we compare breeding strategies in model clonal crop breeding programs by stochastic simulation with various genetic architectures of heterosis due to dominance.

The first consideration in clonal hybrid breeding is that clonal crops may be diploid, as are cassava and white yam (*Dioscorea rotundata*), but are often various degrees of autopolyploid, as in potato (*Solanum tuberosum*), sweetpotato, sugarcane (*Saccharum* spp.), and banana (*Musa* spp.). The quantitative genetics of autopolyploids are an active area of research, and the increased transmissibility of dominance value in autopolyploids with random mating compared to diploids suggests that breeding strategies to harness heterosis due to dominance may differ between diploids and autopolyploids (Amadeu et al., 2020). The second consideration is that the multiplication ratio of clonal crops may be low; for example, maize typically produces around 200 seeds per cross (200:1), but white yam currently produces around 4 to 8 propagules per plant (4:1 to 8:1; Aighewi et al., 2015). Therefore, hybrid breeding strategies which require two stages of crossing may face penalties in species with low multiplication ratios due to the additional time needed for multiplication. Finally, clonal crop genotypes can be routinely reproduced identically by asexual reproduction rather than inbreeding to full homozygosity (McKey et al., 2010). Many clonal crops are difficult or impossible to self and display severe inbreeding depression; some populations lose viability even without complete homozygosity (Lebot, 2019). It has long been recognized that hybrid breeding does not require fully inbred parental genotypes to harness heterosis; rather, fully inbred parents are required to identically reproduce hybrid genotypes in inbred-hybrid crops that cannot be clonally propagated (Schnell, 1961; Lamkey & Edwards, 1999). However, occasional concern that clonal breeding would benefit from fully inbred lines remains (Ceballos et al., 2015; Powell et al., 2020).

The key reason to pursue a hybrid breeding strategy is to utilize heterosis and avoid inbreeding depression due to dominance while also increasing additive value. The mean additive value of traits can be increased by increasing the frequency of favorable alleles, but for traits with both additive and dominance gene action, there is a breeding opportunity to increase mean total genetic value by also maintaining or increasing frequency of heterozygous genotypes. Fundamentally, dominance value (*d*) refers to deviation of heterozygote genetic value from mean homozygote value at a locus (Falconer & Mackay, 1996). For evolutionary reasons, dominance may tend to be positive in the direction of fitness— i.e., across loci which exhibit dominance, heterozygote value is often greater than mean homozygote value on average (Lynch & Walsh, 1998; Manna et al., 2011; Yang et al., 2017).

In traits of crops that do not exhibit dominance, selection on individual value with random mating increases the mean genetic value of populations, because the frequency of favorable alleles can be increased without regard for their transitory allocation into homozygous or heterozygous genotypes in the next breeding cycle (Hallauer & Darrah, 1985). Each unit of increase in the frequency of a favorable allele produces linear increase in mean genetic value. In traits with adequate dominance, the allocation of alleles into heterozygous genotypes nonlinearly affects mean genetic value (Schnell, 1961). Maintaining or increasing the frequency of heterozygous genotypes that exhibit dominance increases mean genetic value because these heterozygous genotypes have disproportionately higher values than the less fit, lower-value homozygous genotype (Wei & Van der Steen, 1991). At a locus with dominance, the lower-value homozygous genotype is often referred to as a deleterious recessive genotype, and the decrease in population fitness due to deleterious recessive loci is sometimes called genetic load (Fisher, 1935; Muller, 1950; Falconer & Mackay, 1996). Ultimately, fixing the favorable homozygous genotype leads to higher mean genetic value than maintaining heterozygous genotypes in absence of complete dominance or overdominance—so maximizing heterosis is suboptimal with incomplete dominance— but if the favorable homozygote is not fixed it is prudent to avoid the deleterious recessive state (Rembe et al., 2019). Even in absence of true overdominance, linkage disequilibrium of dominant alleles in breeding populations can lead to pseudooverdominance (Jones, 1917). If the haplotypes are not broken over the breeding time horizon, they prevent stacking of favorable alleles and effectively behave as an overdominant locus (Bingham, 1998; Werner et al., 2020).

The biologically dominant gene action of individual alleles of complex traits leads to population-wide heterosis and inbreeding depression (Hallauer et al., 2010; Lamkey & Edwards, 1999; Labroo et al., 2021). Here, we borrow from the framework of heterosis and inbreeding depression presented by Falconer & Mackay (1996) and Lamkey & Edwards (1999). As defined by Falconer & Mackay (1996), inbreeding depression is the difference in value between any population at Hardy-Weinberg equilibrium (P_HWE_) and the population if fully inbred (homozygous; *P*_*I*_), or *P*_*I*_ - *P*_*HWE*_. Heterosis can then be considered the opposite of inbreeding depression due to dominance, *P*_*HWE*_ *-P*_*I*_. Lamkey & Edwards (1999) further partition heterosis into values which are relevant to RRS programs. Panmictic heterosis is the difference in the inter-pool hybrid value (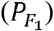) to the mean of the intra-pool genotypes at Hardy-Weinberg equilibrium (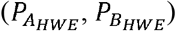), or 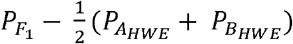. Baseline heterosis refers to the difference in value of the intra-pool genotypes at Hardy-Weinberg equilibrium to the value of the intra-pool genotypes if fully inbred to homozygosity (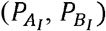) or 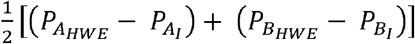. Inbred-midparent heterosis is the sum of panmictic and baseline heterosis. Lamkey & Edwards (1999) specifically define inbreeding depression as the reversal of baseline heterosis, but here we consider the more general definition of Falconer & Mackay (1996). We acknowledge that heterosis due to epistasis is possible, and that heterosis due to epistasis is not the reversal of inbreeding depression, but we do not consider epistasis in this study (Lynch, 1991; Lynch & Walsh, 1998).

As stated, increasing favorable allele frequencies can increase the additive value of populations. Recurrent selection (RS) is a breeding strategy which increases the frequency of favorable alleles (Hallauer et al., 2010). In RS, a single pool of genotypes is formed. The genotypes are evaluated, and the best genotypes are selected. The selected genotypes are then randomly intermated to restart the breeding cycle, which concentrates favorable alleles in the next generation. However, with random mating in a single pool, it is challenging to increase the frequency of heterozygotes beyond 0.5, because Hardy-Weinberg equilibrium is nearly constantly restored by random mating relative to the previous generation (Falconer & Mackay, 1996). Therefore, when traits have appreciable dominance, reciprocal recurrent selection (RRS) can be a viable alternative strategy to RS (Comstock et al., 1949; Hallauer et al., 2010). In RRS, germplasm is split into at least two pools. Within each pool, intra-pool genotypes may or may not be fully inbred. Intra-pool genotypes also may or may not be evaluated for their per-se performance. Next, the intra-pool genotypes are crossed to genotypes of the opposing pool to form single-cross inter-pool F_1_ hybrids; typically, a sample of intra-pool genotypes is used because the number of all possible crosses becomes impractically large. The inter-pool hybrids are evaluated. Then, intra-pool parents of hybrids are usually selected based on estimates of their average inter-pool performance in F_1_ hybrids, or general combining ability (GCA; Comstock et al., 1949; Schnell, 1961). The two pools remain strictly separated with no mixing of pools during recycling, and over breeding cycles, this process leads to the formation of heterotic pools (Duvick et al., 2004). Heterotic pools arise because selection on GCA not only increases the frequency of favorable alleles, but also drives and drifts apart the frequencies of alleles between pools, particularly those which exhibit dominance (Rembe et al., 2019). Upon inter-pool crossing, this difference in allele frequency produces an excess of heterozygous genotypes in the F_1_ hybrids compared to the frequency of heterozygous genotypes in the parent pools (Lamkey & Edwards, 1999). The excess of heterozygosity leads to population-wide heterosis as excess dominance value is expressed in the inter-pool hybrids over the intra-pool parents. This panmictic heterosis occurs regardless of whether the intra-pool genotypes are fully inbred. If the intra-pool genotypes are inbred, upon inter-pool crossing both panmictic heterosis and baseline heterosis are observed in their hybrids, as heterozygosity exceeds not only the diverged pools if they were outbred but also the fully inbred lines.

Despite the widespread popularity of RRS (e.g. in maize breeding), additional assessment of its efficiency is needed to inform decision-making in diverse crops. In absence of heterosis, or even with low amounts of heterosis, RRS is thought to be less efficient than RS in improving mean genetic value of breeding populations because RRS usually requires a longer cycle length (*L*; Longin et al., 2014). RRS also usually has higher costs per genotype generated than RS because RRS requires maintaining separate pools of germplasm and evaluating both intra- and inter-pool material (Longin et al., 2014). However, in the presence of adequate dominance, RRS is thought to be more efficient in producing genetic gain than RS because RRS prevents expression of deleterious homozygous recessive states in F_1_ hybrids by increasing the frequency their heterozygous genotypes. In other words, RRS harnesses and exploits heterosis due to dominance, which partly entails avoiding inbreeding depression due to dominance.

To avoid the challenges of RRS while still making some use of heterosis, animal breeders have developed intermediate strategies (Leroy et al., 2016; Swan & Kinghorn, 1992). Of these strategies, the most relevant to challenges in plant breeding may be terminal crossing (Leroy et al., 2016). Terminal crossing can be thought of as RS within two pools, which are subsequently crossed to obtain panmictic heterosis via drift. In terminal crossing, germplasm is divided into two pools. Within each pool, intra-pool genotypes are evaluated for per-se performance. Then, intra-pool genotypes are “terminally” crossed to the opposing pool to form single-cross inter-pool F_1_ hybrids, and the inter-pool hybrids are evaluated for use as products. However, intra-pool parents are selected and recycled as parents using estimates of their intra-pool per-se performance rather than their GCA. As in RRS, the two pools remain strictly separated during recycling. Terminal crossing has a shorter cycle length than RRS because parents can be recycled without waiting for their hybrid progeny phenotypes, and terminal crossing can be logistically simpler than RRS because testcrossing is not necessary. As mentioned, terminal crossing can also exploit some panmictic heterosis because allele frequencies within pools come to diverge by drift. However, terminal crossing builds less panmictic heterosis than RRS when dominance is present because it relies on drift and does not actively select for divergence between pools as would GCA.

The use of genomic selection (GS) to decrease cycle length can increase the competitiveness of RRS compared to other strategies, especially to establish new hybrid breeding programs (Kinghorn et al., 2010; Rembe et al., 2019). Reciprocal recurrent genomic selection can achieve cycle lengths equal to one-pool recurrent genomic selection and two-pool terminal crossing with genomic selection because parents can be recycled on estimates of their value using their relatives’ phenotypes in a genomic prediction model rather than estimates using the parent’s phenotypes (Kinghorn et al., 2010; Powell et al., 2020). Therefore, in all strategies, parents can be recycled as soon as they can be genotyped and predicted accurately rather than as soon as they can be phenotyped accurately, which is the case with phenotypic selection (PS).

A recently developed strategy to address dominance is cross performance, particularly genomic prediction of cross performance (Werner et al., 2020; Wolfe et al., 2021). In genomic prediction of cross performance, a single pool of genotypes is formed. The genotypes and phenotypes are evaluated and used to generate a genomic prediction model which typically includes both additive and dominance effects. Then, the predicted effects are used to calculate the mean performance of all possible crosses in the pool, and the best crosses are selected. Finally, the selected crosses are made to restart the breeding cycle. Key concepts with cross performance are that mating is non-random in a single pool and that the parental selection units are the crosses rather than the individuals. Non-random mating allows combinations of alleles within a locus (i.e. genotypes) to be “cut-and-paste” from parents into progeny, so more heterozygosity and thus more dominance value is maintained than with random mating. In the presence of dominance, genomic prediction of cross performance has been demonstrated to outperform selection on genomic estimated breeding value with random mating in a single pool (Werner et al., 2020). However, the various possible genomics-assisted hybrid breeding strategies have not been compared previously.

Finally, the long-term benefit and short-term cost of controlling the inbreeding rate in breeding populations is well understood, particularly with use of pedigree selection or GS (Woolliams et al., 2015). However, it is unknown whether the relative performance of hybrid breeding strategies reacts to different degrees of inbreeding control. We contend that inbreeding control can be viewed as a method to manage inbreeding depression in a population, as demonstrated by Fernández et al. (2021). The relative efficiencies of various breeding strategies to address inbreeding depression may differ depending on the inbreeding rate, which is explored indirectly here via the selection intensity. Inbreeding is caused by selection and drift over breeding cycles, which lead to overrepresentation of homozygous genotypes in breeding generations compared to the base population at Hardy-Weinberg equilibrium. Even if populations are at Hardy-Weinberg equilibrium in terms of genotype frequencies, and thus not inbred *per se*, they may still be inbred relative to the base population. Inbreeding due to concentration or fixation of favorable alleles, which can increase overall genetic value, is desirable. However, inbreeding due to drift can increase the frequency of unfavorable alleles and their homozygotes inadvertently. Inbreeding control attempts to limit inbreeding due to drift and thus can prevent inbreeding depression. This is because inbreeding control prevents random loss of heterozygosity which decreases mean genetic value in the presence of directional dominance. Of course, inbreeding control also limits drift of allele frequencies in favorable directions, which often leads to short-term costs. Inbreeding control also informs long-term comparisons of breeding strategies. In its absence, different strategies may completely deplete genetic variance at different timepoints, with no further gain, and long-term comparison is simply a record of these different timepoints. The optimal or acceptable inbreeding rate fundamentally depends on the time horizon of a breeding pipeline (Moeinizade et al., 2019). Different hybrid breeding strategies may have different performance at different time horizons, so inbreeding control may be needed to prevent exhaustion of genetic variance and reveal these differences.

In summary, several possible breeding strategies to improve traits with heterosis and inbreeding depression due to dominance exist. We shall now proceed to their comparison. We consider how various amounts of inbreeding depression and heterosis in a population affect breeding strategy efficiencies across ploidies. We test phenotypic strategy efficiencies for species with a juvenility period (i.e. delayed flowering) and low multiplication ratio. We explore the impact of intra-pool evaluation in RRS programs, as well as the impact of intra-pool doubled haploid development.

## Materials and Methods

Stochastic simulations were conducted in the R 4.0.4 computing environment with the package AlphaSimR 1.0.1 on the International Maize and Wheat Improvement Center High-Performance Computing Cluster and the University of Wisconsin-Madison Center For High Throughput Computing (R Core Team, 2021; Gaynor et al., 2021). The general procedure was that 180 starting populations with different genetic architectures were simulated, then combinations of breeding strategies, selection intensities, and estimation methods were applied to each population for 100 breeding cycles in ten replicates (Fig. 1). The responses were then measured with variously assumed cycle lengths.

**Figure 1.**
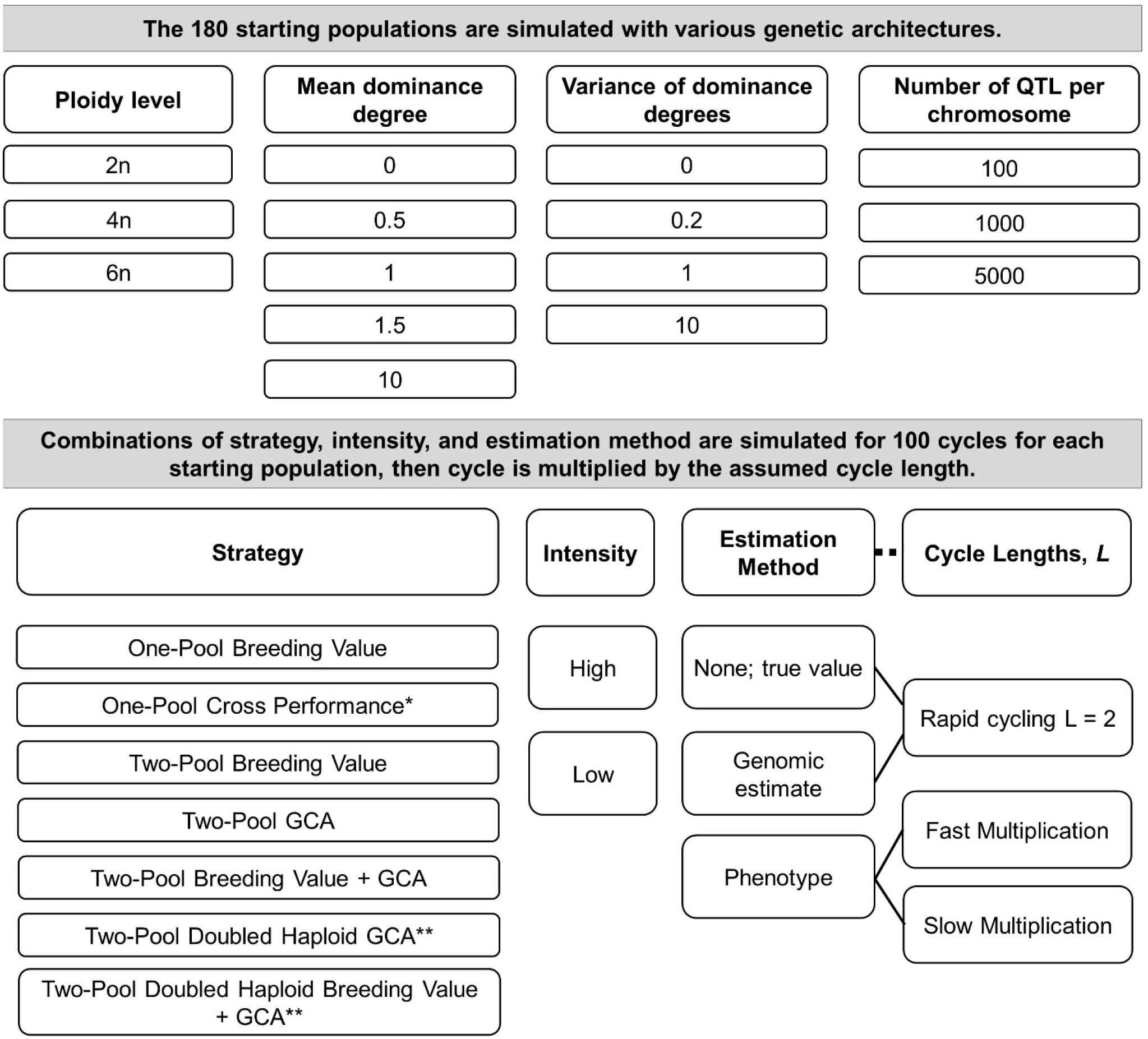
Overview of the study methods. Populations at three ploidy levels with varied amounts of population heterosis were generated by simulating all combinations of the ploidy level, number of QTL, mean dominance degrees, and variance of dominance degrees shown. Linkage disequilibrium was not controlled. For further details of these parameters’ relationship to inbreeding depression and heterosis, please see Gaynor et al., 2018. After simulating the 180 starting populations, a combination of breeding strategy, selection intensity, and estimation method was run on each population, except that strategies with doubled haploids were only run for ploidy = 2 (**). Because multiple cohorts per cycle were not simulated, cycle length was varied by multiplying cycle number by the appropriate value and not by running an independent simulation (dashed line). The combination of strategy, intensity, estimation method, and cycle lengths defined a scenario. All combinations of the scenario factors were assessed, except that the cycle lengths depended on the estimation method (solid lines) and a phenotypic estimate of One-Pool Cross Performance was not considered (*). Cycle lengths (*L*) by strategy and estimation method are given in Table 1.

### Genetic architecture simulation

The following steps were common to all scenarios. A genome with haploid n = 10 chromosomes was simulated using the AlphaSimR *runMacs()* command, which calls the Markovian Coalescent Simulator of Chen et al. (2009). The “*GENERIC*” species history was used, which implied starting effective population size (*N*_*e*_) of 100 * ploidy / 2, following the scaling recommendations of Arnold et al. (2012), and a mutation rate of 2.5 * 10^−8^ mutations per base pair. Following the genome simulation, a founder population of 100 non-inbred hermaphroditic individuals was drawn. A single AD trait with additive and dominance effects was simulated with a starting mean genetic value of zero and additive genetic variance of one using the *addTraitAD()* command. The *useVarA* option was set to *TRUE*, so the starting additive genetic variance in the base population was one for all scenarios, but the dominance variance and thus total genetic variance varied depending on the dominance parameters. Although some types of epistasis can contribute to inbreeding depression and heterosis, epistasis was not considered in this study. We also did not consider environment or genotype x environment effects to reduce the complexity of the study. We assumed no historic population split, which could affect the relative efficiency of the strategies (Lamkey & Edwards, 1999).

To create trait genetic architectures for each scenario, all combinations of the following factors and their levels were simulated: number of quantitative trait loci (QTL) per chromosome, *nQtlPerChr*, of 100, 1000, or 5000; mean dominance degree, *meanDD*, of 0, 0.5, 1, 1.5, or 10; variance of the dominance degrees, *varDD*, of 0, 0.2, 1, or 10; and, ploidy of 2, 4, or 6 (Fig. 1; Supplemental File 1). The methods of simulating allelic effects in AlphaSimR are described in the vignette “Traits in AlphaSimR”; as such, polyploid values were assigned assuming digenic dominance interactions only (Gaynor, 2021; Gallais, 2003). Varying the number of QTL, mean dominance degree, variance of the dominance degrees, and ploidy led to 180 populations (3 * 5 * 4* 3) with varied amounts of initial population heterosis (H_0_) as well as varied starting dominance and total variance, all of which were recorded (Supplemental Fig. 1; Gaynor et al., 2018).

H_0_ was the difference in the starting population at Hardy-Weinberg equilibrium from the starting population if fully inbred to homozygosity; it was divided by the starting genetic standard deviation to allow comparison across populations with traits at different scales. This measure of heterosis is not named in the framework of Lamkey & Edwards (1999), but it corresponds to the reversal of inbreeding depression as defined by Falconer & Mackay (1996). With all else equal, the amount of H_0_ increases as the mean dominance degree and the square root of the number of QTL increase and decreases as the variance of the dominance degrees increases; however, the effect of the variance of the dominance degrees is relatively smaller (Supplemental Fig. 1; Gaynor et al., 2018). We did not control linkage disequilibrium, which also affects H_0_, so simulating populations with identical parameters as in this study may lead to slightly different H_0_ as their linkage disequilibrium varies (Gaynor et al., 2018). Occasional negative H_0_ was observed in architectures with *meanDD* = 0 and *varDD* > 0 due to random sampling of dominance degrees, which sometimes led to negative directional dominance in the starting population and higher mean values of inbred than outbred genotypes. Each single trait modeled can be interpreted as representing an index of quantitative traits.

### Breeding scenarios

Each simulation was initiated by drawing 40 individuals from the same founder population with a given genetic architecture for each of ten replicates. In other words, founder populations were not varied within genetic architectures, and stochasticity within architectures was only due to Mendelian sampling and (at times) random phenotypic error. As such, there was more stochasticity across genetic architectures—which used different founder populations and traits—than within genetic architectures. For simulations with two pools, the 40 individuals were randomly split into two pools of 20 (Cowling et al., 2020). Then, a combination of strategy, selection intensity, and estimation method was applied for 100 cycles. Responses were subsequently interpreted with variously assumed cycle lengths. A scenario was defined as a combination of strategy, estimation method, selection intensity, and assumed relative cycle lengths (Fig. 1). Most combinations of the following were assessed: a strategy of One-Pool Breeding Value, One-Pool Predicted Cross Performance, Two-Pool Breeding Value, Two-Pool GCA, or Two-Pool Breeding Value + GCA; an estimation method of phenotypic value, genomic estimated value, or none (true value); and, high or low selection intensity (Fig. 1). For ploidy = 2 only, we considered two additional selection strategies to address inbred-hybrid crops: Two-Pool Doubled Haploid GCA and Two-Pool Doubled Haploid Breeding Value + True GCA. Two-Pool Breeding Value referred to a terminal crossing program. Scenarios with a phenotypic estimation method and the One-Pool Predicted Cross Performance strategy were not considered; although phenotypic cross performance can be estimated as the mean of the parental phenotypes, this scenario was too computationally intensive with phenotypic program sizes used.

Phenotypes in the study referred to single phenotypic values per entry with a fixed error variance and an initial broad-sense heritability of 0.5, which represent replicated phenotypes. The broad sense heritability of the phenotypes subsequently changed with genetic variance over cycles. The phenotypic estimate of value referred to these single phenotypic values, which were used for selection, though for Two-Pool GCA the single phenotypic records were used to calculate GCA.

Strategy cycle length was assumed to depend on the estimation method. Strategies which used true values or genomic estimates were assumed to have a cycle length of two, which was considered a realistic rapid-cycling length. Some rapid-cycling GS programs may achieve a one-season cycle length, but this is uncommon due to practical constraints (Gaynor et al., 2017). Phenotypic strategies were considered to have different cycle lengths depending on whether fast or slow multiplication was possible. Scenarios with slow multiplication were also assumed to have slow flowering, as occurs in white yam (A. Amele, pers. comm.). Fast multiplication indicated that adequate material for phenotypic evaluation and crossing was available in the season following crossing, and slow multiplication implied adequate material was available after two seasons following crossing. Doubled haploid production was assumed to require one season. All cycle lengths under all assumed constraints are reported in Table 1.

We assumed that a single cohort and breeding stage occurred per season, although typical programs may run multiple cohorts at different stages in parallel per season (Covarrubias-Pazaran et al., 2021). As such, to modify the cycle length, the cycle numbers for a given strategy, estimation method, and intensity were multiplied by the appropriate value. For example, the PS scenarios with fast and slow multiplication were obtained from the same simulations, and fast and slow multiplication cycle lengths were imposed by multiplying the cycle number by the strategy cycle length. We assumed that both phenotypic and genotypic information became available post-flowering. Genotypic information was obtained from a simulated SNP-chip with 1000 markers; the number of markers was not varied across genetic architectures. If genomic estimated values were used, the training set for two-pool programs was comprised of the 2,000 most recently evaluated inter-pool individuals, and the training set for one-pool programs was comprised of the 2,000 most recently evaluated intra-pool individuals. To control resources across strategies, we varied program size by decreasing the number of progeny per cross first, then decreasing the number of crosses if necessary. We assumed that the costs of making crosses and growing out non-evaluated plots were negligible. The cost of evaluation plots was assumed to be equal across strategies. For further comparisons, we defined all costs in terms of evaluation plots. We assumed the cost of generating a doubled haploid line was three times the cost of an evaluation plot. We assumed that the cost of phenotyping an individual was equal to the cost of genotyping an individual. With use of outbred intra-pool parents, genotyping both intra-pool parents and their inter-pool segregating progeny was necessary. In the doubled haploid scenarios, we assumed that both intra- and inter-pool genotypes were genotyped, even though the inter-pool progeny genotypes could be inferred from their doubled haploid parents under the assumed cycle lengths. Scenarios which used true values were identical in size to scenarios with genomic estimated values; cost is not a realistic consideration to obtain true values, and the true value scenarios were used to consider a situation with perfect accuracy.

A description of each strategy follows. For conciseness, the program sizes are represented by variables, and the values of variables for each scenario are given in Supplemental Table 1. Parents were randomly mated in the first cycle, and in all subsequent cycles a crossing plan conferring maximum avoidance of inbreeding was used (Kimura & Crow, 1963).

- One-Pool Breeding Value: The parents are made into *x* crosses with *y* progeny per cross, totaling z individuals. The *z* progeny are phenotyped. Then, 2 individuals per family (cross) are selected using the estimate of value for the scenario strategy. The cycle restarts with the selected individuals. Genomic estimates were made from a directional dominance model fit on the training population of intra-pool genotypes using the *RRBLUP_D()* function (Xiang et al., 2016). Code is in Supplemental Files 2—7.
- Two-Pool Breeding Value: Within each pool, the parents are made into *x* crosses with *y* progeny per cross, totaling z intra-pool progeny per pool. The *z* intra-pool progeny are phenotyped. From each pool, two individuals are then selected randomly. For both pools, all *z* intra-pool progeny per pool are crossed to both individuals selected from the opposing pool, and each inter-pool cross produces one progeny, creating *w* inter-pool progeny. The inter-pool progeny are phenotyped. Within each pool, 2 individuals per family (cross) are selected on the scenario surrogate of intra-pool breeding value. The cycle restarts with the selected individuals. Genomic estimates were made from a directional dominance model fit on the training population of inter-pool genotypes using the *RRBLUP_D()* function. We did not explore use of other models or use of intra-pool information in the training set. Code is in Supplemental Files 8—13.
- One-Pool Predicted Cross Performance: The parents are made into *x* crosses with *y* progeny per cross, totaling z individuals. The *z* progeny are evaluated. The expected mean progeny value for each possible biparental cross is calculated from the expected genotype distribution for each locus under the assumption that gametes pair independently and that the frequency of these gametes follows a binomial distribution. In the case of autopolyploids, these assumptions are consistent with strict bivalent pairing of chromosomes in meiosis, which is the assumption used in this study. True expected mean progeny value is calculated using true QTL and their effects, whereas genomic estimated expected mean progeny value is using SNP markers and their estimated effects (https://github.com/gaynorr/QuantGenResources/blob/main/CalcCrossMeans.cpp). To conduct maximum avoidance with cross performance, the pairs of families (crosses) which satisfy a maximum avoidance of inbreeding plan are identified. Within those pairs of families, the values of inter-family crosses of their individual members are calculated. Then the two best crosses from each set of paired families are selected. The cycle restarts with the selected crosses. Genomic estimates were made from a directional dominance model fit on the training population of intra-pool genotypes using the *RRBLUP_D()* function. Code is in Supplemental Files 14—17.
- Two-Pool GCA: Within each pool, the parents are made into *x* crosses with *y* progeny per cross, totaling z intra-pool progeny per pool. From each pool, two individuals are selected randomly. For both pools, all *z* intra-pool progeny per pool are crossed to both individuals selected from the opposing pool, and each inter-pool cross produces one progeny, creating *w* inter-pool progeny. The inter-pool progeny are phenotyped. Then, within each pool, 2 individuals per family (cross) are selected as parents on GCA. The cycle restarts with the selected individuals. Genomic estimates of GCA were made from a model with parent-specific allelic additive effects fit on the training population of inter-pool genotypes using the *RRBLUP_GCA()* function. Code is in Supplemental Files 18—23.
- Two-Pool Breeding Value + GCA: these strategies have the same structure as Two-Pool GCA, except that the intra-pool progeny are evaluated before testcrossing. The top ∼75% of individuals per family (cross) are selected on the appropriate estimate of breeding value according to scenario, and only the selected individuals are used in testcrossing. With use of genomic estimated values, intra-pool breeding values were estimated with use of a directional dominance model, *RRBLUP_D()*, on a training set of inter-pool genotypes. Intra-pool GCA were estimated with the same training set but the *RRBLUP_GCA()* model. Code is in Supplemental Files 24— 29.
- Two-Pool Doubled Haploid GCA: these strategies have the same structure as Two-Pool GCA, except that all intra-pool progeny were used to create a single doubled haploid line in the season before testcrossing. Code is in Supplemental Files 30—35.
- Two-Pool Doubled Haploid Breeding Value + GCA: these strategies had the same structure as Two-Pool GCA, except that all intra-pool progeny were used to create a single doubled haploid line in the season following intra-pool crossing. The intra-pool doubled haploid lines were evaluated before testcrossing and the top ∼75% of individuals per family (cross) were selected on the appropriate estimate of breeding value according to scenario, and only these selected individuals were used in testcrossing. With use of genomic estimated values, intra-pool breeding values were estimated with use of a directional dominance model, *RRBLUP_D()*, on a training set of inter-pool genotypes. Intra-pool GCA was estimated with the same training set but the *RRBLUP_GCA()* model. Code is in Supplemental Files 36—41.

### Responses and analysis

The responses reported were as follows:

- For one-pool scenarios, genetic gain was the mean genetic value at a given timepoint in the intra-pool genotypes following their evaluation (*G*_*t*_) minus the mean genetic value of the founder population (*G*_0_), or *G*_*t*_ - *G*_0_. For the two-pool scenarios, the method was the same except the inter-pool genotypes were used. This allowed comparison of genetic gain in the product pools of both scenarios. Genetic gain was also reported for the intra-pool genotypes in the Two-Pool GCA, Two-Pool Doubled Haploid GCA, Two-Pool Breeding Value + GCA, and Two-Pool Doubled Haploid Breeding Value + GCA scenarios. Genetic gain was divided by the initial population genetic standard deviation.
- Mean additive value and mean dominance value were reported at a given cycle in the respective product pools for one-pool and two-pool scenarios and scaled to the starting population genetic standard deviation.
- Inbreeding depression was reported for the product pools of the scenarios as previously described (Falconer & Mackay, 1996).
- For scenarios with selection on true values, the genomic inbreeding coefficient *f* was reported for the product pools relative to their initial populations based on a genomic (**G**) additive relationship matrix (Van Raden 2008; Method 1) with allele frequencies from the initial population. For diploids, the mean diagonal of **G** equals 1 + *f* (Powell et al., 2010; Endelman and Jannink, 2012). The more general relationship for ploidy □is that the mean diagonal of **G** equals 1 + (□–1)*f* (Gallais 2003). Please note that the inbreeding coefficient was used only to compare inbreeding for identical strategies at high vs. low selection intensity and requires subtlety in interpretation across populations with different levels of homozygosity due to structure.
- Panmictic heterosis was reported for the two-pool strategies as previously described (Lamkey & Edwards, 1999).

We wish to highlight that the methods used do not permit meaningful comparisons of absolute or scaled values across ploidies. For example, observing that a breeding program for autohexaploids leads to greater mean genetic value than a diploid at a given cycle does not necessarily imply that more gain is possible in autohexaploids.

Responses were reported for all scenarios after 15 and 50 years of breeding, at which timepoints genetic variance was non-zero for all scenarios. Genetic variance was later exhausted at different timepoints among scenarios. Responses were also reported for PS at the same cycle numbers (8 and 25) as GS and true scenarios. This was done to demonstrate the effect of using GS as an estimation method, without using it to reduce cycle length, on the relative performance of PS and GS. For clarity, results were grouped by the question of interest. The core strategies to explore the optimal breeding strategy across H_0_ were One-Pool Breeding Value, One-Pool Cross Performance, Two-Pool Breeding Value, and Two-Pool GCA. The core strategies were also used to explore the optimal estimation method—i.e. genomic estimated or phenotypic—under the experimental assumptions. The non-core strategies, Two-Pool Breeding Value + GCA, Two-Pool GCA, Two-Pool Doubled Haploid GCA, and Two-Pool Doubled Haploid Breeding Value + GCA, were used to assess whether combined selection on intra-pool breeding value and inter-pool GCA increased gain with or without fully inbred intra-pool parents. The non-core strategies were also used to assess whether use of fully inbred diploid intra-pool parents increased the rate of genetic gain.

To analyze and plot the results, each response at the timepoint of interest (15 years, 50 years, or 25 cycles) for the questions of interest (core or non-core strategies) was linearly modeled in base R as follows:

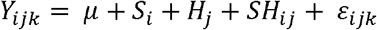

Where *Y*_*ijk*_ was the response value for the *i*^th^ scenario *S*, the *j*^th^ H_0_ value *H*, their *ij*^th^ interaction *SH*, and the *ijk*^th^ error of the simulation replicate. The scenario of a response was the combination of strategy, estimation method, selection intensity, and assumed cycle length. All effects were assumed to be fixed, normally distributed, and independently distributed. The coefficient of determination (*R*^*2*^) value, slope, slope standard error, intercept, and intercept standard error was recorded for each regression (Supplemental File 42). The regressions, the 95% confidence interval of their predicted means, and, at times, raw data points were plotted using the R package ggplot2 (Wickham, 2011). The intersections of the regressions which occurred within the surveyed H_0_ values and, when possible, their standard errors were also calculated (Supplemental File 43). The standard errors of the intersections were estimated by maximum likelihood with the R package nlme and used to calculate the 95% confidence interval of the intersection (whuber, 2020; Pinheiro et al., 2017). In accordance with recent guidelines of the statistical community, significance testing was not conducted and confidence intervals were interpreted (Wasserstein & Lazar, 2016; Alexander & Davis, 2022). We assumed regressions could be meaningfully distinguished at a given value of H_0_ if their confidence intervals did not overlap.

Only selected responses are plotted in the figures and supplementary figures, but plots of all responses for all scenarios in the study are available for reference in Supplemental File 44.

## Results

### Genetic gain in the core strategies

The relative performance of the core strategies depended on H_0_, the time horizon, the selection intensity in the program, the relative cycle lengths among strategies, the estimation method, ploidy level, and their interactions. Typically, the comparative advantage of Two-Pool GCA increased with increased H_0_, time horizon, and selection intensity, as well as with use of GS, but it decreased with increased ploidy level or increased cycle length.

With use of GS in the clonal diploids, at high intensity Two-Pool GCA was the best strategy after 15 years if H_0_ was greater than 9.3, and One-Pool Breeding Value or One-Pool Cross Performance was the best strategy if H_0_ was lower (Fig. 2). After 50 years, Two-Pool GCA was the best predicted strategy at all positive H_0_ values, and its relative advantage increased as H_0_ increased (Fig. 2). In contrast, at low intensity, one-pool strategies were always better than Two-Pool GCA after 15 years (Fig. 2). After 50 years at low intensity Two-Pool GCA only outperformed One-Pool Breeding Value if H_0_ was greater than 17.7, a substantially greater amount of H_0_ than at high intensity (Fig. 2). High intensity programs had greater genetic gain than low intensity programs on average, but low intensity one pool strategies outperformed high intensity one pool strategies if H_0_ was relatively high (Fig. 2). (Of course, two-pool strategies still outperformed the best one-pool strategy over the range at which low intensity one pool strategies outperformed high intensity one pool strategies.)

**Figure 2.**
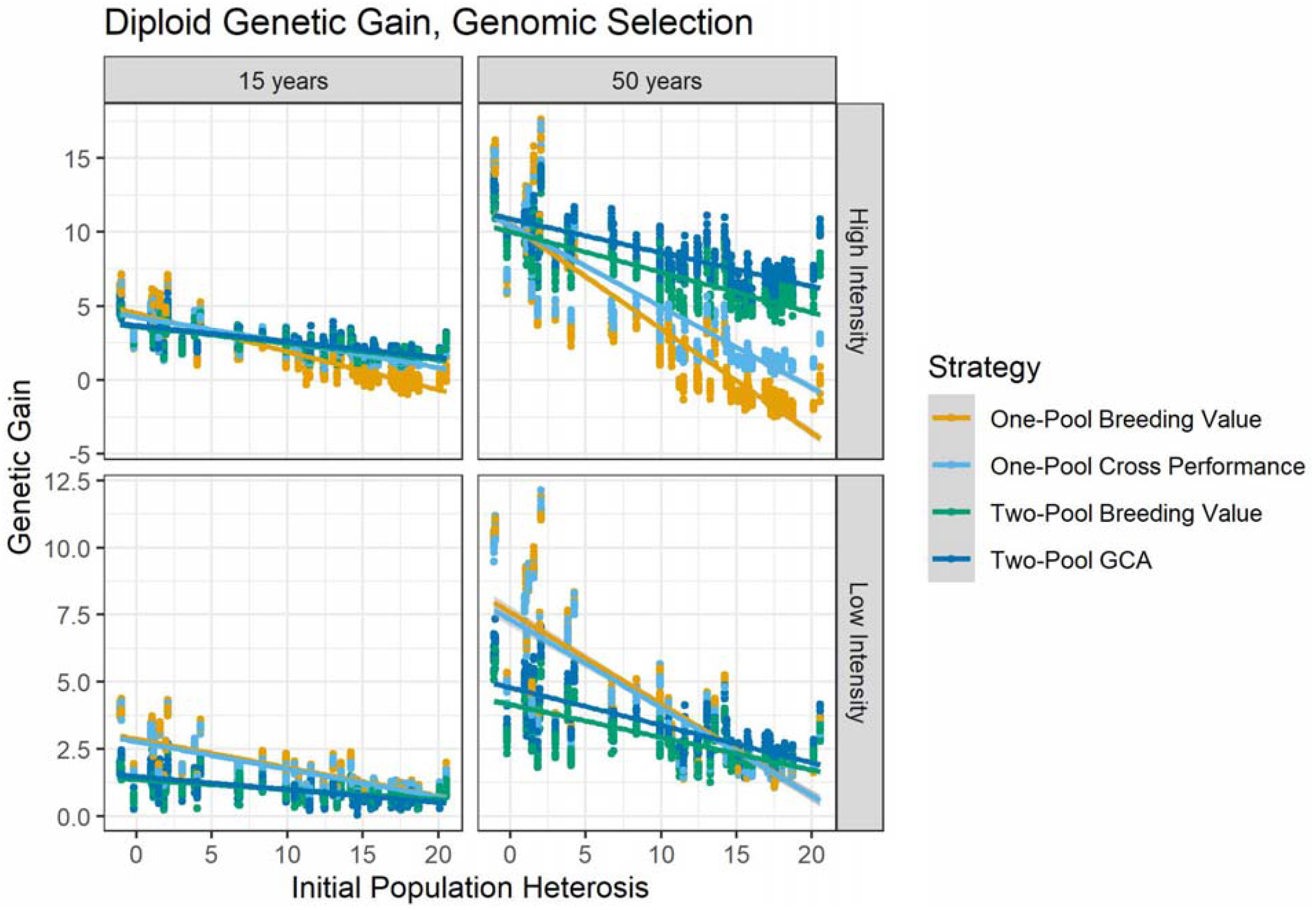
Genetic gain in diploids after 15 and 50 years with use of GS regressed on breeding scenario, initial population heterosis, H_0_, and their interaction. Colored lines indicate regressions by breeding strategy with GS and cycle length 2, and grey bands indicate the standard error of the predicted means. Dots indicate raw data points and dot color indicates strategy as in the lines. At high intensity after 15 years, the differences among strategies were marginal, and after 50 years Two-Pool GCA provided the most gain over almost all H_0_ values. At low intensity, two-pool strategies required more H_0_ and time to outperform the one-pool strategies than at high intensity.

With use of PS and fast multiplication in clonal diploids, Two-Pool GCA was not the best strategy after 15 years at any H_0_ value (Supplemental Fig. 2). After 50 years, it required H_0_ greater than 13.9 to outperform other strategies, and the amount of overperformance was relatively less than with GS (Supplemental Fig. 3). With PS and slow multiplication, Two-Pool GCA never outperformed other strategies over the time horizons surveyed (Supplemental Fig. 2,Supplemental Fig. 3).

With use of GS in the clonal autopolyploids, Two-Pool GCA showed fewer advantages than in diploids, and One-Pool Breeding Value or One-Pool Cross Performance were typically better strategies (Fig. 3). At high intensity after 15 years, One-Pool Breeding Value or One-Pool Cross Performance were the best strategies for both autotetraploids and autohexaploids. One-Pool Cross Performance was the better strategy at high H_0_, and One-Pool Breeding Value was the better strategy at low H_0_. After 50 years at high intensity in the autotetraploids, One-Pool Breeding Value or One-Pool Cross Performance provided the most gain if H_0_ was less than or equal to 31.0 ± 2.4; if H_0_ was greater, Two-Pool GCA or Two-Pool Breeding Value provided the most gain, but the advantages were small (Fig. 3). In the autohexaploids, the same strategy pattern was apparent but the intersection occurred at H_0_ of 61.7 ± 5.0. At low selection intensity, One-Pool Breeding Value or One-Pool Cross Performance provided the most gain at both timepoints for both autotetraploids and autohexaploids (Fig. 3).

**Figure 3.**
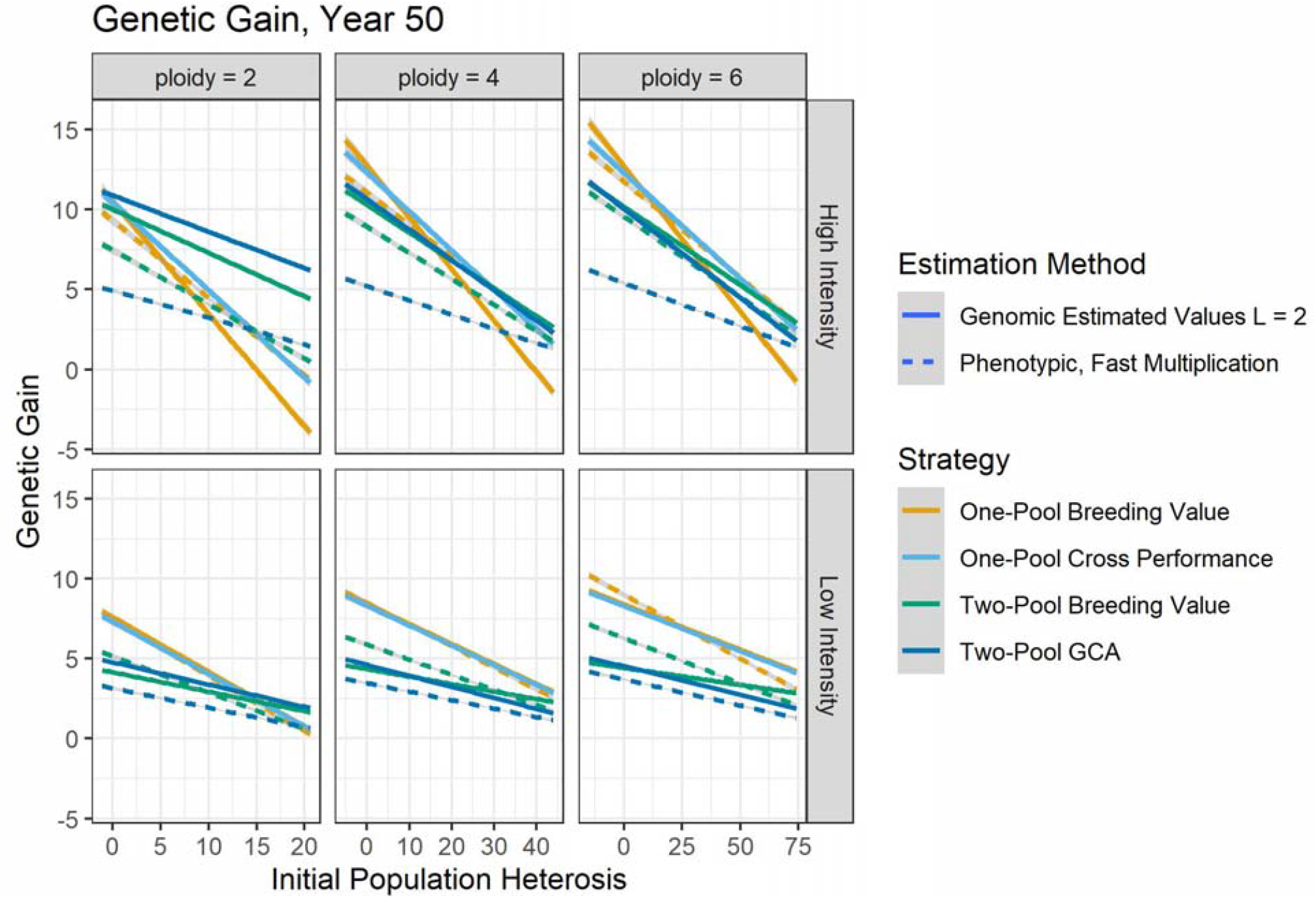
Genetic gain for each ploidy level after 50 years of breeding with use of genomic and phenotypic selection and various strategies as a function of H_0_, breeding scenario, and their interaction. Line color indicates strategy, and grey bands indicated the standard error of the predicted mean. Line type indicates estimation method with the accompanying set of cycle lengths (*L*). In clonal diploids at high intensity, genomic selection on Two-Pool GCA is the best strategy regardless of H_0_, but this advantage is not apparent in the autopolyploids. Instead, the autopolyploids tend to benefit from one-pool strategies. Use of GS typically increases or does not change genetic gain at high intensity, particularly for diploids. It is not appropriate to compare amounts of genetic gain across ploidy levels.

For the clonal diploids, use of the best GS strategy increased genetic gain compared to the best PS strategy with fast multiplication after 50 years (Fig. 3). If GS was not used to reduce cycle length, and all strategies were compared at 25 cycles, then at small values of H_0_, the best PS strategy produced more gain and the best GS strategy produced more gain with greater H_0_ (Supplemental Fig. 3). This indicates the dependency of the relative performance of GS and PS on their relative cycle length as well as H_0_. For the clonal autopolyploids, at high intensity the best GS strategy was better than or equal to the best PS strategy (Fig. 3). The advantage of GS decreased as H_0_ decreased. At low intensity in autotetraploids, the best GS strategy was indistinguishable from the best PS strategy. At low intensity in autohexaploids, PS outperformed GS if H_0_ was low, and vice versa if H_0_ was high.

Less absolute genetic gain was observed as H_0_ increased (Fig. 2—3). Based on the slopes of the regression lines, one-pool strategies were more sensitive to H_0_ than two-pool strategies (Supplemental File 42; Fig. 2—3). As genetic gain increased due either to a longer time horizon or higher intensity, the sensitivity of genetic gain to H_0_ also increased.

### Additive and dominance value in the core strategies

Regardless of ploidy level, strategy, selection intensity, or timepoint, the regression of additive value on H_0_ produced a negative slope, while the regression of dominance value on H_0_ produced a positive slope (Supplementary File 42; Supplemental Fig. 5—8). If no dominance was simulated, then both dominance value and H_0_ were always zero. In general, one-pool strategies produced more additive value than two-pool strategies regardless of ploidy, timepoint, or intensity (Supplemental Fig. 5, Supplemental Fig. 7). In diploids, Two-Pool GCA produced more dominance value than other strategies at high but not low intensity and as timepoint increased, particularly with use of GS (Supplemental Fig. 6). In autopolyploids, there was typically little difference in dominance value among strategies (Supplemental Fig. 8).

### Inbreeding coefficient with true values for the core strategies

The inbreeding coefficient was recorded for scenarios with an estimation method of none (true values) only. Within a given ploidy level and timepoint, the regression of inbreeding coefficient on H_0_ for each strategy differed depending on the selection intensity (Supplemental File 42). After 15 years, regardless of strategy and ploidy, strategies had higher inbreeding coefficients with high selection intensity and lower inbreeding coefficients with low selection intensity across H_0_ values (Supplemental Fig. 9). After 50 years, in diploids One-Pool Cross Performance and Two-Pool Breeding Value had higher inbreeding coefficients with high selection intensity and lower inbreeding coefficients with low intensity, but crossover was observed for Two-Pool GCA and One-Pool Breeding Value (Supplemental Fig. 10). For both, high intensity tended to lead to higher inbreeding coefficients when H_0_ was smaller, but low intensity led to high inbreeding coefficients with higher H_0_. In autopolyploids, after 50 years all strategies tended to lead to higher inbreeding coefficients under high selection intensity than low selection intensity (Supplemental Fig. 10). The difference in the inbreeding coefficient by intensity was less in autopolyploids than diploids.

### Inbreeding depression with the core selection strategies

Subsequent to the simulation of an initial amount of inbreeding depression, the amount of inbreeding depression in the population potentially could change as allele frequencies changed due to selection and other forces. Regardless of ploidy level, strategy, selection intensity, or timepoint, the regression of population inbreeding depression on H_0_ produced a positive slope as expected, given that populations with greater H_0_ sustained greater amounts of inbreeding depression regardless of breeding cycle (Supplemental File 42). In general, with comparisons at the same number of cycles, the amount of inbreeding depression for a given ploidy level, estimation method, intensity, and timepoint did not dramatically differ by strategy although some differences were detected (Supplemental Fig. 11—12). Greater reduction of population inbreeding depression was not associated with greater genetic gain.

### Panmictic heterosis with the core selection strategies

Panmictic heterosis was zero for the one-pool strategies by definition. For the two-pool strategies, the regression of panmictic heterosis on H_0_ produced positive slopes, indicating that the amount of panmictic heterosis strategies built increased with the amount of H_0_ regardless of ploidy (Supplemental File 42; Fig. 4). Two-Pool GCA tended to build more panmictic heterosis than Two-Pool Breeding Value, and their relative difference decreased as H_0_ decreased. In general, Two-Pool GCA built increasingly more panmictic heterosis than Two-Pool Breeding Value as selection intensity and timepoint increased. However, the difference in panmictic heterosis between Two-Pool GCA and Two-Pool Breeding Value decreased as ploidy level increased.

**Figure 4.**
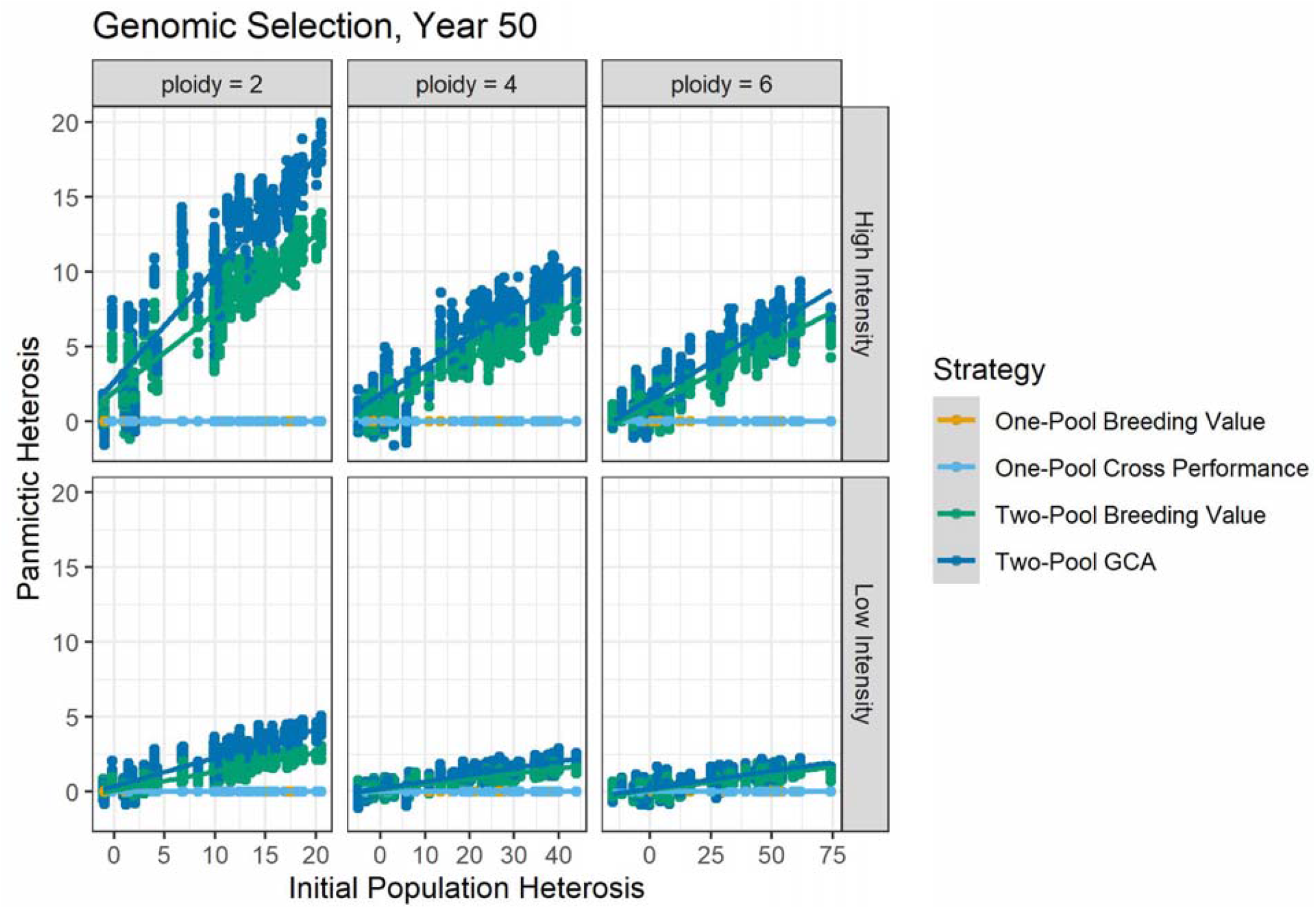
Panmictic heterosis for each ploidy as a function of initial population heterosis, H_0_, after 50 years of breeding with each strategy and use of genomic selection with cycle length of 2. Colored lines indicate strategy and grey bands indicate the standard error of their predicted means. Colored dots indicate the corresponding strategy raw data points. Two-Pool GCA tended to build more panmictic heterosis that Two-Pool Breeding Value, especially in diploids at high intensity, because Two-Pool GCA leads to increased divergence of allele frequencies between pools by selection. Two-Pool Breeding Value builds panmictic heterosis primarily by drift, and one-pool strategies do not build panmictic heterosis. Two-pool strategies lead to clear panmictic heterosis in autopolyploids even though neither two-pool strategy was optimal in terms of genetic gain. Comparisons of absolute values across ploidies are not likely to be biologically relevant.

### Breeding Value + GCA strategies

Strategies in which intra-pool evaluation was used to advance genotypes to intra-pool crossing, Two-Pool Breeding Value + GCA and Two-Pool Doubled Haploid Breeding Value + GCA, showed increased genetic gain with PS and unchanged genetic gain with GS compared to strategies without intra-pool evaluation, Two-Pool GCA and Two-Pool Doubled Haploid GCA (Supplemental Fig. 13). The same pattern was observed across ploidies for Two-Pool Breeding Value + GCA and Two-Pool GCA. More interestingly, with selection on GCA, intra-pool genetic value tended to decrease over cycles (compared to the initial intra-pool genotypes) regardless of whether intra-pool evaluation was used at high H_0_ (Supplemental Fig. 14). However, intra-pool genetic value tended to increase over cycles at low H_0_. Intra-pool evaluation increased intra-pool genetic values compared to its absence with PS and fast multiplication without use of doubled haploids, but intra-pool evaluation had no effect on intra-pool genetic values with GS or with PS and use of doubled haploids (Supplemental Fig. 14).

### Doubled Haploid GCA strategies

The use of intra-pool fully inbred lines generally led to unchanged genetic gain after 50 years with GS, but in some cases increased genetic gain with PS. (Supplemental Fig. 13). With PS, Two-Pool Doubled Haploid GCA increased gain compared to Two-Pool GCA but had similar performance to Two-Pool Breeding Value + GCA and Two-Pool Doubled Haploid Breeding Value + GCA (Supplemental Fig. 13). Intra-pool fully inbred lines typically had lower mean genetic values than intra-pool outbred clones in both the short and long term (Supplemental Fig. 14). The difference in doubled haploid and outbred intra-pool genotypes was greater as H_0_ increased as they suffered additional inbreeding depression (Supplemental Fig. 14). Population inbreeding depression typically did not differ between Two-Pool Doubled Haploid GCA and Two-Pool GCA, nor between Two-Pool Doubled Haploid Breeding Value + GCA and Two-Pool Breeding Value + GCA (Supplemental Fig. 15).

## Discussion

Although Two-Pool GCA sometimes provided substantially greater rates of genetic gain per unit cost than other strategies in clonal diploids, its relative performance depended on heterosis and inbreeding depression due to dominance in the trait population, the time horizon, the selection intensity in the program, the relative achievable cycle lengths among strategies, the estimation method, ploidy level, and their interactions. The use of GS rather than PS drastically increased the competitiveness of Two-Pool GCA, indicating that GS unlocks novel opportunities to utilize heterosis. Increased selection intensity increased the relative performance of Two-Pool GCA to other strategies, perhaps indicating that Two-Pool GCA is more competitive at higher inbreeding rates. In typical diploid programs with high selection intensities, if Two-Pool GCA could achieve equal cycle lengths as other strategies, then Two-Pool GCA tended to increase the rate of genetic gain per unit cost at lower amounts of H_0_ than if Two-Pool GCA required a longer cycle length. However, in autopolyploids, Two-Pool GCA usually did not increase the rate of genetic gain compared to One-Pool Breeding Value or One-Pool Cross Performance. Autopolyploid Two-Pool GCA tended to provide an advantage in genetic gain at higher values of H_0_ than in diploids, if at all, and the amount of relative increase was less than in diploids. As in other studies, the use of GS tended to increase gain compared to PS likely due to increased accuracy, faster inbreeding, and decreased cycle length across H_0_; use of GS to reduce of the cycle length was a determining factor in whether it outperformed PS at the heritabilities used (Powell et al., 2020; Gaynor et al., 2017; Heslot et al., 2015; Heffner et al., 2010; Longin et al., 2015).

### Clonal diploids

In clonal diploids, Two-Pool GCA appeared to outperform other strategies in some conditions because of its exceptional ability to increase the dominance value of F_1_ hybrid populations, as well as the additive value. Fundamentally, this is because use of Two-Pool GCA can increase not only the frequency of favorable alleles but also the frequency of heterozygote genotypes relative to Hardy-Weinberg equilibrium in F_1_ hybrids of two pools, leading to panmictic heterosis (Lamkey & Edwards, 1999). The latter is achieved by selection on GCA, which differs from breeding values in a single pool because dominance value *(d*) is weighted by allele frequencies in the opposite pool (Schnell, 1965; Rembe et al., 2019). Selection on GCA drives apart allele frequencies between pools, which results in a sustained increase in heterozygosity and therefore dominance value in the F_1_ hybrids. Although both additive and dominance value are transmissible with selection on breeding value and random mating in a single pool, the frequency of heterozygotes is limited by Hardy-Weinberg equilibrium, which is overcome by non-random mating in two pools (Hardy, 1908; Weinberg, 1908). Reducing population heterosis (inbreeding depression) was neither required nor a strategic advantage to make genetic gain, and at longer time horizons genetic variance was exhausted due to drift and selection well before any changes in population heterosis or inbreeding depression were observed. Generally, the advantages of Two-Pool GCA in clonal diploids increase as:

- the amount of H_0_ due to dominance increases, because ability to increase dominance value becomes relatively more important
- the time horizon increases, because formation of heterotic pools with diverged allele frequencies requires selection over breeding cycles
- its relative cycle length to the other strategies decreases, because cycle length directly impacts the rate of genetic gain, and Two-Pool GCA has a necessarily longer cycle length than the other strategies with PS but not GS
- the selection intensity increases, perhaps because higher selection intensities lead to more inbreeding which lead to greater reductions in heterozygosity due to selection and drift which are better alleviated by GCA compared to other strategies, or because higher selection intensities more rapidly drove apart allele frequencies between pools
- its relative cost to the other strategies decreases; however, we did not investigate different levels of relative cost among strategies because this was demonstrated by Longin et al. (2014) and its particulars are highly program-specific.

The amount of trait population heterosis can be estimated experimentally in breeding populations, but it is typically unknown. Better methods and increased effort to estimate heterosis in breeding programs would be useful to inform decision-making. However, for clonal diploids which can utilize rapid-cycling GS, the benefit of Two-Pool GCA was robust to H_0_ under the study assumptions. Two-Pool GCA provided the most gain over most H_0_ values and timepoints surveyed, and if H_0_ was relatively low Two-Pool GCA only modestly decreased gain in the short term. Programs for which Two-Pool GCA is relatively more expensive than assumed here may require more H_0_ to reap its benefit. In contrast to GS, moving to Two-Pool GCA without adequate population heterosis or time presented a risk of decreased genetic gain for phenotypic programs. Interestingly, clonal crops using PS with a low multiplication ratio never benefited from Two-Pool GCA over the time horizons in the study, highlighting this consideration for clonal species and the usefulness of efforts to increase the multiplication ratio (Aighewi et al., 2015). It would be useful to confirm the optimal GS strategies for programs with low multiplication ratios, particularly with multiple cohorts running in parallel per season. Please see Supplemental File 5 for discussion of Two-Pool Breeding Value and One-Pool Cross Performance in diploids, which may be useful for programs which cannot transition to Two-Pool GCA.

In applied diploid inbred-hybrid RRS programs of seed crops, intra-pool genotypes are often first selected as parents of hybrids on their per-se value (Lee & Tracy, 2009). In clonal crops with relatively lower multiplication ratios, increased performance of intra-pool parents may not drastically increase hybrid propagule or seed production, so it was unclear whether resource allocation to intra-pool evaluation is efficient. For the costs and proportions of individuals advanced assumed in the study, we observed that a round of intra-pool advancement on breeding value before intra-pool recycling on GCA typically increased genetic gain with PS or did not change the rate of genetic gain with GS in the inter-pool hybrids. Intra-pool evaluation led to a shift from dominance to additive gain compared to forgoing intra-pool evaluation. As such, breeders likely have some flexibility in whether to conduct intra-pool evaluation. For example, with multiple traits, it is common to cull intra-pool parents for markers and highly heritable traits; unless negative genetic correlations are present in the trait index, this decision likely would not decrease genetic gain for inter-pool traits, assuming it does not increase cycle length. For the GS scenarios here, it was likely suboptimal to predict intra-pool breeding values from a training set of inter-pool individuals, and predicting intra-pool breeding values from intra-pool individuals may increase genetic gain.

Interestingly, the effect of recycling on GCA on intra-pool mean value over cycles depended on H_0_: it tended to decrease intra-pool value as H_0_ increased but increase intra-pool value as H_0_ decreased. In absence of dominance, intra-pool breeding value is equal to GCA, so intra-pool genotypes selected for GCA are nearly the same as those which would be selected on breeding value at low H_0_ (Rembe et al., 2019). This likely led to increases in intra-pool genetic value. As dominance increases, and as allele frequencies differ between pools, the values of intra-pool breeding value and GCA diverge. At high H_0_, selection on GCA led the parental pools to suffer inbreeding depression as they were driven to homozygous states, thus decreasing their value over breeding cycles. Conducting intra-pool advancement on breeding value sometimes slightly increased intra-pool parents’ value compared to forgoing intra-pool evaluation. However, at the proportion of individuals advanced (75%), intra-pool selection did not prevent decrease in intra-pool value when population heterosis was high. In practice, if population heterosis is high and it is necessary to maintain or increase intra-pool value with Two-Pool GCA, it may be necessary to select intra-pool parents more stringently on their breeding values or even to recycle intra-pool parents on an index of intra-pool breeding value and GCA (Longin et al., 2006).

Another concern in clonal diploids is whether RRS programs benefit from using fully inbred parents, as is done in other species. We did not observe substantial increases in genetic gain with use of inbred parents in RRS, especially with intra-pool evaluation. With all else equal, it is expected that inbreeding depression (loss of baseline heterosis) suffered in the intra-pool parents is fully reversed in the inter-pool hybrids, as well as the addition of the panmictic heterosis value, so intra-pool inbreeding is unnecessary to harness heterosis. The cost and time to generate inbred lines are likely higher than assumed in our study, given that doubled haploid technologies do not exist for most clonal species.

Furthermore, the simulated inbred line values may correspond to total non-viability in some species or populations, especially those with high population inbreeding depression. It has been proposed that use of inbred parents could enable seed systems in clonal crops and reduce the cost of propagation, the time and cost required to transport clones across national borders, and the spread of disease (McKey et al., 2010; Ceballos et al., 2015). These are worthy considerations that are considered externalities in the current study, but they are completely independent of the use of RRS and could equally be availed in one-pool strategies. Programs considering line development should thoroughly assess their germplasm’s tolerance of full inbreeding as well as the tradeoffs in time and resources needed for line development.

### Clonal autopolyploids

In contrast to clonal diploids, Two-Pool GCA typically did not outperform other strategies in clonal autopolyploids. Instead, One-Pool Breeding Value or One-Pool Cross Performance was the safest option depending on H_0_. A larger range of H_0_ values were considered in autopolyploids than diploids; RRS did not benefit autopolyploids at the same and some greater amounts of H_0_ which benefited diploids. This is likely because autopolyploids inherit multiple chromosome copies per gamete, and therefore autopolyploids sustain greater heterozygosity across all gametes, genotypes, and matings at segregating loci even in response to selection on One-Pool Breeding Value (Supplemental Fig. 16; Bartlett & Haldane, 1934; Bever & Felber, 1992). The relative advantage of Two-Pool GCA in diploids is due to its ability to increase heterozygosity of inter-pool populations at loci with dominance. Because the frequency of heterozygotes compared to homozygotes at segregating loci in autopolyploid populations is already relatively high compared to diploids, there is not only less value to be gained by increasing heterozygote frequency with Two-Pool GCA but also less value lost to the smaller increase in deleterious recessive homozygote frequency under selection on One-Pool Breeding Value (Supplemental Fig. 16, Supplemental Table 2). Though this study considered clonal species, these conclusions should be applicable to non-clonal autopolyploids.

Consistent with this hypothesis, the relative overperformance of one-pool strategies compared to Two-Pool GCA was greater in autohexaploids than autotetraploids: autohexaploids inherit more chromosome copies per gamete (3) than autotetraploids (2), leading to greater heterozygosity at segregating loci. We expect that the relative genetic gain per unit cost of Two-Pool GCA to One-Pool Breeding Value would be further reduced at higher autoploidies. Another line of support for this hypothesis was that the relative performance of Two-Pool GCA to other strategies increased with GS at high intensity. High-intensity GS likely increased inbreeding and genetic drift compared to low-intensity GS or high-intensity PS, so the ability of Two-Pool GCA to relieve homozygosity became more important. However, One-Pool Cross Performance was similarly capable of relieving inbreeding in this situation and is less logistically demanding. Finally, Two-Pool GCA built more panmictic heterosis than Two-Pool Breeding Value, but the difference was less in autopolyploids than diploids. This indicates breeding for heterosis with GCA was less effective in autopolyploids, since it more narrowly outperformed incurrence of heterosis due to drift.

It is possible that further increasing the inbreeding rate in autopolyploids (e.g. by reducing the number of parents or using truncation selection without inbreeding control) could increase the relative performance of Two-Pool GCA to other strategies, but this would not necessarily increase genetic gain. However, further investigation of strategy relative performance over additional inbreeding rates is warranted. Tangentially, the accuracy of autopolyploid genomic estimates tended to be similar to diploids at low H_0_, but increasingly lower than diploids at high H_0_, suggesting that allelic effects may be harder to predict in autopolyploids than diploids as dominance increases. This is sensible because more dominance effects are present in autopolyploids per phenotypic observation. However, it did not seem to be the main cause of the decreased advantage of Two-Pool GCA in the autopolyploids, which also appeared with use of true values. It may be worth noting that the lack of advantage to selection on Two-Pool GCA only applies to autopolyploids, not to allopolyploids for which chromosome copies are not independently assorted.

The lack of advantages of Two-Pool GCA in autopolyploids does not imply that autopolyploids cannot or do not exhibit heterosis. Selection on Two-Pool GCA or Two-Pool Breeding Value led to clear panmictic heterosis in the autopolyploids simulated in the study. Empirical evidence of panmictic heterosis in autohexaploid sweetpotato, for example, is readily available for fresh root yield (Diaz et al., 2021). The point is that even if autopolyploids exhibit heterosis or inbreeding depression, RRS did not provide increased gain per unit cost compared to RS on breeding value in a single, merged pool under the study assumptions. In the case of sweetpotato, two pools exhibiting panmictic heterosis emerged when a single breeding population was split into two locations (M. Andrade, pers. comm.). Over approximately twenty years, the pools were selected separately by truncation (W. Gruneberg, pers. comm.), and therefore allele frequencies likely came to diverge between pools due to selection and drift. Reunion of the pools then led to population-level panmictic heterosis in the F_1_ hybrids (Diaz et al., 2021). The existence of panmictic heterosis in autohexaploids does not imply that Two-Pool GCA or Two-Pool Breeding Value is the optimal breeding strategy for autohexaploids. The observed panmictic heterosis in sweetpotato could also be availed by intermating the two pools and conducting RS on breeding value in the single, merged pool. However, further comparisons of strategy efficiencies with pre-existing diverged pools would be informative in both diploids and autopolyploids.

The relatively decreased homozygosity of autopolyploids compared to diploids with selection on breeding value does not imply that autopolyploids suffer less inbreeding depression than diploids in the event that they do experience homozygosity of unfavorable alleles. This misconception may arise from failure to differentiate the inbreeding rate and inbreeding depression value. Autopolyploids in fact may experience more inbreeding depression in response to increased homozygosity than diploids, which can be observed in simulated autopolyploids produced by chromosome doubling with digenic dominance. Although few comparable estimates of inbreeding depression in real data are available, one such dataset is that of Yao et al. (2020), which compared genotypically matched diploid and autotetraploid maize. In a selfing series of each, Yao et al. observed similar inbreeding depression in the diploids and autotetraploids at the same selfing generation (2020). Since autotetraploids are less inbred than diploids at a given selfing generation, their similar inbreeding depression suggest that autotetraploid inbreeding depression was more severe per unit increase in homozygosity. Of course, it cannot be concluded that the maize autotetraploids used experienced only inbreeding depression due to digenic dominance, and the inbreeding depression observed could be due to loss of higher-order dominance interactions as well.

### Assumptions, limitations, and future research directions

The conclusions of this study depend on the assumptions made and parameters used. Further exploration of these factors is welcomed, and we encourage breeding programs to simulate and optimize their specific situation when information is readily available. Exploration of ranges of values is helpful to explore factors which affect the relative performance of breeding strategies, but once identified, the number of real-world constraints on breeding programs is much smaller than all possible constraints on breeding programs.

The breeding schemes used are not optimal but are rather a baseline for comparison of population improvement methods. For example, we did not optimize accuracy within the breeding strategies and estimation methods, which may require different designs for optimal accuracy. Particularly, testcrossing is necessary with phenotypic Two-Pool GCA but is suboptimal for genomic estimated Two-Pool GCA (Fristche-Neto et al., 2017; Seye et al., 2020). We did not optimize tester choice or number and simply used two random testers. With GS and Two-Pool Breeding Value, prediction of intra-pool genotypes from an inter-pool training set was suboptimal compared to use of intra-pool training genotypes, which has been demonstrated in prediction of purebred animals from crossbreds (Wei & Van der Werf, 1994; Moghaddar et al., 2014; Hidalgo et al., 2016). However, to address the lack of optimization of accuracy, we simulated all scenarios with true values to control accuracy across strategies and did not observe radically different trends of the breeding strategies with respect to population heterosis. The scenarios with true values have controlled accuracy but less genetic drift than GS scenarios, because true values are like using phenotypes with broad-sense heritabilities of one (Daetwyler et al., 2007; Sonesson et al., 2012).

We did not optimize each scenario to a given time horizon. The number of parents used were certainly not optimal for the time horizons explored, because unused genetic variance remained for all scenarios. It is possible that different strategies could produce different amounts of gain at optimal intensities for the times considered, and it may be that this also varies by genetic architecture. Somewhat arbitrarily, we also assumed a fixed number of parents per strategy rather than a fixed number of parents per pool.

We did not fully explore all possible genetic architectures, particularly those including epistasis or higher-order autopolyploid dominance. We note that positive directional dominance could arise from selection and was not necessarily present in the starting population for situations when Two-Pool GCA to presented advantages over one-pool strategies—e.g., with an initial mean dominance degree of zero and non-zero variance of dominance degrees (Falconer & Mackay, 1996; Varona et al., 2018). We did not consider environment or genotype x environment effects, which may affect the relative performance of GS and PS and depletion of genetic variance. We assumed a fixed marker density and genome size. We assumed biallelic loci. We do not expect that multiallelic loci in autopolyploids would likely lead to increased advantages of Two-Pool GCA, because with linkage disequilbrium haplotypes of biallelic loci effectively behave as a single multiallelic locus. We did not vary the probability of autopolyploid multivalents.

We assessed H_0_ as a predictor of various responses. H_0_ appeared to explain the variance of responses among strategies well, but it is possible that its components—mean dominance degree, the variance of the dominance degrees, and the square root of the number of QTL—could reveal different patterns of strategy performance if used as predictors rather than H_0_. We plotted genetic gain of the core strategies with use of true values after 50 years with use of each component as a predictor of responses with both other components held constant in all possible combinations (Supplemental Fig. 17—25). In general, we observed similar patterns as with use of H_0_ for mean dominance degree and the square root of the number of QTL, with the relative performance of Two-Pool GCA increasing as each of these increased. The relative performance of Two-Pool GCA increased as mean dominance degree increased regardless of whether incomplete dominance, complete dominance, or overdominance was simulated; notably, overdominance did not decrease the relative advantage of Two-Pool GCA (Rembe et al., 2019). However, for the variance of dominance degrees, if the mean dominance degree was low then advantage of Two-Pool GCA increased as the variance of dominance degrees increased, even though the variance of dominance degrees has an inverse relationship with H_0_. This seemed to be because selection on GCA led to directional dominance in the breeding population when loci with positive dominance degrees were present. This trend reversed to expectation as mean dominance degree and the number of QTL increased.

With use of maximum avoidance at high vs. low intensity, there were necessarily more full siblings per family at high vs. low intensity. Availability of additional full siblings at high intensity may have increased the accuracy of prediction of dominance values (Misztal et al., 1998), which could affect the relative performance of Two-Pool GCA. However, the difference in relative performance between Two-Pool GCA and other strategies at high vs. low intensity was also apparent with use of true values at perfect accuracy, indicating the influence of the inbreeding rate.

Although we completely disregarded product development strategies or prediction of inter-pool crosses in additional to GCA for RRGS, we presume that population improvement strategies which produce populations with higher means and similar distributions will lead to extraction of higher-value products with all else, such as product evaluation strategy, equal. Allocation of resources among stages was not explored.

The study considered plausible values for the cost of phenotyping, genotyping, and phenotyping to genotyping among strategies, but these may differ among applied programs. Particularly, the cost of two-pool vs. one-pool breeding depends strongly on crop biology. We assumed that the cost of controlled inter-pool crossing was negligible, which may not be the case in some crops.

Multiple frameworks to model dominance in polyploids are available; here, only digenic dominance is considered, while other frameworks allow for additional intra-locus interactions (Gallais, 2003). It does not seem likely that other valuations of various possible heterozygotes or inclusion of additional intra-locus interactions would change the relative performances of the strategies presented here, because the superfluity of Two-Pool GCA seems to arise from the increased frequency of heterozygotes in autopolyploids regardless of their valuation. However, further study may reveal unexpected results.

We note that heterosis in autopolyploids is not maximized with single crosses among two diverged pools, i.e. heterosis is progressive (Groose et al., 1989; Washburn & Birchler, 2014; Washburn et al., 2019; Labroo et al., 2021). Autopolyploid heterosis due to dominance is progressive because autopolyploids have fewer parents than inherited gametes. If allele frequencies diverge randomly across the genome among parents, additional heterosis occurs by making multi-parental crosses because additional heterozygosity can be stacked into the progeny genome. We do not expect that utilization of progressive heterosis in autopolyploids would change the relative performance of the strategies because the additional heterosis is likely relatively small compared to the potential additional time needed to make additional crosses as well as the resources needed to maintain additional pools. However, testing this hypothesis is warranted. We note that progressive heterosis due to digenic dominance can be observed by the simulation methods of the study (https://github.com/gaynorr/AlphaSimR_Examples/blob/master/misc/ProgressiveHeterosis.R).

As mentioned repeatedly, comparisons of gain across ploidies from simulation should not be made because they are not guaranteed to reflect biological reality. Real data, which are likely population-specific, would be needed. For example, we assume that the minimum homozygote and maximum heterozygote value are the same in diploids and polyploids, but there is evidence that this is unrealistic in some populations because polyploid populations produced by colchicine doubling sometimes have higher mean values than their diploid progenitors (Sattler et al., 2016). For example, in the case of potato, our findings strongly suggest that Two-Pool GCA is not likely to be the optimal breeding strategy for autotetraploid potato, whereas Two-Pool GCA is likely to be the optimal breeding strategy for diploid potato if GS is used or H_0_ is adequate. However, we cannot determine from simulation alone whether overall genetic gain is likely to be higher in autotetraploid or diploid potato.

## Statements and Declarations

### Author Contribution Statement

The authors confirm contribution to the paper as follows: study conception: GCP, JBE, RCG; development of theory and algorithms: RCG, JBE; study design: all authors; coding: RCG, MRL, JBE; data collection: MRL, DCG, GCP; analysis and interpretation of results: all authors; figure design: all authors; manuscript editing: all authors. All authors reviewed the results and approved the final version of the manuscript.

*Acknowledgments*

We thank the CGIAR and the Roots, Tubers, and Bananas community for helpful discussion and motivating questions regarding hybrid breeding, particularly Asrat Amele, Elizabeth Parkes, Godwill Makunde, Ismail Kayondo, Ismail Rabbi, Jean-Luc Jannink, Maria Andrade, Marnin Wolfe, Paterne Agre, Randall Holley, Reuben Ssali, Wolfgang Grüneberg, and Xiaofei Zhang.

We thank Jaime Campos Serna and Rachel Lombardi for maintaining computing resources which enabled the study. This research was performed using the compute resources and assistance of the CIMMYT HPCC and the UW-Madison Center For High Throughput Computing (CHTC) in the Department of Computer Sciences. The CHTC is supported by UW-Madison, the Advanced Computing Initiative, the Wisconsin Alumni Research Foundation, the Wisconsin Institutes for Discovery, and the National Science Foundation, and is an active member of the OSG Consortium, which is supported by the National Science Foundation and the U.S. Department of Energy’s Office of Science.

### Data availability

All code and results generated for the current study are available in the Supplementary Information. Raw data are available at https://doi.org/10.7910/DVN/7RVFL8. The initial simulated populations used are available upon request.

