## Supplemental Figures for "Clonal breeding strategies to harness heterosis: insights from stochastic simulation"

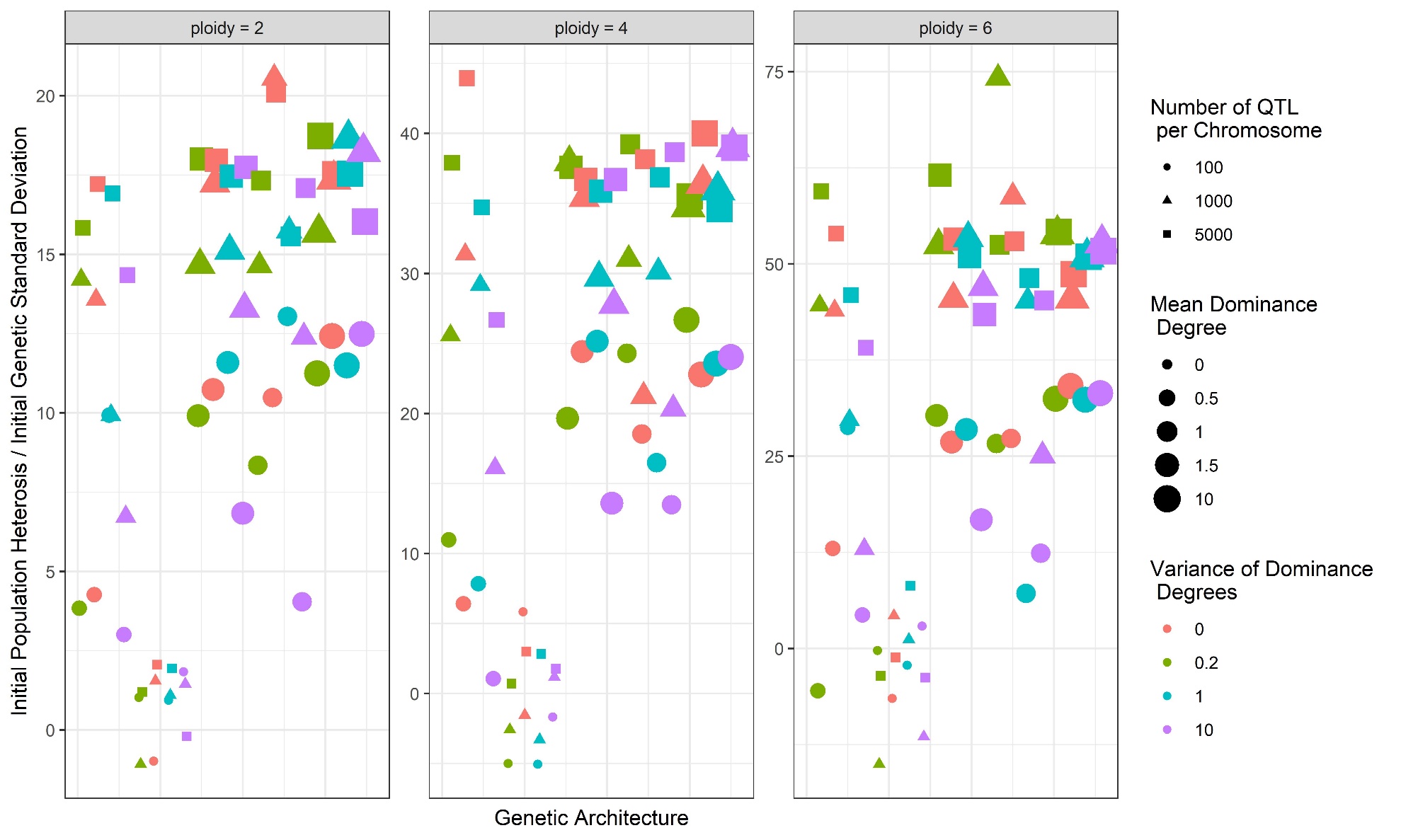

Supplemental Figure 1. Scaled initial population heterosis (H_0_) resulting from each combination of ploidy, number of QTL per chromosome (point shape), mean dominance degree (point size), and variance of the dominance degrees (point color). Ten chromosomes were assumed. A larger range of H_0_ values were simulated as ploidy increased. In general, H_0_ increases with increased mean dominance degree or number of QTL per chromosome, but slightly decreases with increased variance of dominance degrees. Linkage disequilibrium also affects H_0_. Multiple genetic architectures can lead to the same heterosis value.

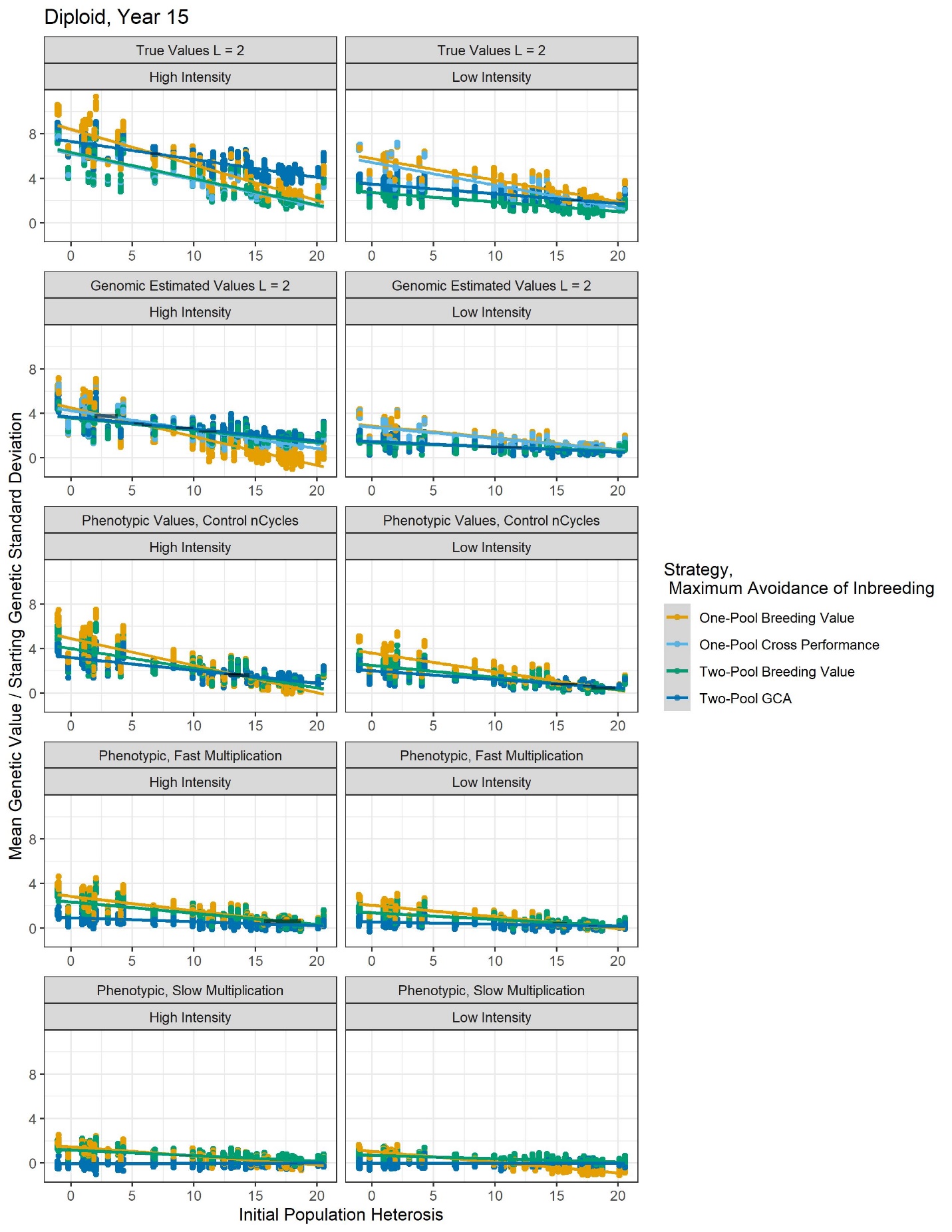

Supplemental Figure 2. Diploid genetic gain in the product pool at year 15. Symbols are as described in Fig. 2. With PS, Two-Pool GCA is never the best strategy regardless of H_0_.

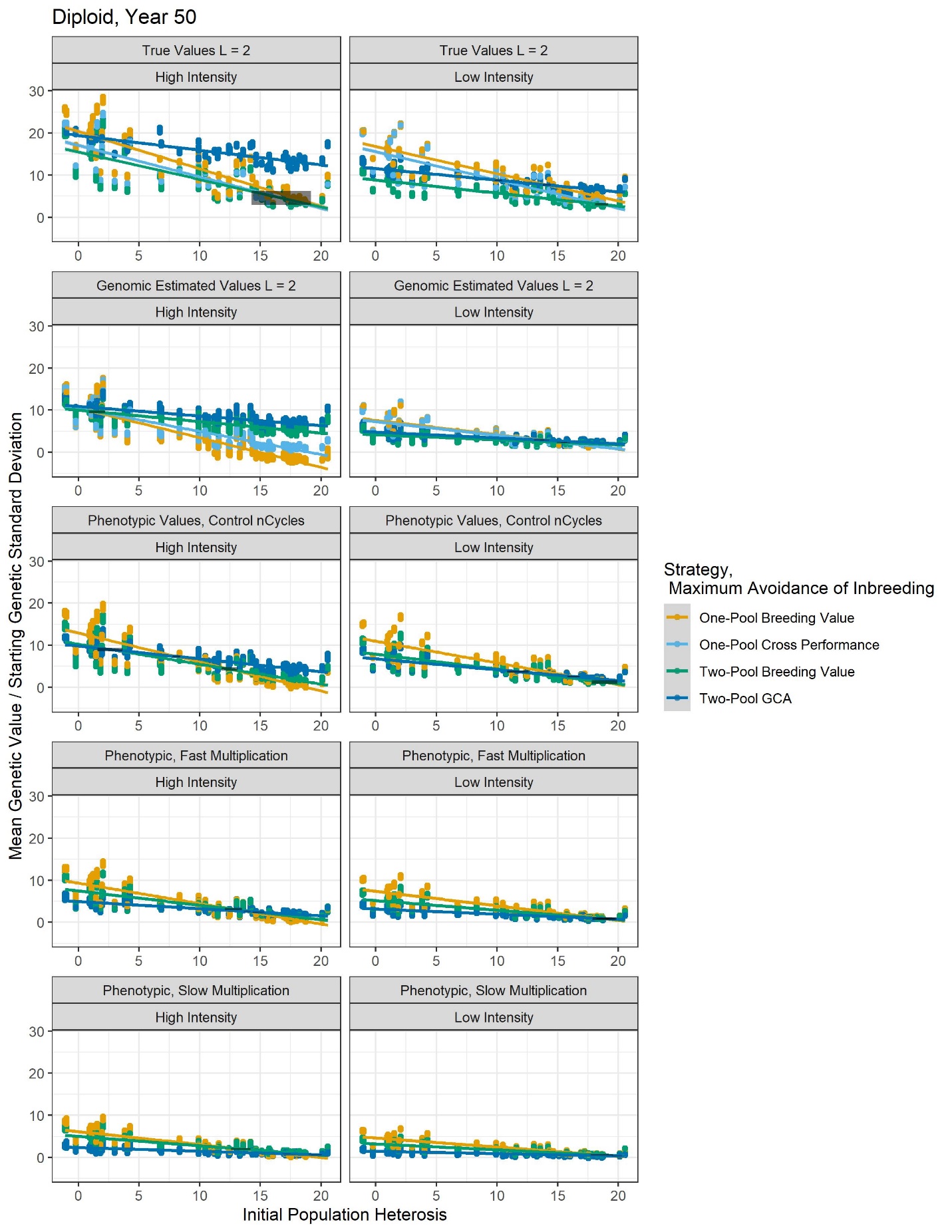

Supplemental Figure 3. Diploid genetic gain at year 50. Symbols are as described in Fig. 2. With PS, Two-Pool GCA provides marginal gain at high intensity only if H_0_ is adequate with fast multiplication.

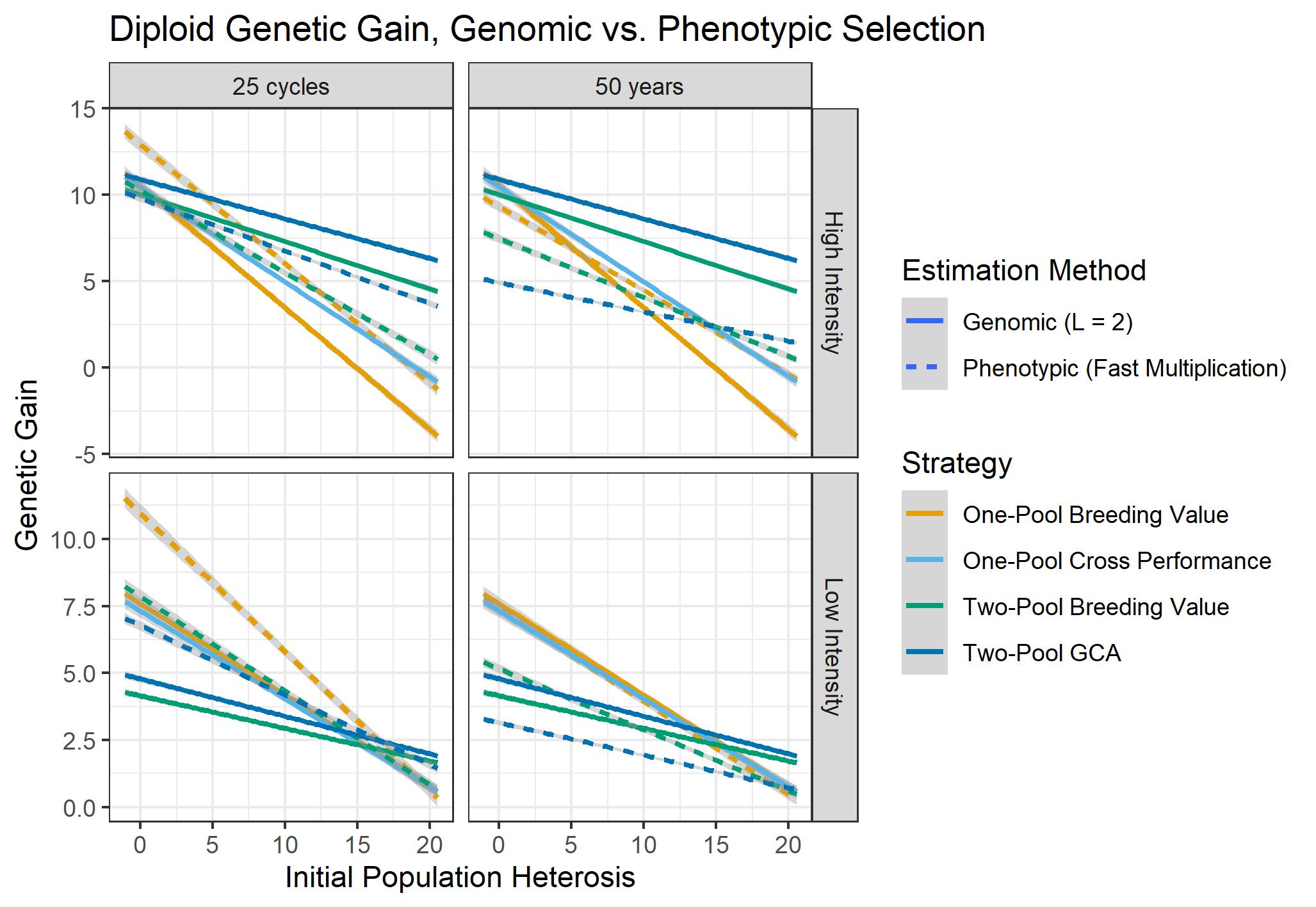

Supplemental Figure 4. Diploid genetic gain with genomic vs. phenotypic selection after a fixed number of cycles (25) and, realistically, after a fixed number of years (50). After 50 years, the best genomic selection strategy outperforms the best phenotypic strategy at high intensity and matches or exceeds the best phenotypic strategy at low intensity. However, if compared after 25 cycles, there is crossover in whether phenotypic or genomic selection is higher-performing. This highlights the utility of genomic selection to reduce cycle length, because comparing at a fixed number of cycles removes the effect of differing cycle lengths between strategies. Failure to use genomic selection to reduce cycle length can erase its benefits in some situations.

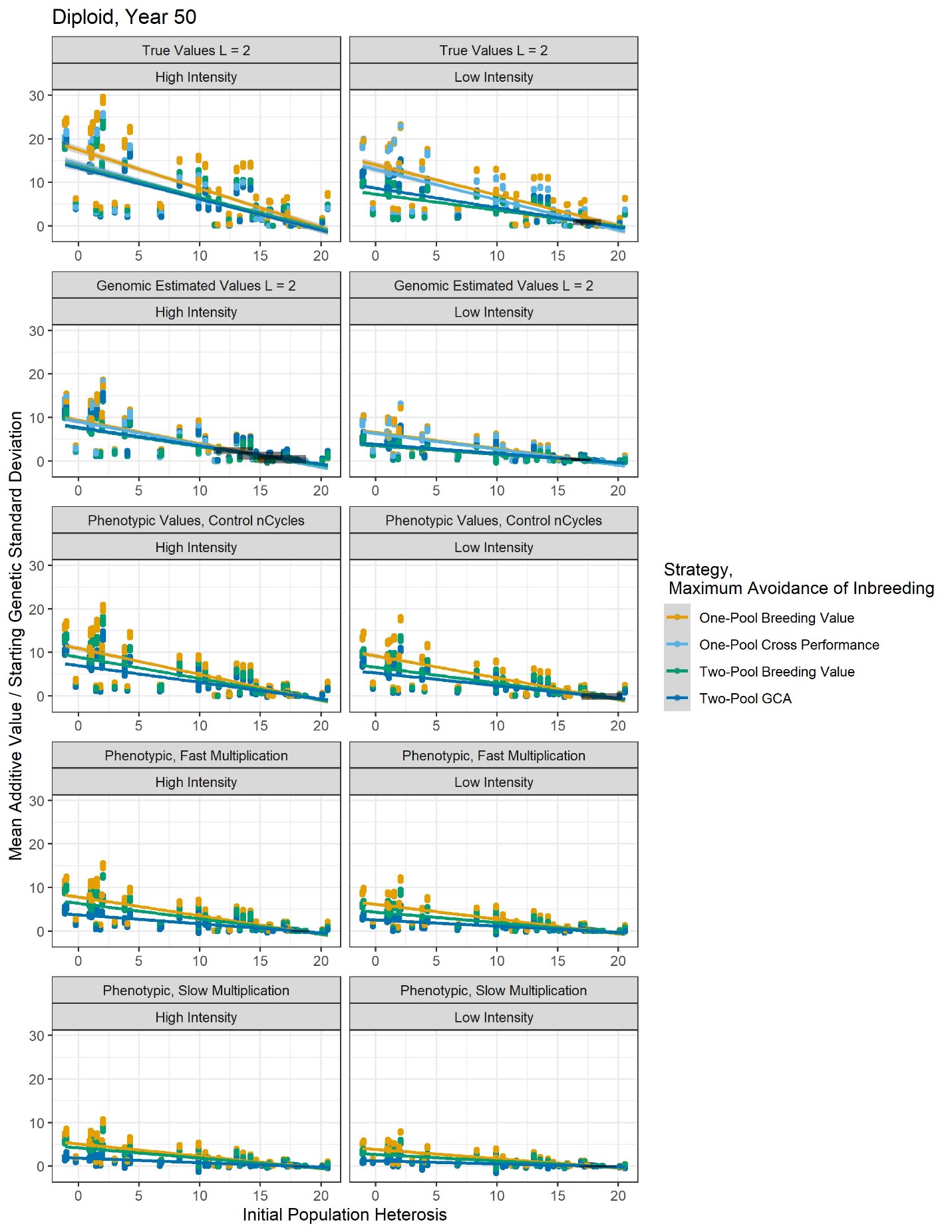

Supplemental Figure 5. Diploid additive value by strategy across H_0_ after 50 years of breeding. One-pool strategies typically have higher additive values than two-pool strategies.

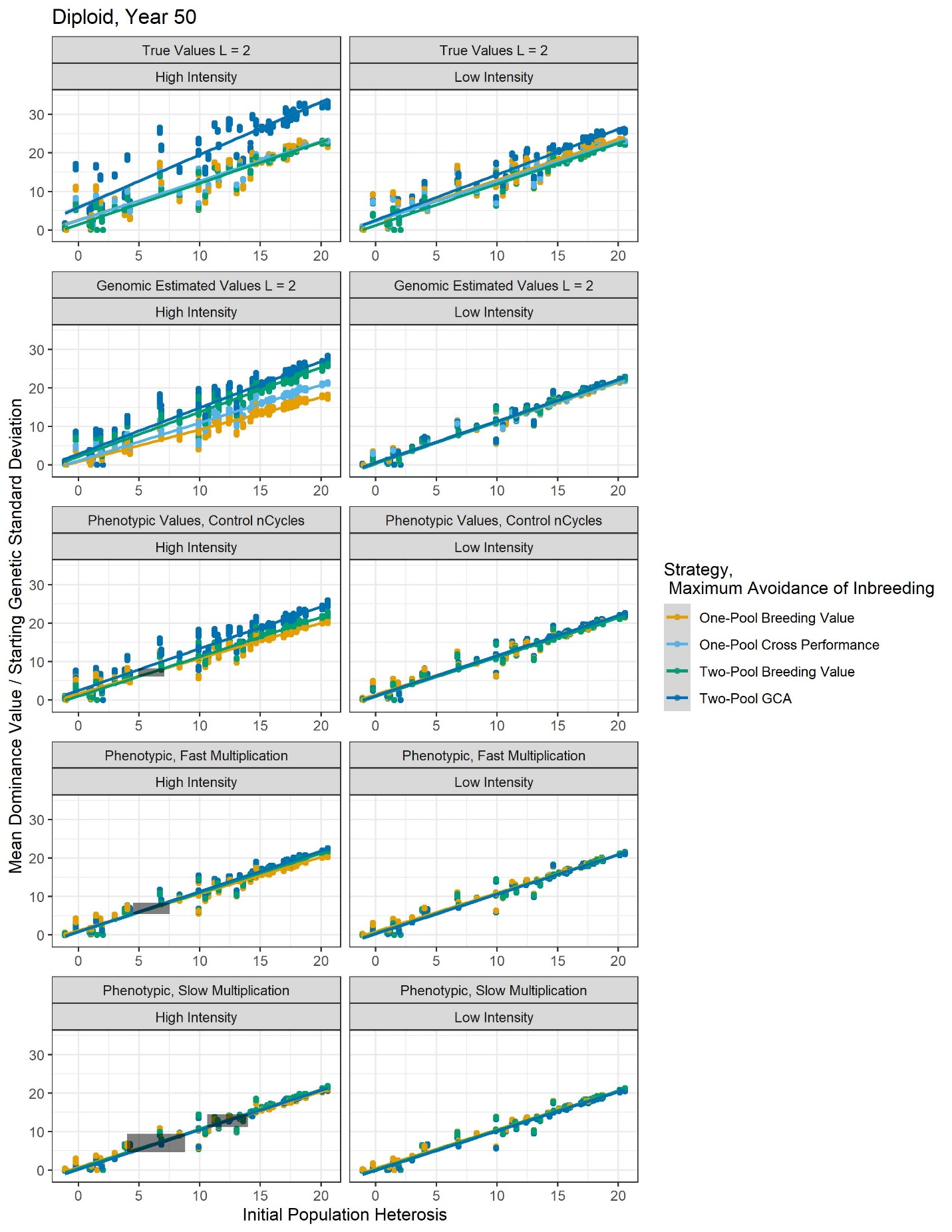

Supplemental Figure 6. Diploid dominance value by strategy across H_0_ after 50 years of breeding. Two-pool strategies typically have higher dominance values than one-pool strategies at high intensity.

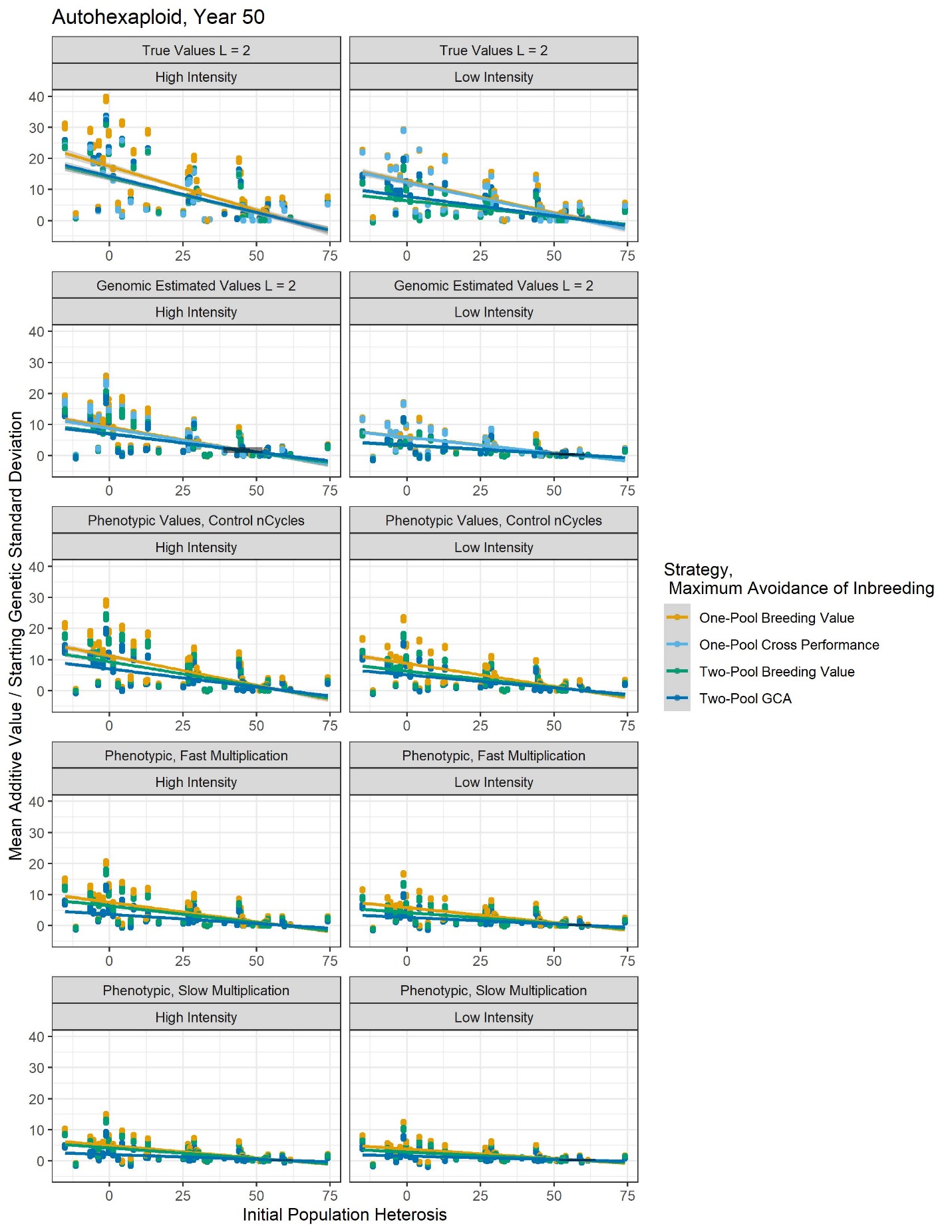

Supplemental Figure 7. Autohexaploid additive value by strategy across H_0_ after 50 years of breeding. One-pool strategies typically have higher additive values than two-pool strategies.

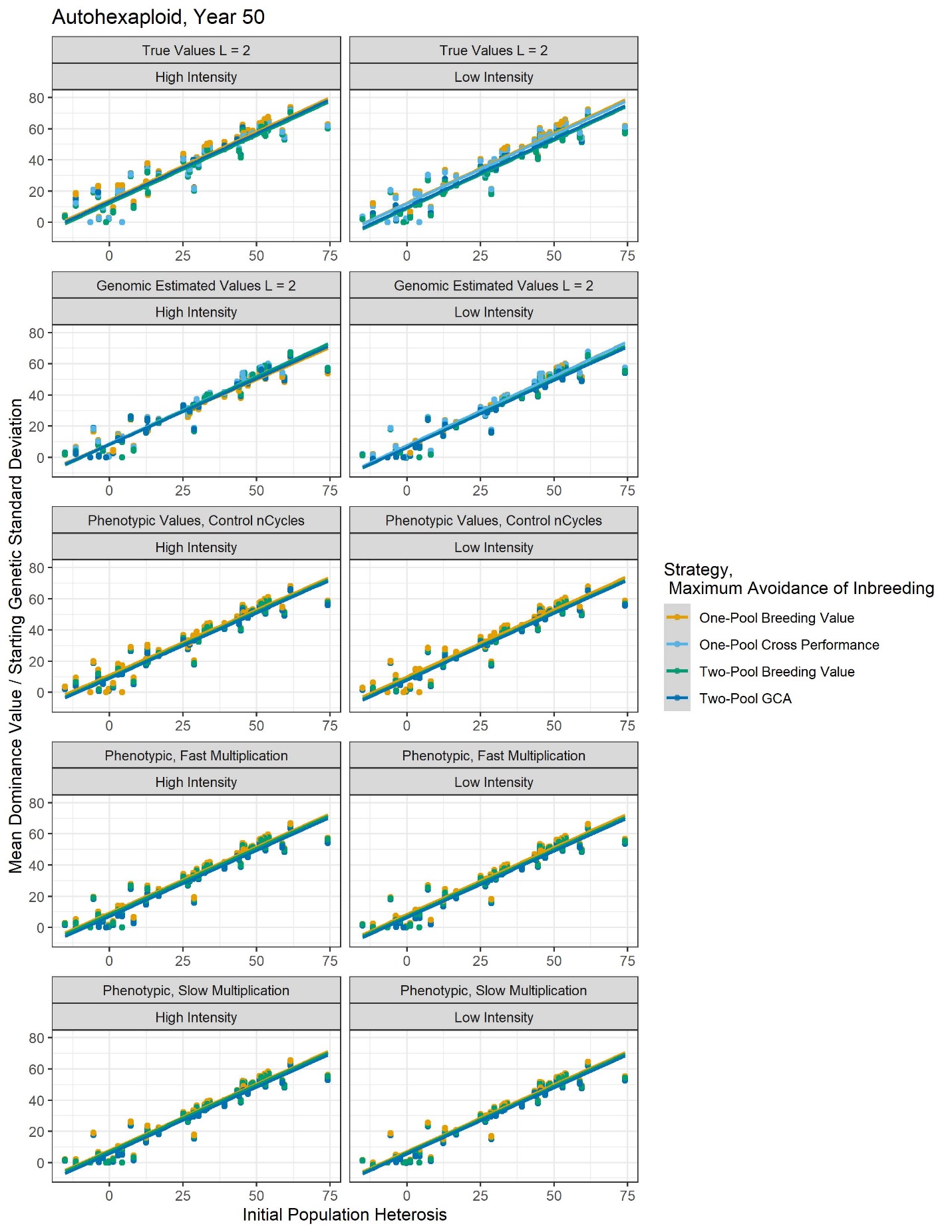

Supplemental Figure 8. Autohexaploid dominance value by strategy across H_0_ after 50 years of breeding. There is no difference in dominance value by strategy.

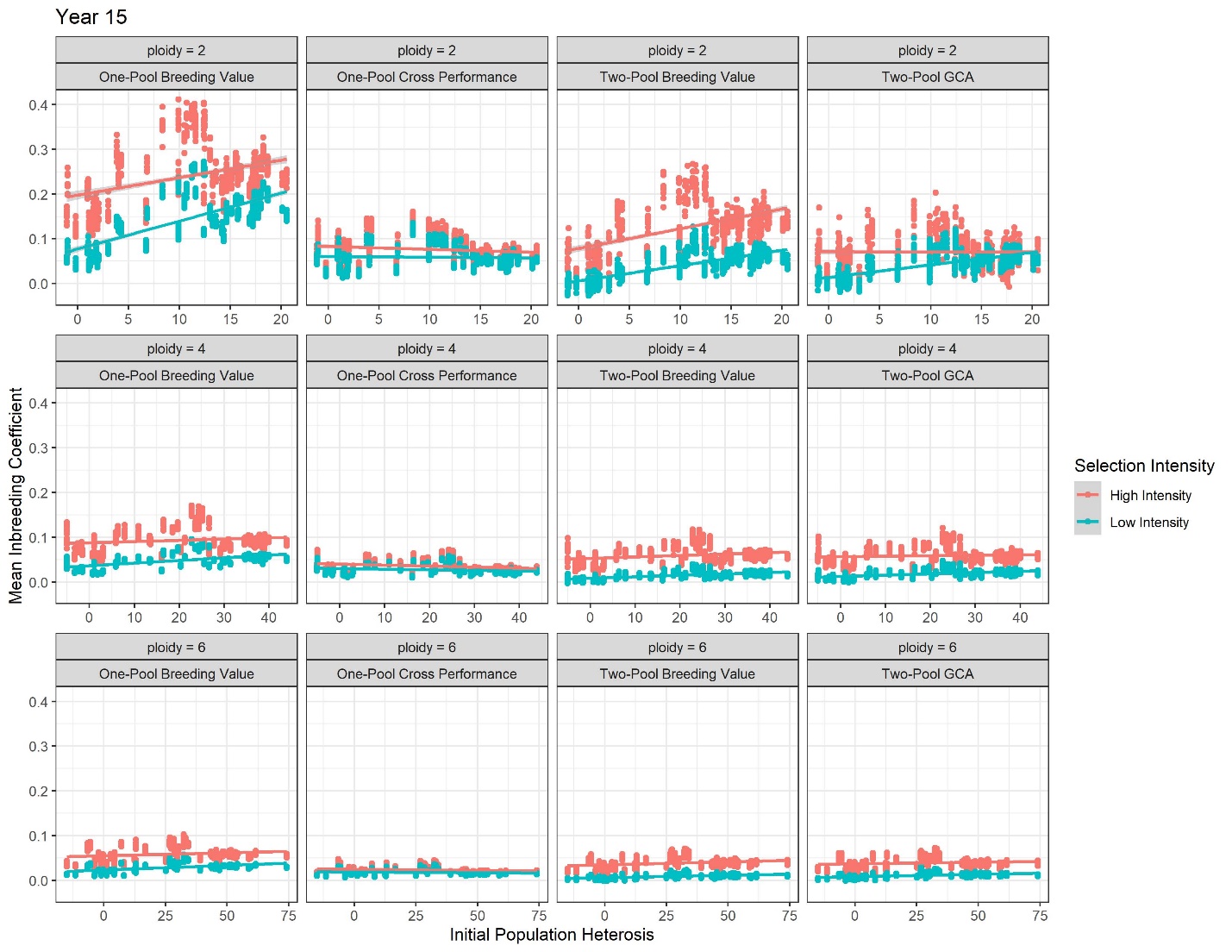

Supplemental Figure 9. Mean inbreeding coefficient by ploidy, strategy, and intensity as a function of H_0_ with use of true values after 15 years. High intensity programs typically have higher inbreeding coefficients than low intensity programs, but the difference is less as ploidy increases.

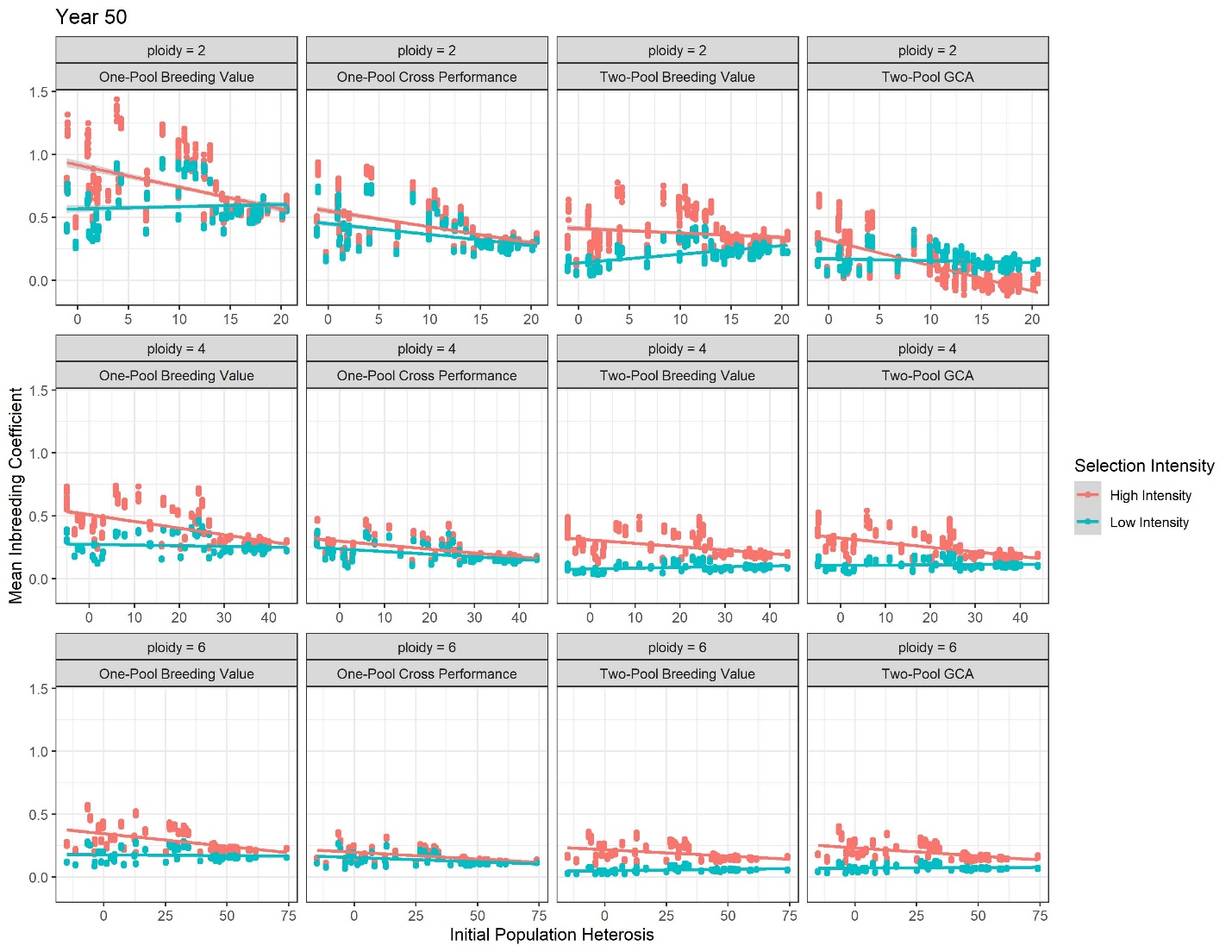

Supplemental Figure 10. Mean inbreeding coefficient by ploidy, strategy, and intensity as a function of H_0_ with use of true values after 50 years. High intensity programs typically have higher inbreeding coefficients than low intensity programs, but crossover occurs for some strategies. For Two-Pool GCA, at high intensity the product pool is likely driven to increased heterozygosity at high H_0_ compared to low intensity, so the inbreeding coefficient decreases. The difference in inbreeding depression by intensity is less as ploidy increases.

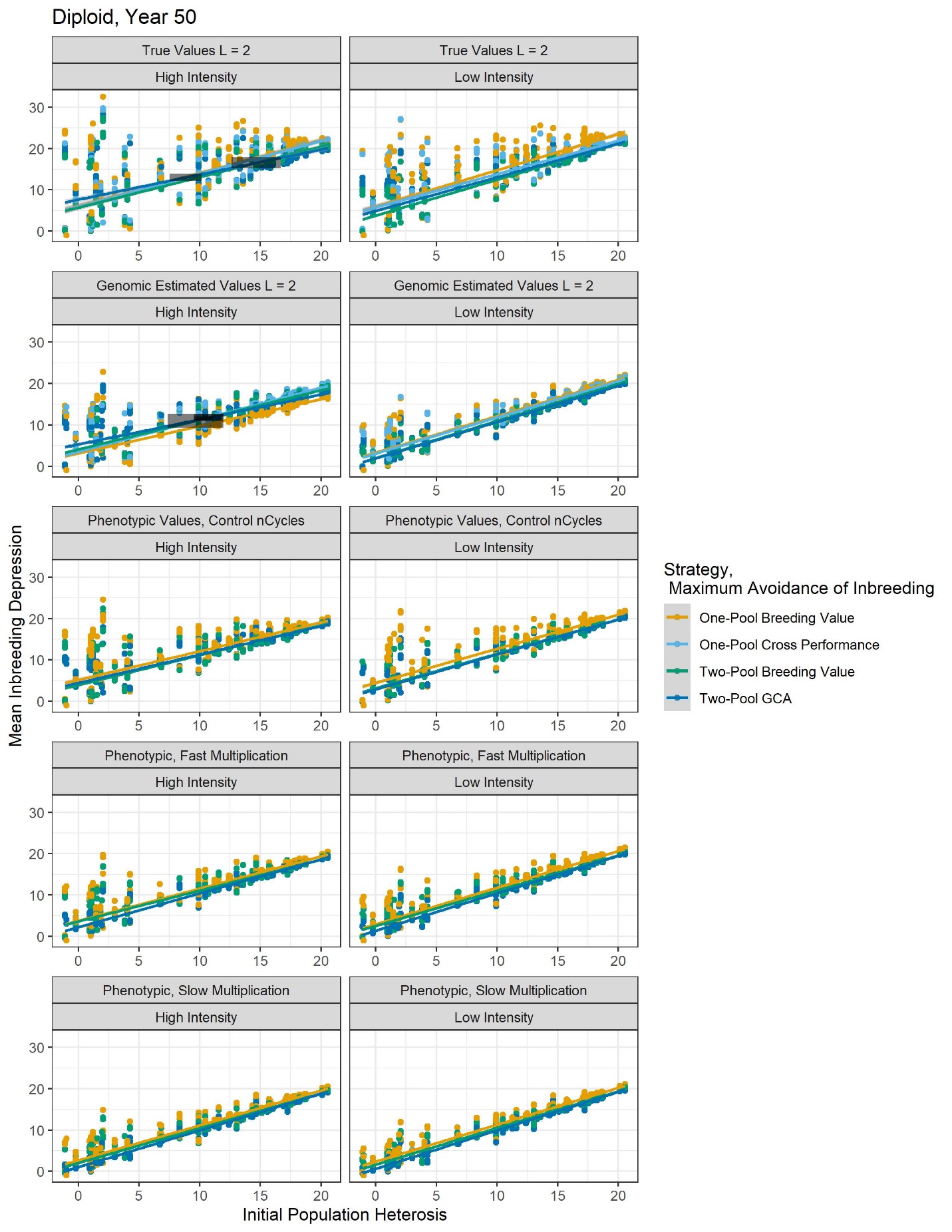

Supplemental Figure 11. Mean diploid population inbreeding depression at year 50 as a function of H_0_ by strategy. In general, strategies do not differ much in their reduction of population inbreeding depression.

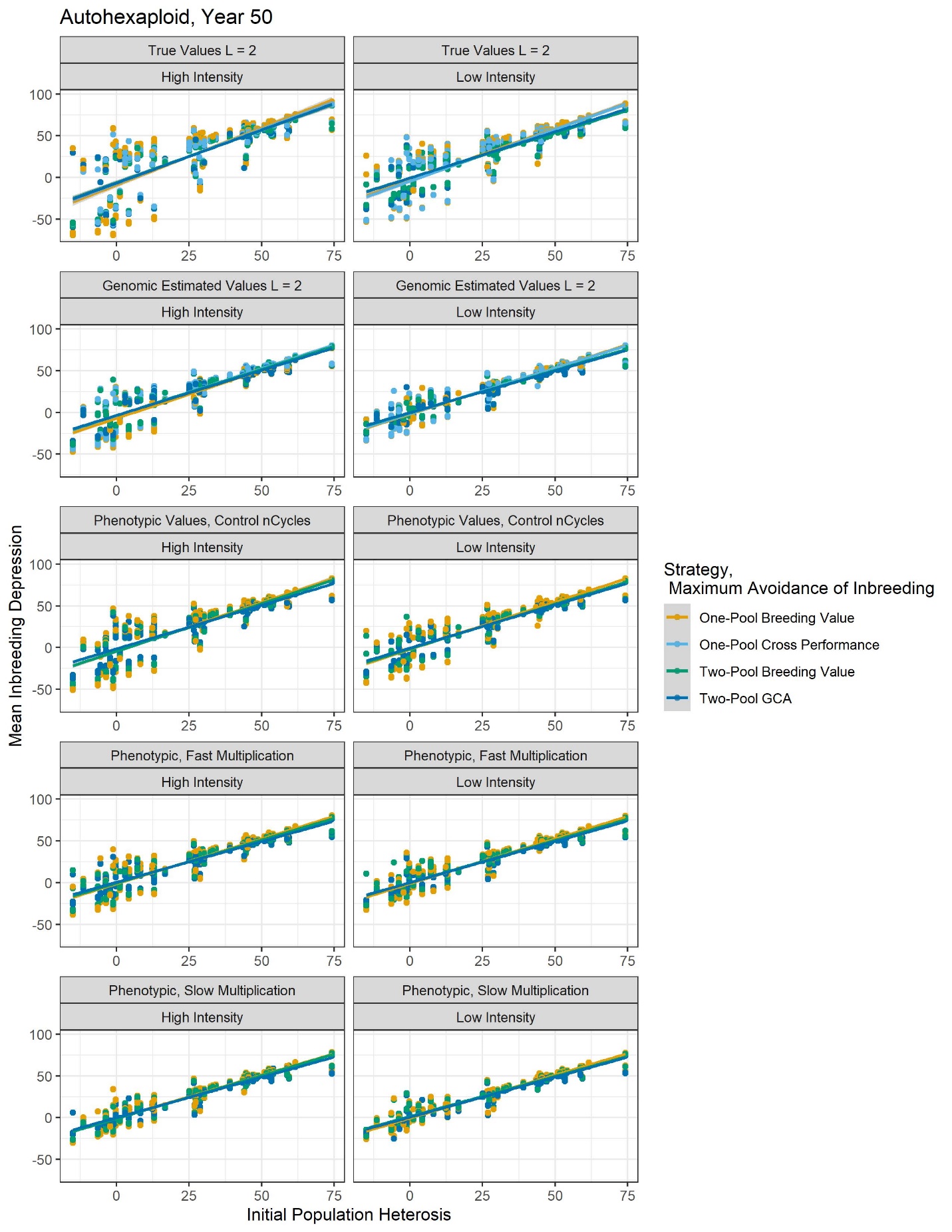
Supplemental Figure 12. Mean autohexaploid population inbreeding depression at year 50 as a function of H_0_ by strategy. Strategies do not differ in their capacity to reduce population inbreeding depression.

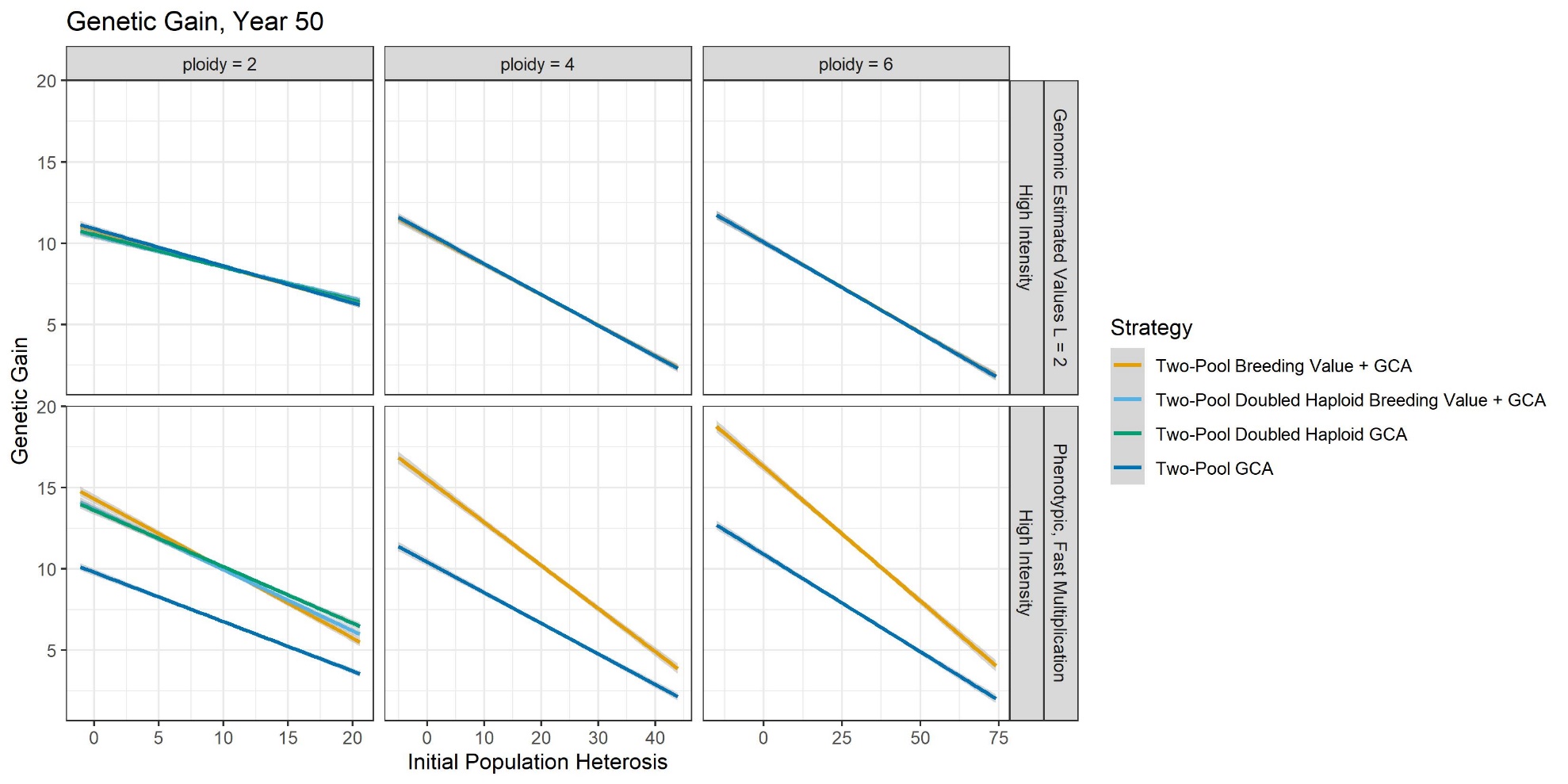

Supplemental Figure 13. Genetic gain in the non-core Breeding Value + GCA strategies as a function of H_0_ after 50 years at high intensity with GS and PS. With use of GS, intra-pool evaluation did not affect genetic gain. This was likely due to use of inter-pool genotypes to predict the intra-pool breeding values. With PS and fast multiplication, intra-pool evaluation increased genetic gain if doubled haploids were not used, possibly due to the use of outbred testers. Strategies which used doubled haploids were simulated for diploids only.

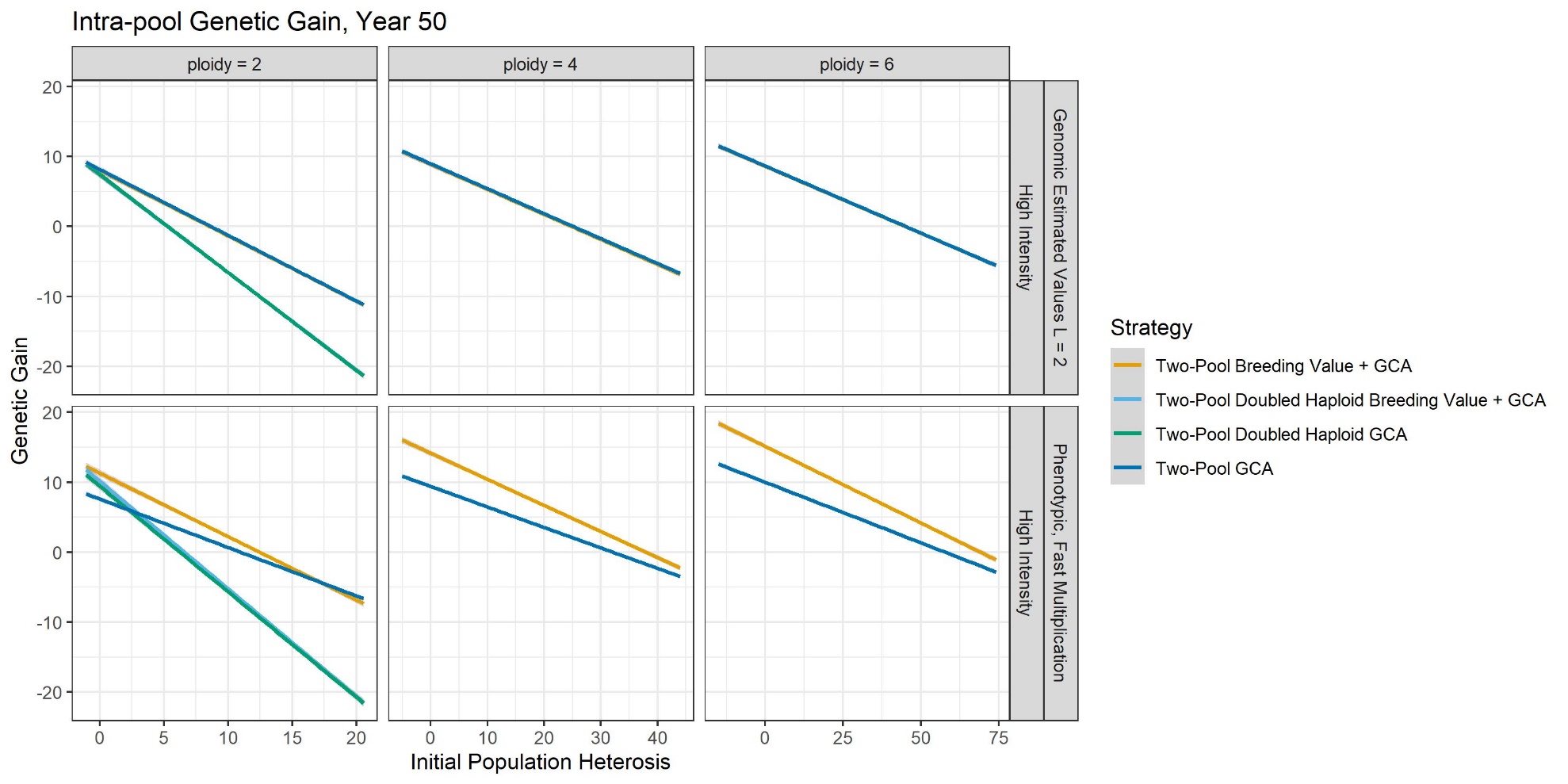

Supplemental Figure 14. Genetic gain in the intra-pool genotypes of Breeding Value + GCA strategies as a function of H_0_ after 50 years at high intensity with GS and PS. Notably, as H_0_ increased, doubled haploid intra-pool genotypes suffered increased inbreeding depression. Whether selection on GCA created positive, negative, or zero genetic gain over cycles depended on H_0_, and as H_0_ increased intra-pool genotypes had negative genetic gain. Use of intra-pool evaluation increased intra-pool genetic gain with PS if doubled haploids were not used, and intra-pool evaluation had no effect if doubled haploids were used. In the top three panels, the dark blue line covers the orange line. In the top left panel, the teal line covers the light blue line. Strategies which used doubled haploids were simulated for diploids only.

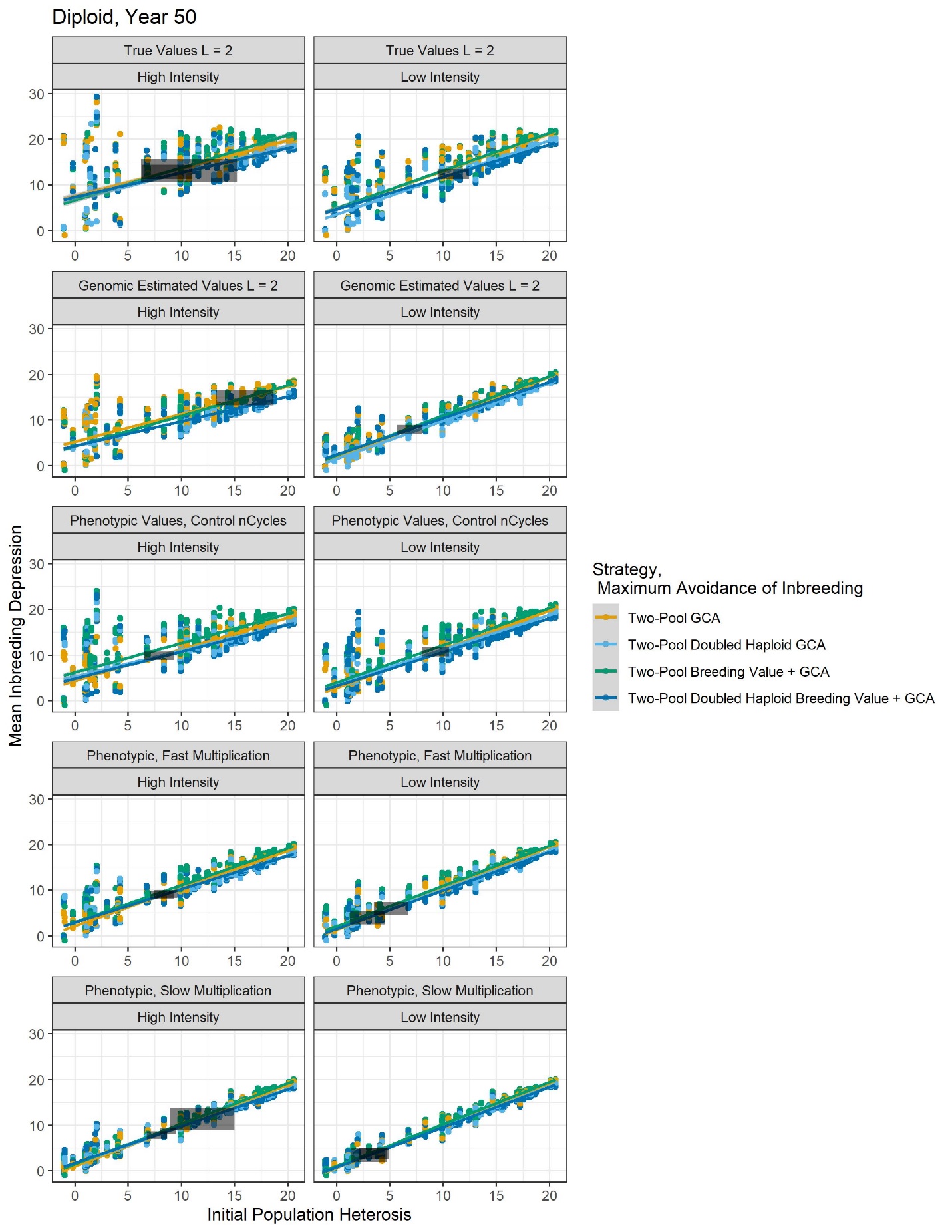

Supplemental Figure 15. Population inbreeding depression after 50 years with the non-core breeding strategies in diploids. Strategies tended to maintain similar amounts of inbreeding depression.

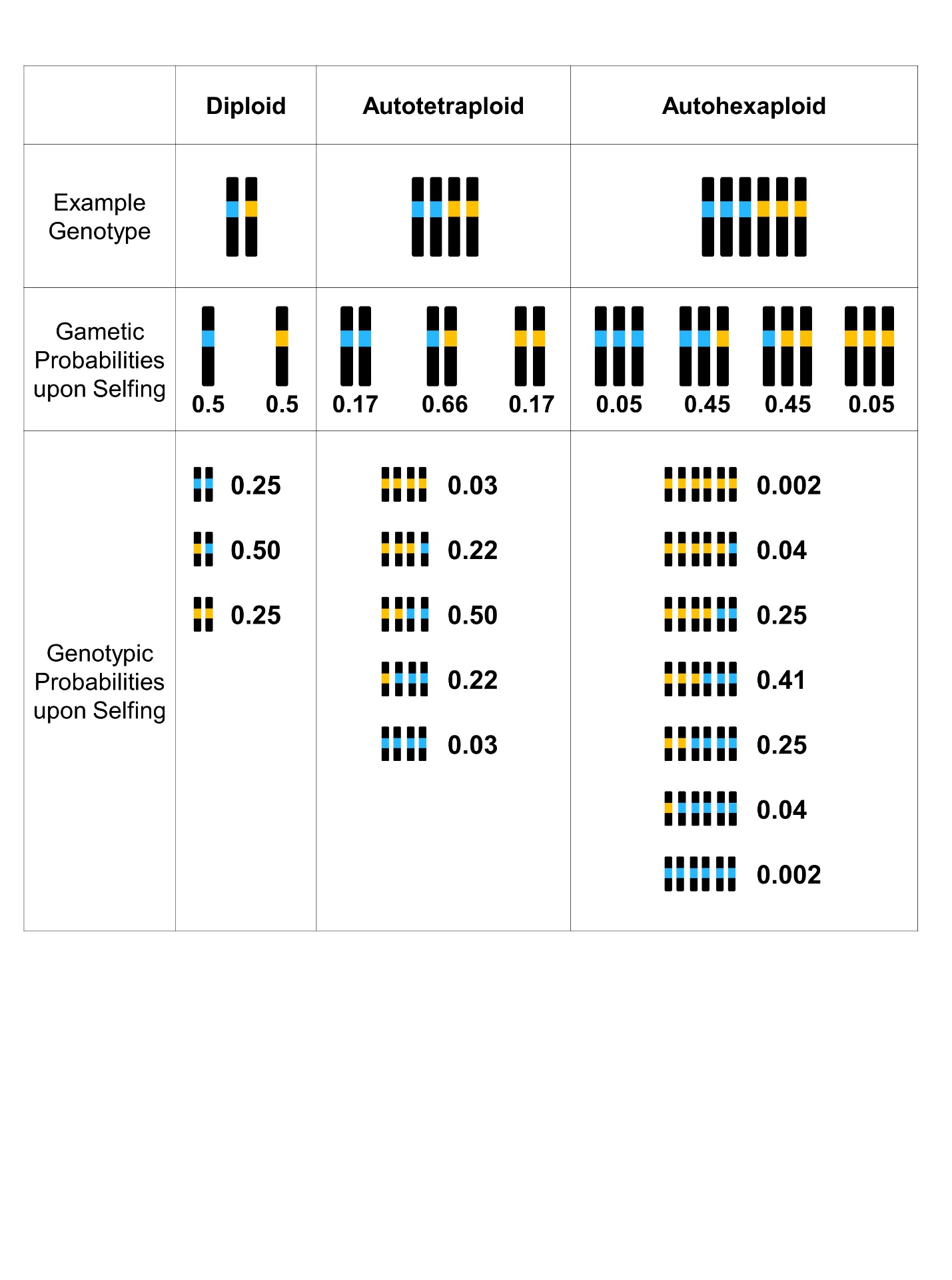

Supplemental Figure 16. Illustration of how autopolyploid meiosis leads to decreased probability of fully homozygous genotypes compared to diploids. Example genotypes are indicated by colored chromosome sets, with blue and orange rectangles indicating reference and alternate alleles. As an example, gametic and genotypic probabilities upon selfing are given in chromosome subtext. This decreased homozygosity at the gametic level leads to decreased homozygosity at the population level, as seen in Table B. Even though this decreased homozygosity implies that autopolyploids have a slower inbreeding rate than diploids, it does not imply that autopolyploids suffer less inbreeding depression than diploids.

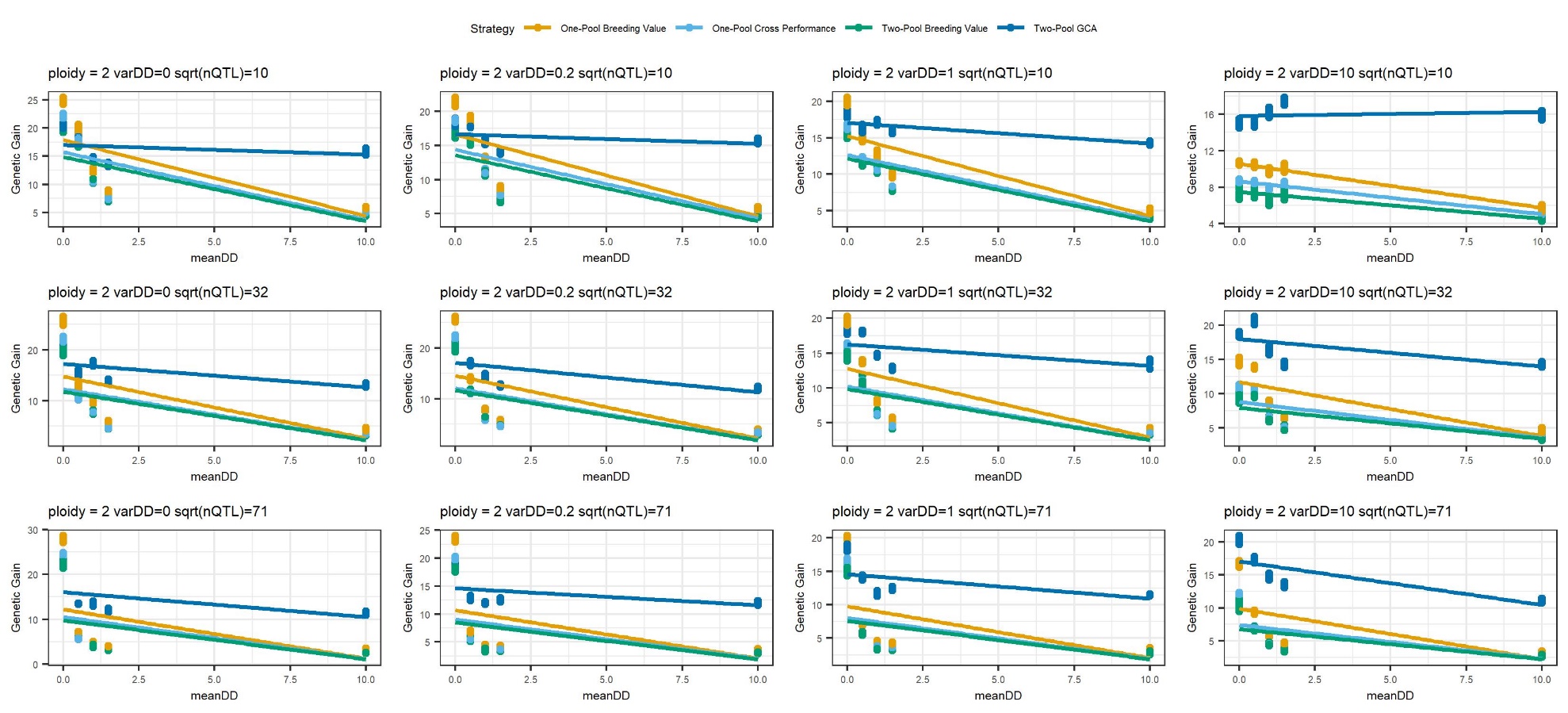

Supplemental Figure 17. For use of true values at high intensity after 50 years in diploids, the relative performance of the core strategies as a function of mean dominance degree (meanDD) instead of H_0_, at each level of the variance of dominance degrees (varDD) and the square root of the number of QTL per chromosome (sqrt(nQTL)).

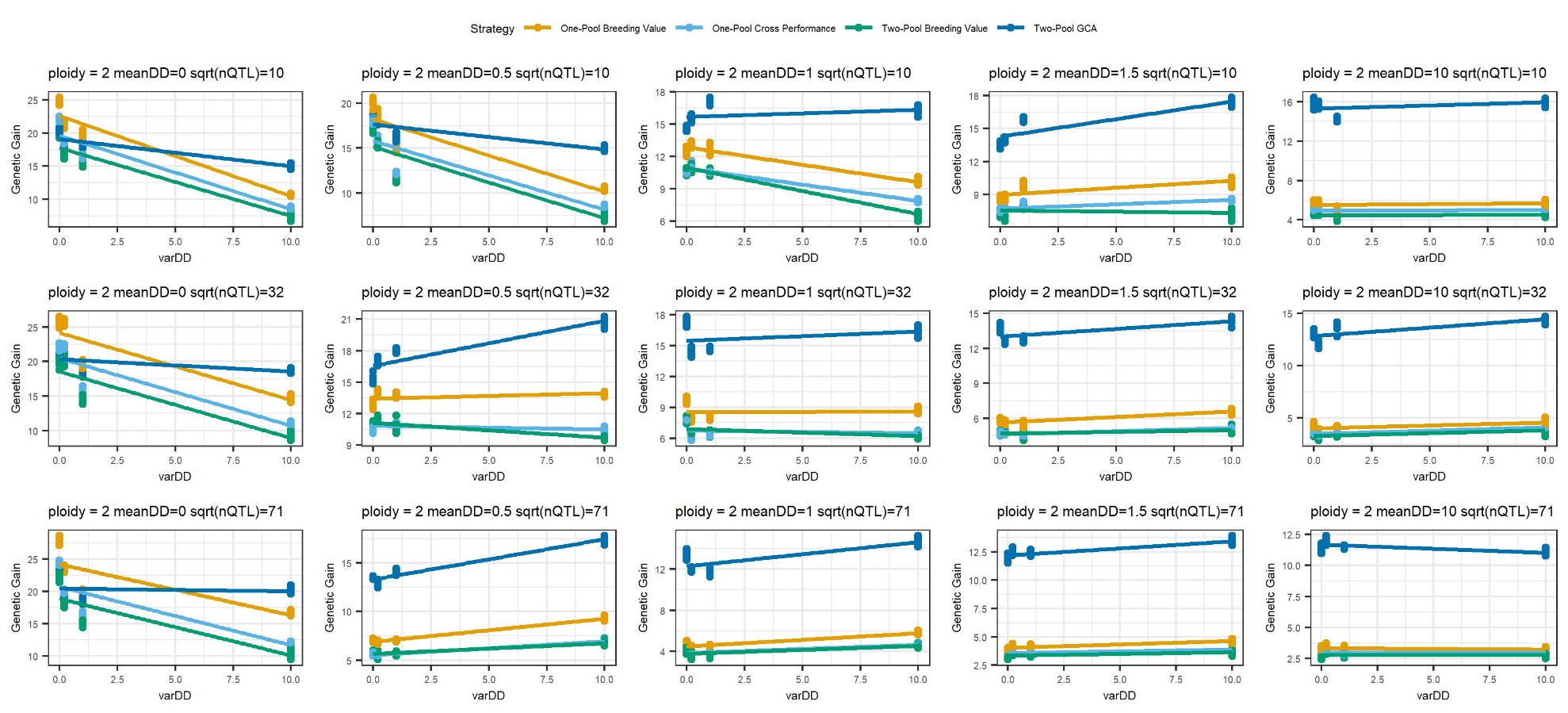

Supplemental Figure 18. For use of true values at high intensity after 50 years in diploids, the relative performance of the core strategies as a function of the variance of dominance degrees (varDD) instead of H_0_, at each level of the mean dominance degree (meanDD) and the square root of the number of QTL per chromosome (sqrt(nQTL)).

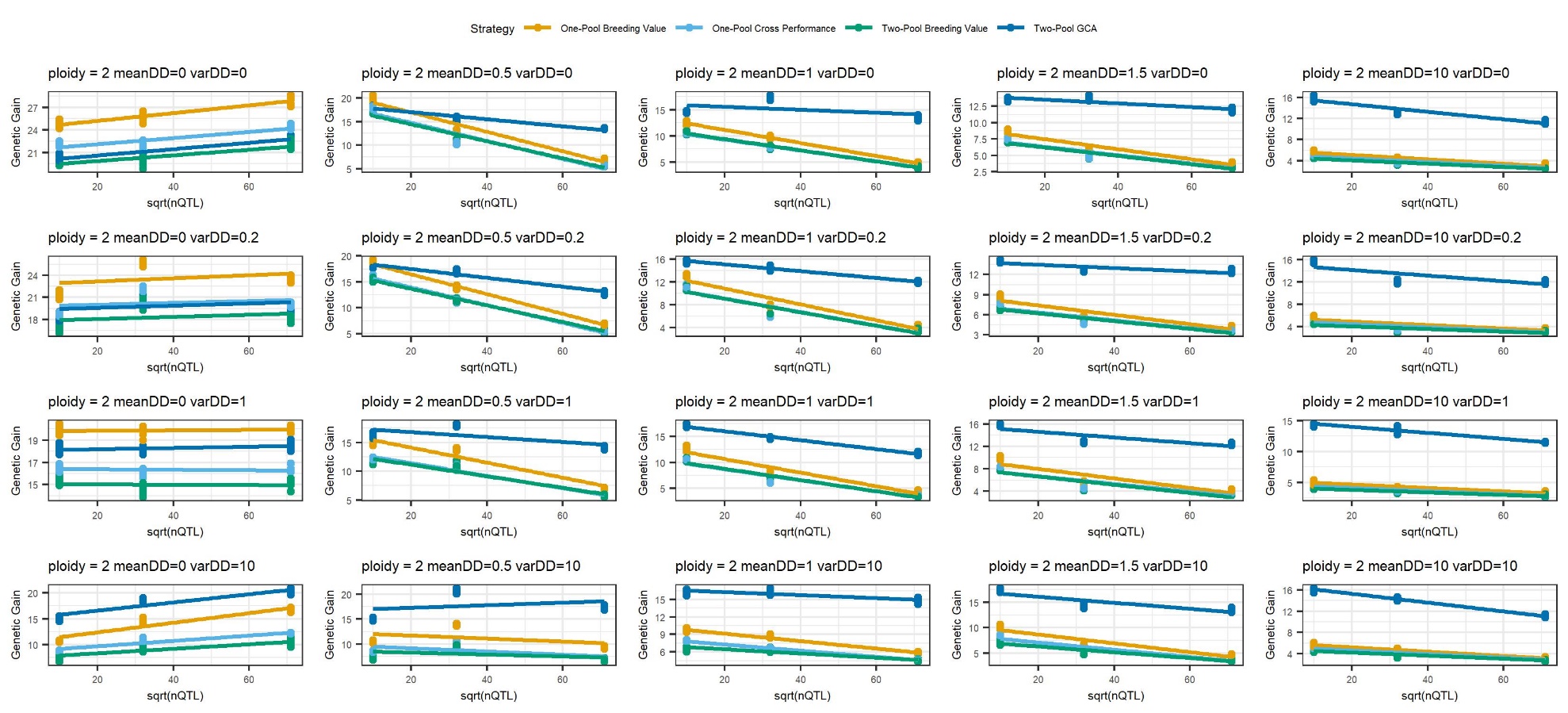

Supplemental Figure 19. For use of true values at high intensity after 50 years in diploids, the relative performance of the core strategies as a function of the square root of the number of QTL per chromosome (sqrt(nQTL) instead of H_0_, at each level of the variance of dominance degrees (varDD) and the mean dominance degree (meanDD).

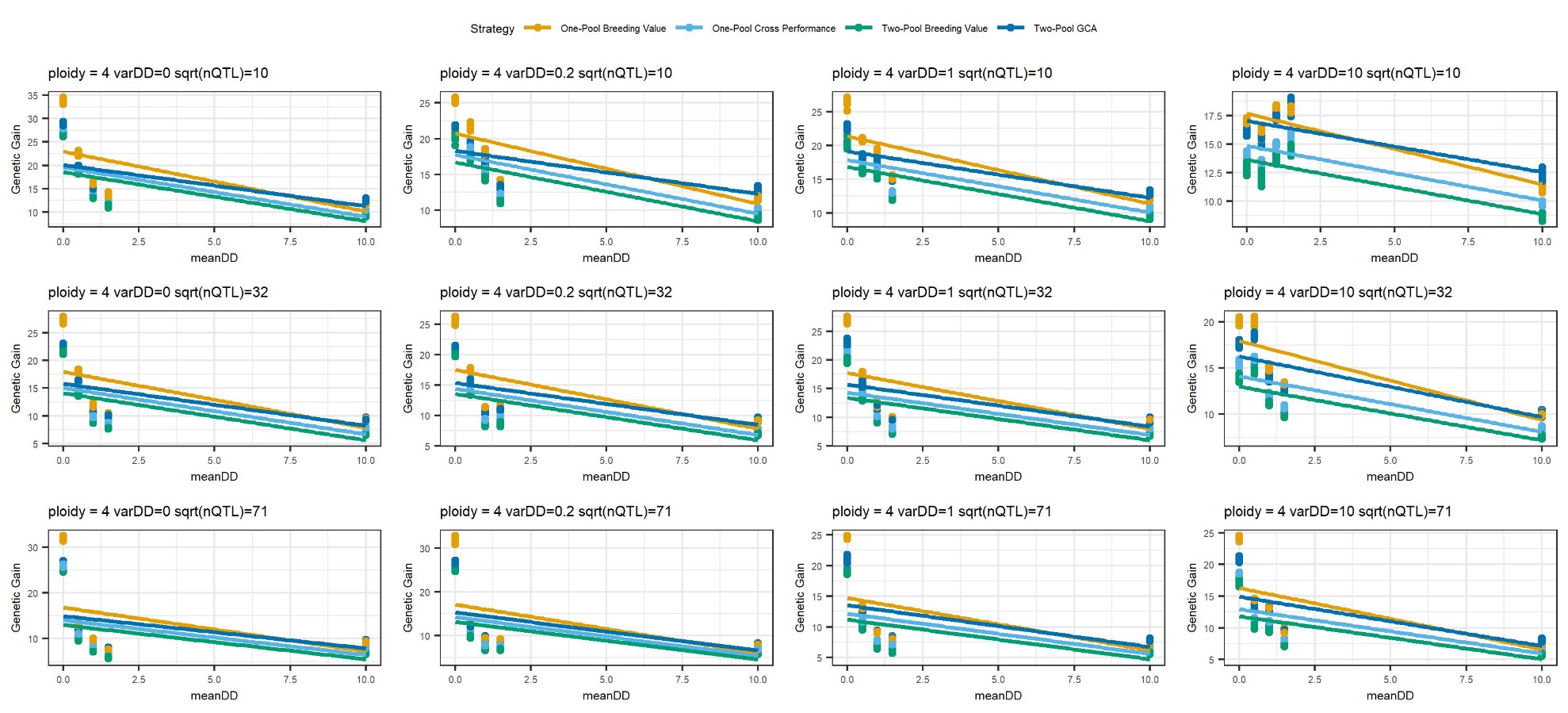

Supplemental Figure 20. For use of true values at high intensity after 50 years in autotetraploids, the relative performance of the core strategies as a function of mean dominance degree (meanDD) instead of H_0_, at each level of the variance of dominance degrees (varDD) and the square root of the number of QTL per chromosome (sqrt(nQTL)).

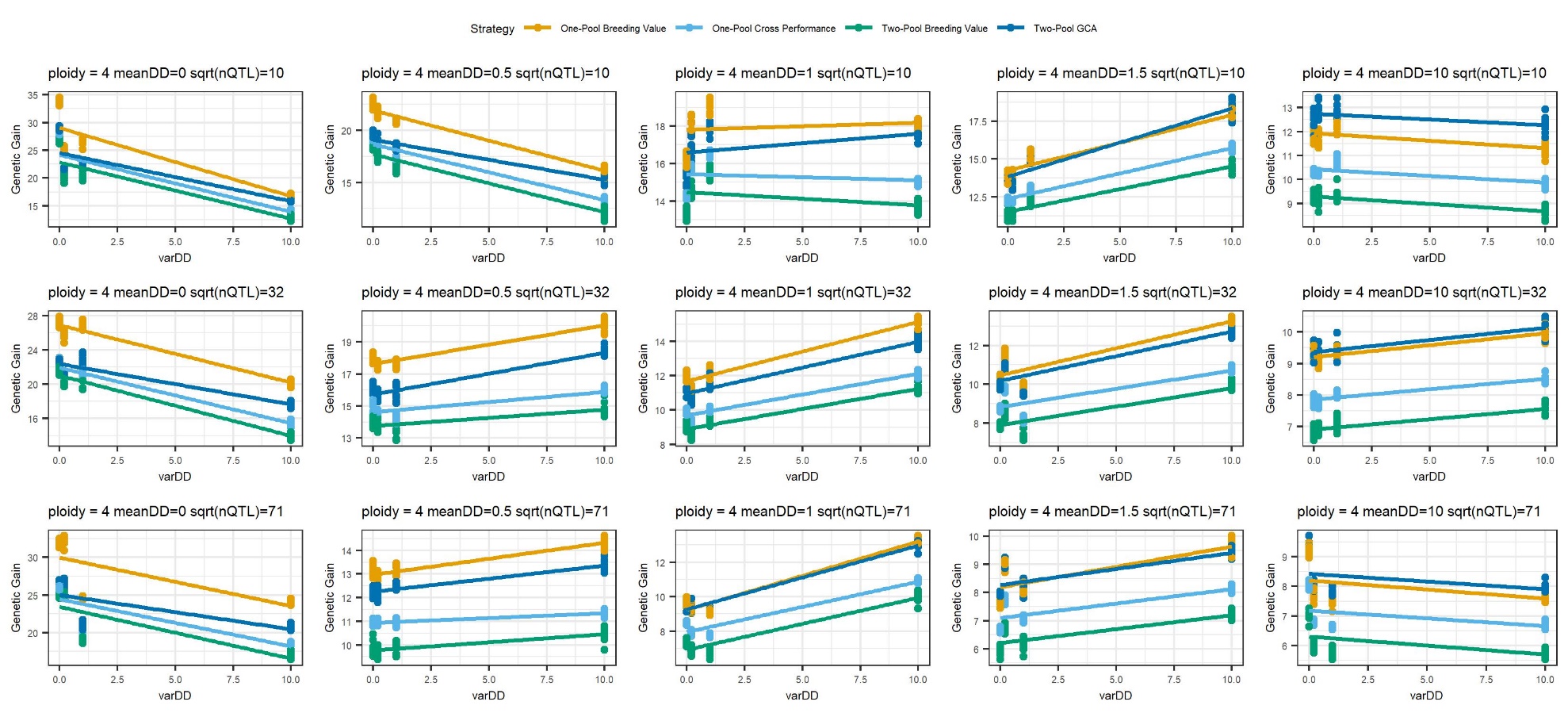

Supplemental Figure 21. For use of true values at high intensity after 50 years in autotetraploids, the relative performance of the core strategies as a function of the variance of dominance degrees (varDD) instead of H_0_, at each level of the mean dominance degree (meanDD) and the square root of the number of QTL per chromosome (sqrt(nQTL)).

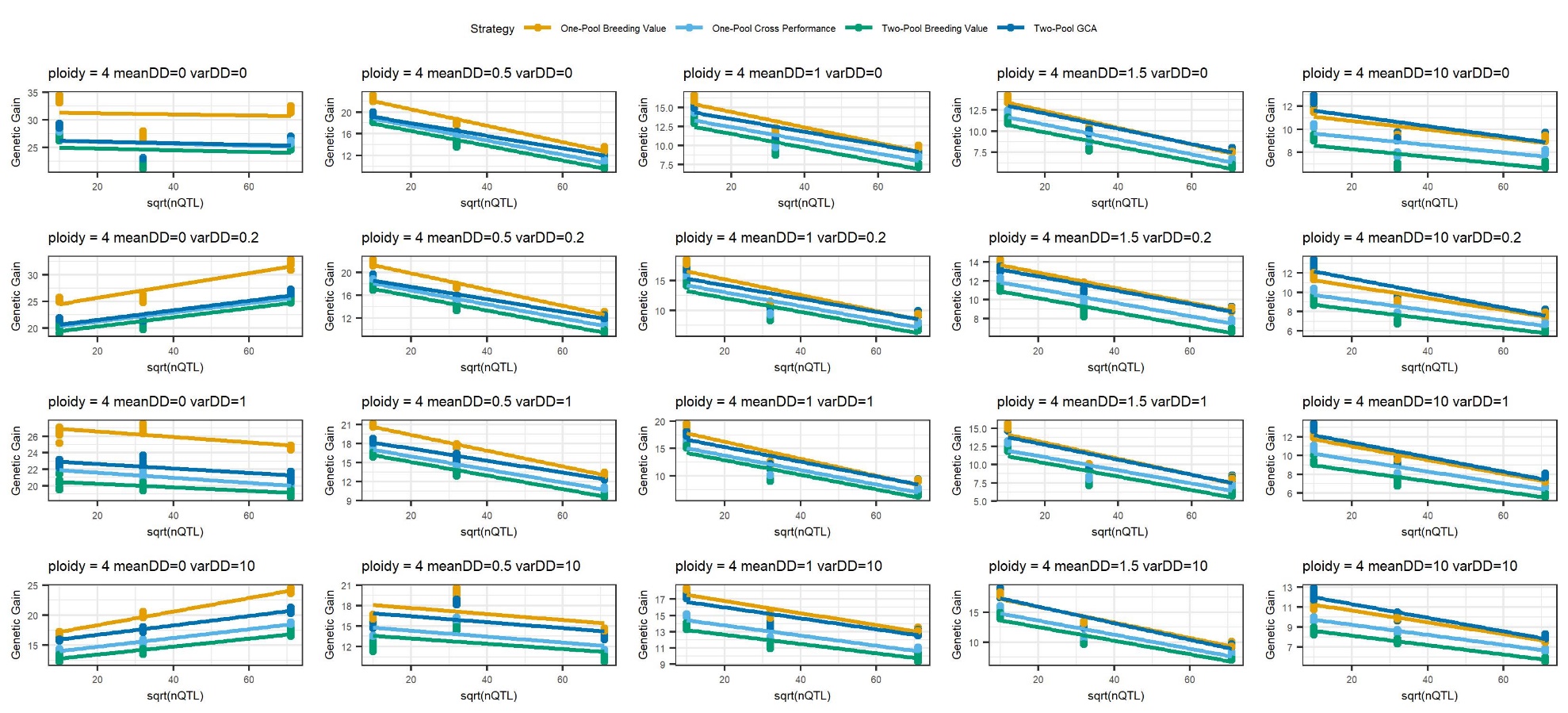

Supplemental Figure 22. For use of true values at high intensity after 50 years in autotetraploids, the relative performance of the core strategies as a function of the square root of the number of QTL per chromosome (sqrt(nQTL) instead of H_0_, at each level of the variance of dominance degrees (varDD) and the mean dominance degree (meanDD).

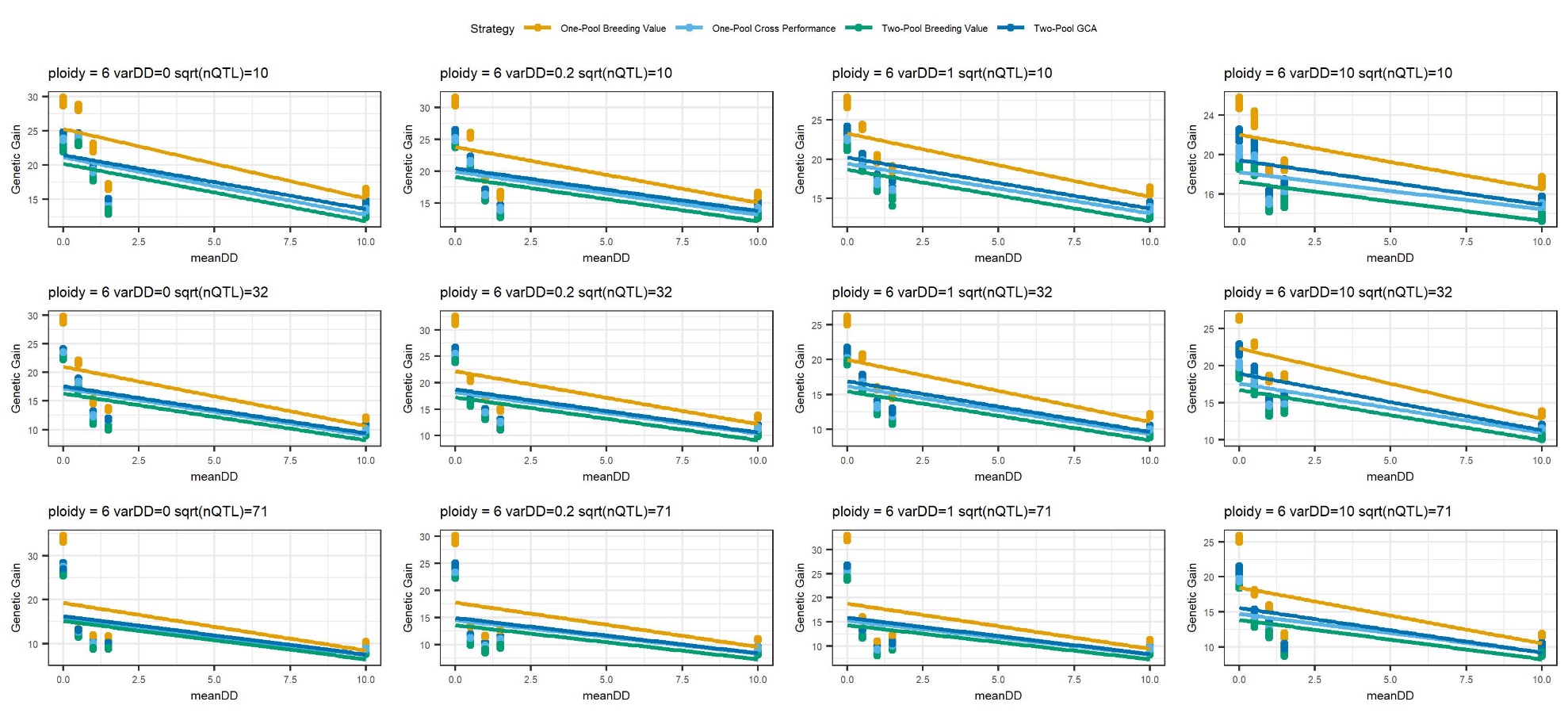

Supplemental Figure 23. For use of true values at high intensity after 50 years in autohexaploids, the relative performance of the core strategies as a function of mean dominance degree (meanDD) instead of H_0_, at each level of the variance of dominance degrees (varDD) and the square root of the number of QTL per chromosome (sqrt(nQTL)).

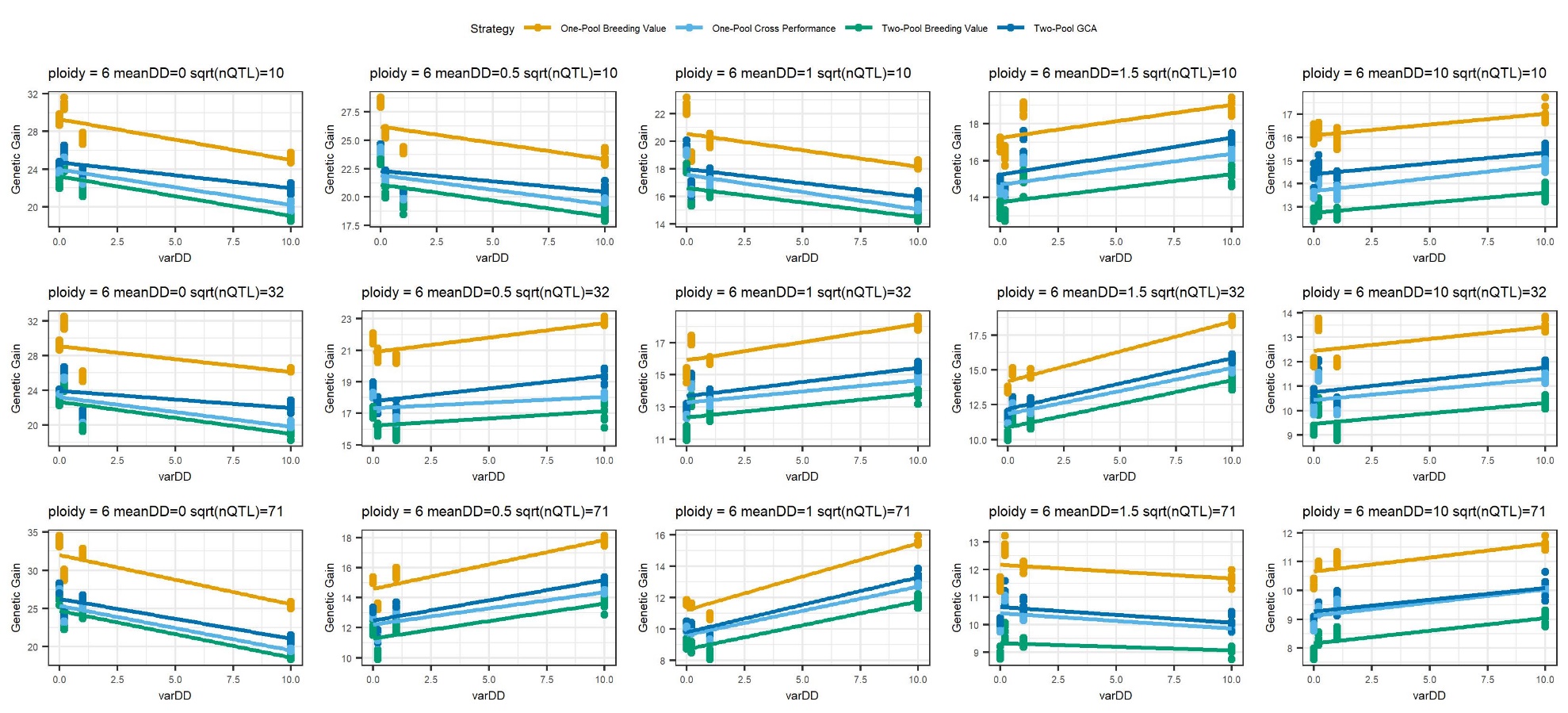

Supplemental Figure 24. For use of true values at high intensity after 50 years in autohexaploids, the relative performance of the core strategies as a function of the variance of dominance degrees (varDD) instead of H_0_, at each level of the mean dominance degree (meanDD) and the square root of the number of QTL per chromosome (sqrt(nQTL)).

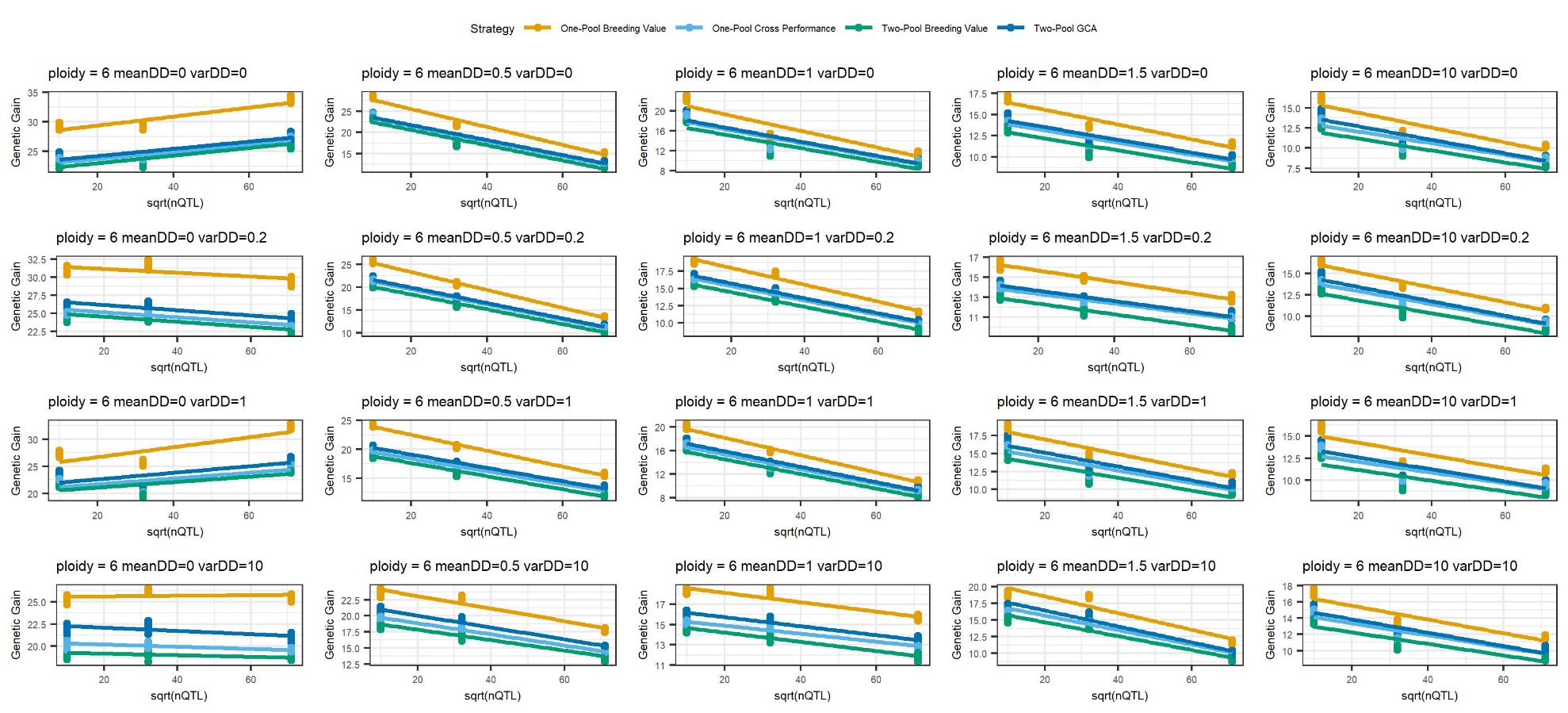

Supplemental Figure 25. For use of true values at high intensity after 50 years in autohexaploids, the relative performance of the core strategies as a function of the square root of the number of QTL per chromosome (sqrt(nQTL) instead of H_0_, at each level of the variance of dominance degrees (varDD) and the mean dominance degree (meanDD).
