## Supplemental Tables for "Clonal breeding strategies to harness heterosis: insights from stochastic simulation"

**Supplemental Table 1.** Program sizes for given strategies, estimation methods, and selection intensities.

| **True, Low Intensity** | One-Pool Breeding Value | One-Pool Predicted Cross Performance | Two-Pool Breeding Value | Two-Pool GCA | Two-Pool Breeding Value + GCA | Two-Pool Doubled Haploid GCA | Two-Pool Doubled Haploid Breeding Value + GCA |
| --- | --- | --- | --- | --- | --- | --- | --- |
| Number of Pools | 1 | 1 | 2 | 2 | 2 | 2 | 2 |
| Number of Intrapool Crosses per Pool, *x* | 100 | 100 | 50 | 50 | 50 | 50 | 40 |
| Number of Progeny per Intrapool Cross, *y* | 13 | 13 | 4 | 5 | 5 | 3 | 4 |
| Total Number of Intrapool Progeny, *z* | 1300 | 1300 | 400 | 500 | 500 | 300 | 320 |
| Total Number of Interpool Parents, *v* | 0 | 0 | 400 | 500 | 400 | 300 | 240 |
| Total Number of Interpool Crosses | 0 | 0 | 800 | 1000 | 800 | 600 | 480 |
| Number of Progeny per Interpool Cross | 0 | 0 | 1 | 1 | 1 | 1 | 1 |
| Total Number of Interpool Progeny, *w* | 0 | 0 | 800 | 1000 | 800 | 600 | 480 |
| Total Number of Evaluation Plots | 1300 | 1300 | 1200 | 1000 | 1300 | 600 | 800 |
| Total Number of Doubled Haploid Plots | 0 | 0 | 0 | 0 | 0 | 900 | 960 |
| Total Number of Genotyping Plots | 0 | 0 | 0 | 0 | 0 | 0 | 0 |
| Total Cost (Plots) | 1300 | 1300 | 1200 | 1000 | 1300 | 1500 | 1760 |

| **True, High Intensity** | One-Pool Breeding Value | One-Pool Predicted Cross Performance | Two-Pool Breeding Value | Two-Pool GCA | Two-Pool Breeding Value + GCA | Two-Pool Doubled Haploid GCA | Two-Pool Doubled Haploid Breeding Value + GCA |
| --- | --- | --- | --- | --- | --- | --- | --- |
| Number of Pools | 1 | 1 | 2 | 2 | 2 | 2 | 2 |
| Number of Intrapool Crosses per Pool, *x* | 20 | 20 | 10 | 10 | 10 | 10 | 10 |
| Number of Progeny per Intrapool Cross, *y* | 65 | 65 | 20 | 25 | 25 | 15 | 16 |
| Total Number of Intrapool Progeny, *z* | 1300 | 1300 | 400 | 500 | 500 | 300 | 320 |
| Total Number of Interpool Parents, *v* | 0 | 0 | 400 | 500 | 400 | 300 | 240 |
| Total Number of Interpool Crosses | 0 | 0 | 800 | 1000 | 800 | 600 | 480 |
| Number of Progeny per Interpool Cross | 0 | 0 | 1 | 1 | 1 | 1 | 1 |
| Total Number of Interpool Progeny, *w* | 0 | 0 | 800 | 1000 | 800 | 600 | 480 |
| Total Number of Evaluation Plots | 1300 | 1300 | 1200 | 1000 | 1300 | 600 | 800 |
| Total Number of Doubled Haploid Plots | 0 | 0 | 0 | 0 | 0 | 900 | 960 |
| Total Number of Genotyping Plots | 0 | 0 | 0 | 0 | 0 | 0 | 0 |
| Total Cost (plot equivalents) | 1300 | 1300 | 1200 | 1000 | 1300 | 1500 | 1760 |

| **Genomic Estimated, Low Intensity** | One-Pool Breeding Value | One-Pool Predicted Cross Performance | Two-Pool Breeding Value | Two-Pool GCA | Two-Pool Breeding Value + GCA | Two-Pool Doubled Haploid GCA | Two-Pool Doubled Haploid Breeding Value + GCA |
| --- | --- | --- | --- | --- | --- | --- | --- |
| Number of Pools | 1 | 1 | 2 | 2 | 2 | 2 | 2 |
| Number of Intrapool Crosses per Pool, *x* | 100 | 100 | 50 | 50 | 50 | 50 | 40 |
| Number of Progeny per Intrapool Cross, *y* | 13 | 13 | 4 | 5 | 5 | 3 | 4 |
| Total Number of Intrapool Progeny, *z* | 1300 | 1300 | 400 | 500 | 500 | 300 | 320 |
| Total Number of Interpool Parents, *v* | 0 | 0 | 400 | 500 | 400 | 300 | 240 |
| Total Number of Interpool Crosses | 0 | 0 | 800 | 1000 | 800 | 600 | 480 |
| Number of Progeny per Interpool Cross | 0 | 0 | 1 | 1 | 1 | 1 | 1 |
| Total Number of Interpool Progeny, *w* | 0 | 0 | 800 | 1000 | 800 | 600 | 480 |
| Total Number of Evaluation Plots | 1300 | 1300 | 1200 | 1000 | 1300 | 600 | 800 |
| Total Number of Doubled Haploid Plots | 0 | 0 | 0 | 0 | 0 | 900 | 960 |
| Total Number of Genotyping Plots | 1300 | 1300 | 1200 | 1500 | 1300 | 900 | 800 |
| Total Cost (plot equivalents) | 2600 | 2600 | 2400 | 2500 | 2600 | 2400 | 2560 |

| **Genomic Estimated, High Intensity** | One-Pool Breeding Value | One-Pool Predicted Cross Performance | Two-Pool Breeding Value | Two-Pool GCA | Two-Pool Breeding Value + GCA | Two-Pool Doubled Haploid GCA | Two-Pool Doubled Haploid Breeding Value + GCA |
| --- | --- | --- | --- | --- | --- | --- | --- |
| Number of Pools | 1 | 1 | 2 | 2 | 2 | 2 | 2 |
| Number of Intrapool Crosses per Pool, *x* | 20 | 20 | 10 | 10 | 10 | 10 | 10 |
| Number of Progeny per Intrapool Cross, *y* | 65 | 65 | 20 | 25 | 25 | 15 | 16 |
| Total Number of Intrapool Progeny, *z* | 1300 | 1300 | 400 | 500 | 500 | 300 | 320 |
| Total Number of Interpool Parents, *v* | 0 | 0 | 400 | 500 | 400 | 300 | 240 |
| Total Number of Interpool Crosses | 0 | 0 | 800 | 1000 | 800 | 600 | 480 |
| Number of Progeny per Interpool Cross | 0 | 0 | 1 | 1 | 1 | 1 | 1 |
| Total Number of Interpool Progeny, *w* | 0 | 0 | 800 | 1000 | 800 | 600 | 480 |
| Total Number of Evaluation Plots | 1300 | 1300 | 1200 | 1000 | 1300 | 600 | 800 |
| Total Number of Doubled Haploid Plots | 0 | 0 | 0 | 0 | 0 | 900 | 960 |
| Total Number of Genotyping Plots | 1300 | 1300 | 1200 | 1500 | 1300 | 900 | 800 |
| Total Cost (plot equivalents) | 2600 | 2600 | 2400 | 2500 | 2600 | 2400 | 2560 |

| **Phenotypic, Low Intensity** | One-Pool Phenotypic Value | One-Pool Cross Performance | Two-Pool Phenotypic Value | Two-Pool GCA | Two-Pool Phenotypic Value + GCA | Two-Pool Doubled Haploid GCA | Two-Pool Doubled Haploid Phenotypic Value + GCA |
| --- | --- | --- | --- | --- | --- | --- | --- |
| Number of Pools | 1 | 1 | 2 | 2 | 2 | 2 | 2 |
| Number of Intrapool Crosses per Pool, *x* | 100 | 100 | 50 | 50 | 50 | 50 | 50 |
| Number of Progeny per Intrapool Cross, *y* | 26 | 26 | 9 | 13 | 13 | 5 | 5 |
| Total Number of Intrapool Progeny, *z* | 2600 | 2600 | 900 | 1300 | 1300 | 500 | 500 |
| Total Number of Interpool Parents, *v* | 0 | 0 | 900 | 1300 | 600 | 500 | 300 |
| Total Number of Interpool Crosses | 0 | 0 | 1800 | 2600 | 1200 | 1000 | 600 |
| Number of Progeny per Interpool Cross | 0 | 0 | 1 | 1 | 1 | 1 | 1 |
| Total Number of Interpool Progeny, *w* | 0 | 0 | 1800 | 2600 | 1200 | 1000 | 600 |
| Total Number of Evaluation Plots | 2600 | 2600 | 2700 | 2600 | 2500 | 1000 | 1100 |
| Total Number of Doubled Haploid Plots | 0 | 0 | 0 | 0 | 0 | 1500 | 1500 |
| Total Number of Genotyping Plots | 0 | 0 | 0 | 0 | 0 | 0 | 0 |
| Total Cost (plot equivalents) | 2600 | 2600 | 2700 | 2600 | 2500 | 2500 | 2600 |

| **Phenotypic, High Intensity** | One-Pool Phenotypic Value | Two-Pool Phenotypic Value | Two-Pool GCA | Two-Pool Phenotypic Value + GCA | Two-Pool Doubled Haploid GCA | Two-Pool Doubled Haploid Phenotypic Value + GCA |
| --- | --- | --- | --- | --- | --- | --- |
| Number of Pools | 1 | 2 | 2 | 2 | 2 | 2 |
| Number of Intrapool Crosses per Pool, *x* | 20 | 10 | 10 | 10 | 10 | 10 |
| Number of Progeny per Intrapool Cross, *y* | 130 | 42 | 63 | 50 | 25 | 23 |
| Total Number of Intrapool Progeny, *z* | 2600 | 840 | 1260 | 1000 | 500 | 460 |
| Total Number of Interpool Parents, *v* | 0 | 840 | 1260 | 760 | 500 | 360 |
| Total Number of Interpool Crosses | 0 | 1680 | 2520 | 1520 | 1000 | 720 |
| Number of Progeny per Interpool Cross | 0 | 1 | 1 | 1 | 1 | 1 |
| Total Number of Interpool Progeny, *w* | 0 | 1680 | 2520 | 1520 | 1000 | 720 |
| Total Number of Evaluation Plots | 2600 | 2520 | 2520 | 2520 | 1000 | 1180 |
| Total Number of Doubled Haploid Plots | 0 | 0 | 0 | 0 | 1500 | 1380 |
| Total Number of Genotyping Plots | 0 | 0 | 0 | 0 | 0 | 0 |
| Total Cost (plot equivalents) | 2600 | 2520 | 2520 | 2520 | 2500 | 2560 |

**Supplemental Table 2**. Genotype frequencies for diploids, autotetraploids, and autohexaploids at Hardy-Weinberg equilibrium if allele frequencies *p* = *q* = 0.5. As ploidy increases, the frequency of homozygous genotypes decreases.

| **Diploid Genotype** | **Diploid Genotype Frequency*** | **Autotetraploid Genotype** | **Autotetraploid Genotype Frequency*** | **Autohexaploid Genotype** | **Autohexaploid Genotype Frequency*** |
| --- | --- | --- | --- | --- | --- |
| 0 | 0.25 | 0 | 0.06 | 0 | 0.02 |
|  |  |  |  | 1 | 0.09 |
|  |  | 1 | 0.25 | 2 | 0.23 |
| 1 | 0.5 | 2 | 0.38 | 3 | 0.31 |
|  |  | 3 | 0.25 | 4 | 0.23 |
|  |  |  |  | 5 | 0.09 |
| 2 | 0.25 | 4 | 0.06 | 6 | 0.02 |

***With *p* = *q* = 0.5 at Hardy-Weinberg equilibrium
