## Supplementary material for "Clonal breeding strategies to harness heterosis: insights from stochastic simulation": Tables

**Table 1.** Sets of cycle lengths assumed to be required for each strategy in a given scenario.

|  | **True Values L = 2** | **Genomic Estimated Values L = 2** | **Phenotypic Values, Fast Multiplication** | **Phenotypic Values, Slow Multiplication** |
| --- | --- | --- | --- | --- |
| One-Pool Breeding Value | 2 | 2 | 3 | 5 |
| One-Pool Cross Performance | 2 | 2 |  |  |
| Two-Pool Breeding Value | 2 | 2 | 3 | 5 |
| Two-Pool GCA | 2 | 2 | 4 | 8 |
| Two-Pool Doubled Haploid GCA | 2 | 2 | 5 | 9 |
| Two-Pool Breeding Value + GCA | 2 | 2 | 4 | 8 |
| Two-Pool Doubled Haploid Breeding Value + GCA | 2 | 2 | 5 | 9 |
