## Supplemental File 44 for "Clonal breeding strategies to harness heterosis: insights from stochastic simulation"

**Table of Contents, Additional Plots**

For all plots, initial population heterosis (H_0_) is scaled to initial population genetic standard deviation as described in the main manuscript, and each response is scaled as appropriate to initial population genetic standard deviation (genetic gain, additive value, dominance value, inbreeding depression, panmictic heterosis) or initial population total genetic variance (total genetic variance, additive variance, dominance variance) or with no scaling (accuracy). For each plot, regressions of response value by scenario are plotted as a function of H_0_, scenario, and their interaction.

1. Diploid genetic gain after 15 years, core strategies
2. Diploid additive value after 15 years, core strategies
3. Diploid dominance value after 15 years, core strategies
4. Diploid total genetic variance after 15 years, core strategies
5. Diploid additive genetic variance after 15 years, core strategies
6. Diploid dominance genetic variance after 15 years, core strategies
7. Diploid selection accuracy after 15 years, core strategies
8. Diploid inbreeding depression after 15 years, core strategies
9. Diploid panmictic heterosis after 15 years, core strategies
10. Diploid intra-pool genetic gain after 15 years, core strategies
11. Diploid intra-pool additive value after 15 years, core strategies
12. Diploid intra-pool dominance value after 15 years, core strategies
13. Diploid genetic gain after 50 years, core strategies
14. Diploid additive value after 50 years, core strategies
15. Diploid dominance value after 50 years, core strategies
16. Diploid total genetic variance after 50 years, core strategies
17. Diploid additive genetic variance after 50 years, core strategies
18. Diploid dominance genetic variance after 50 years, core strategies
19. Diploid selection accuracy after 50 years, core strategies
20. Diploid inbreeding depression after 50 years, core strategies
21. Diploid panmictic heterosis after 50 years, core strategies
22. Diploid intra-pool genetic gain after 50 years, core strategies
23. Diploid intra-pool additive value after 50 years, core strategies
24. Diploid intra-pool dominance value after 50 years, core strategies
25. Autotetraploid genetic gain after 15 years, core strategies
26. Autotetraploid additive value after 15 years, core strategies
27. Autotetraploid dominance value after 15 years, core strategies
28. Autotetraploid total genetic variance after 15 years, core strategies
29. Autotetraploid additive genetic variance after 15 years, core strategies
30. Autotetraploid dominance genetic variance after 15 years, core strategies
31. Autotetraploid selection accuracy after 15 years, core strategies
32. Autotetraploid inbreeding depression after 15 years, core strategies
33. Autotetraploid panmictic heterosis after 15 years, core strategies
34. Autotetraploid intra-pool genetic gain after 15 years, core strategies
35. Autotetraploid intra-pool additive value after 15 years, core strategies
36. Autotetraploid intra-pool dominance value after 15 years, core strategies
37. Autotetraploid genetic gain after 50 years, core strategies
38. Autotetraploid additive value after 50 years, core strategies
39. Autotetraploid dominance value after 50 years, core strategies
40. Autotetraploid total genetic variance after 50 years, core strategies
41. Autotetraploid additive genetic variance after 50 years, core strategies
42. Autotetraploid dominance genetic variance after 50 years, core strategies
43. Autotetraploid selection accuracy after 50 years, core strategies
44. Autotetraploid inbreeding depression after 50 years, core strategies
45. Autotetraploid panmictic heterosis after 50 years, core strategies
46. Autotetraploid intra-pool genetic gain after 50 years, core strategies
47. Autotetraploid intra-pool additive value after 50 years, core strategies
48. Autotetraploid intra-pool dominance value after 50 years, core strategies
49. Autohexaploid genetic gain after 15 years, core strategies
50. Autohexaploid additive value after 15 years, core strategies
51. Autohexaploid dominance value after 15 years, core strategies
52. Autohexaploid total genetic variance after 15 years, core strategies
53. Autohexaploid additive genetic variance after 15 years, core strategies
54. Autohexaploid dominance genetic variance after 15 years, core strategies
55. Autohexaploid selection accuracy after 15 years, core strategies
56. Autohexaploid inbreeding depression after 15 years, core strategies
57. Autohexaploid panmictic heterosis after 15 years, core strategies
58. Autohexaploid intra-pool genetic gain after 15 years, core strategies
59. Autohexaploid intra-pool additive value after 15 years, core strategies
60. Autohexaploid intra-pool dominance value after 15 years, core strategies
61. Autohexaploid genetic gain after 50 years, core strategies
62. Autohexaploid additive value after 50 years, core strategies
63. Autohexaploid dominance value after 50 years, core strategies
64. Autohexaploid total genetic variance after 50 years, core strategies
65. Autohexaploid additive genetic variance after 50 years, core strategies
66. Autohexaploid dominance genetic variance after 50 years, core strategies
67. Autohexaploid selection accuracy after 50 years, core strategies
68. Autohexaploid inbreeding depression after 50 years, core strategies
69. Autohexaploid panmictic heterosis after 50 years, core strategies
70. Autohexaploid intra-pool genetic gain after 50 years, core strategies
71. Autohexaploid intra-pool additive value after 50 years, core strategies
72. Autohexaploid intra-pool dominance value after 50 years, core strategies
73. Diploid genetic gain after 15 years, non-core strategies
74. Diploid additive value after 15 years, non-core strategies
75. Diploid dominance value after 15 years, non-core strategies
76. Diploid total genetic variance after 15 years, non-core strategies
77. Diploid additive genetic variance after 15 years, non-core strategies
78. Diploid dominance genetic variance after 15 years, non-core strategies
79. Diploid selection accuracy after 15 years, non-core strategies
80. Diploid inbreeding depression after 15 years, non-core strategies
81. Diploid panmictic heterosis after 15 years, non-core strategies
82. Diploid intra-pool genetic gain after 15 years, non-core strategies
83. Diploid intra-pool additive value after 15 years, non-core strategies
84. Diploid intra-pool dominance value after 15 years, non-core strategies
85. Diploid genetic gain after 50 years, non-core strategies
86. Diploid additive value after 50 years, non-core strategies
87. Diploid dominance value after 50 years, non-core strategies
88. Diploid total genetic variance after 50 years, non-core strategies
89. Diploid additive genetic variance after 50 years, non-core strategies
90. Diploid dominance genetic variance after 50 years, non-core strategies
91. Diploid selection accuracy after 50 years, non-core strategies
92. Diploid inbreeding depression after 50 years, non-core strategies
93. Diploid panmictic heterosis after 50 years, non-core strategies
94. Diploid intra-pool genetic gain after 50 years, non-core strategies
95. Diploid intra-pool additive value after 50 years, non-core strategies
96. Diploid intra-pool dominance value after 50 years, non-core strategies
97. Autotetraploid genetic gain after 15 years, non-core strategies
98. Autotetraploid additive value after 15 years, non-core strategies
99. Autotetraploid dominance value after 15 years, non-core strategies
100. Autotetraploid total genetic variance after 15 years, non-core strategies
101. Autotetraploid additive genetic variance after 15 years, non-core strategies
102. Autotetraploid dominance genetic variance after 15 years, non-core strategies
103. Autotetraploid selection accuracy after 15 years, non-core strategies
104. Autotetraploid inbreeding depression after 15 years, non-core strategies
105. Autotetraploid panmictic heterosis after 15 years, non-core strategies
106. Autotetraploid intra-pool genetic gain after 15 years, non-core strategies
107. Autotetraploid intra-pool additive value after 15 years, non-core strategies
108. Autotetraploid intra-pool dominance value after 15 years, non-core strategies
109. Autotetraploid genetic gain after 50 years, non-core strategies
110. Autotetraploid additive value after 50 years, non-core strategies
111. Autotetraploid dominance value after 50 years, non-core strategies
112. Autotetraploid total genetic variance after 50 years, non-core strategies
113. Autotetraploid additive genetic variance after 50 years, non-core strategies
114. Autotetraploid dominance genetic variance after 50 years, non-core strategies
115. Autotetraploid selection accuracy after 50 years, non-core strategies
116. Autotetraploid inbreeding depression after 50 years, non-core strategies
117. Autotetraploid panmictic heterosis after 50 years, non-core strategies
118. Autotetraploid intra-pool genetic gain after 50 years, non-core strategies
119. Autotetraploid intra-pool additive value after 50 years, non-core strategies
120. Autotetraploid intra-pool dominance value after 50 years, core strategies
121. Autohexaploid genetic gain after 15 years, non-core strategies
122. Autohexaploid additive value after 15 years, non-core strategies
123. Autohexaploid dominance value after 15 years, non-core strategies
124. Autohexaploid total genetic variance after 15 years, non-core strategies
125. Autohexaploid additive genetic variance after 15 years, non-core strategies
126. Autohexaploid dominance genetic variance after 15 years, non-core strategies
127. Autohexaploid selection accuracy after 15 years, non-core strategies
128. Autohexaploid inbreeding depression after 15 years, non-core strategies
129. Autohexaploid panmictic heterosis after 15 years, non-core strategies
130. Autohexaploid intra-pool genetic gain after 15 years, non-core strategies
131. Autohexaploid intra-pool additive value after 15 years, non-core strategies
132. Autohexaploid intra-pool dominance value after 15 years, non-core strategies
133. Autohexaploid genetic gain after 50 years, non-core strategies
134. Autohexaploid additive value after 50 years, non-core strategies
135. Autohexaploid dominance value after 50 years, non-core strategies
136. Autohexaploid total genetic variance after 50 years, non-core strategies
137. Autohexaploid additive genetic variance after 50 years, non-core strategies
138. Autohexaploid dominance genetic variance after 50 years, non-core strategies
139. Autohexaploid selection accuracy after 50 years, non-core strategies
140. Autohexaploid inbreeding depression after 50 years, non-core strategies
141. Autohexaploid panmictic heterosis after 50 years, non-core strategies
142. Autohexaploid intra-pool genetic gain after 50 years, non-core strategies
143. Autohexaploid intra-pool additive value after 50 years, non-core strategies
144. Autohexaploid intra-pool dominance value after 50 years, non-core strategies
145. Mean inbreeding coefficient for true values after 15 years, core strategies
146. Mean inbreeding coefficient for true values after 50 years, core strategies
147. Mean inbreeding coefficient for true values after 15 years, non-core strategies
148. Mean inbreeding coefficient for true values after 50 years, non-core strategies
149. Diploid genetic gain after 15 years, GS vs. PS scenarios, core strategies
150. Diploid genetic gain after 50 years, GS vs. PS scenarios, core strategies
151. Autotetraploid genetic gain after 15 years, GS vs. PS scenarios, core strategies
152. Autotetraploid genetic gain after 50 years, GS vs. PS scenarios, core strategies
153. Autohexaploid genetic gain after 15 years, GS vs. PS scenarios, core strategies
154. Autohexaploid genetic gain after 50 years, GS vs. PS scenarios, core strategies

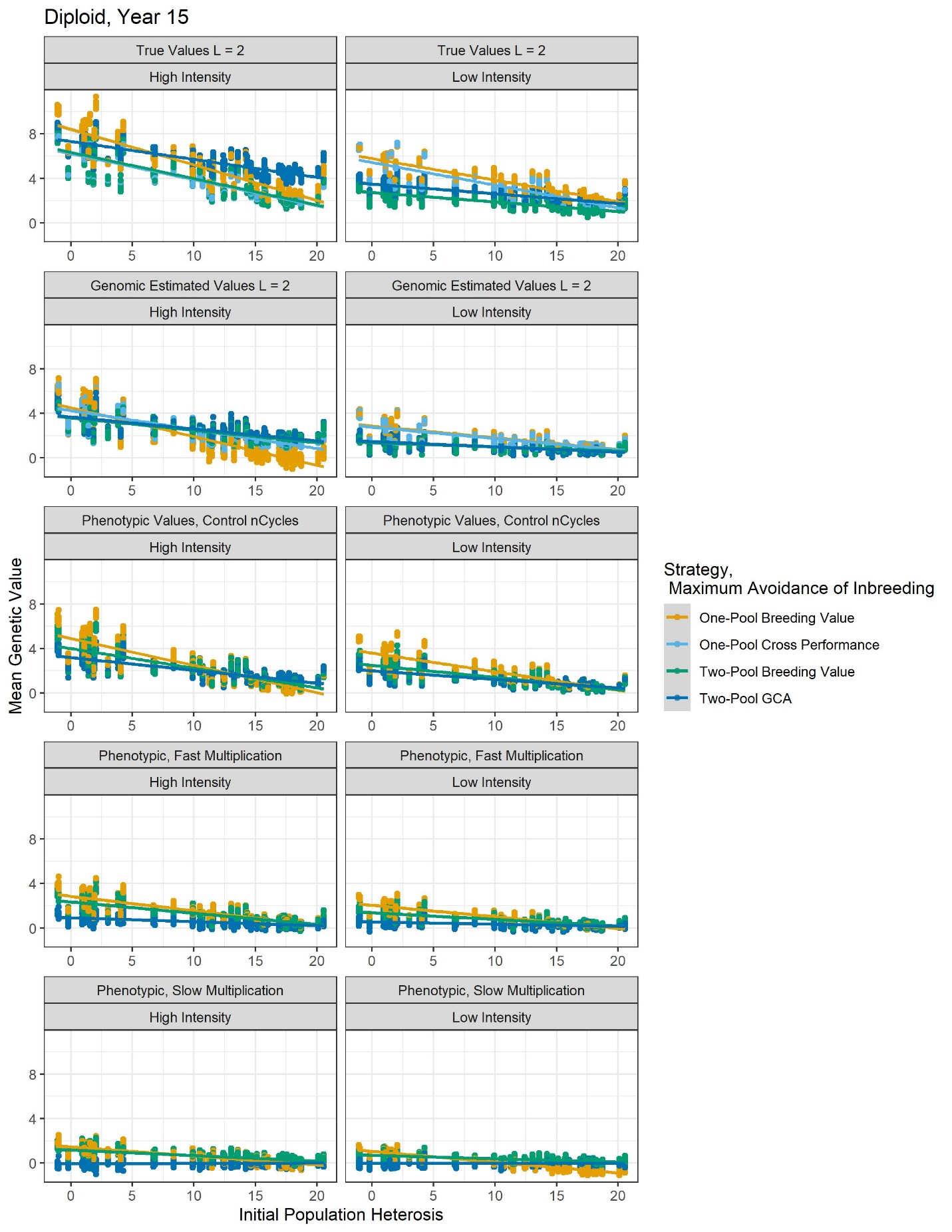
 Additional Plot 1. Diploid genetic gain after 15 years in the core strategies.

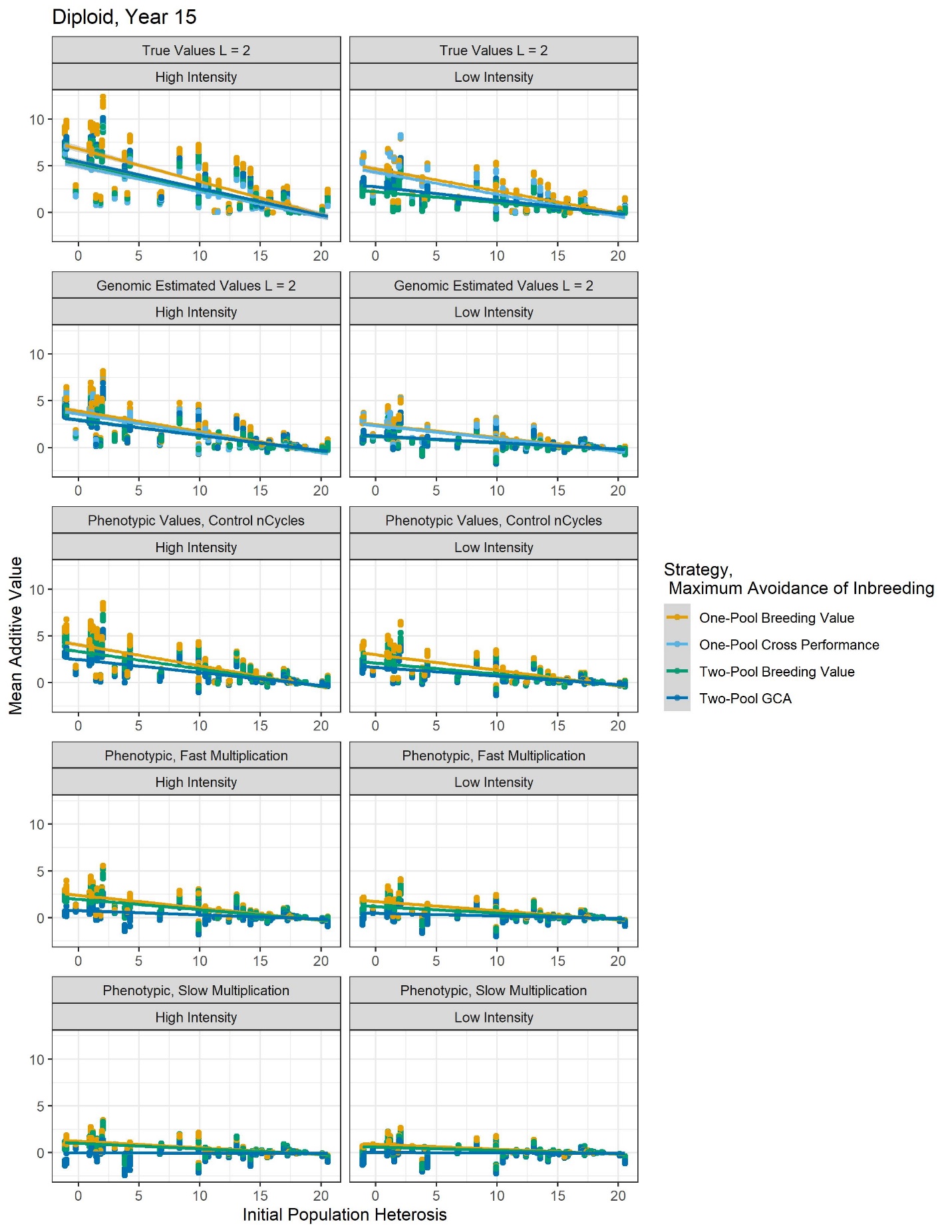

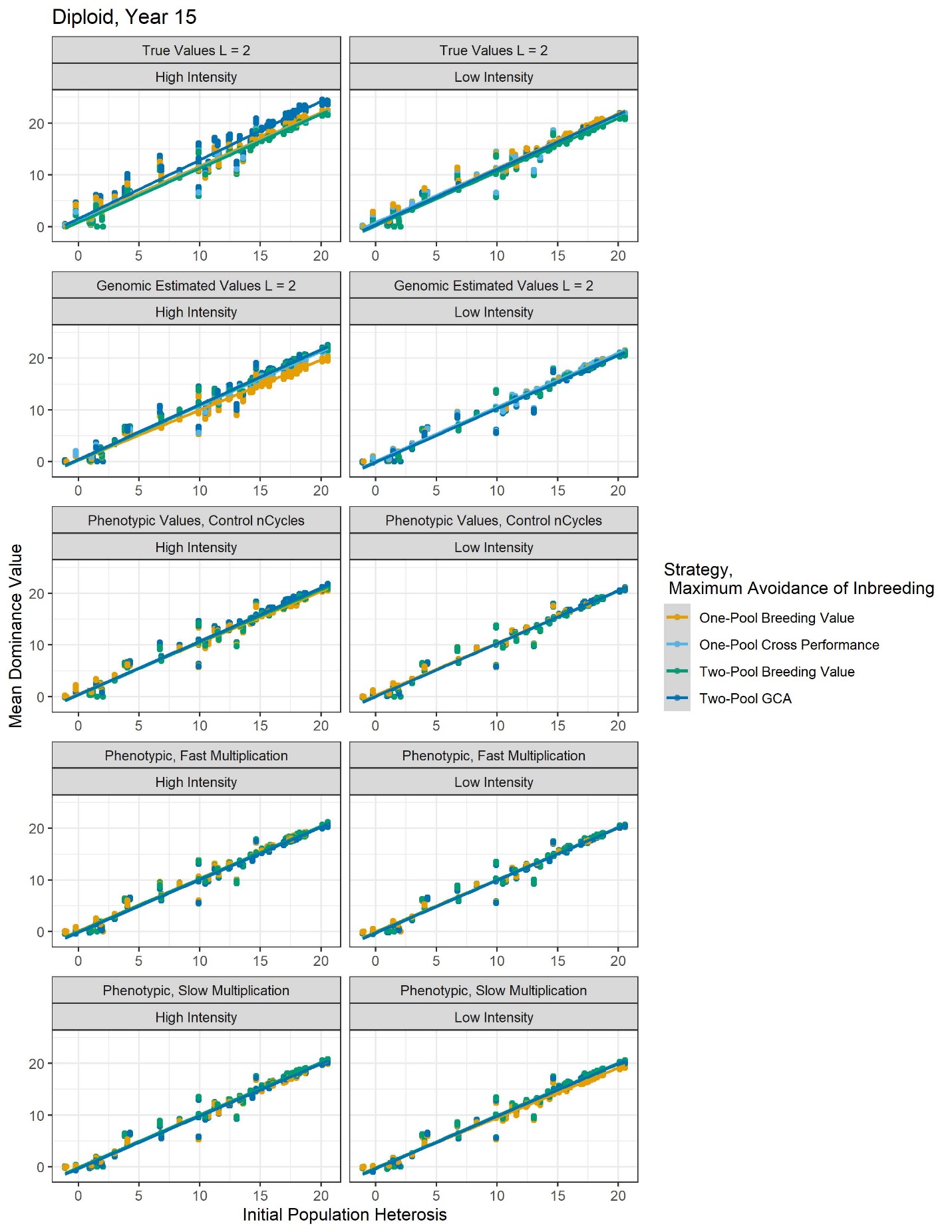

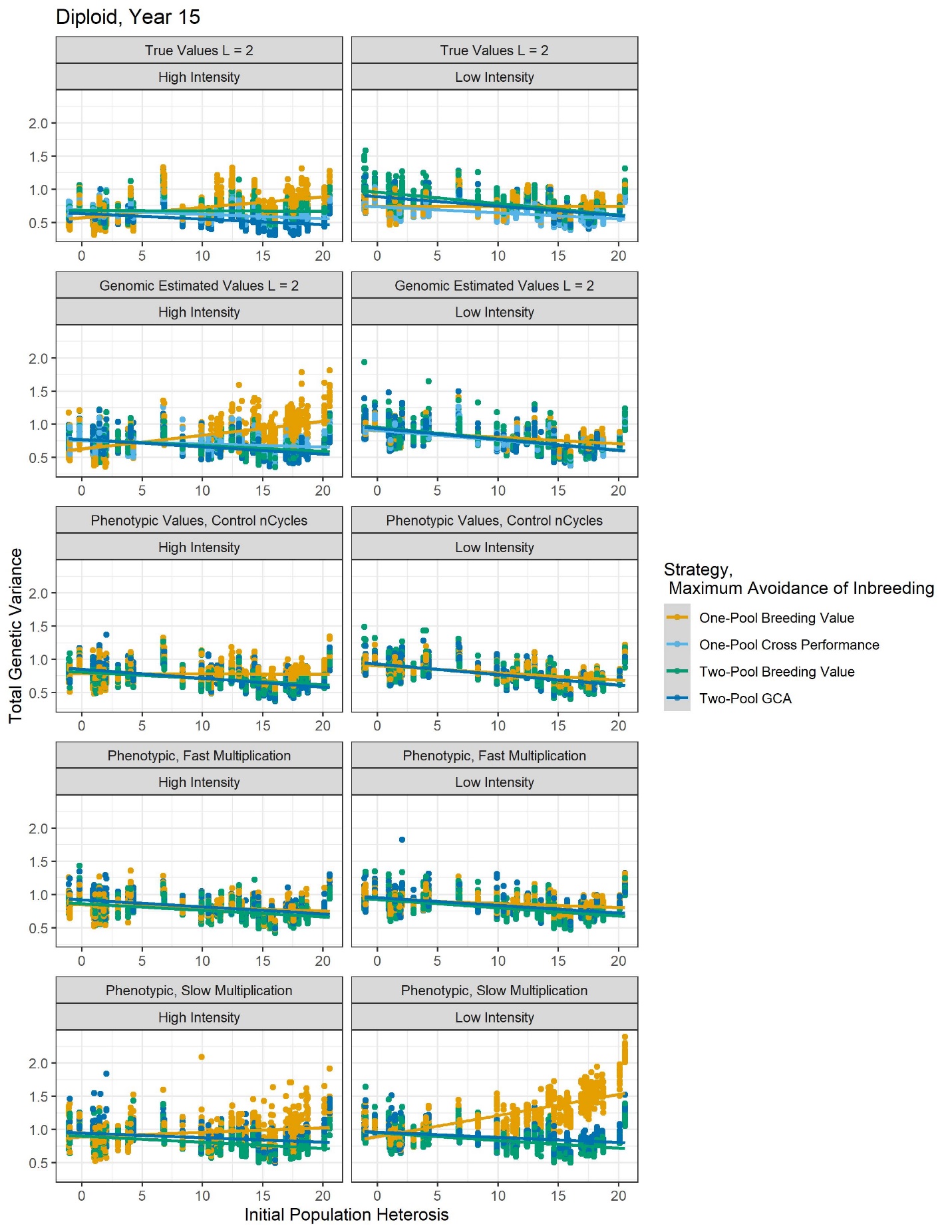

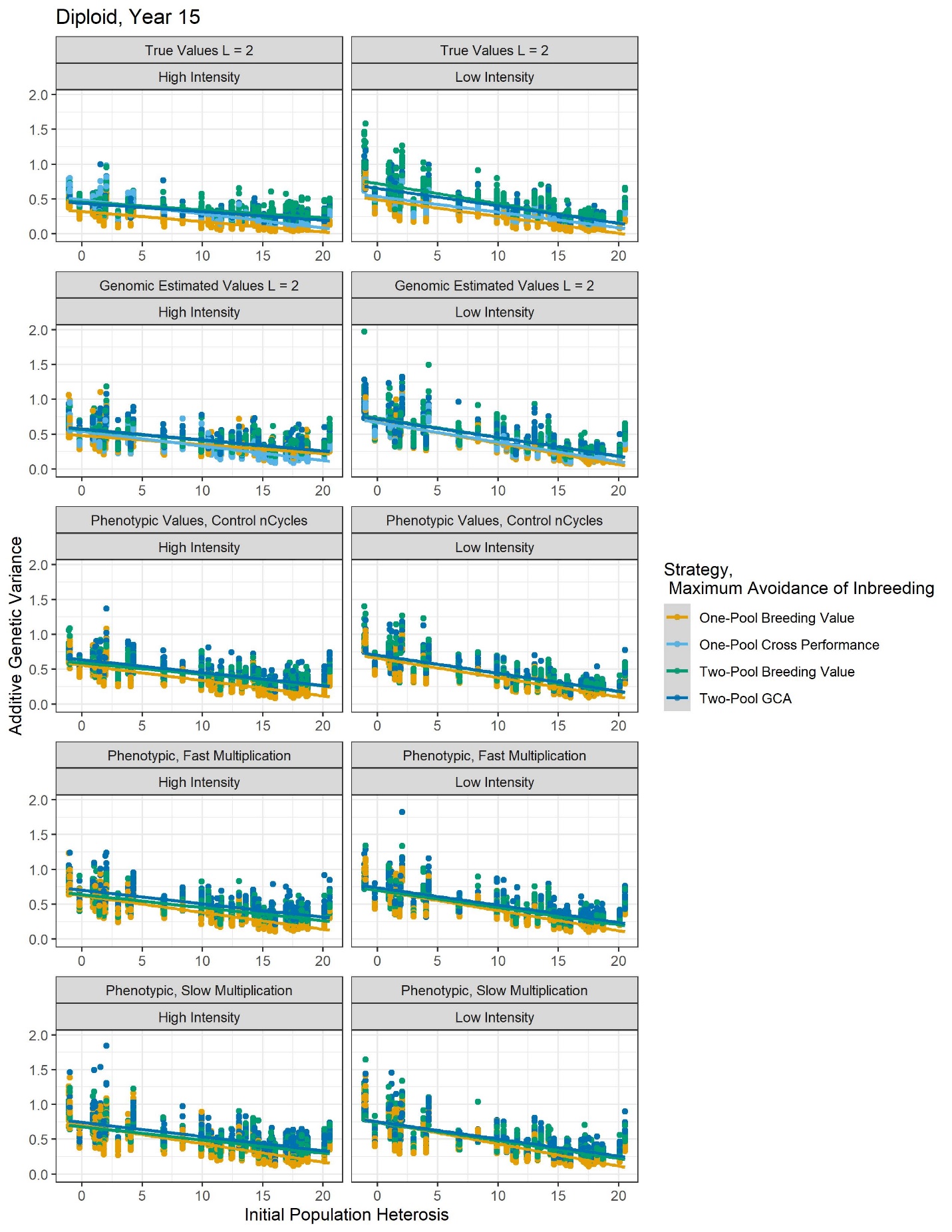

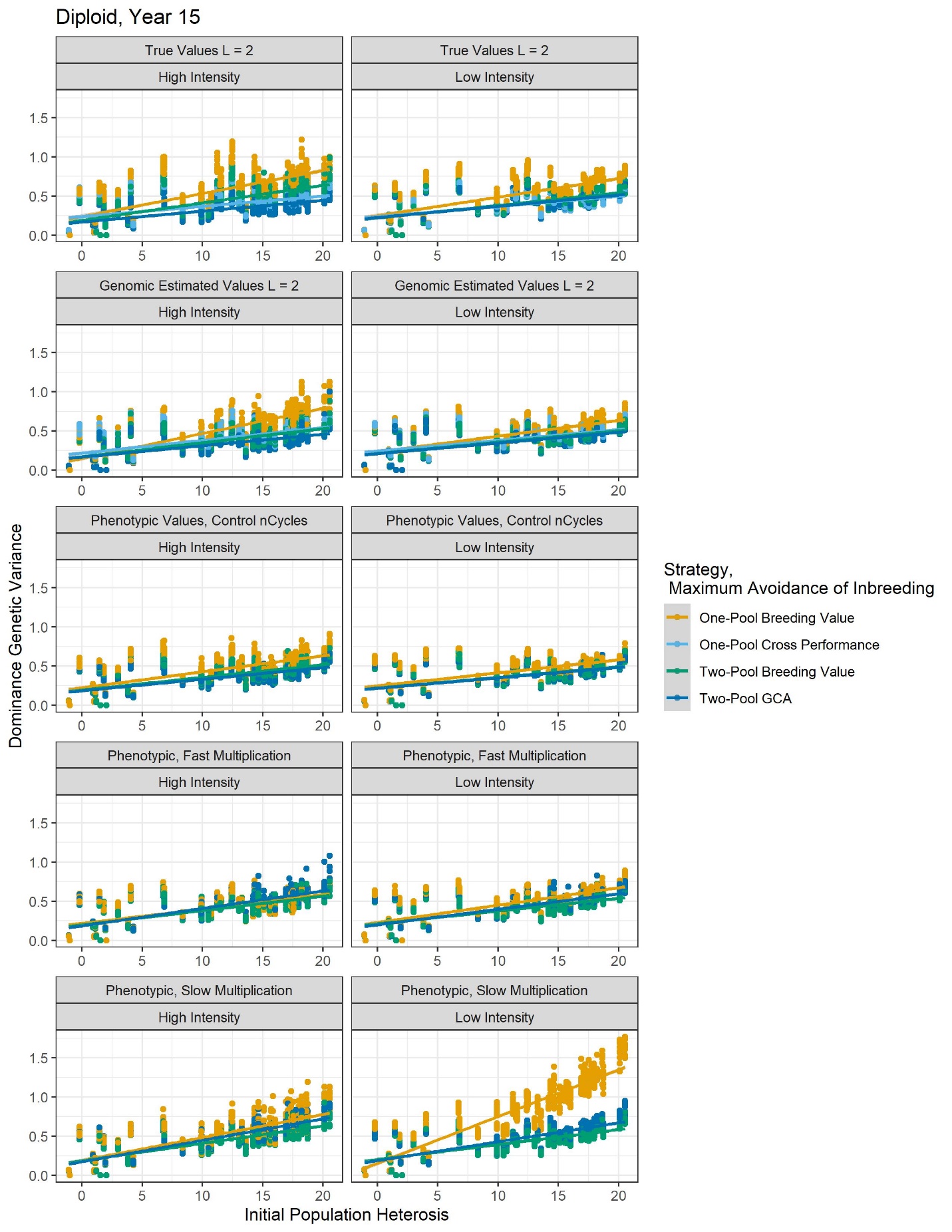

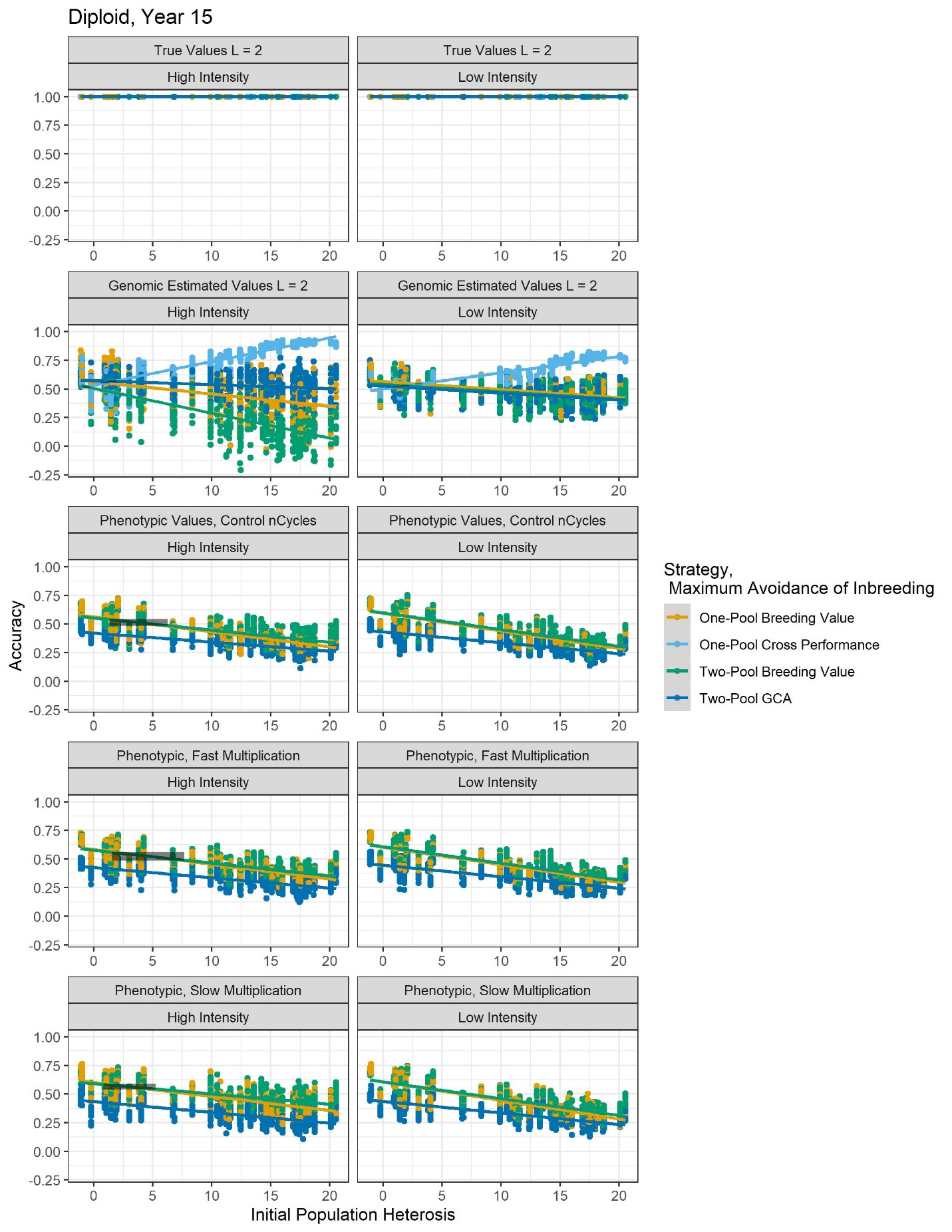

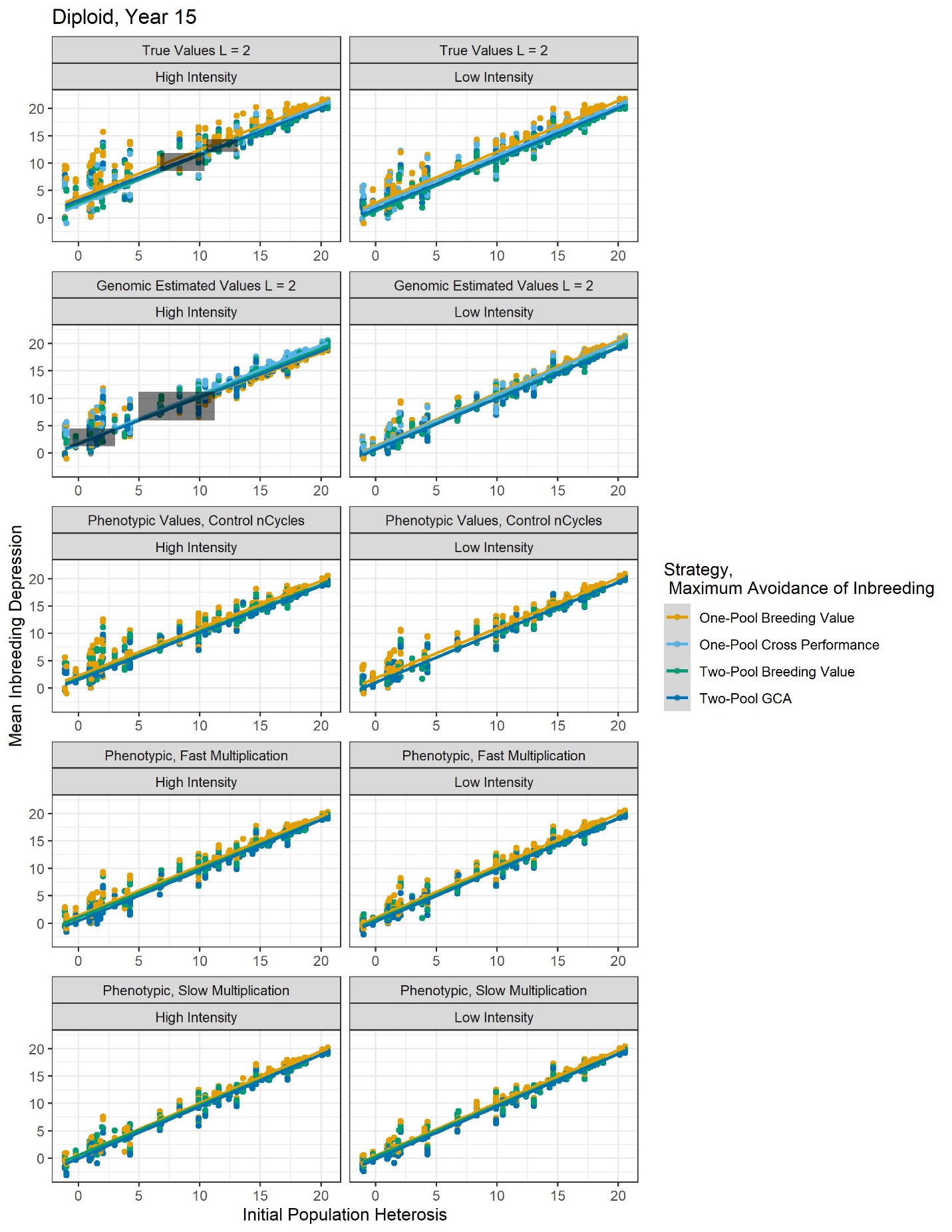

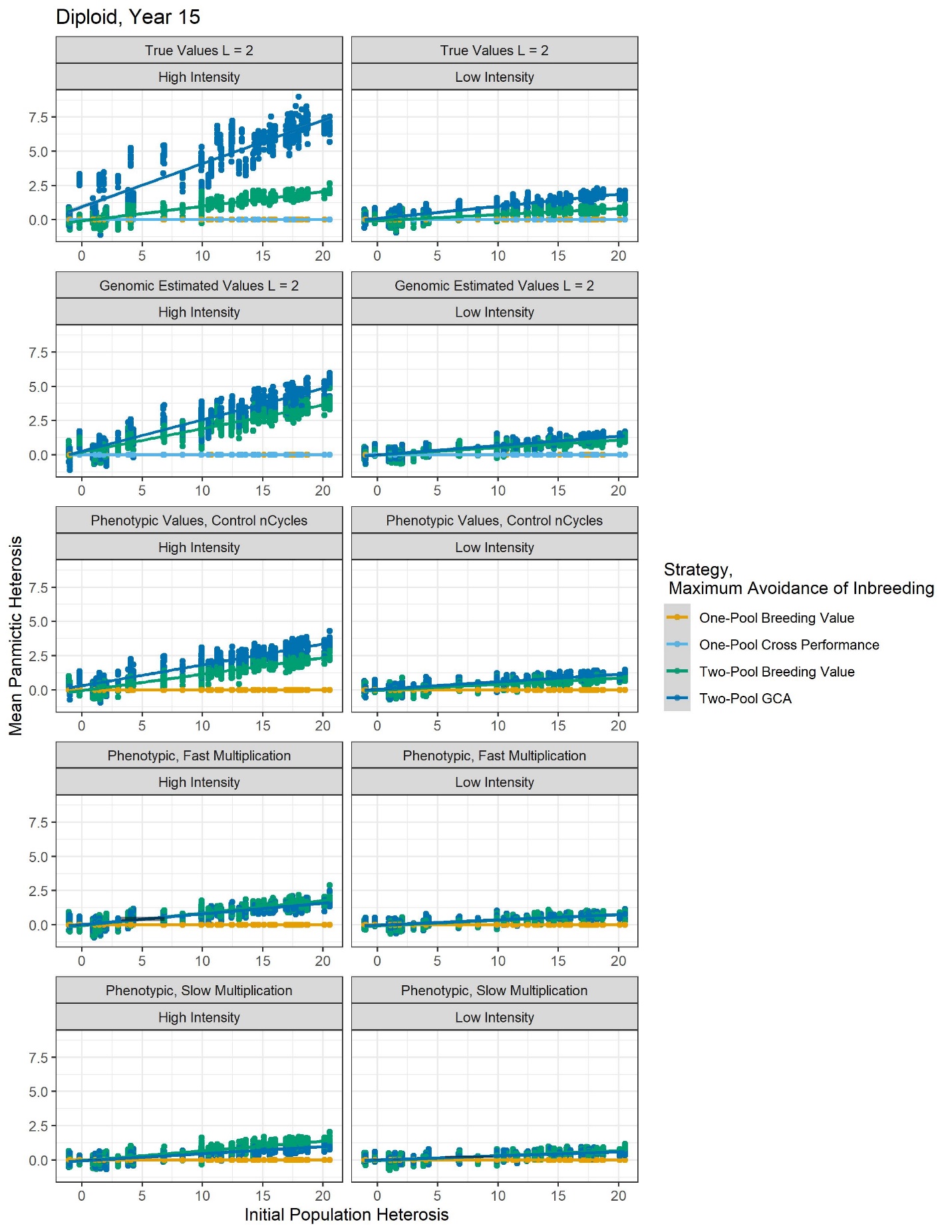

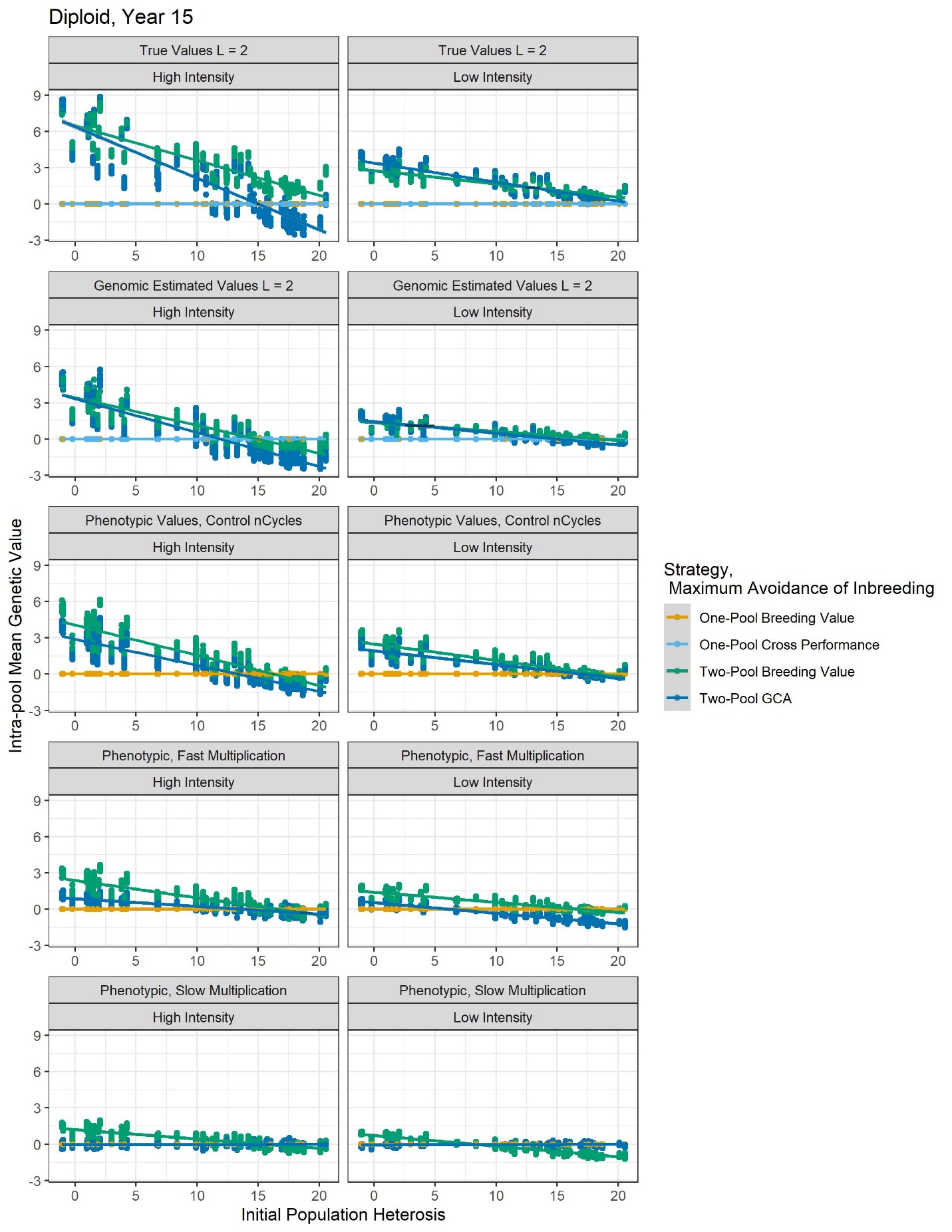

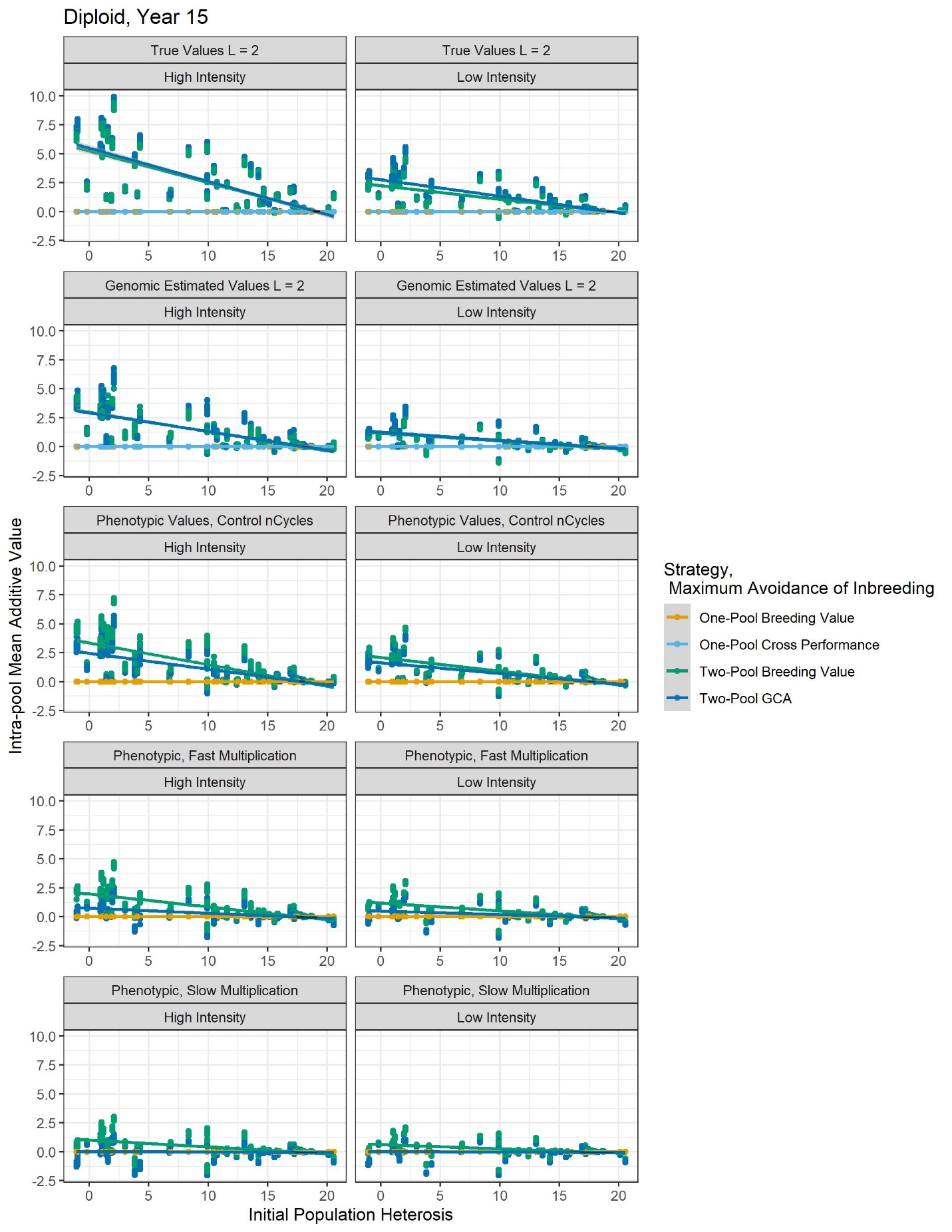

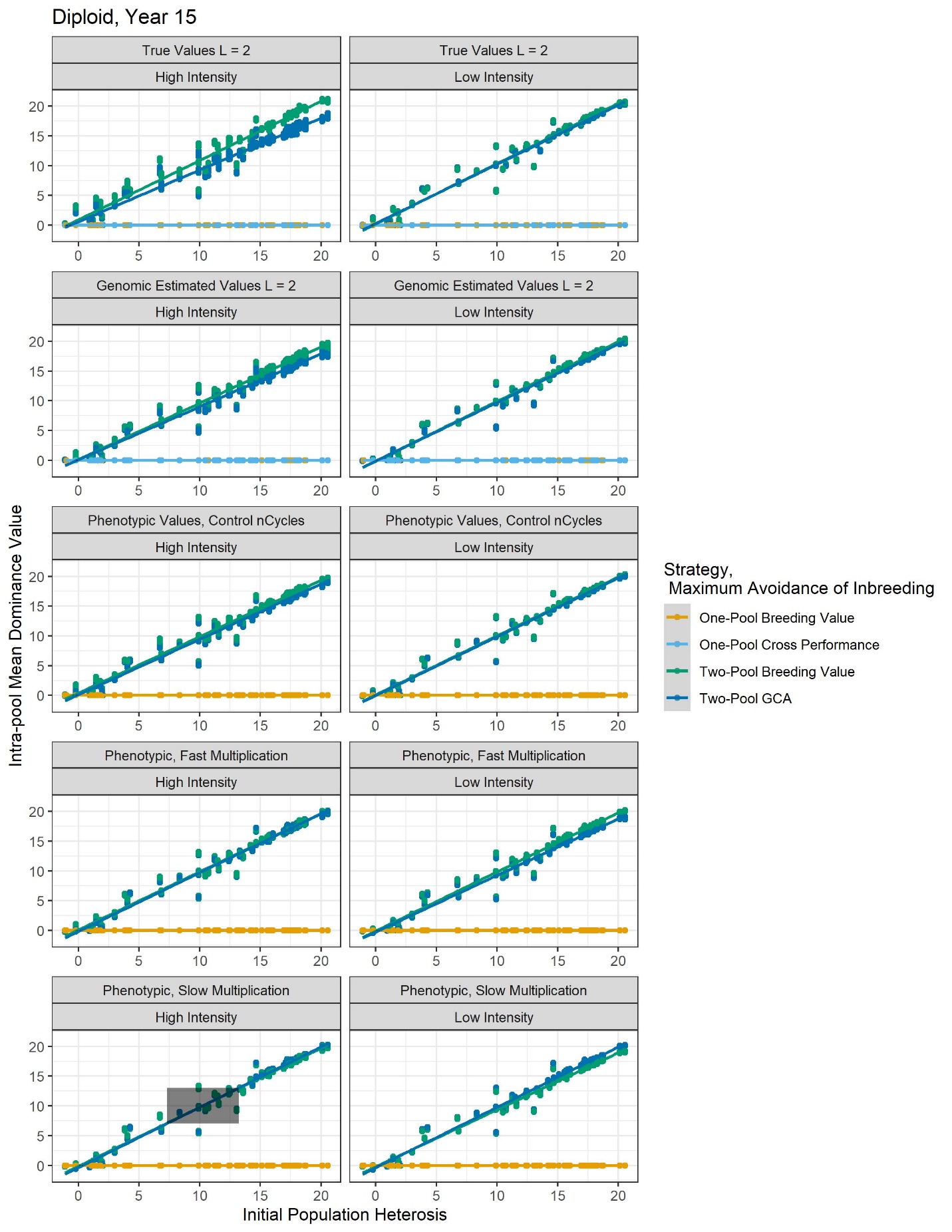

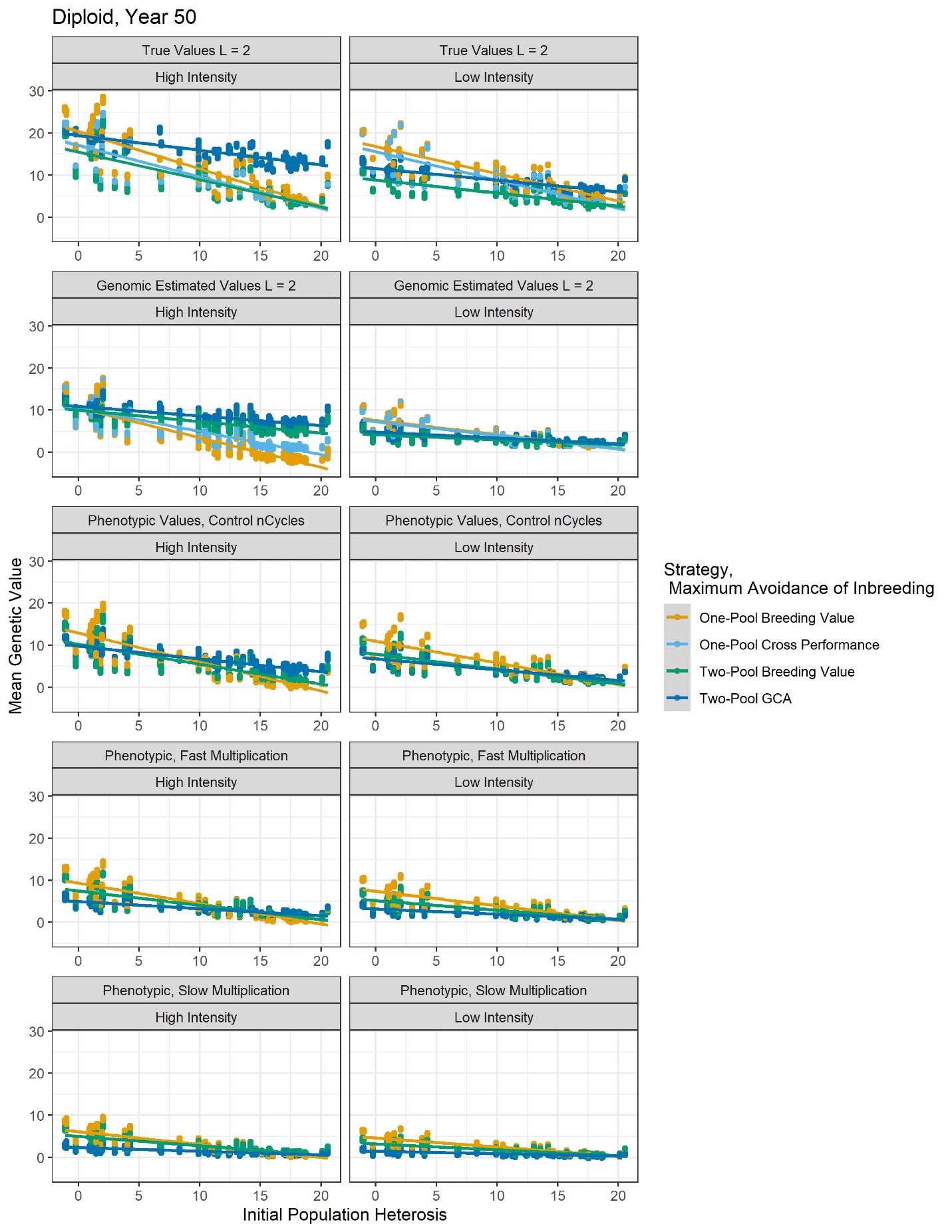

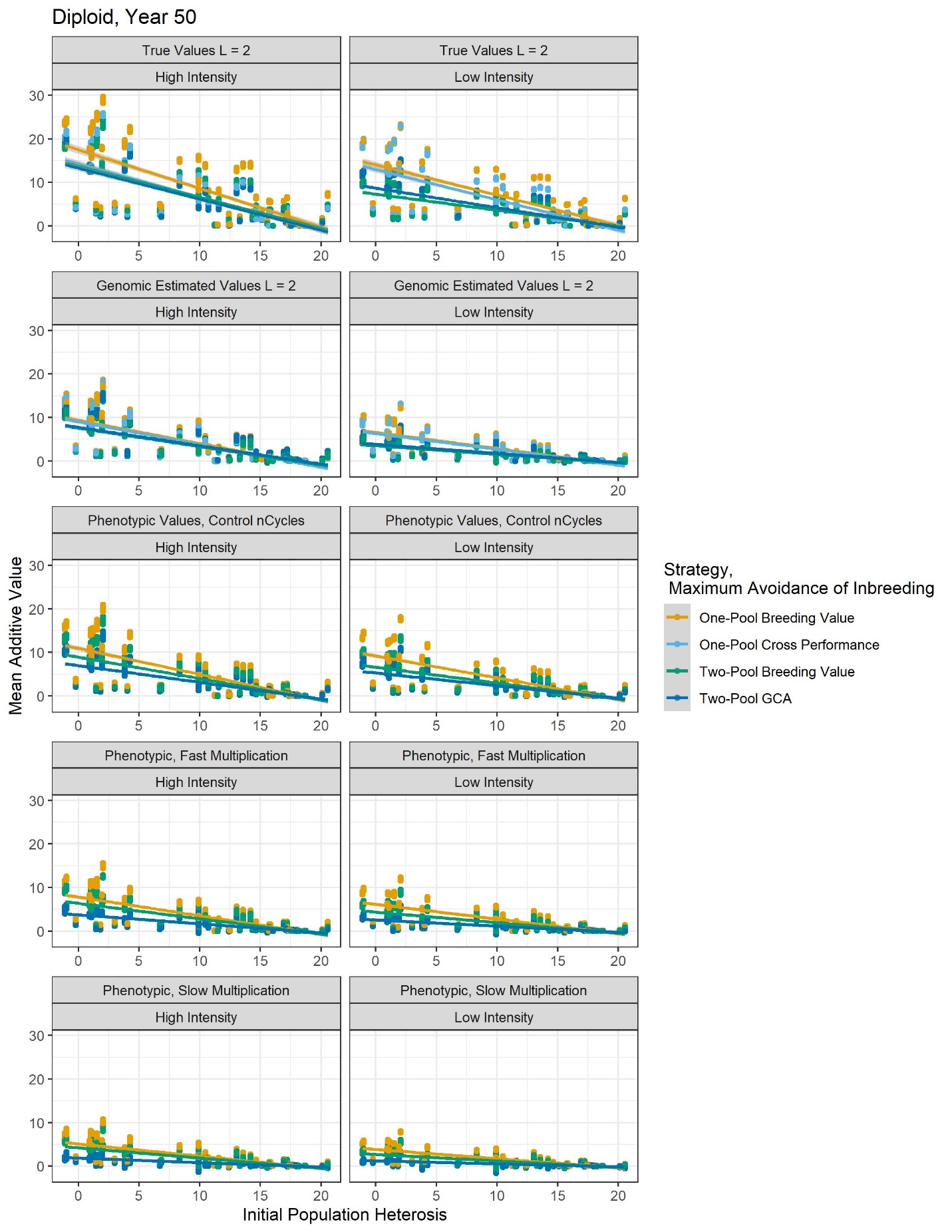

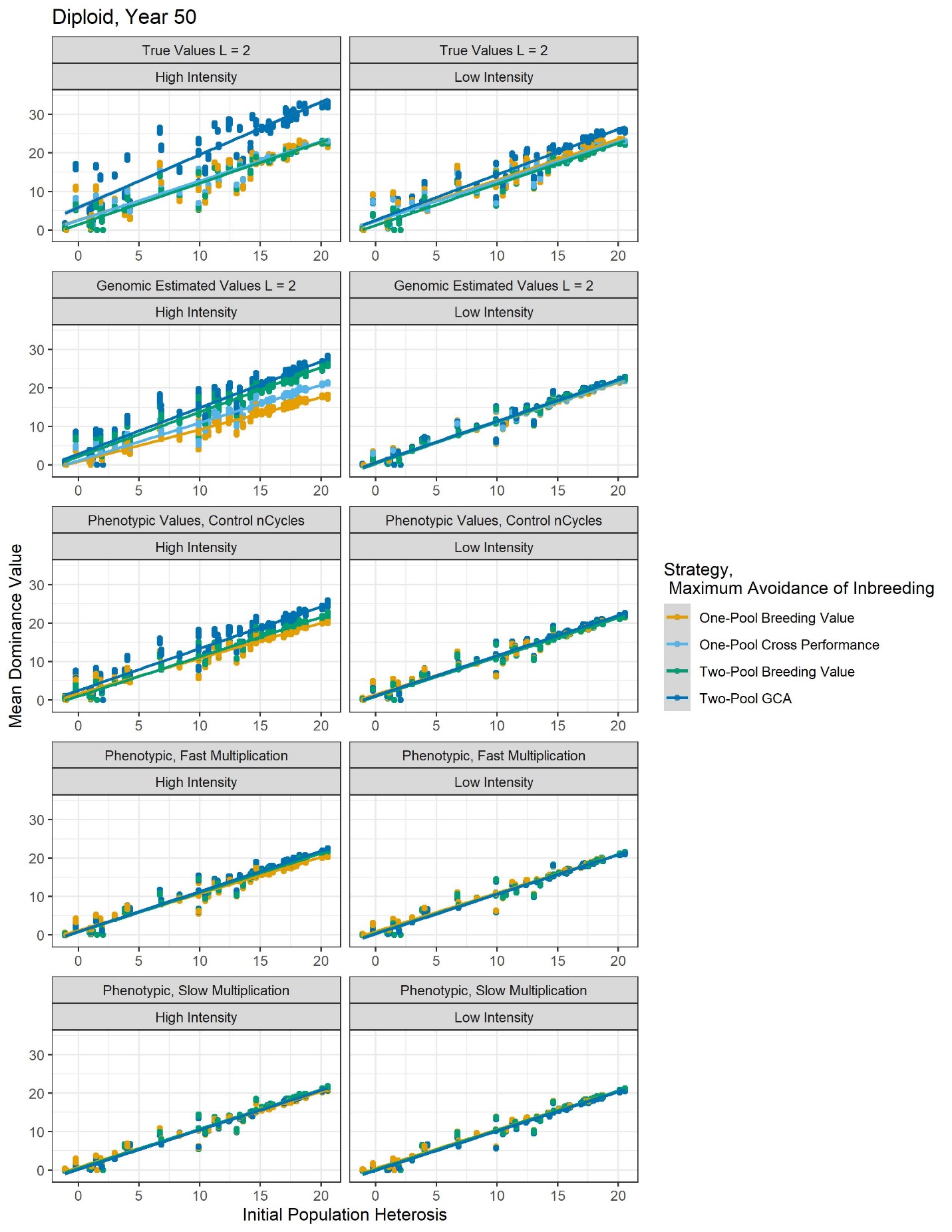

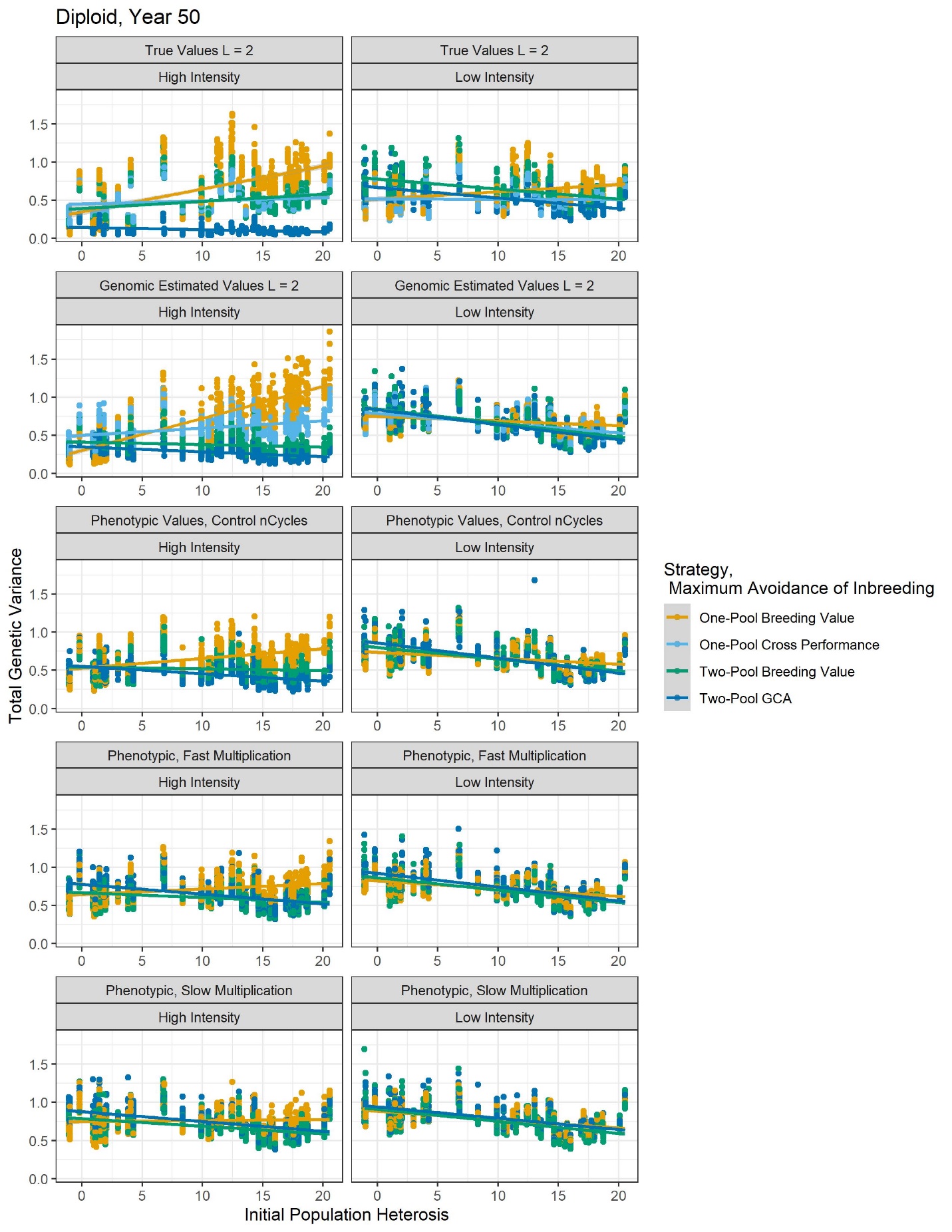

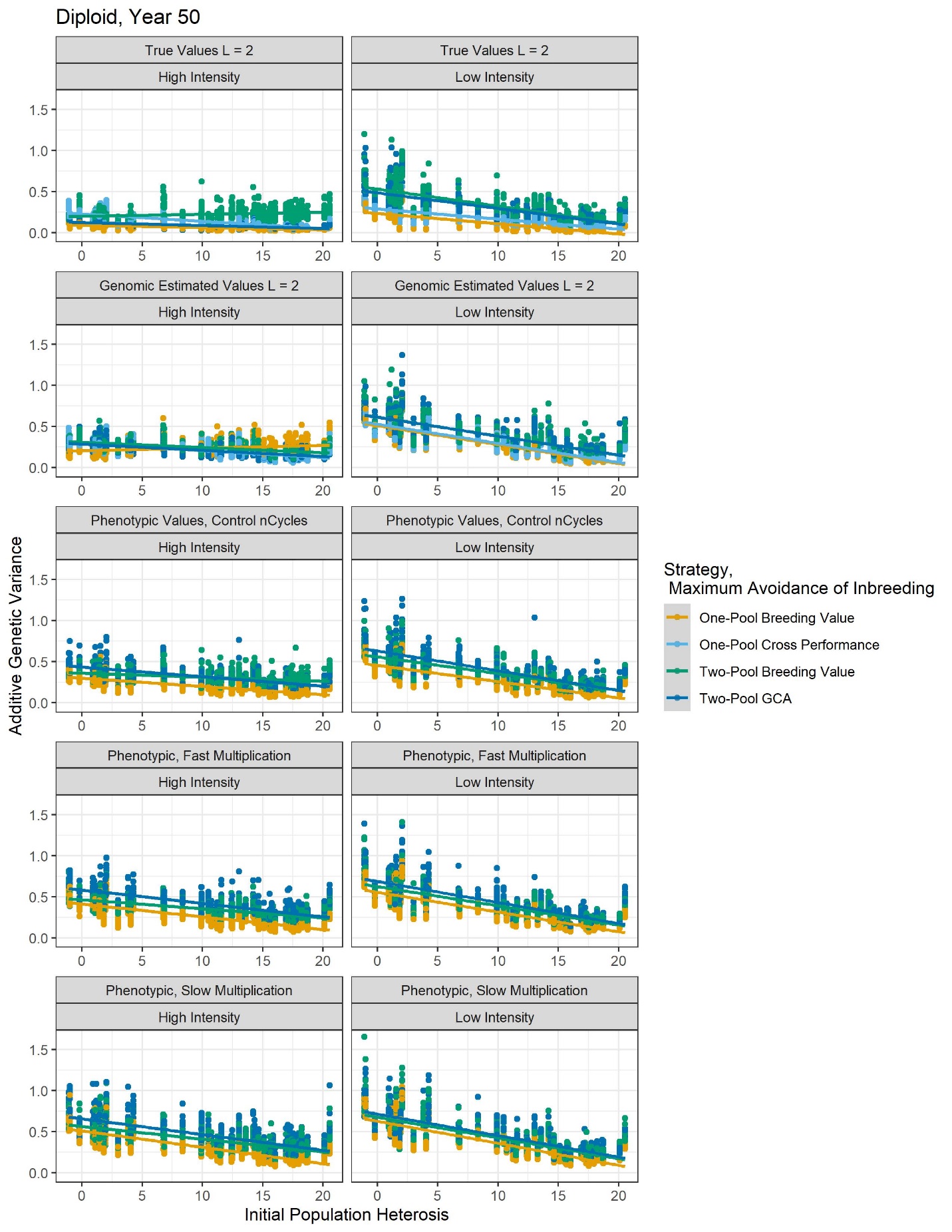

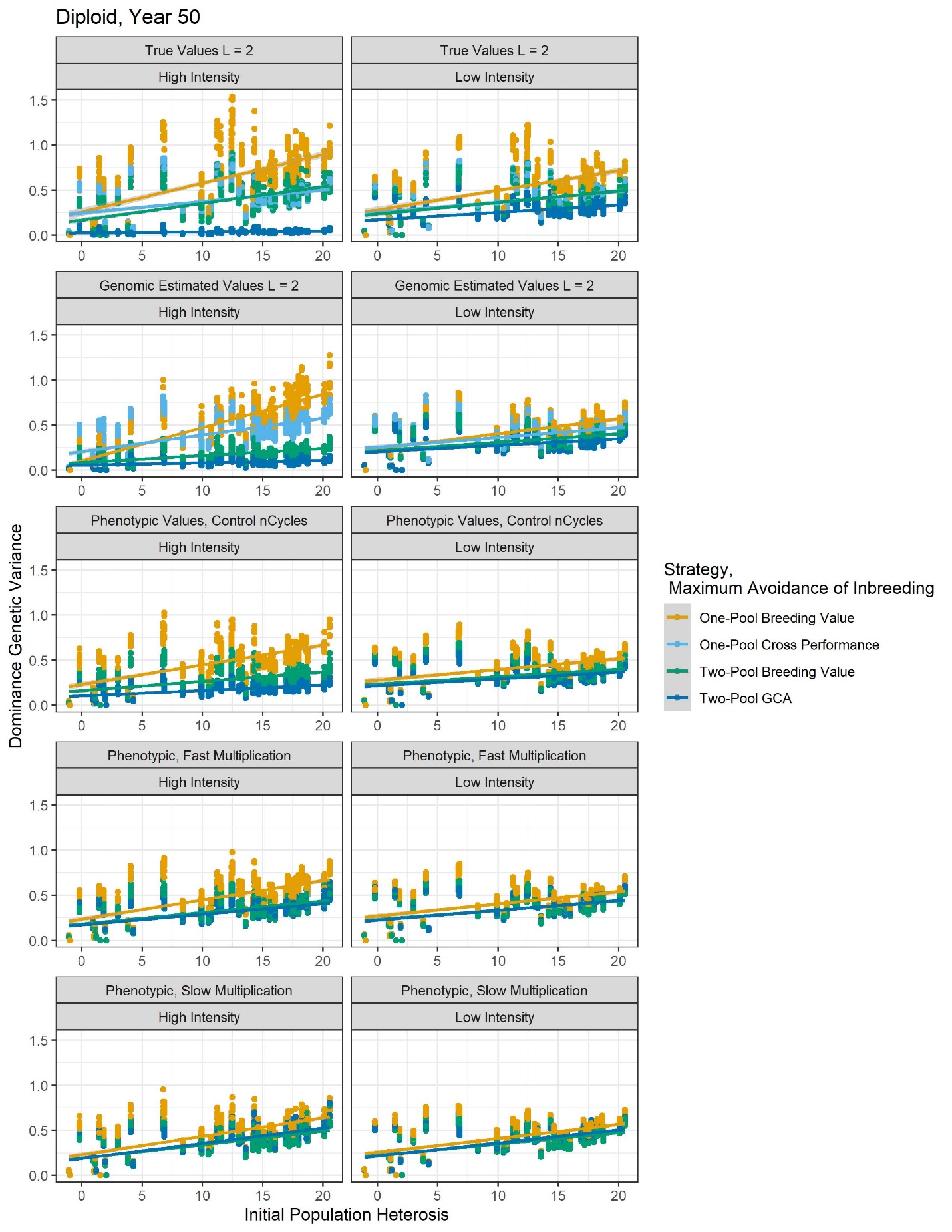

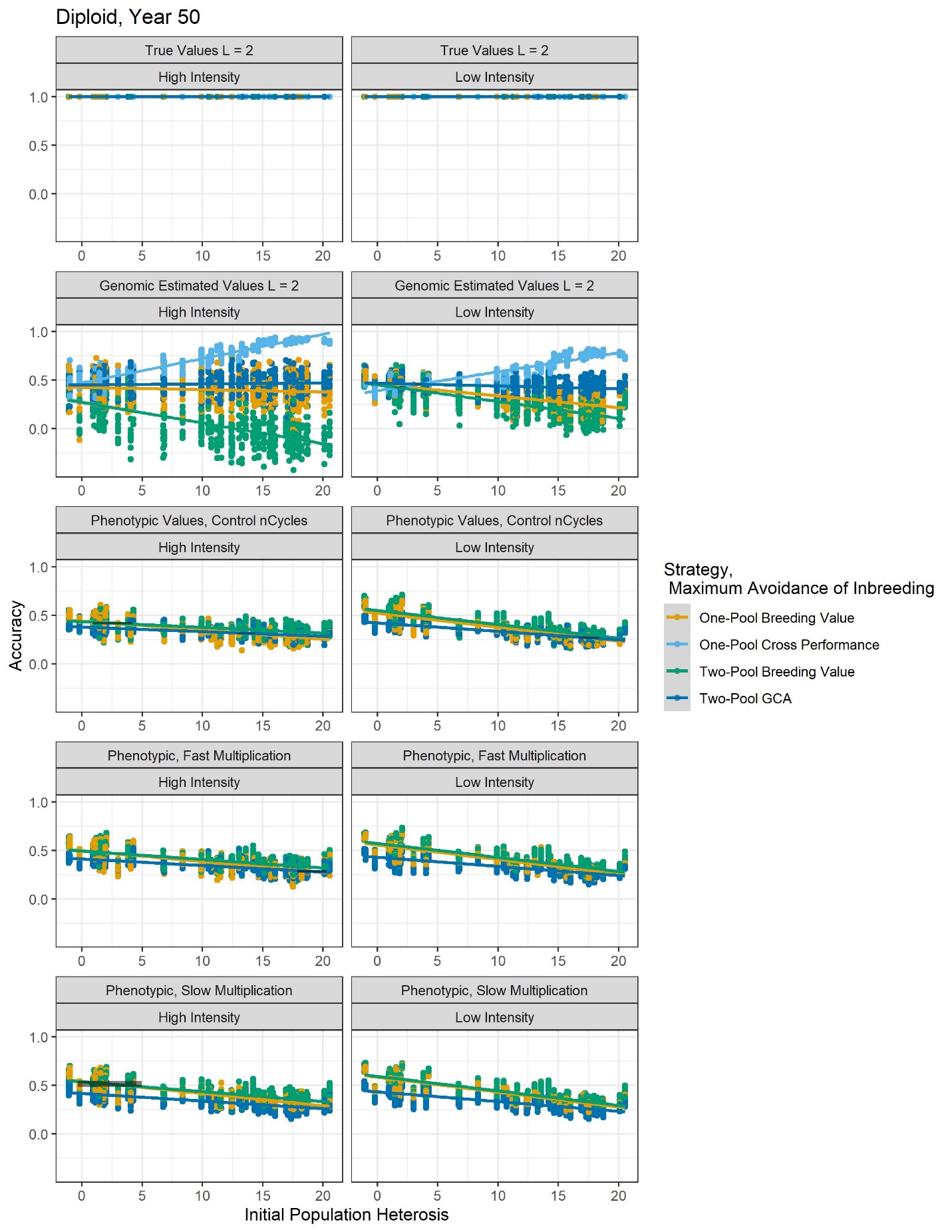

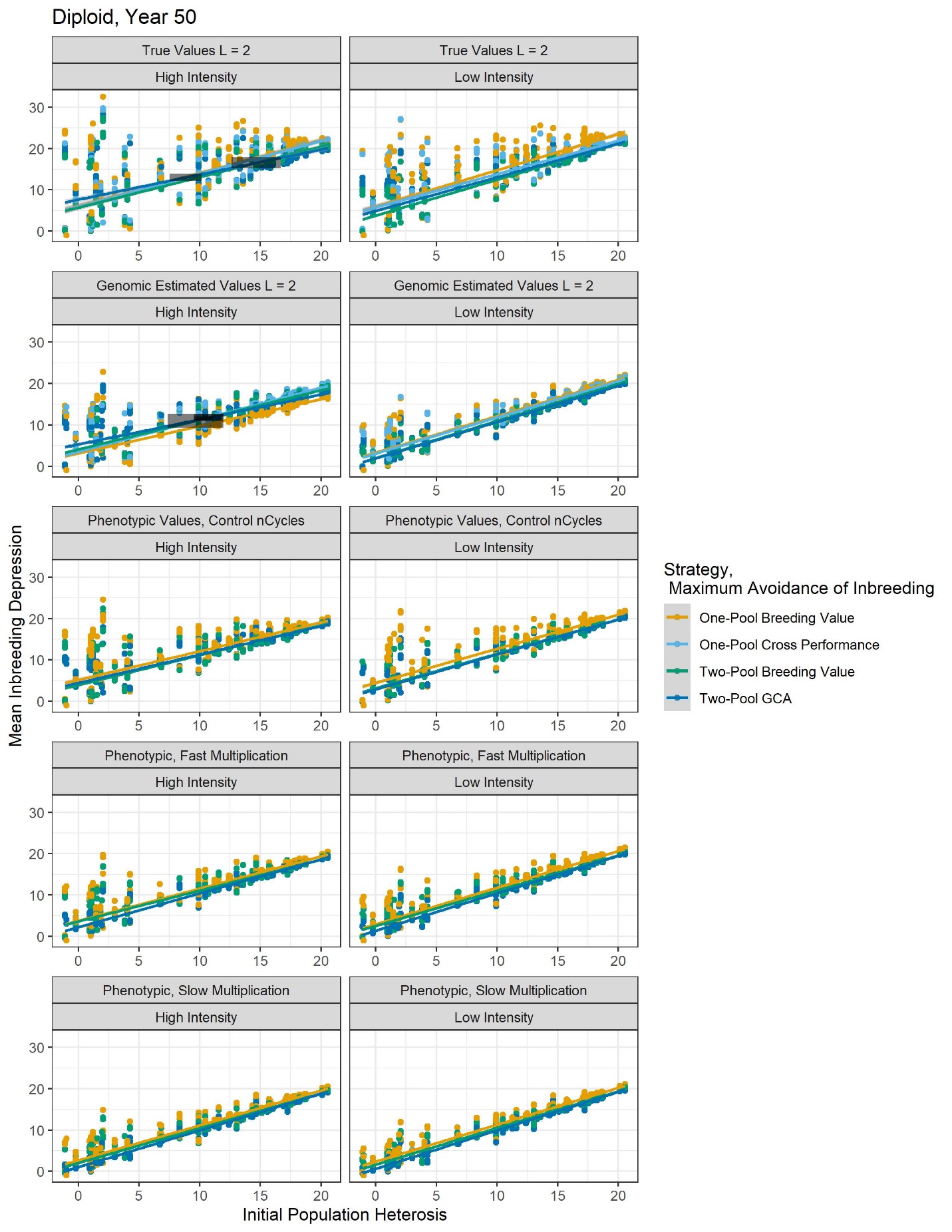

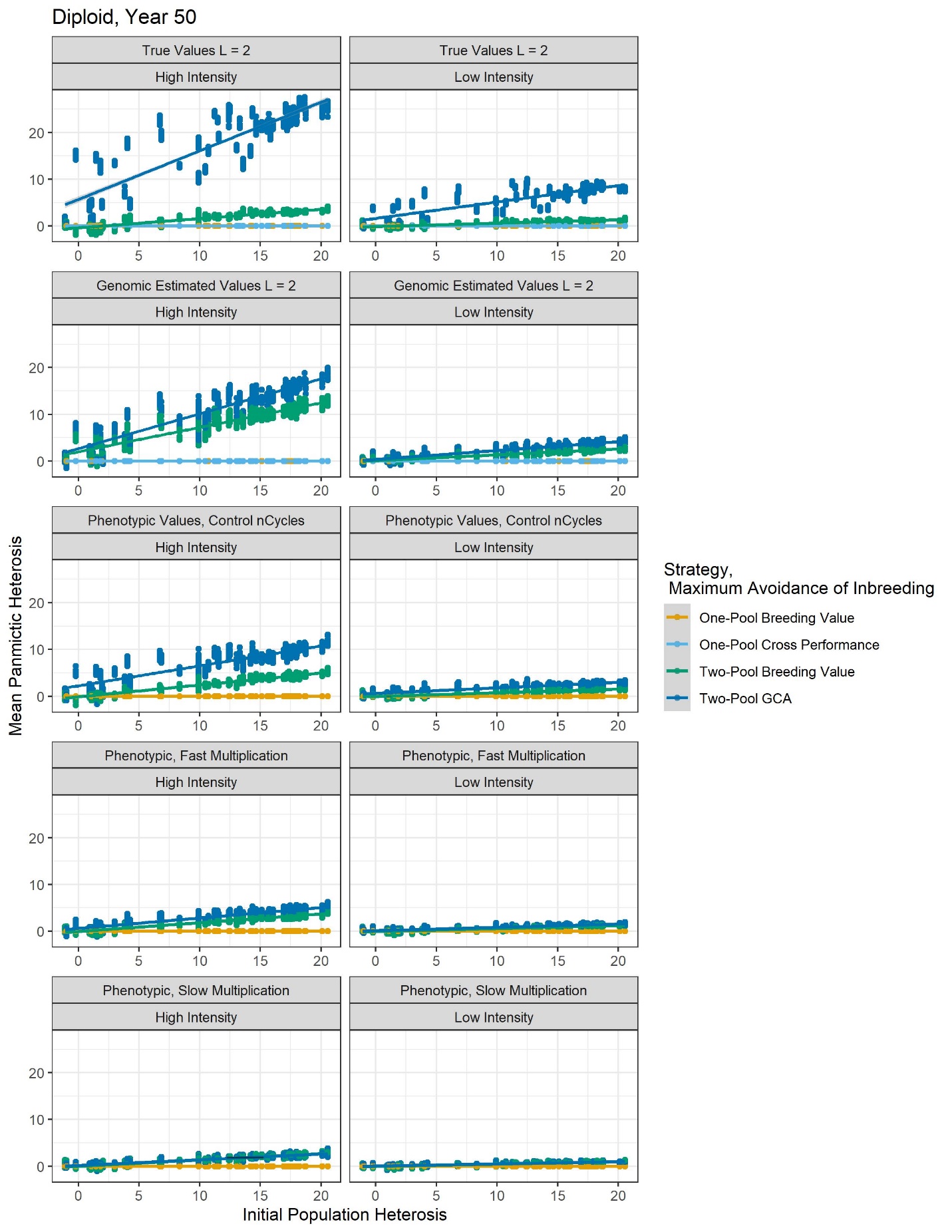

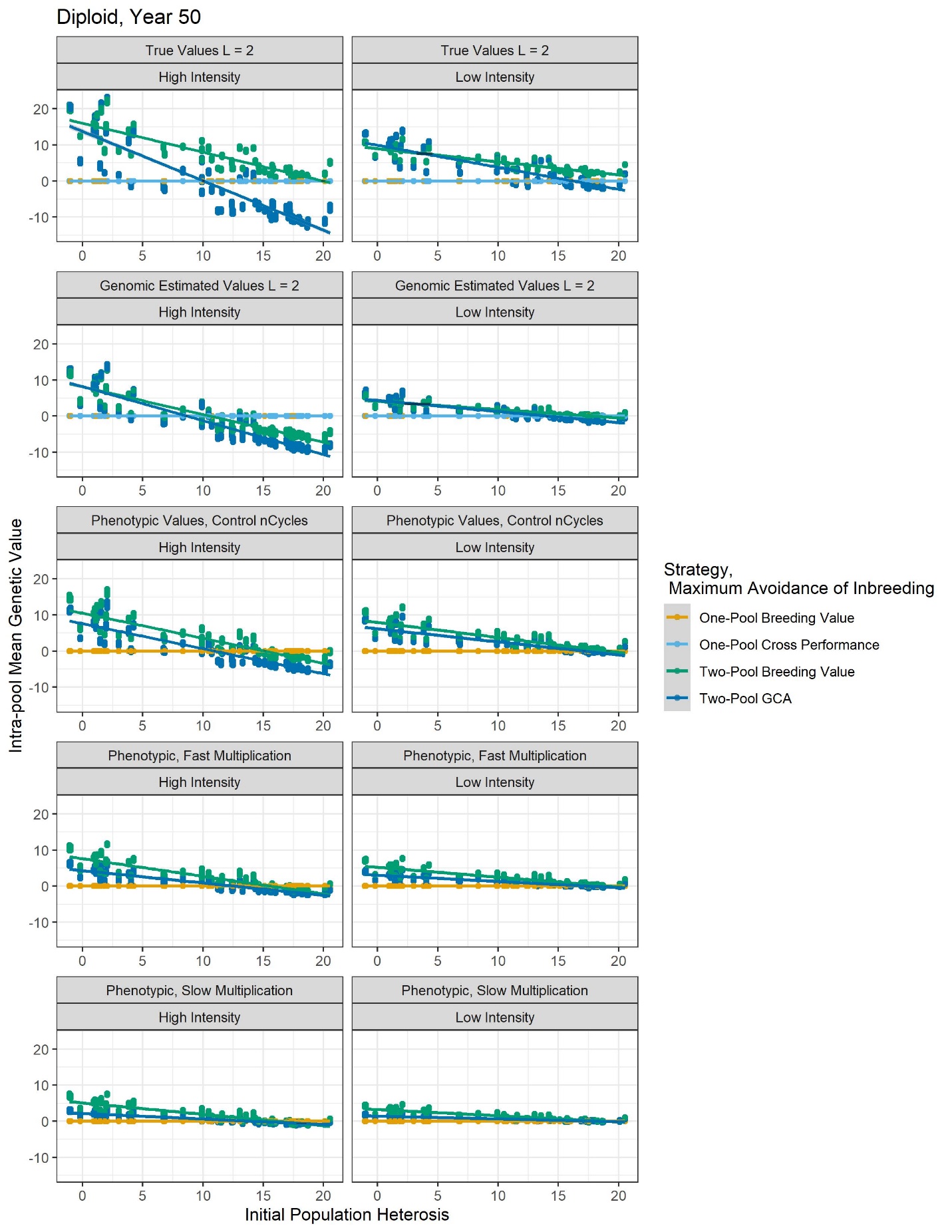

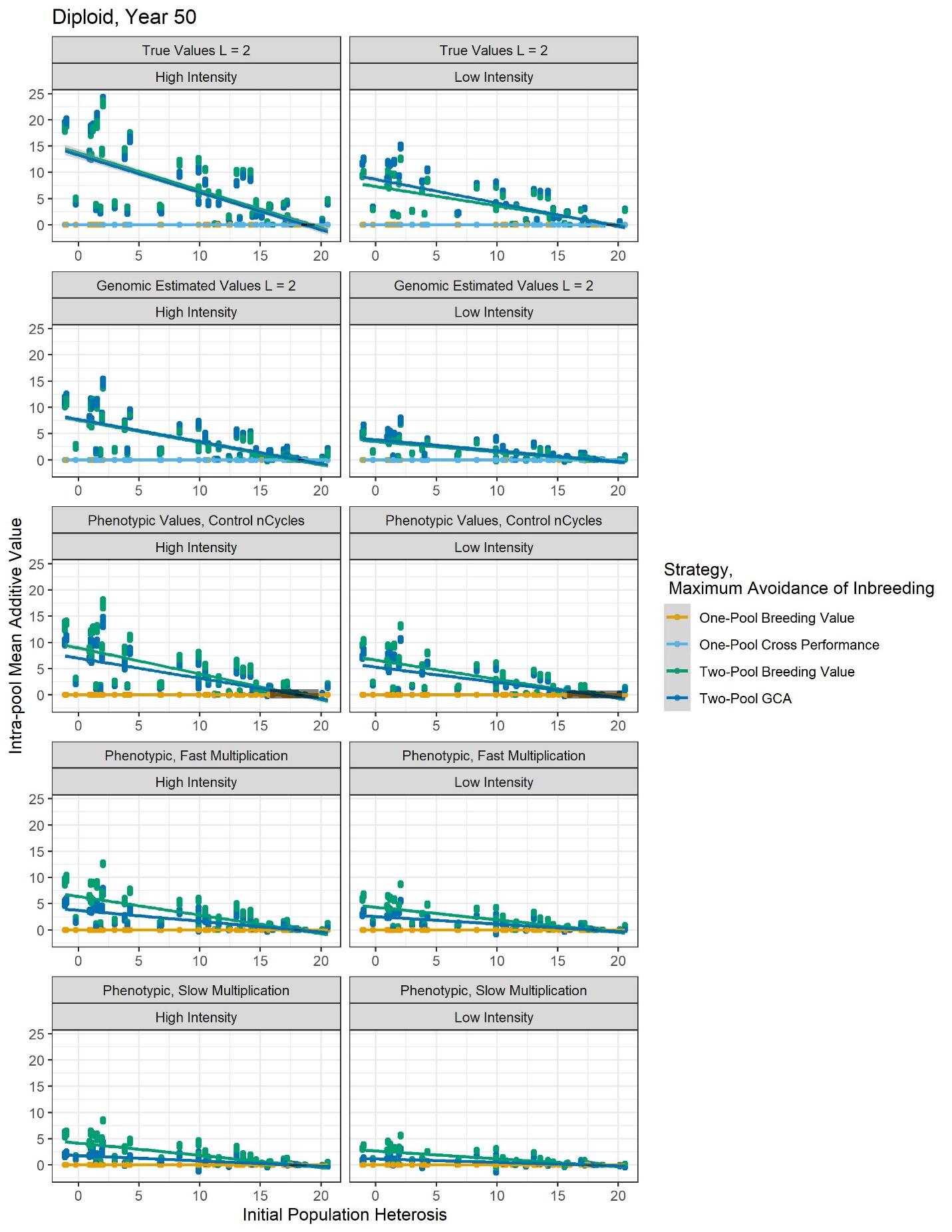

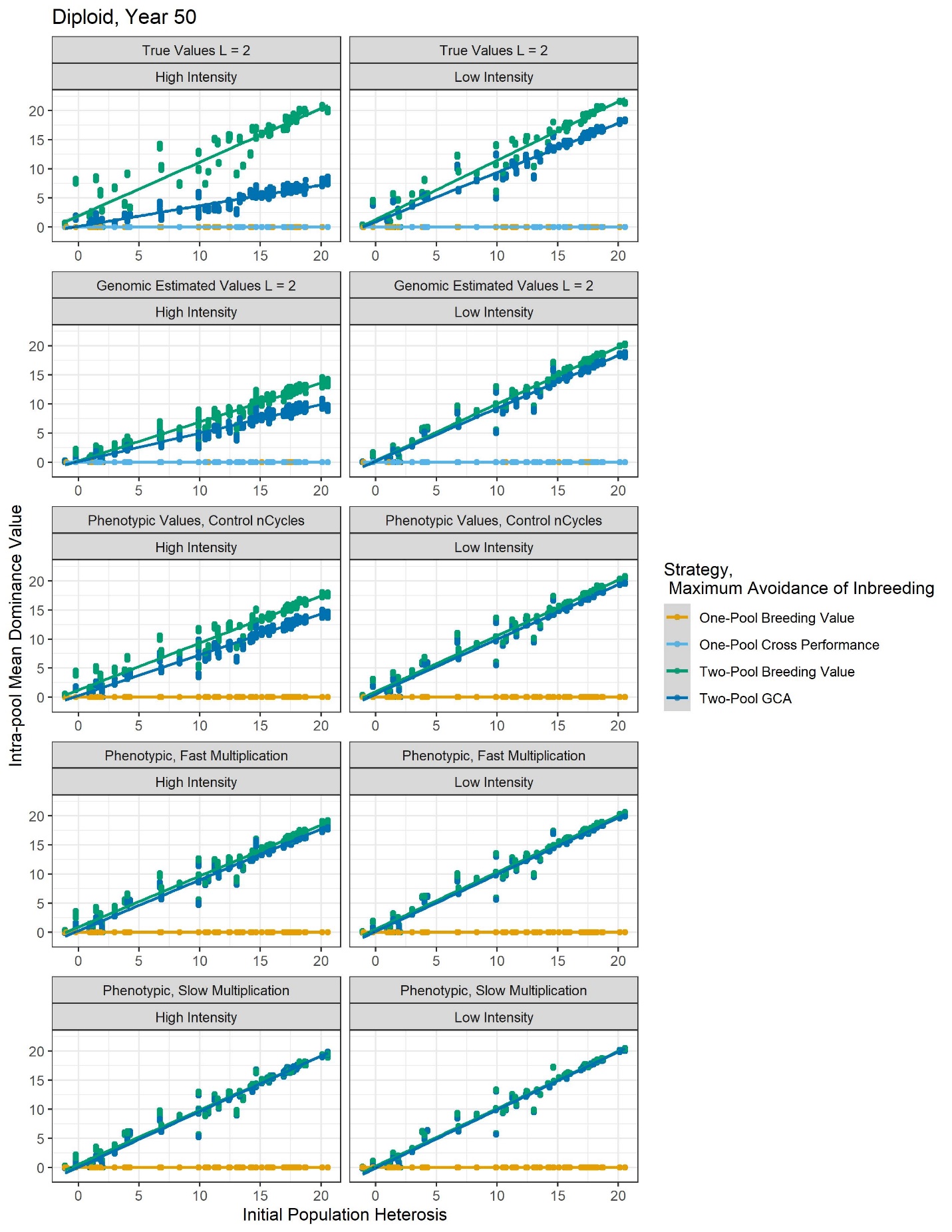

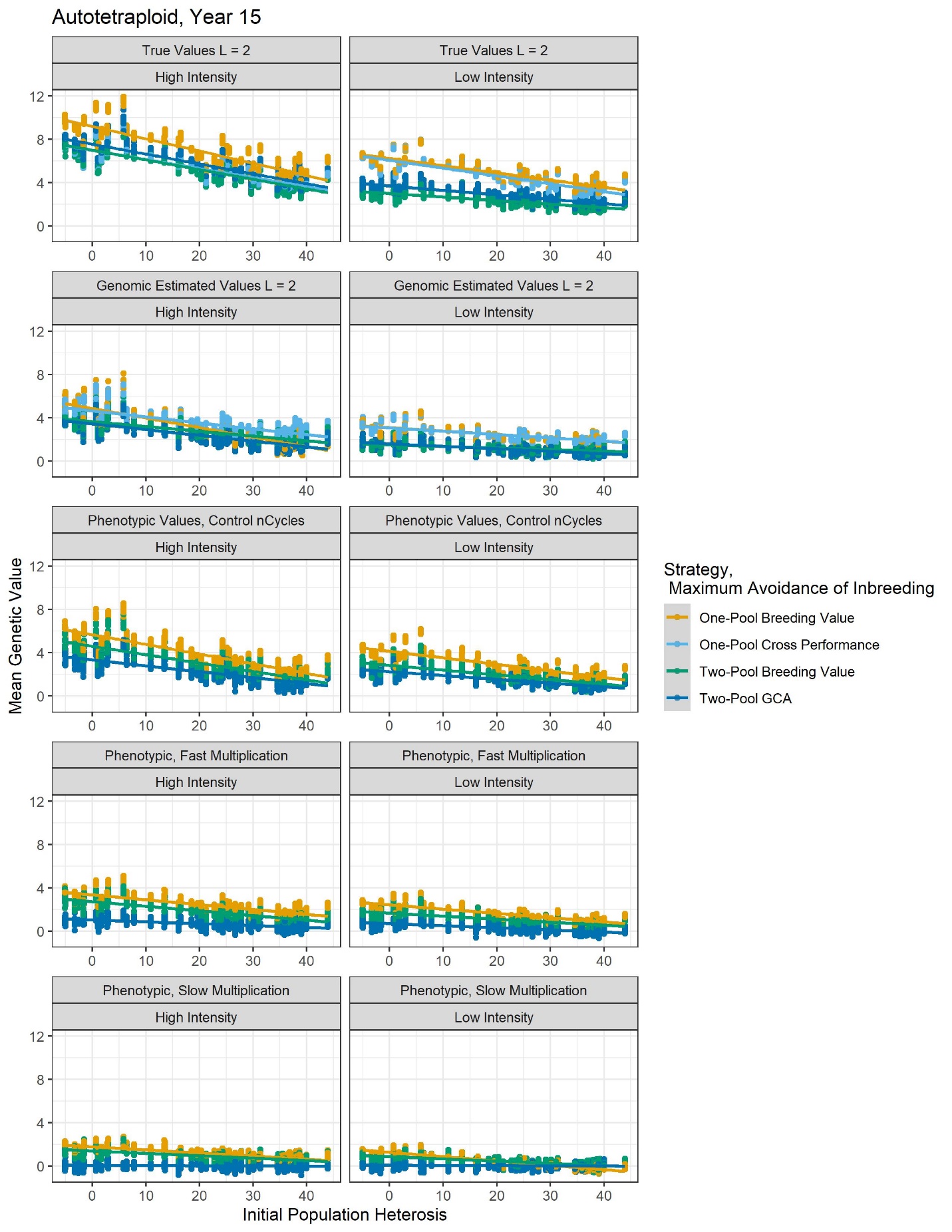

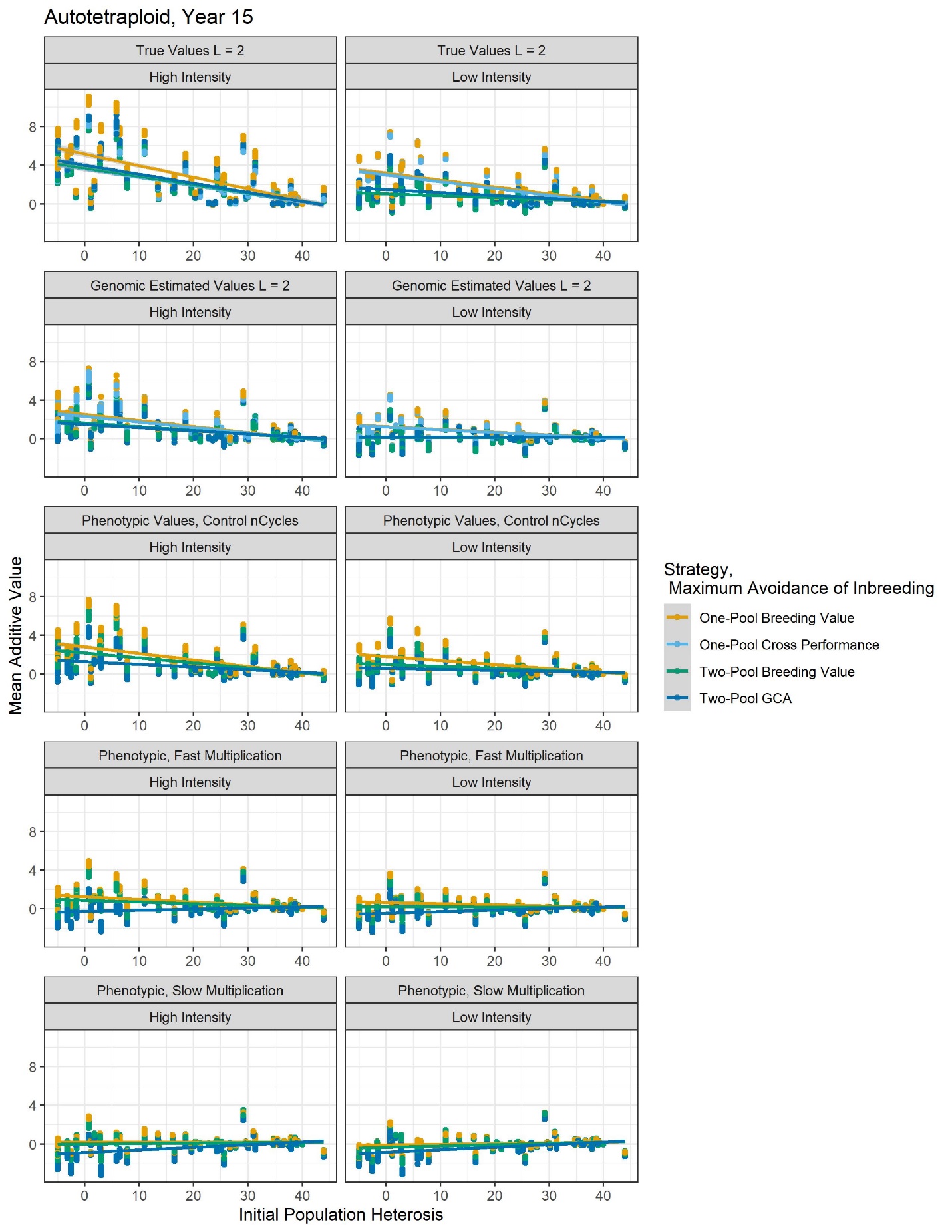

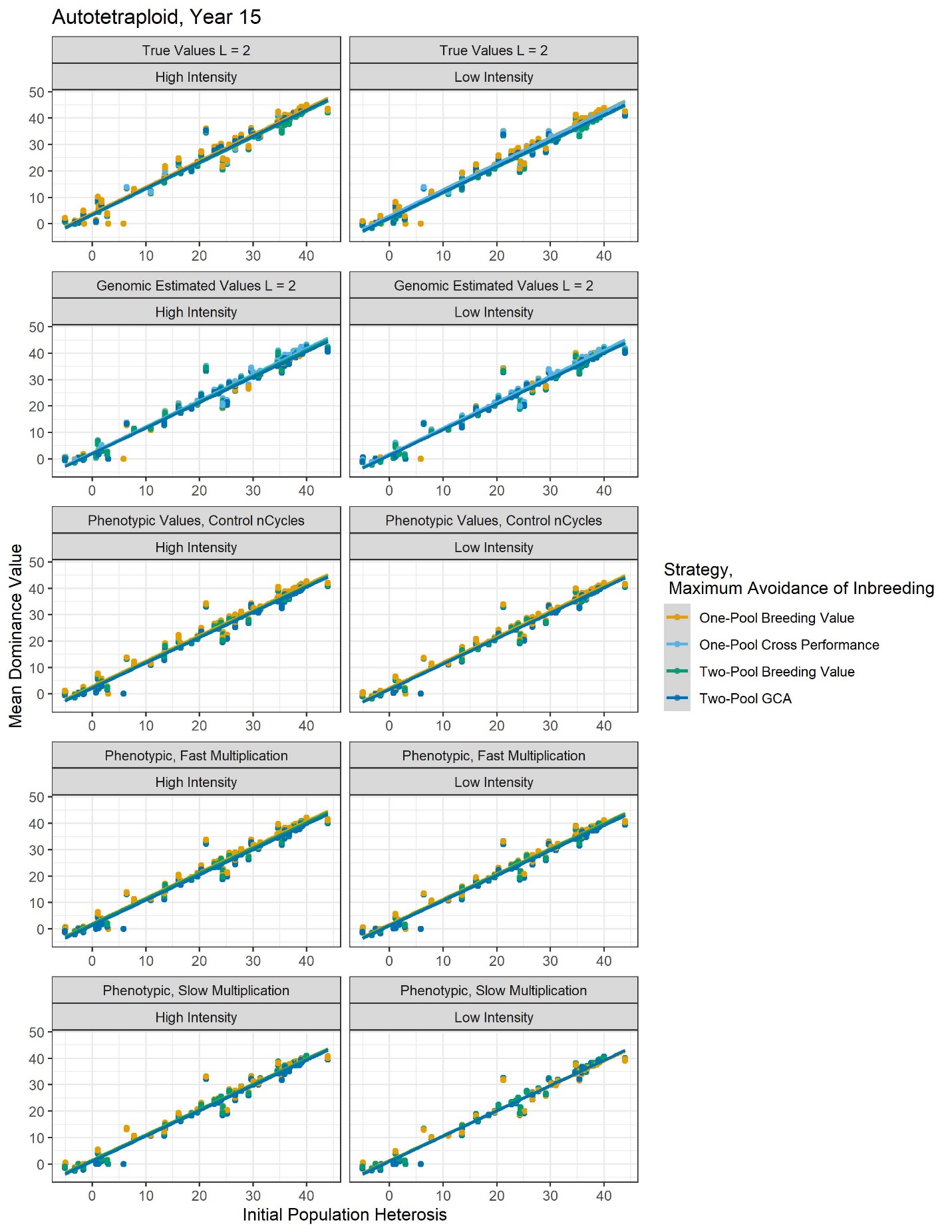

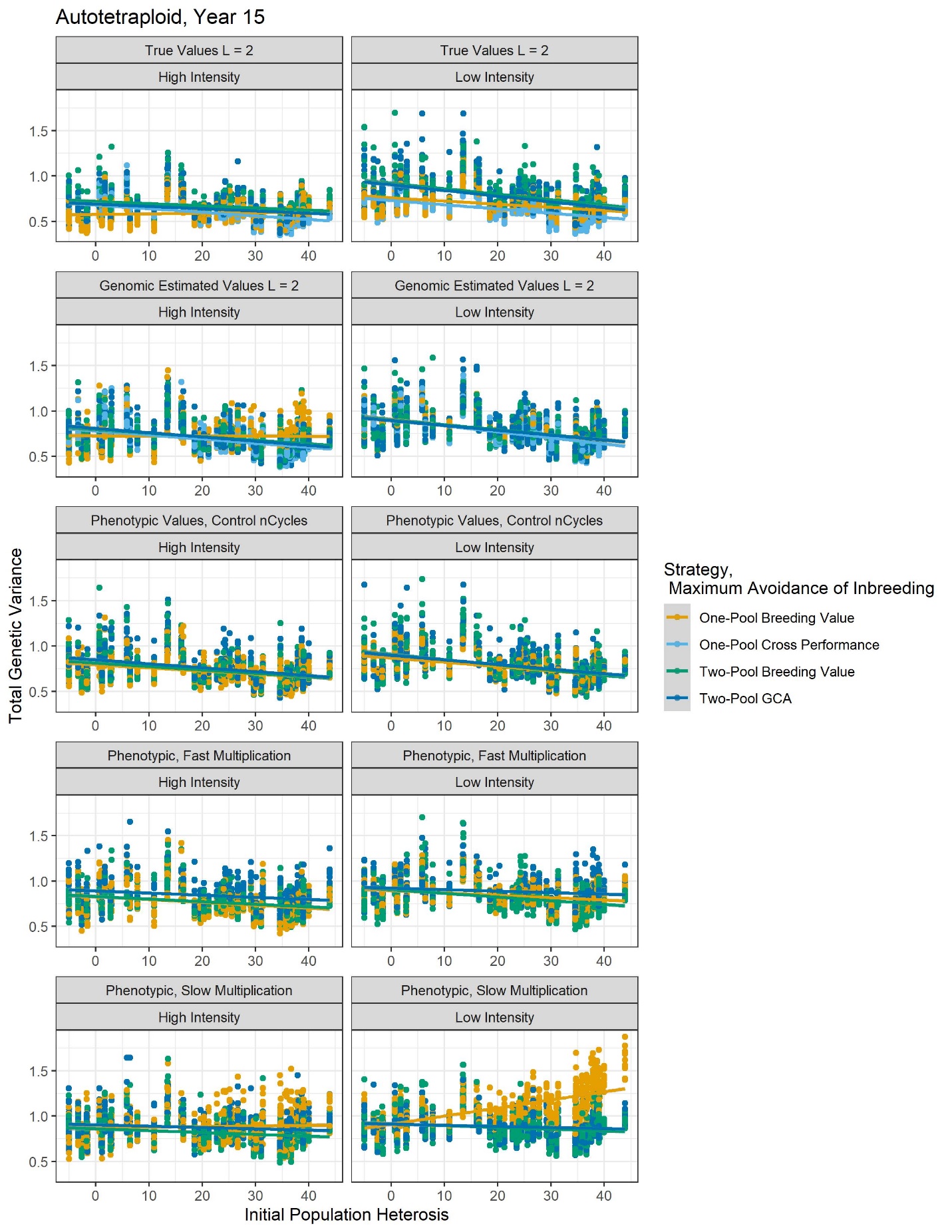

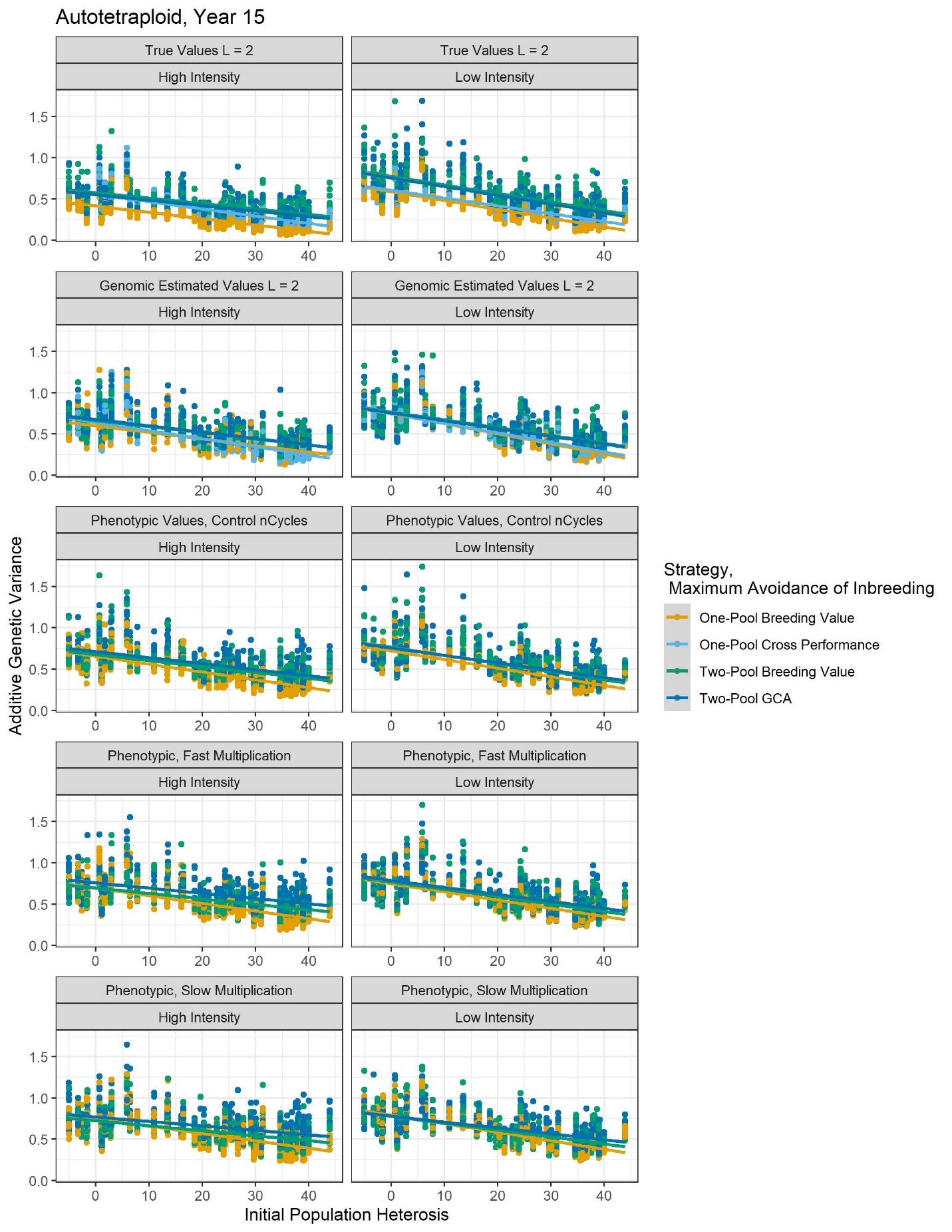

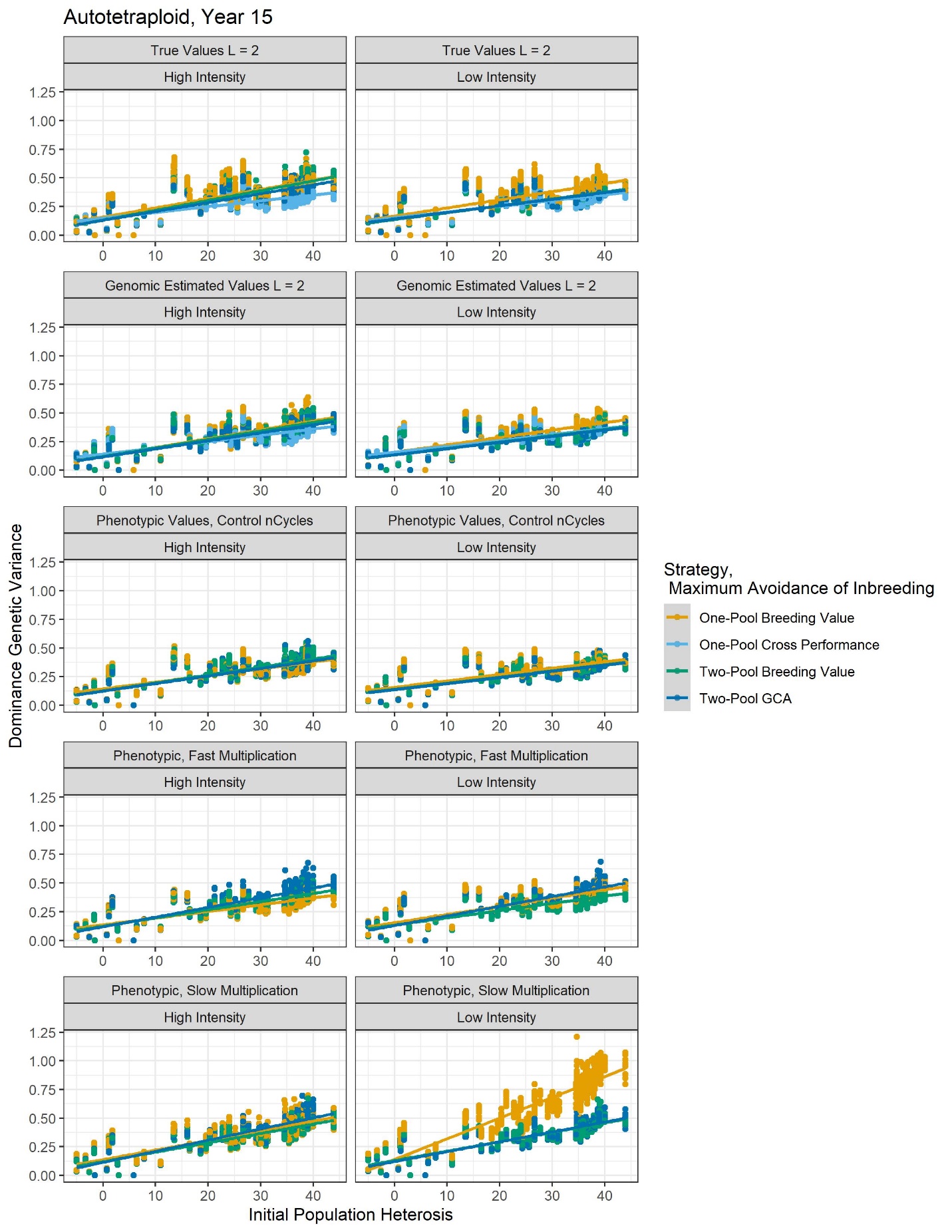

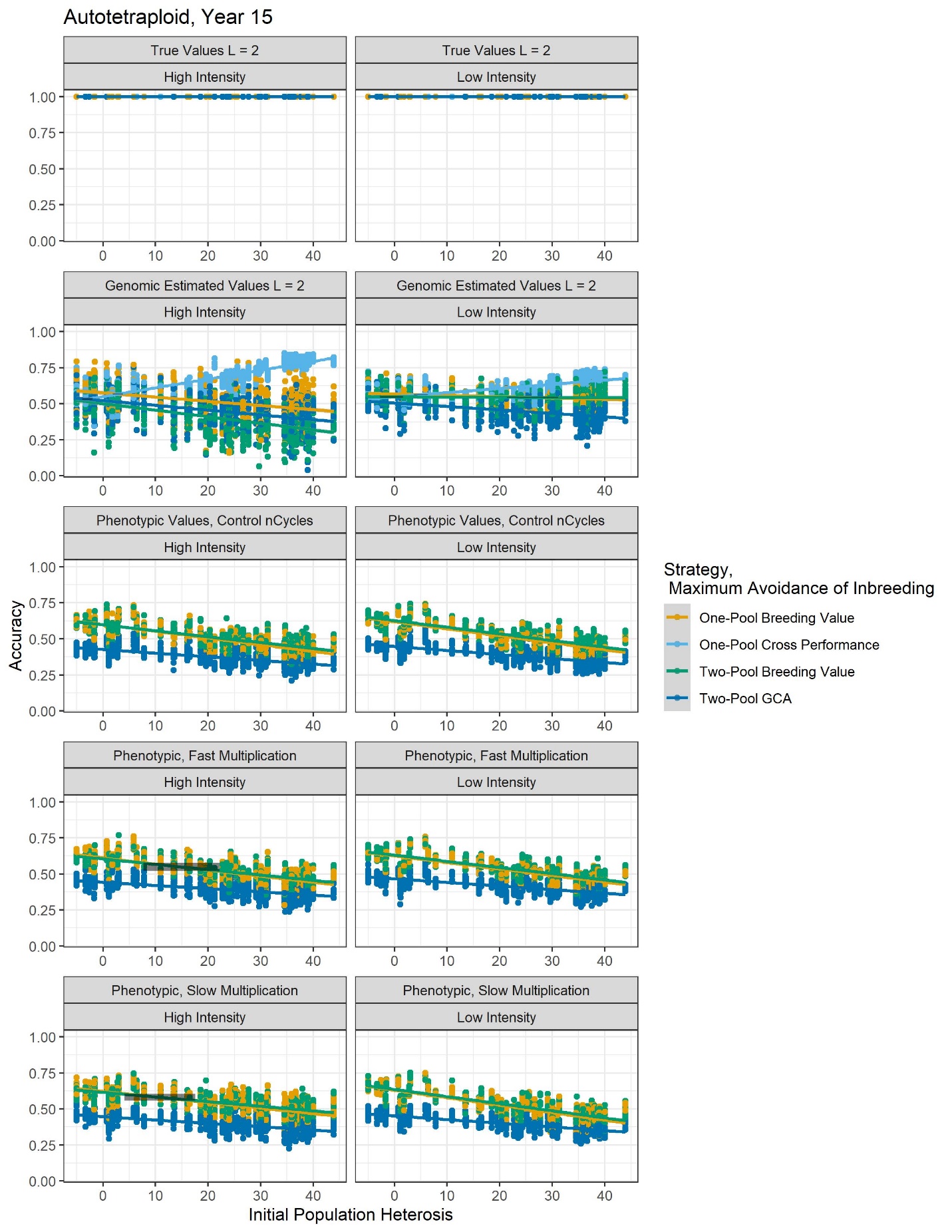

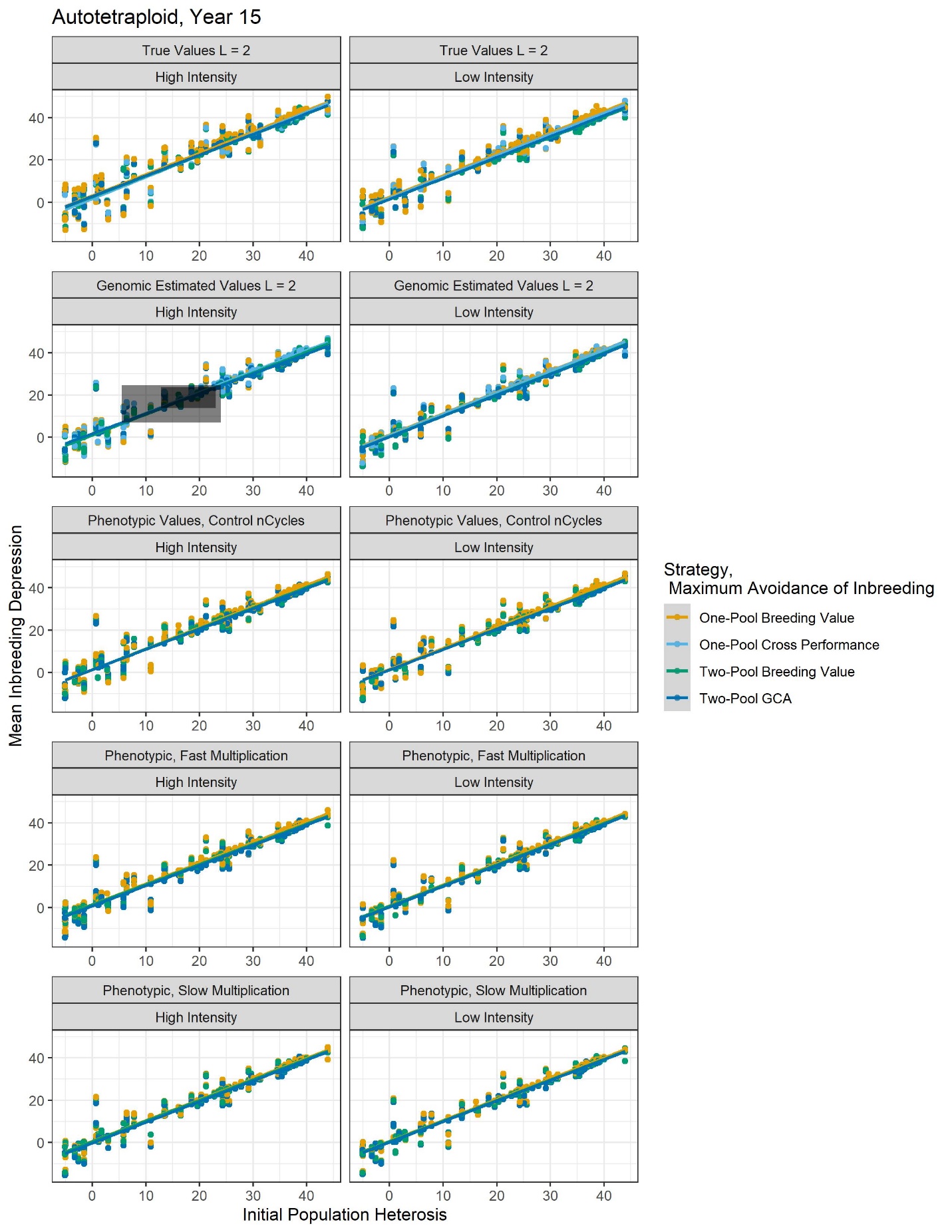

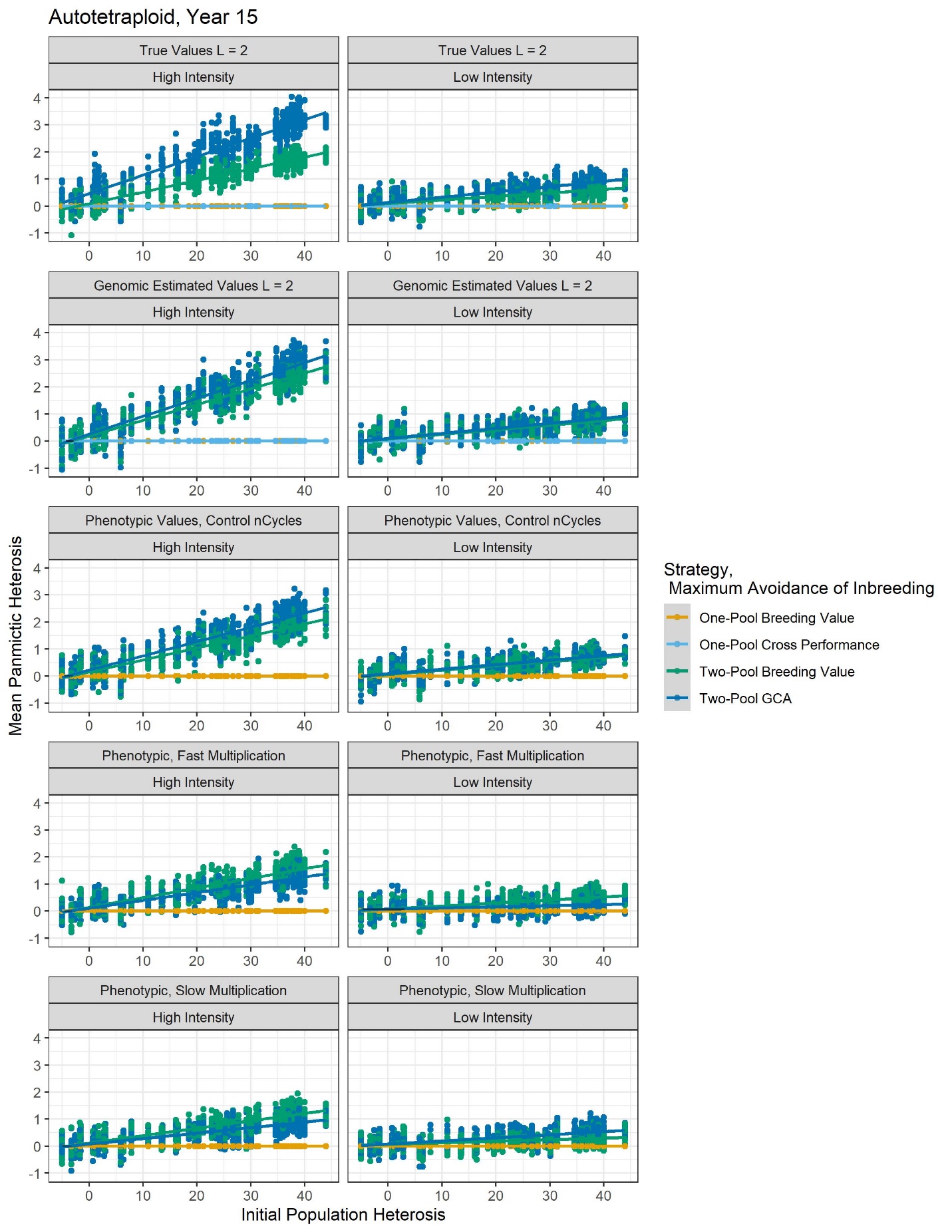

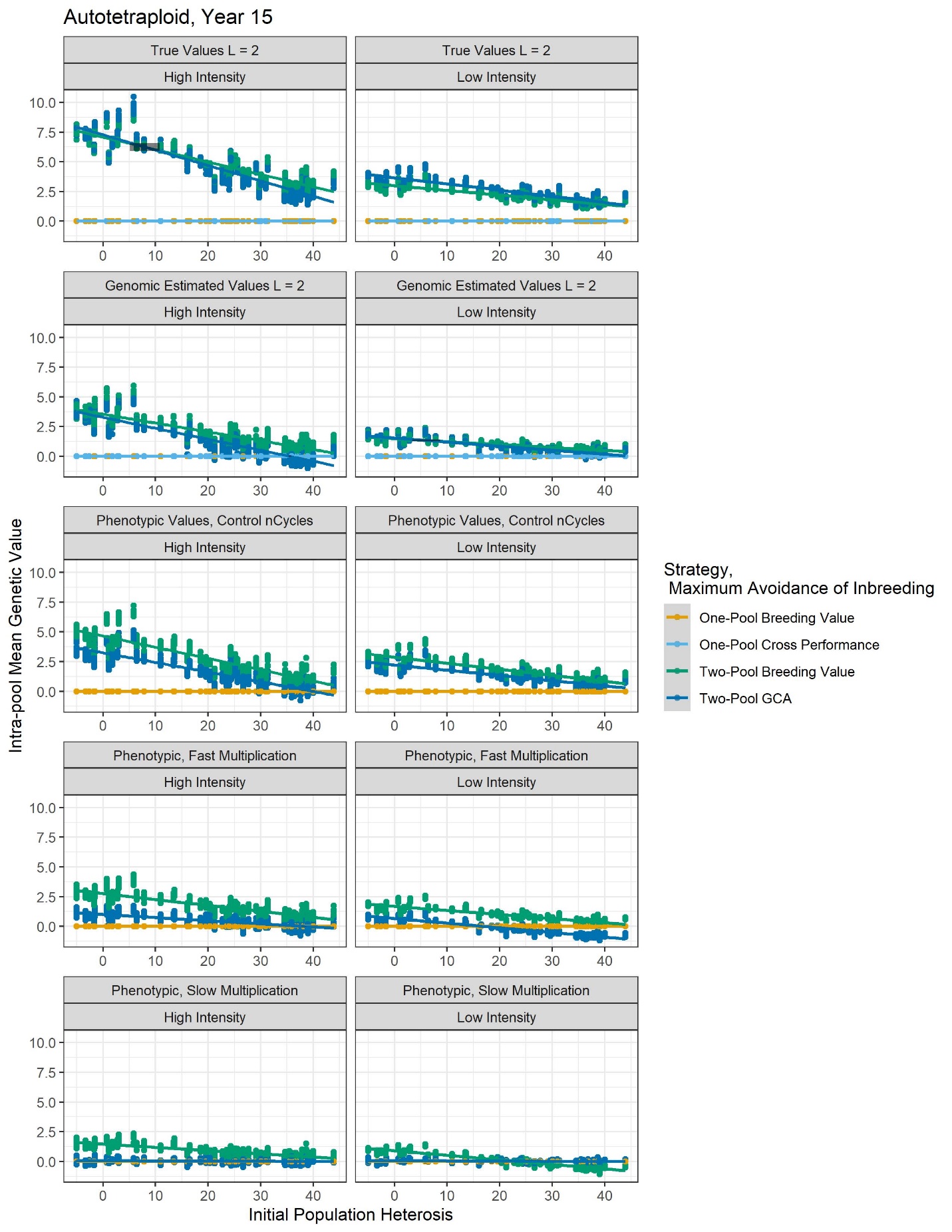

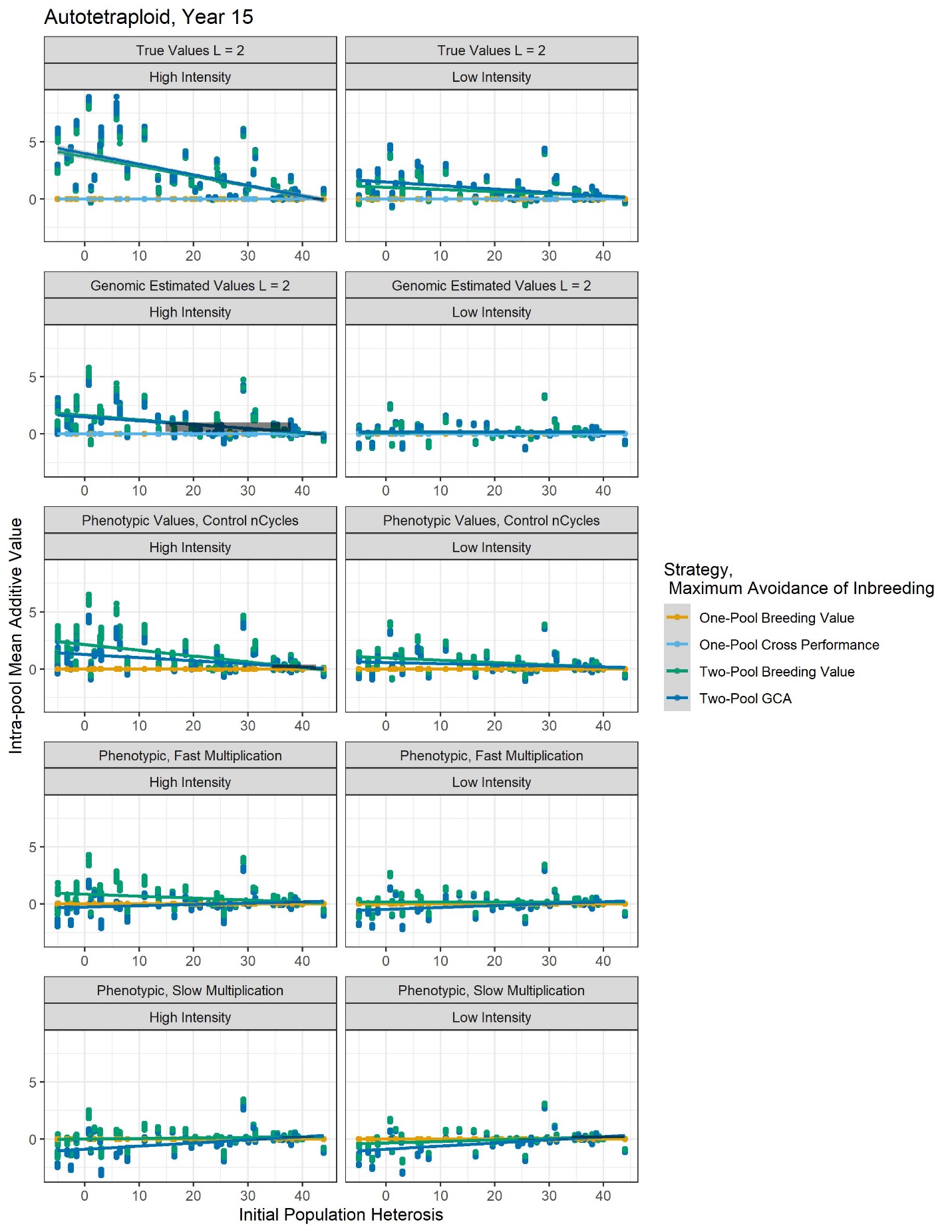

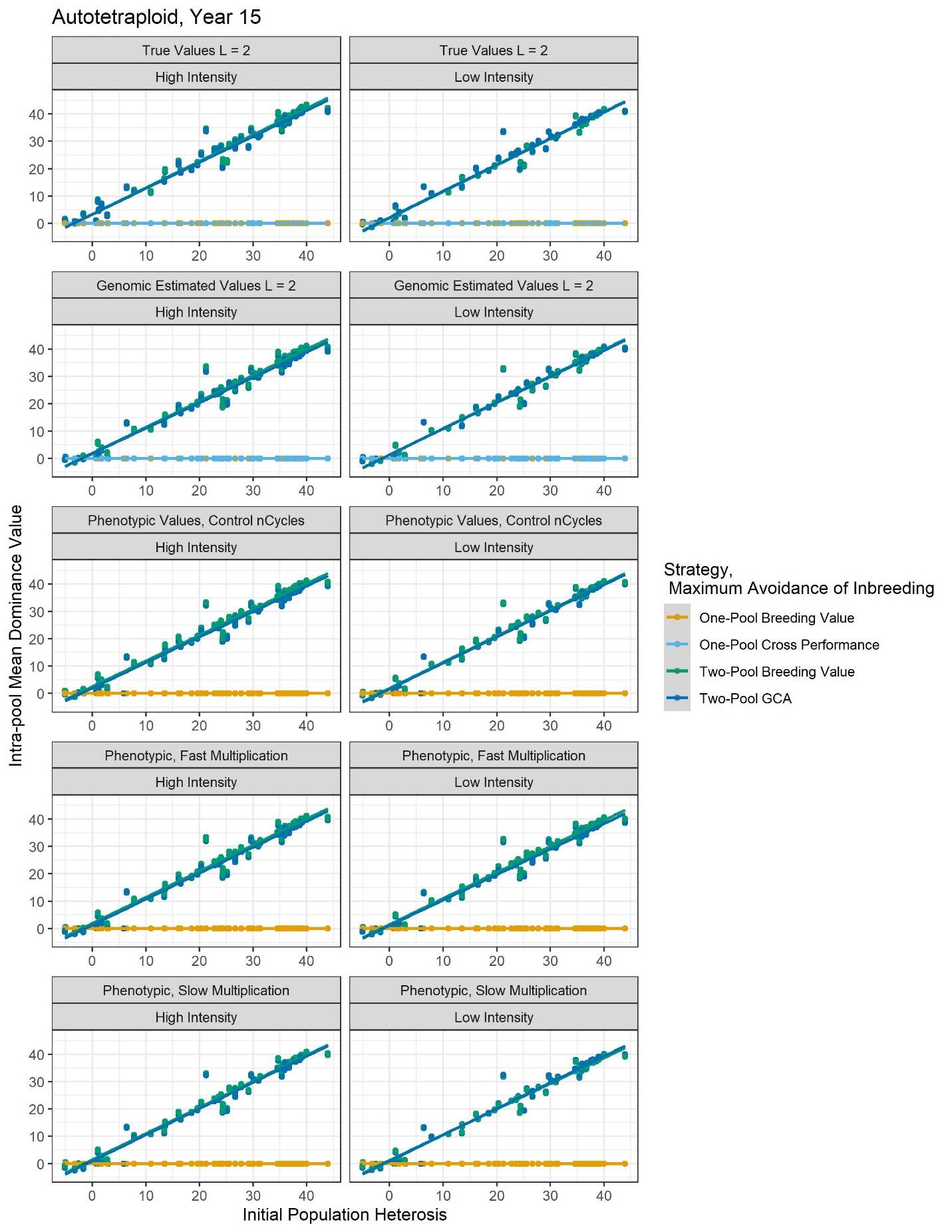

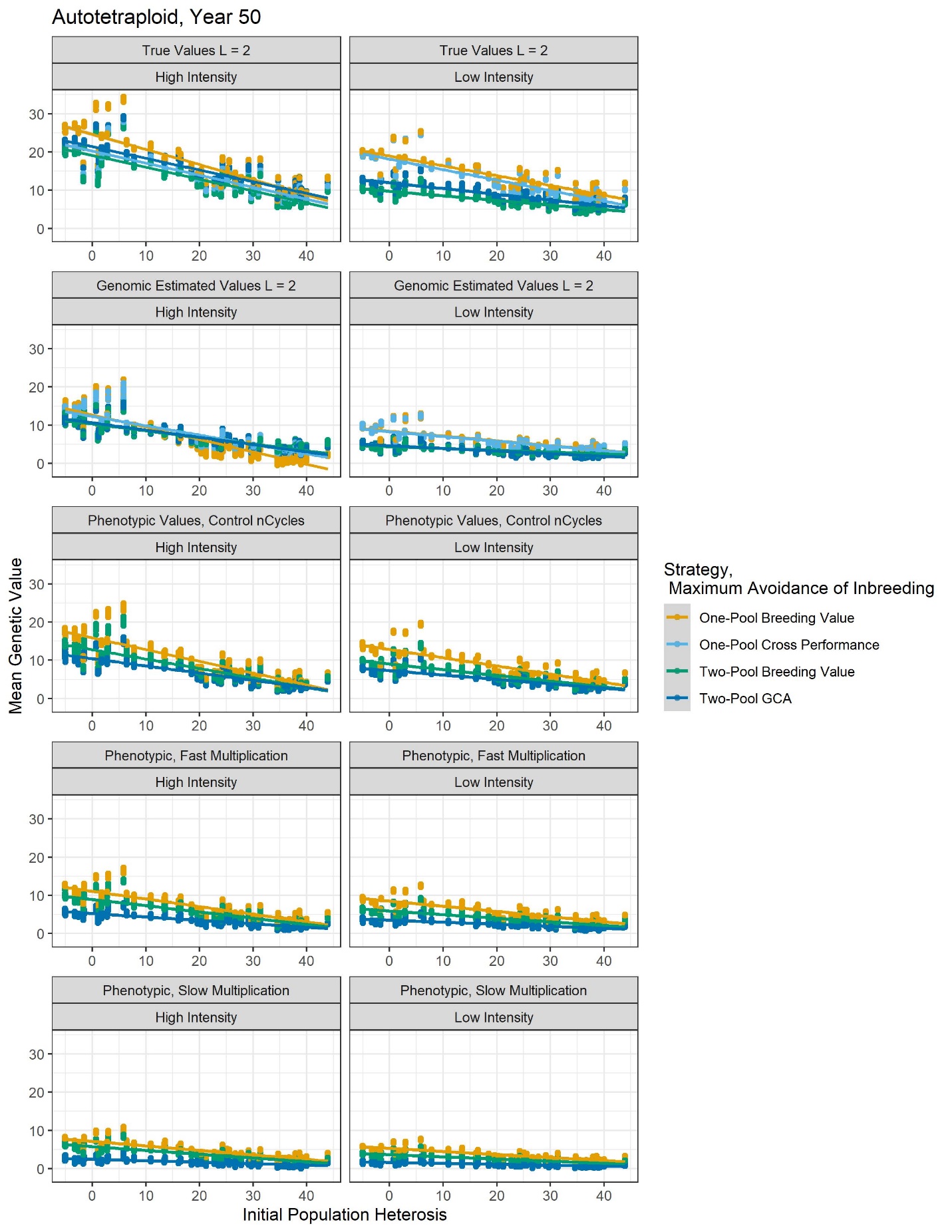

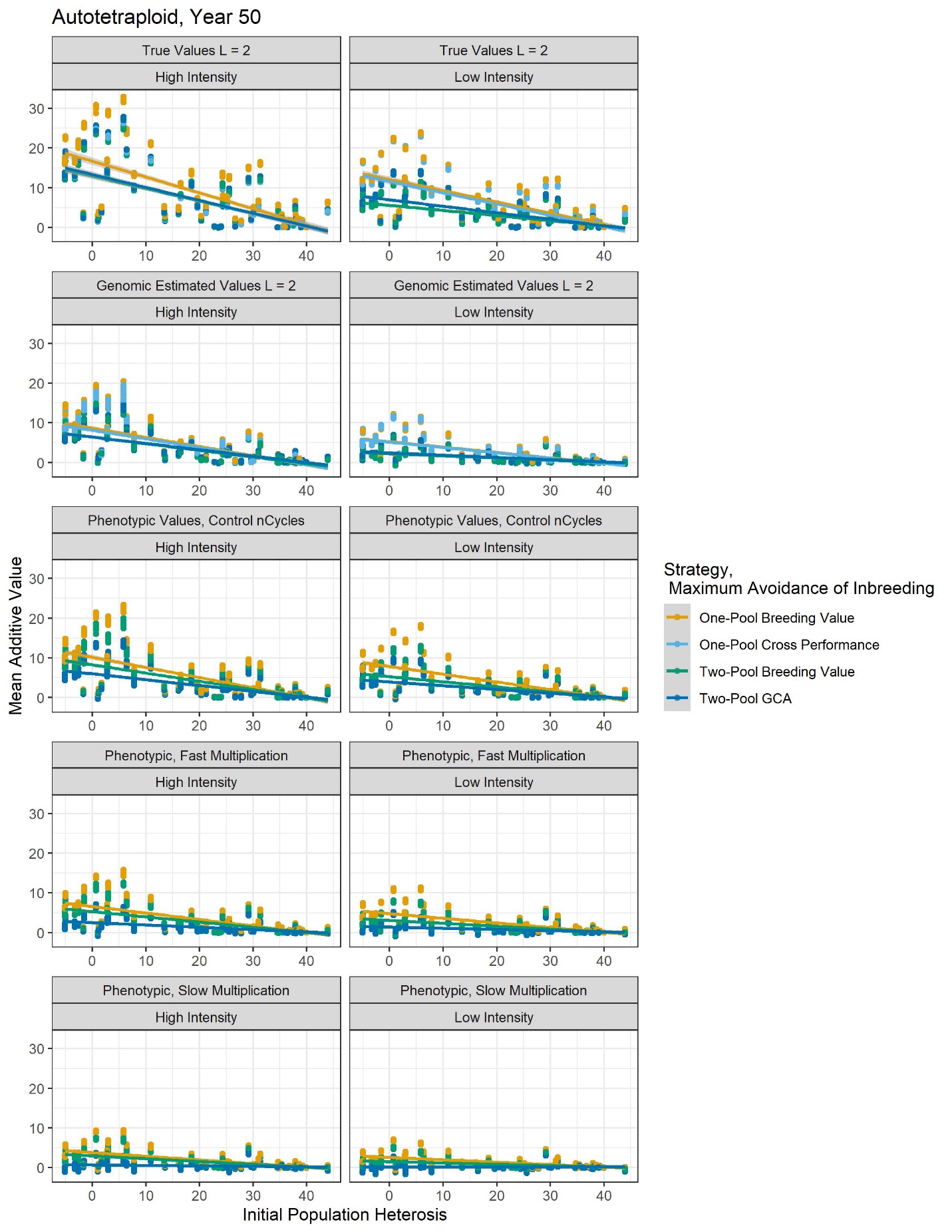

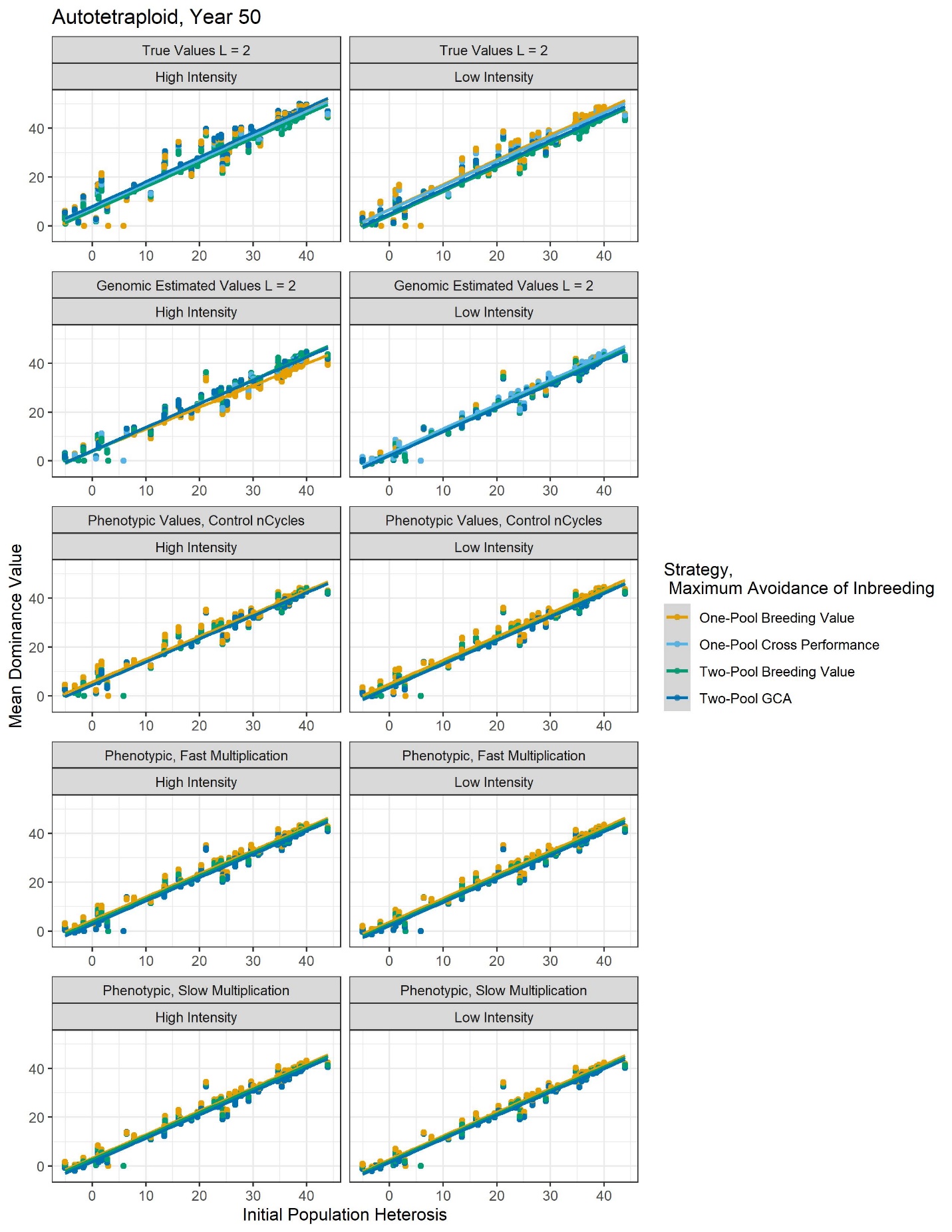

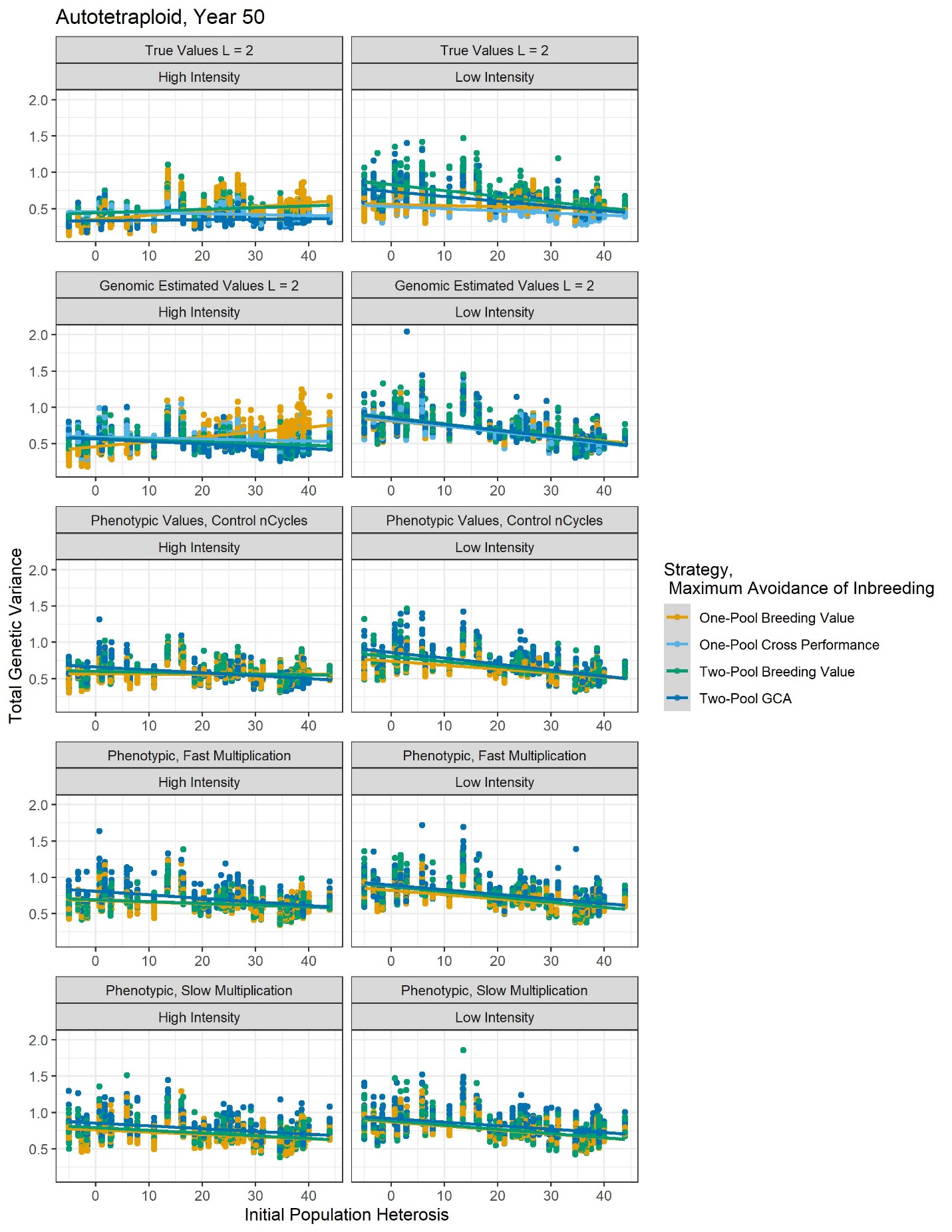

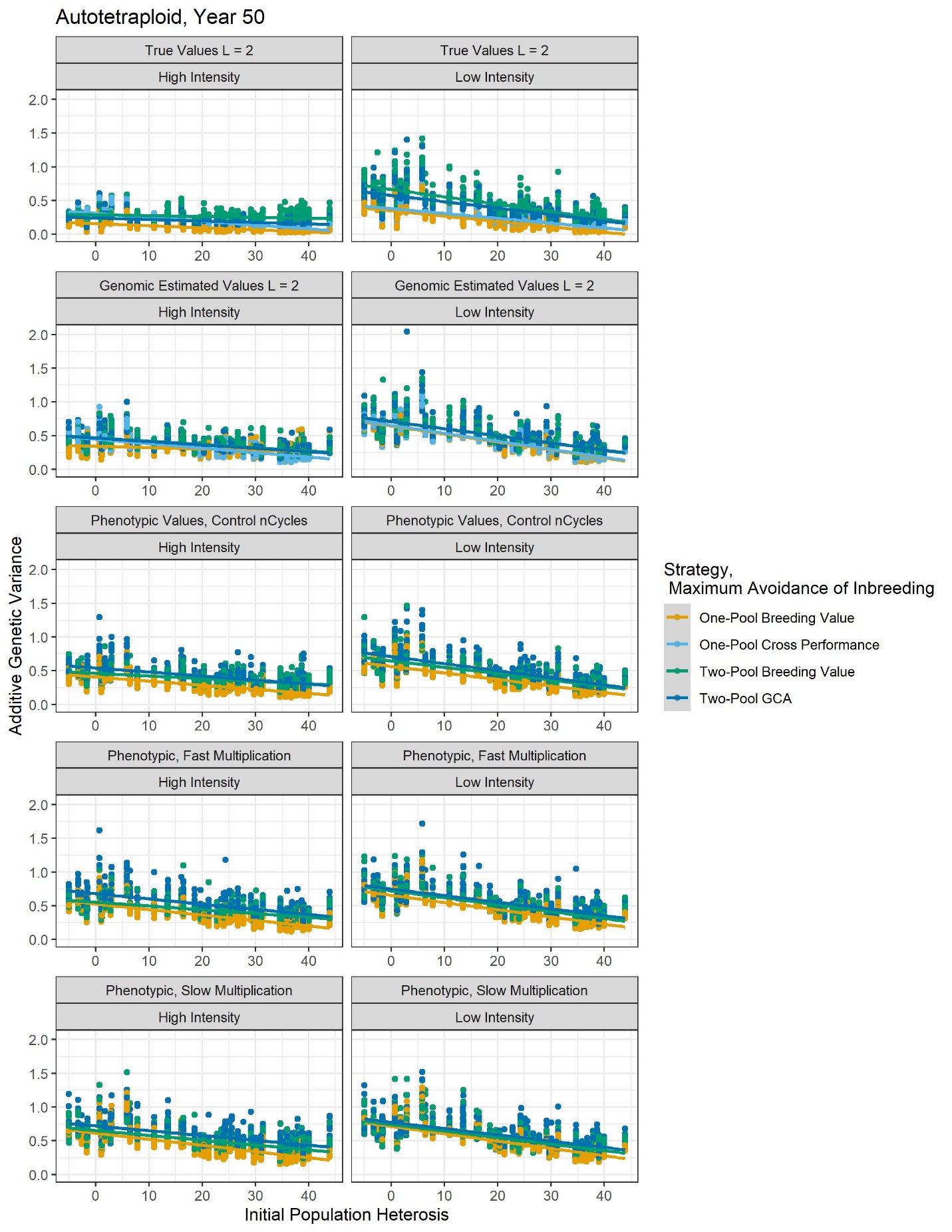

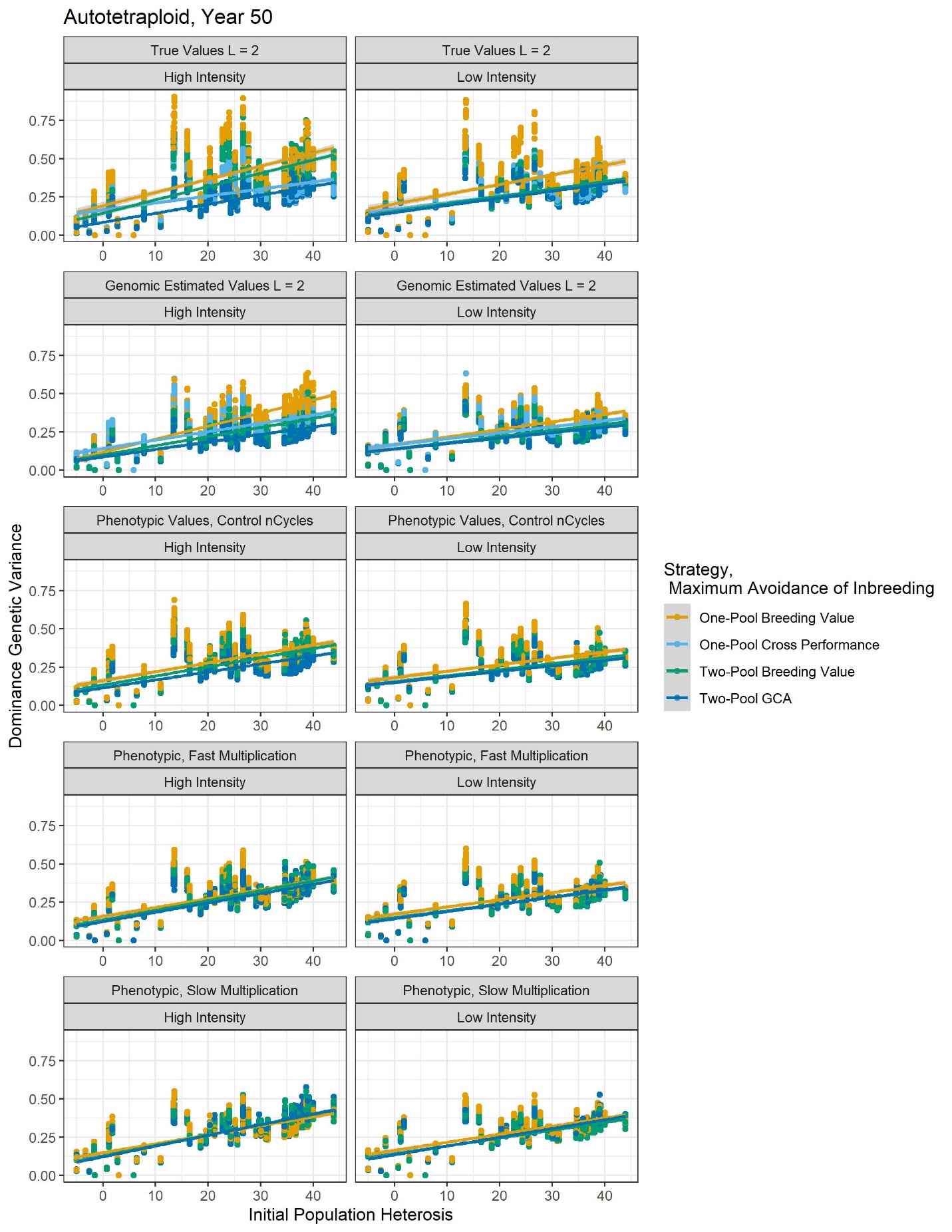

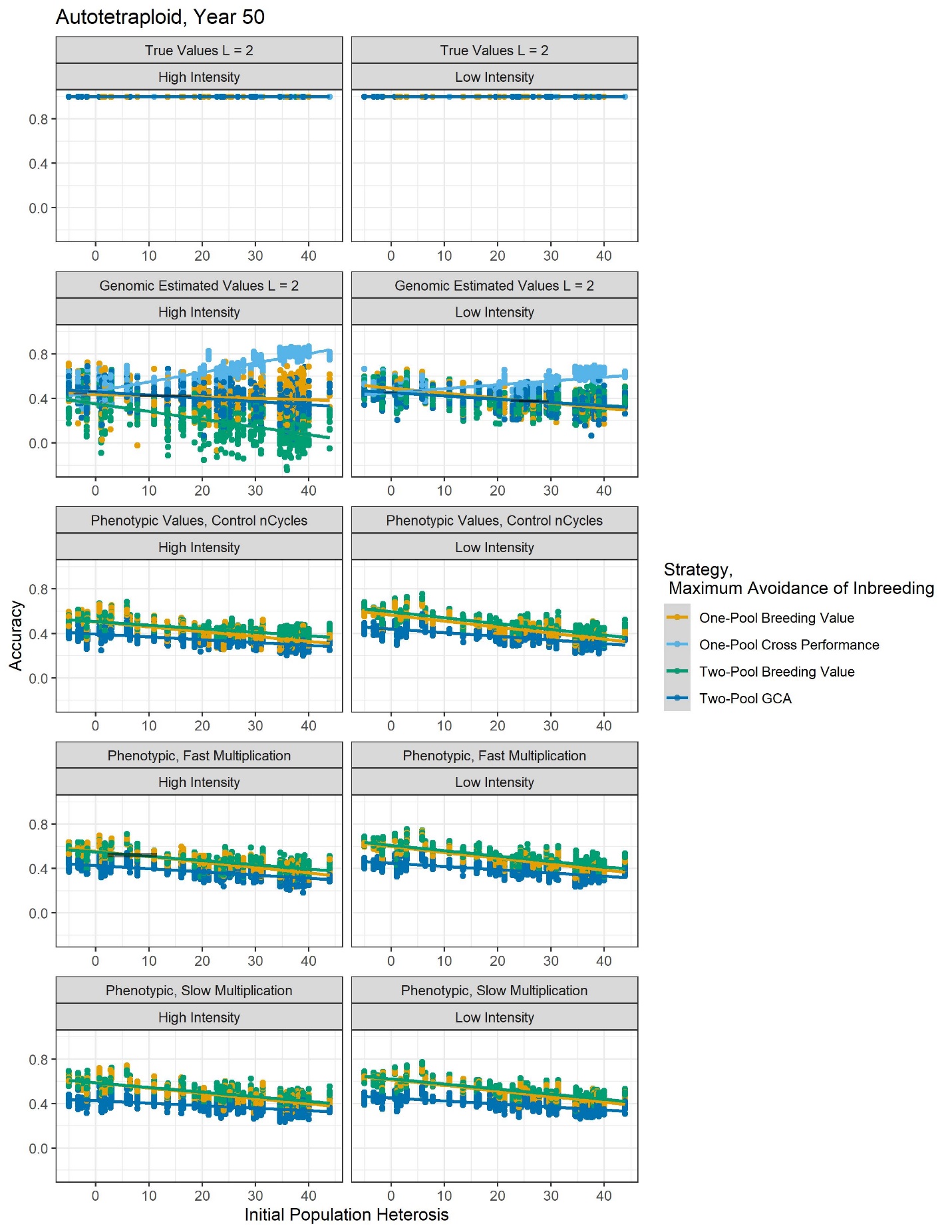

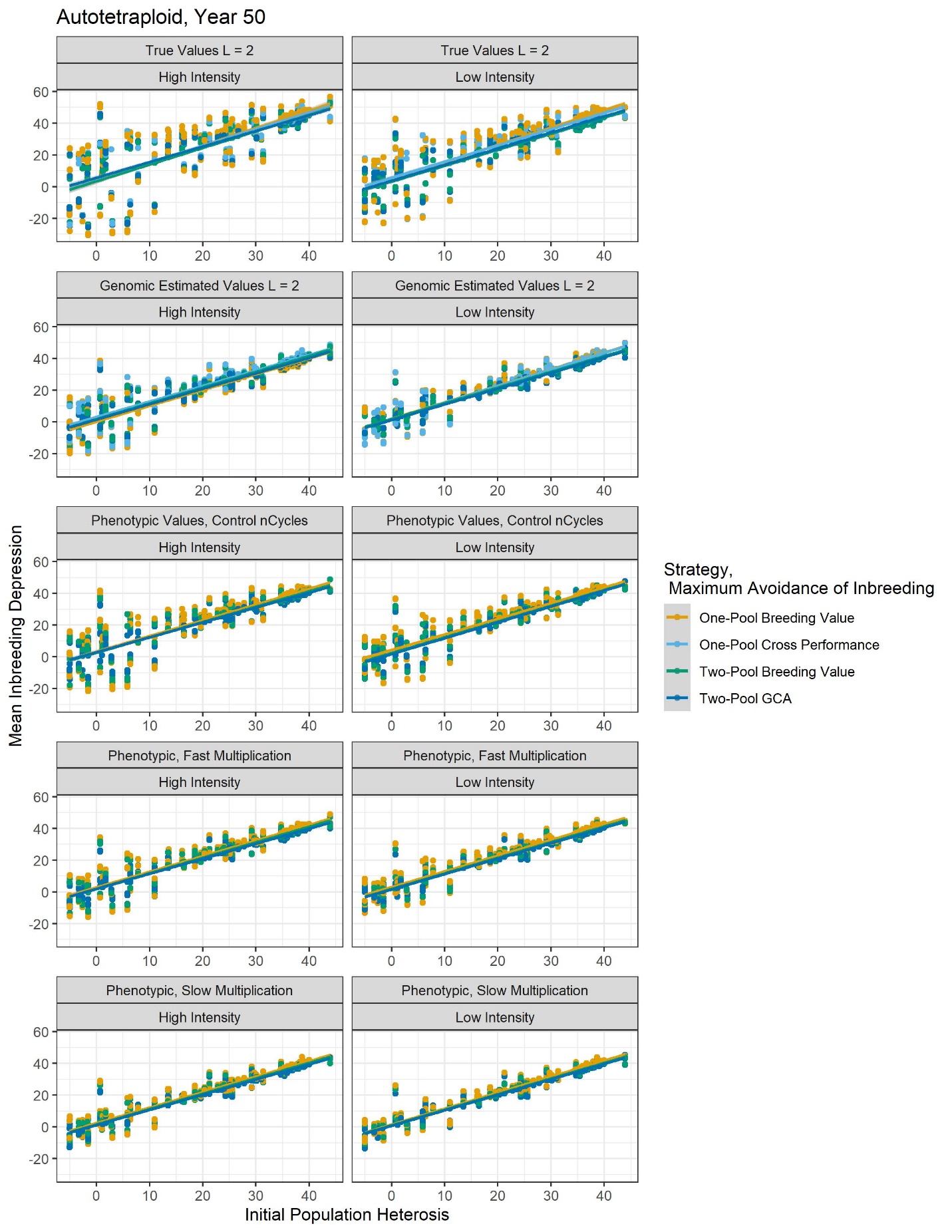

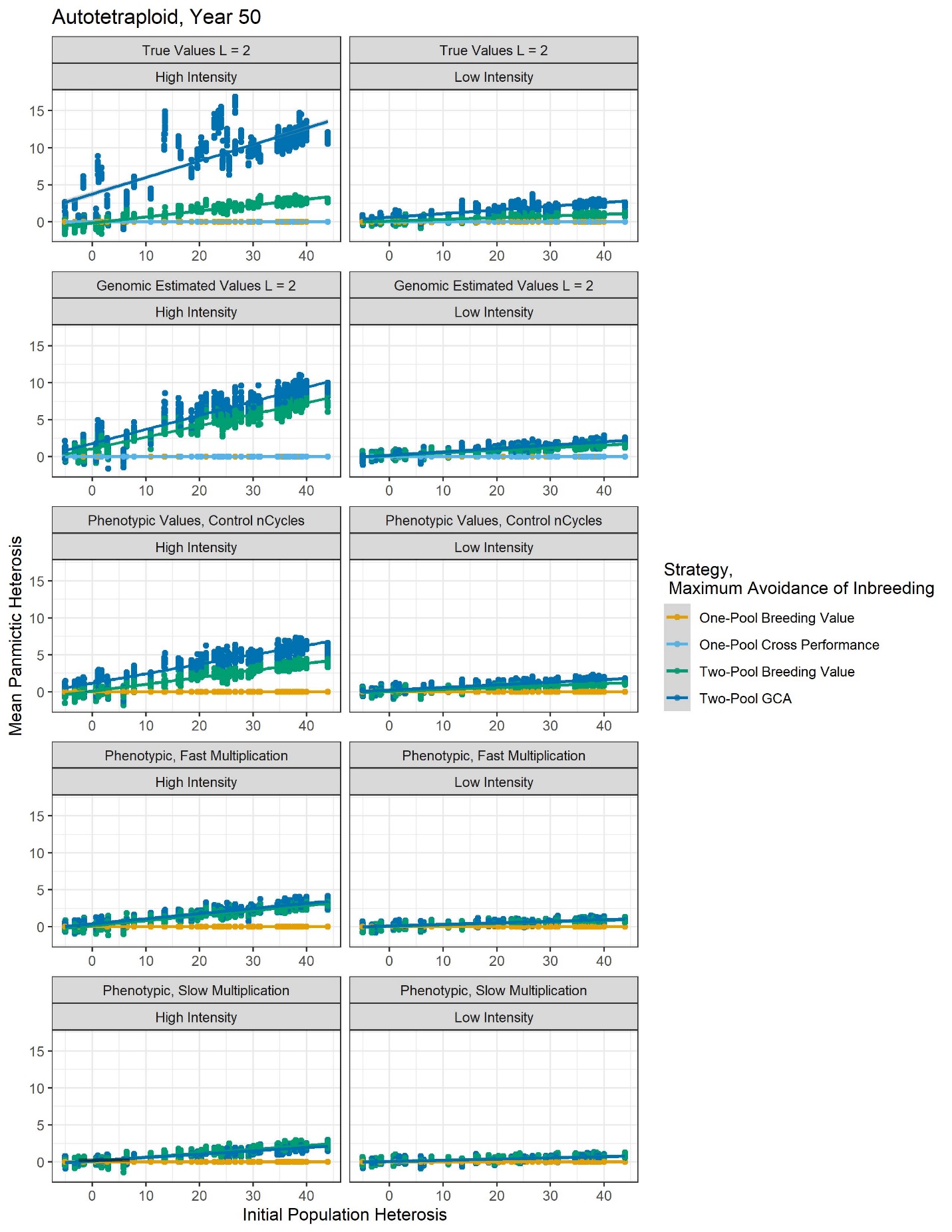

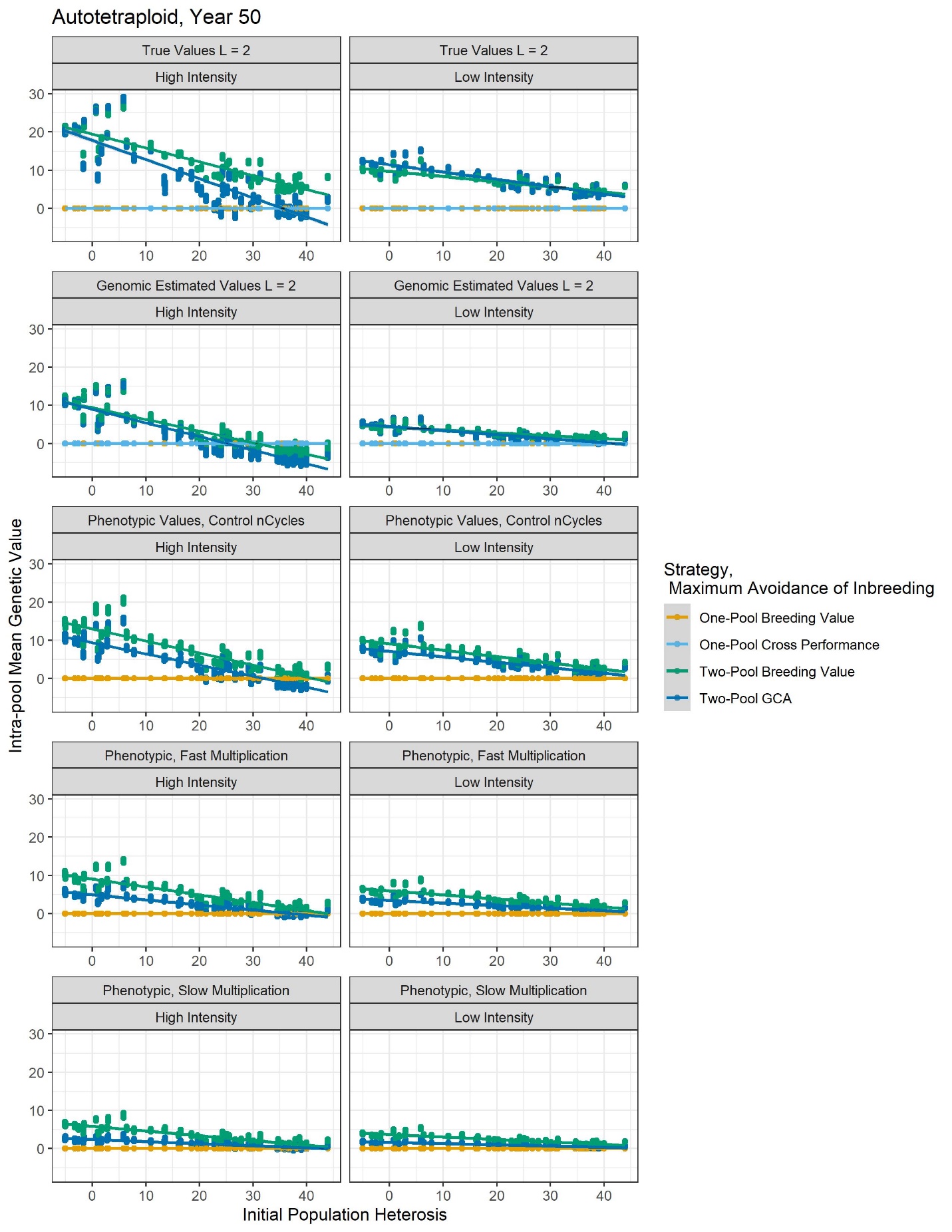

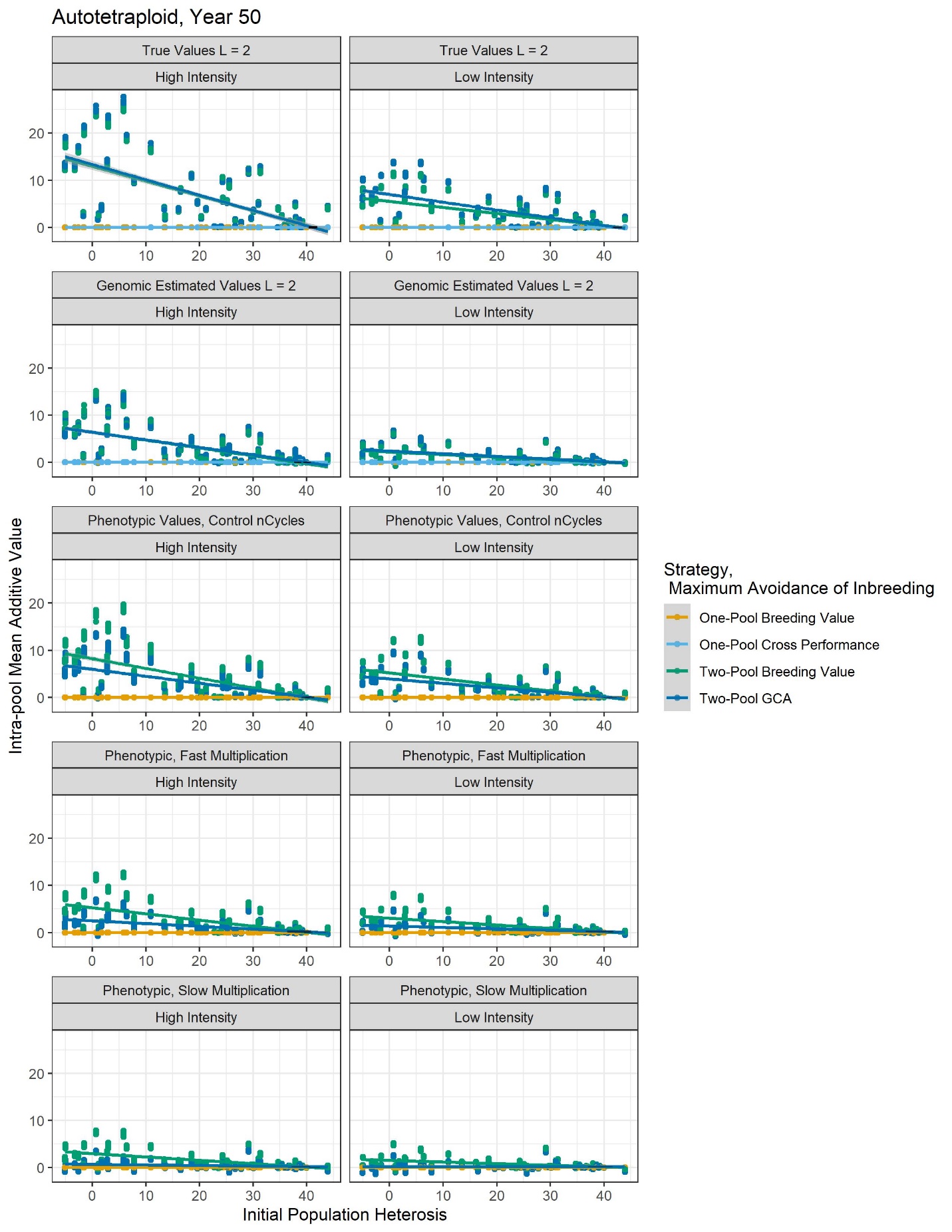

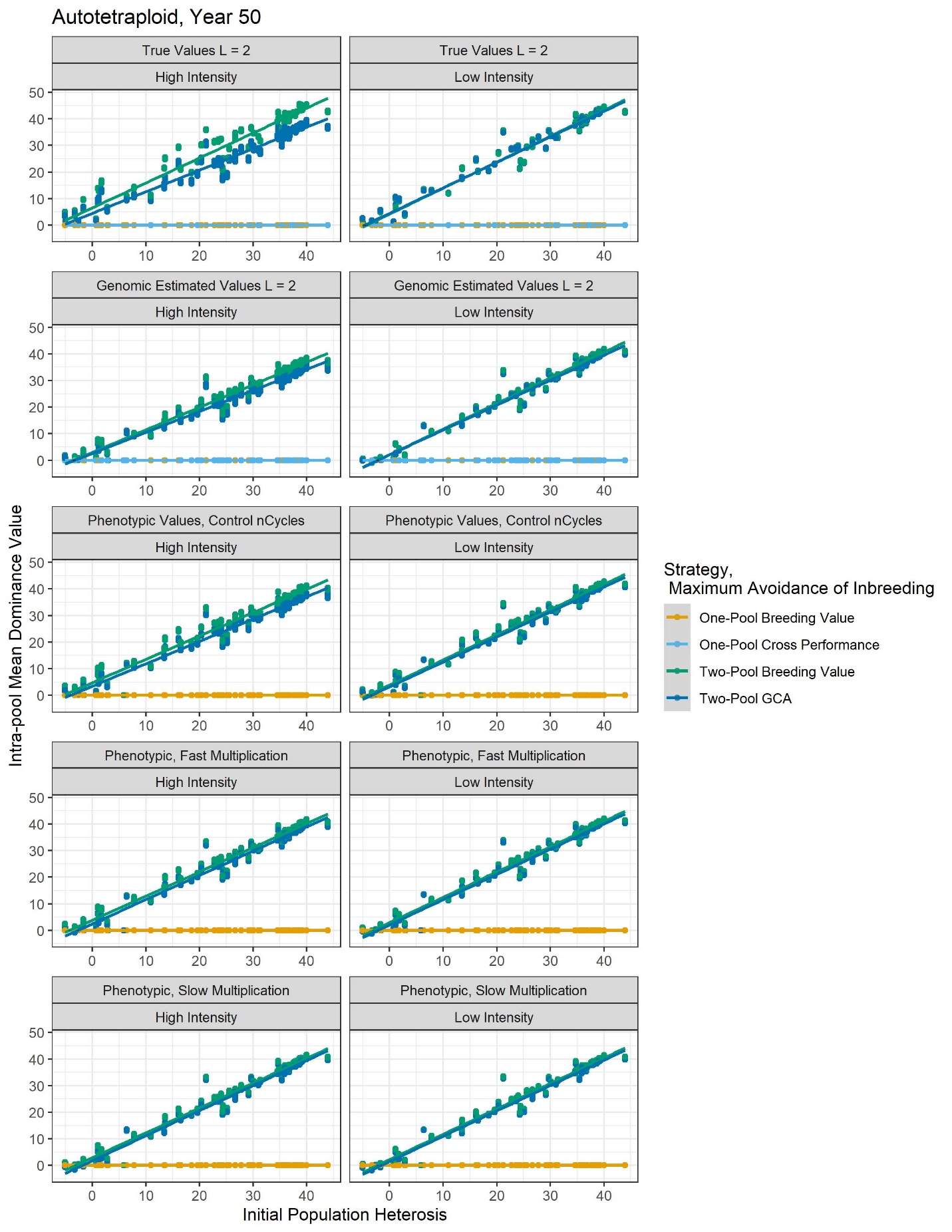

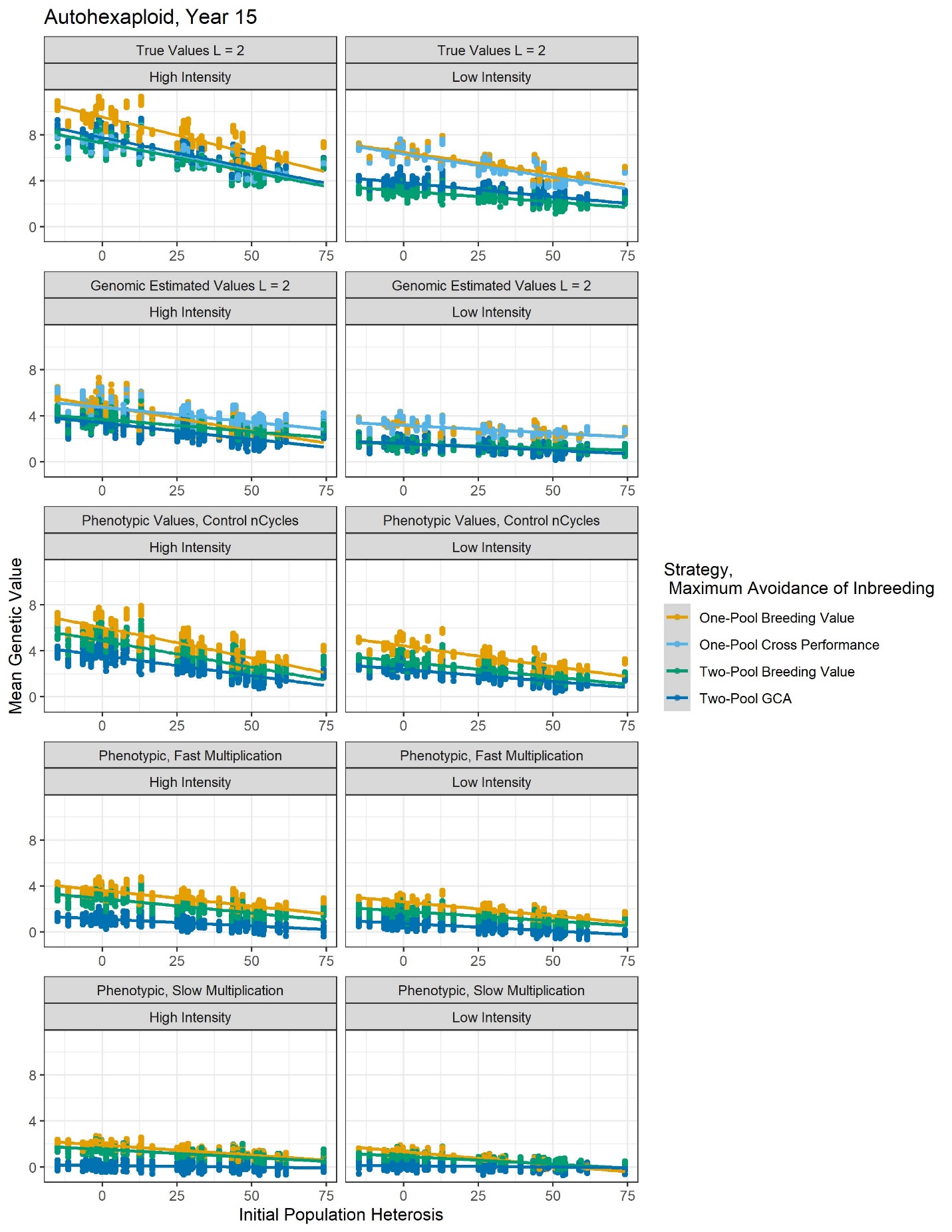

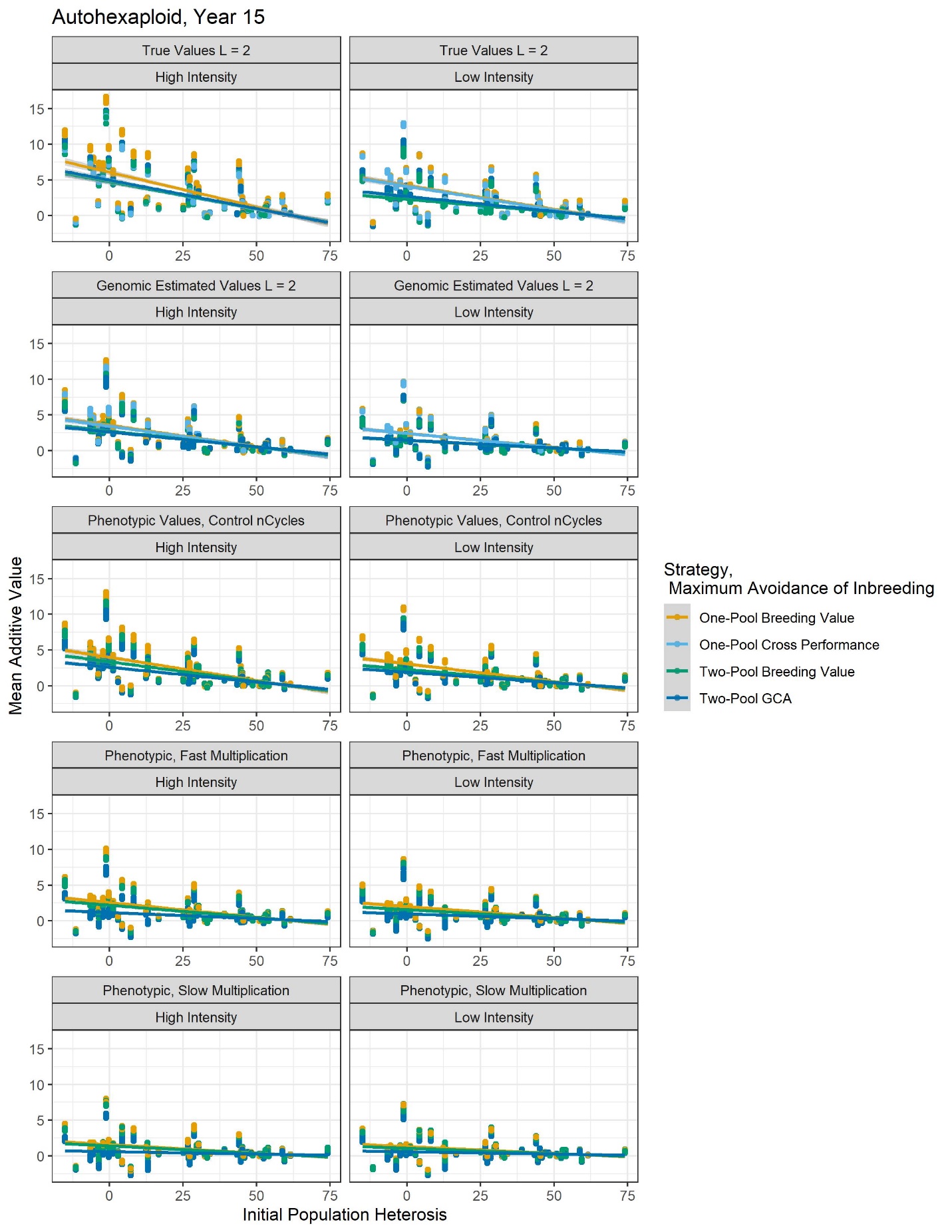

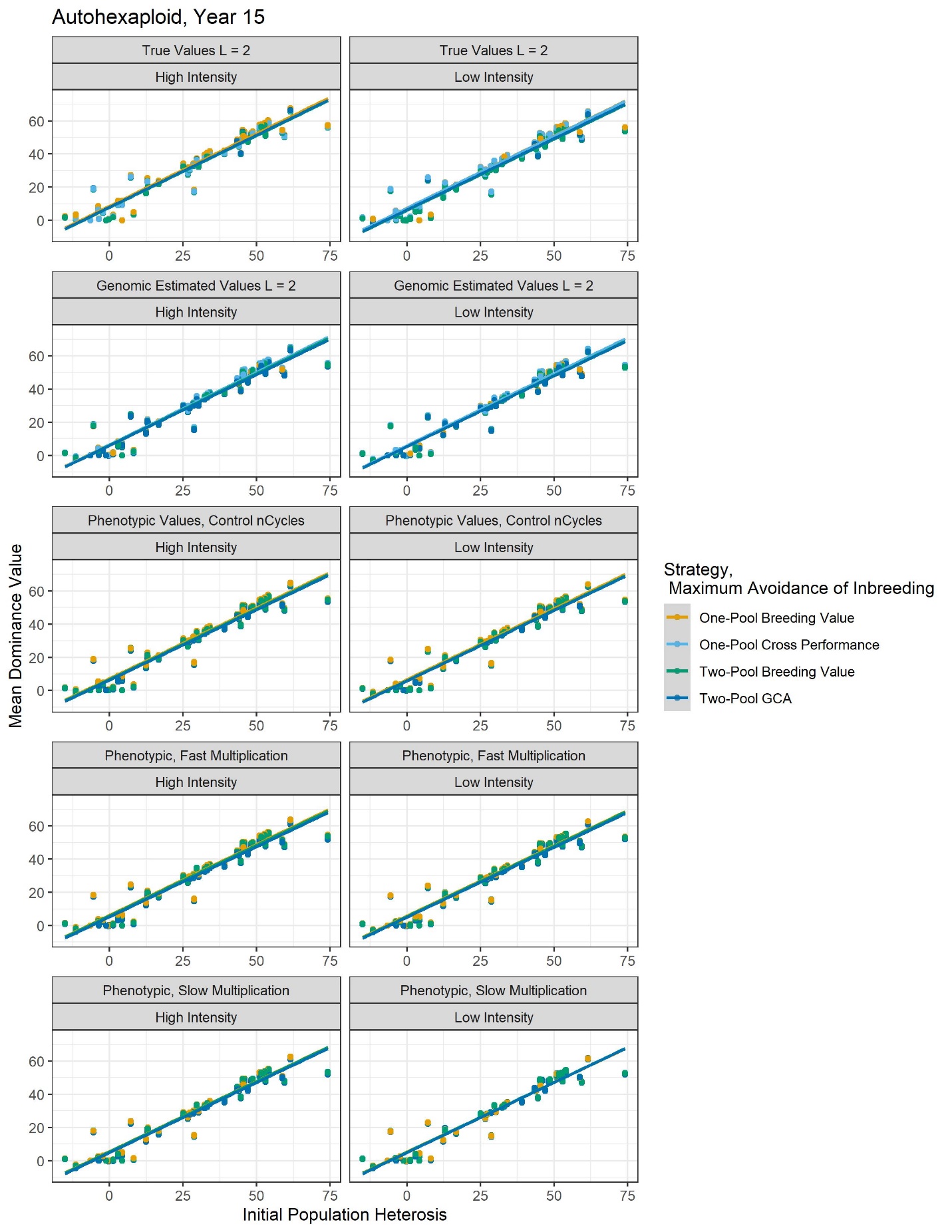

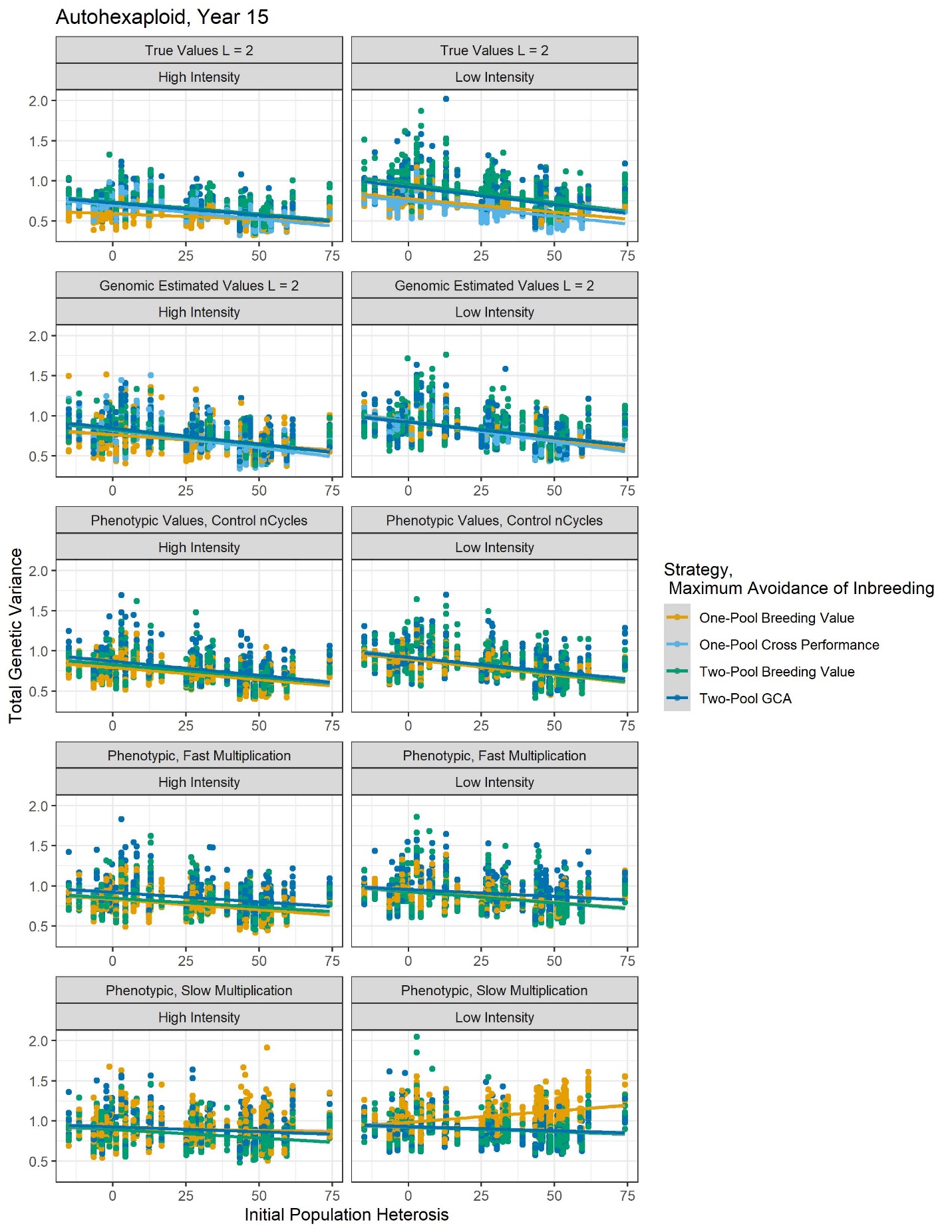

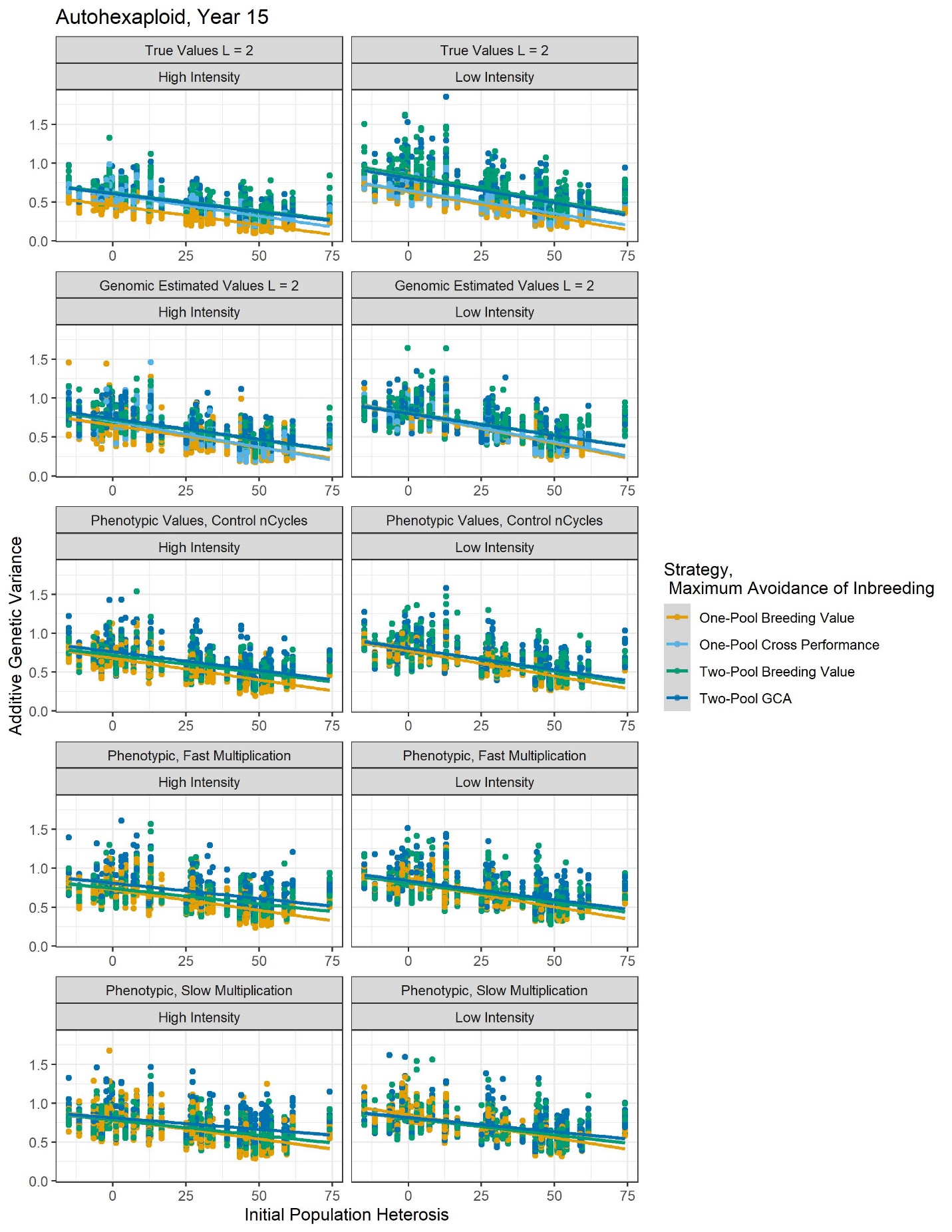

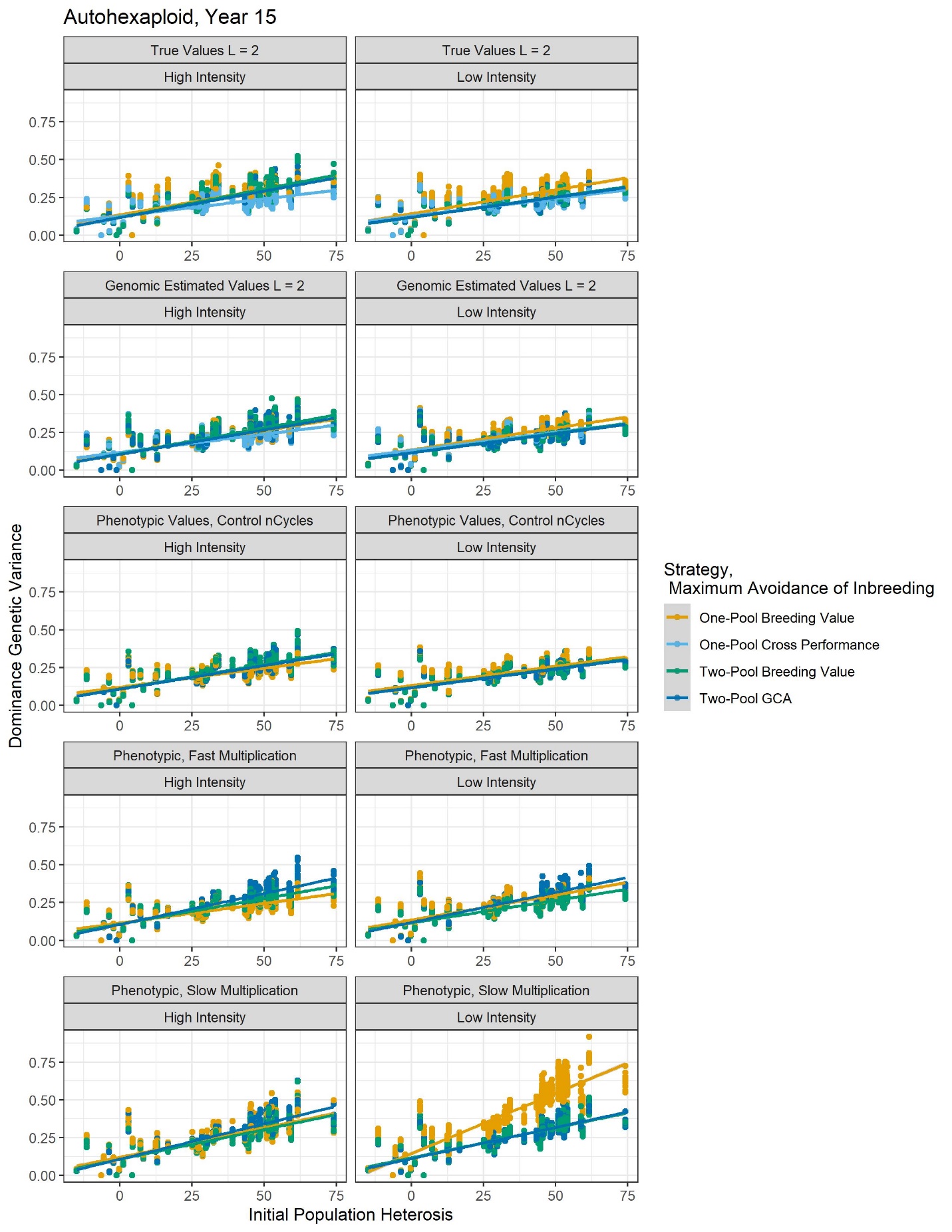

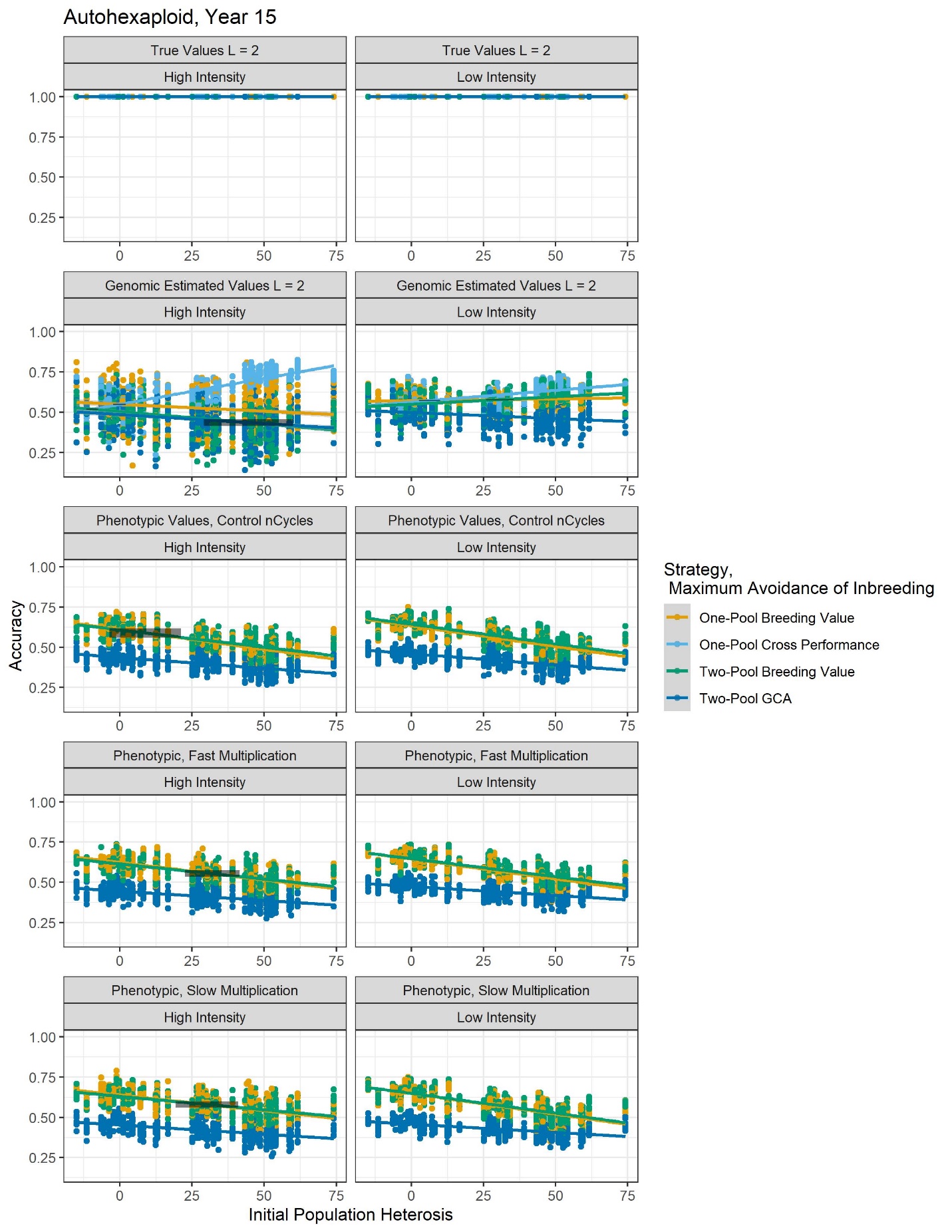

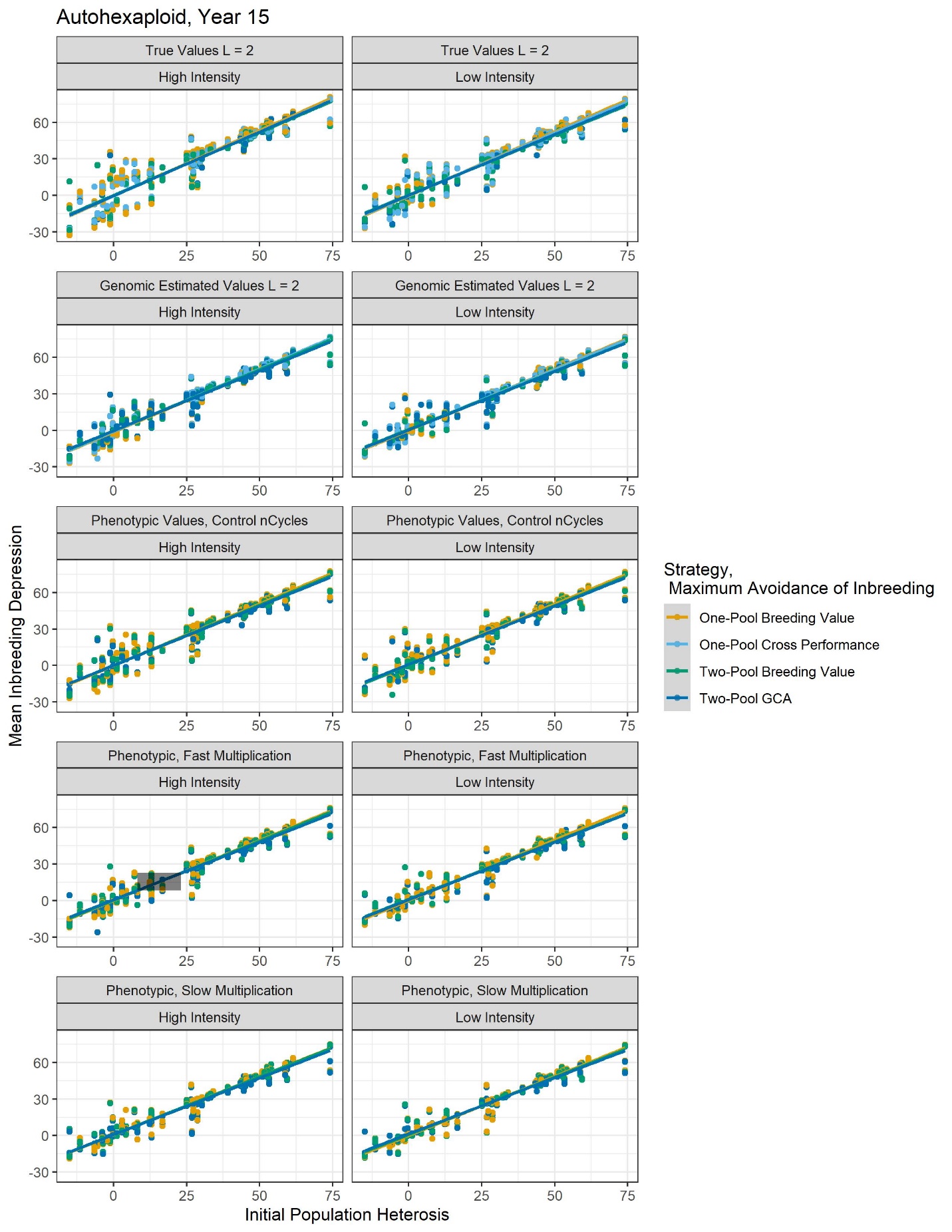

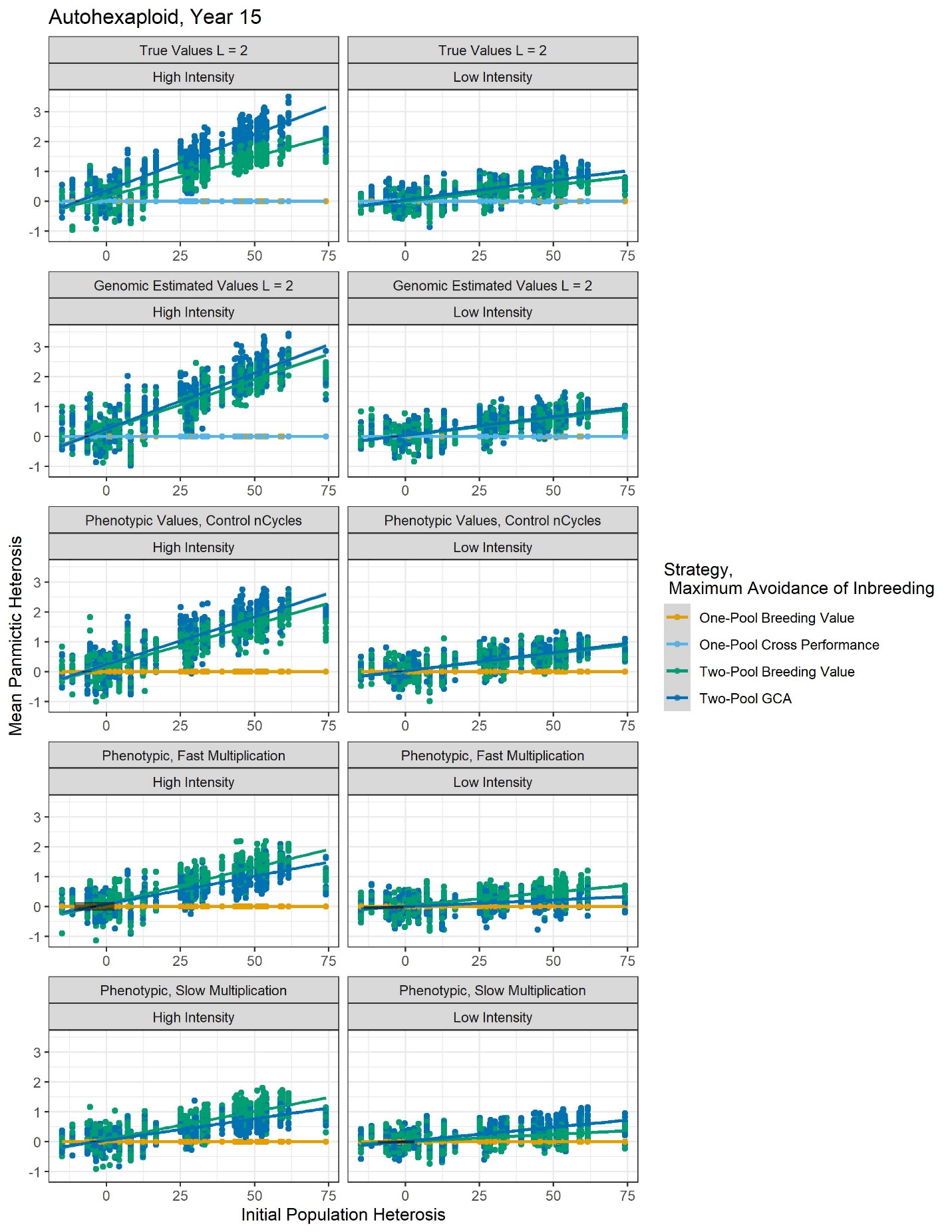

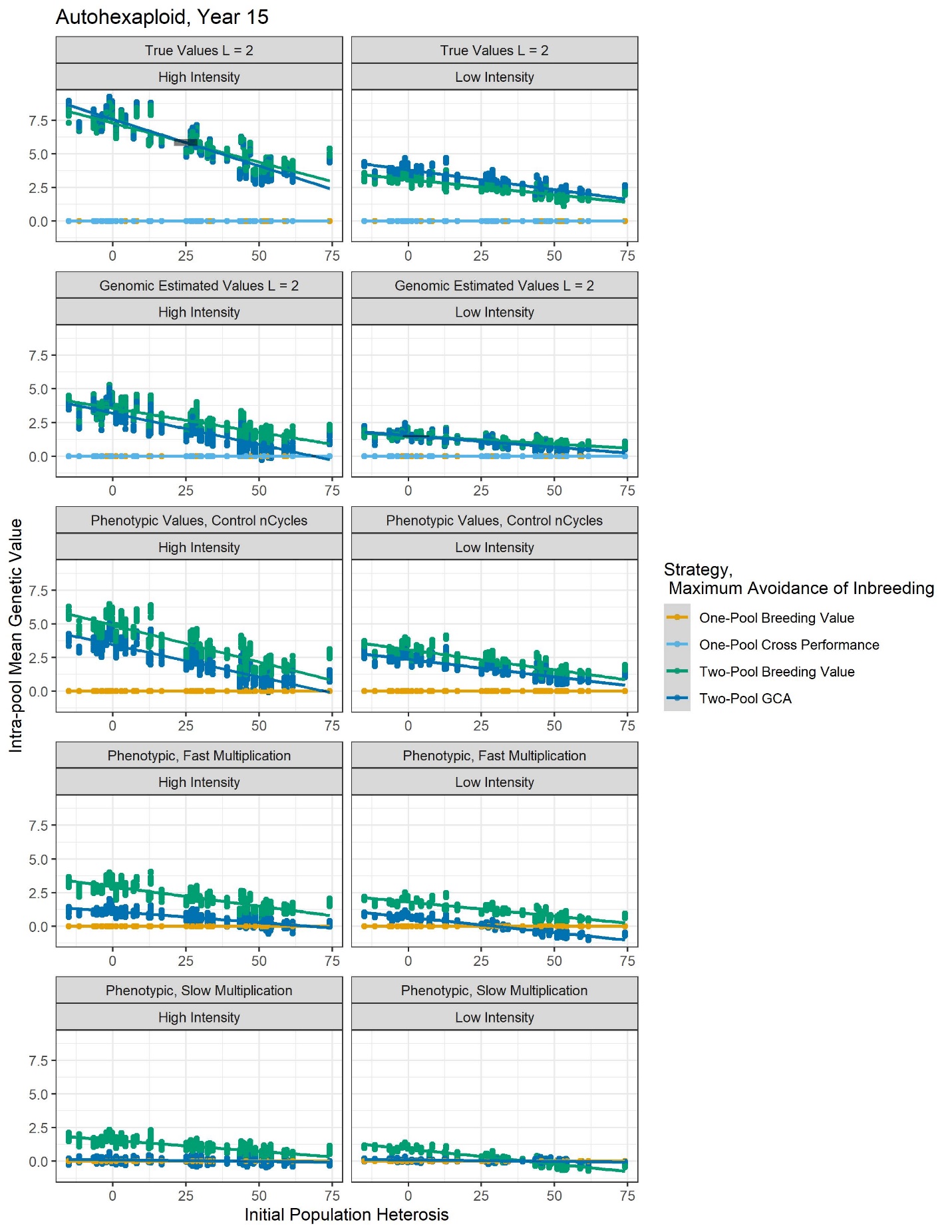

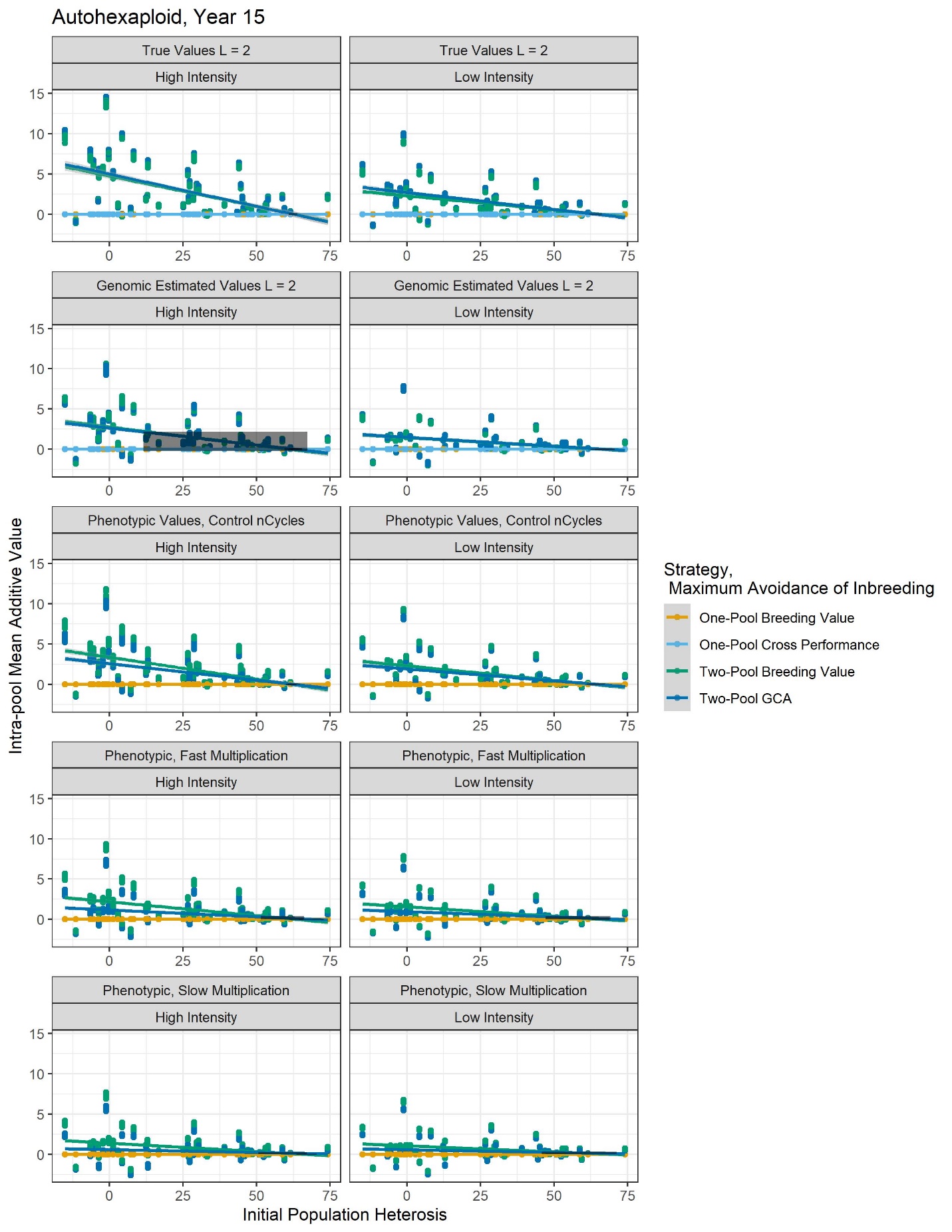

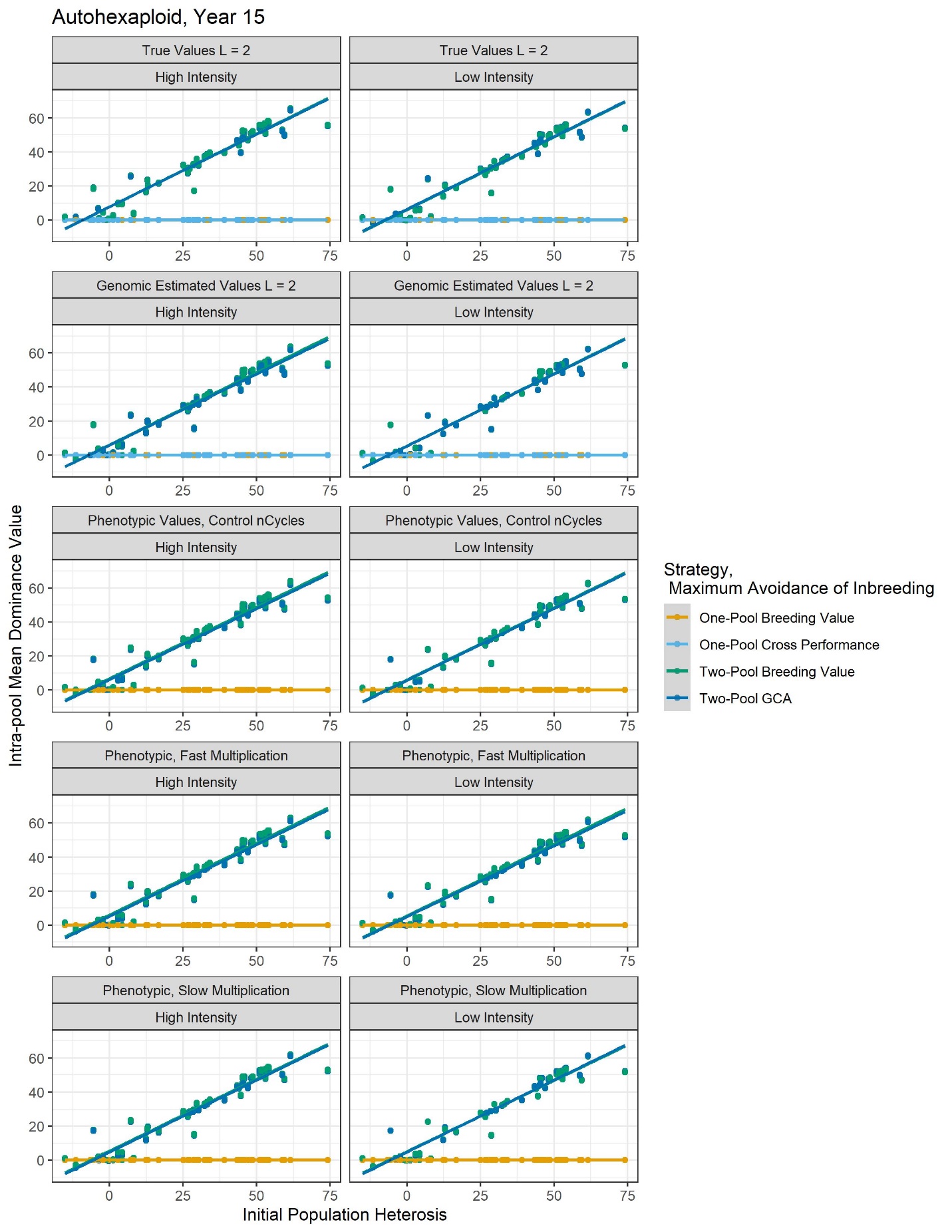

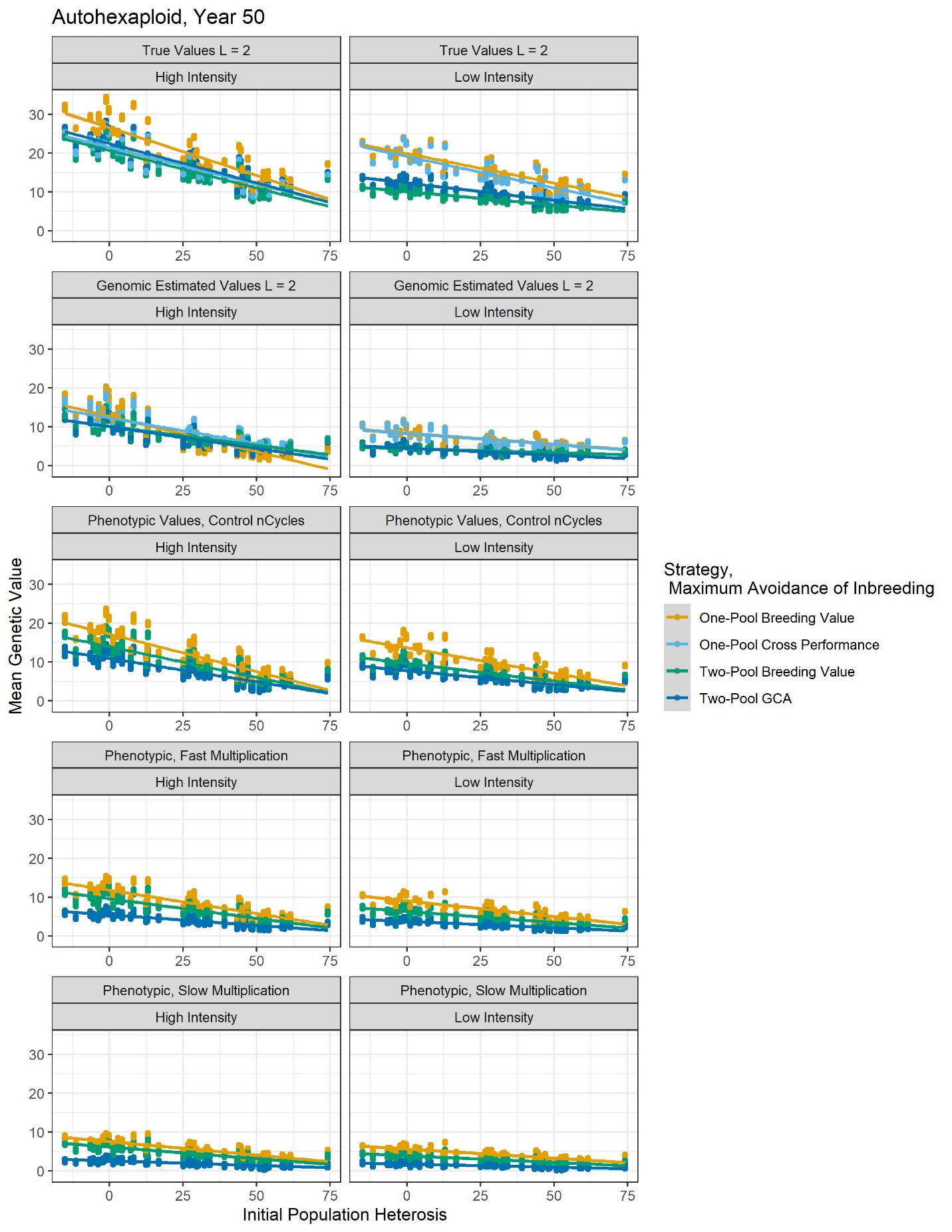

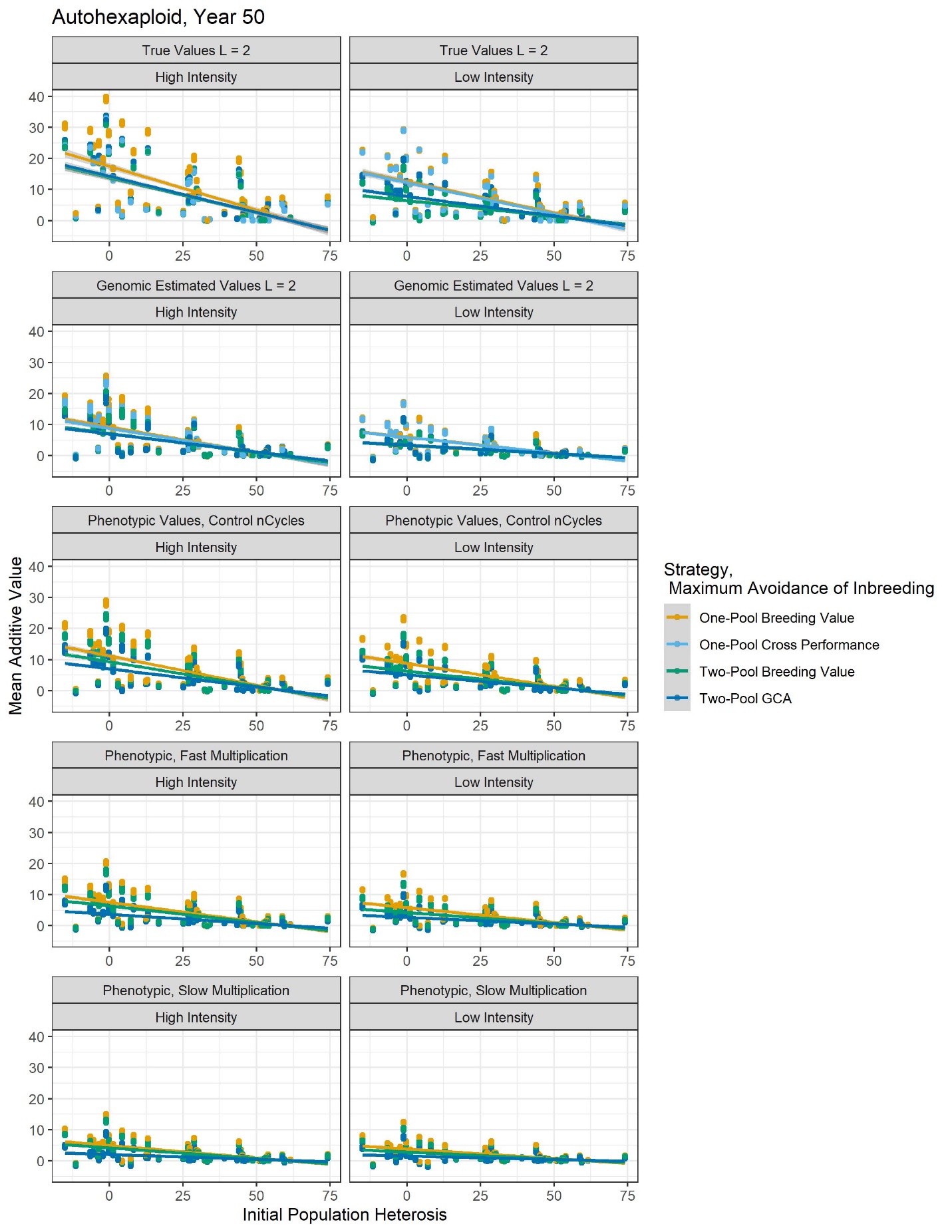

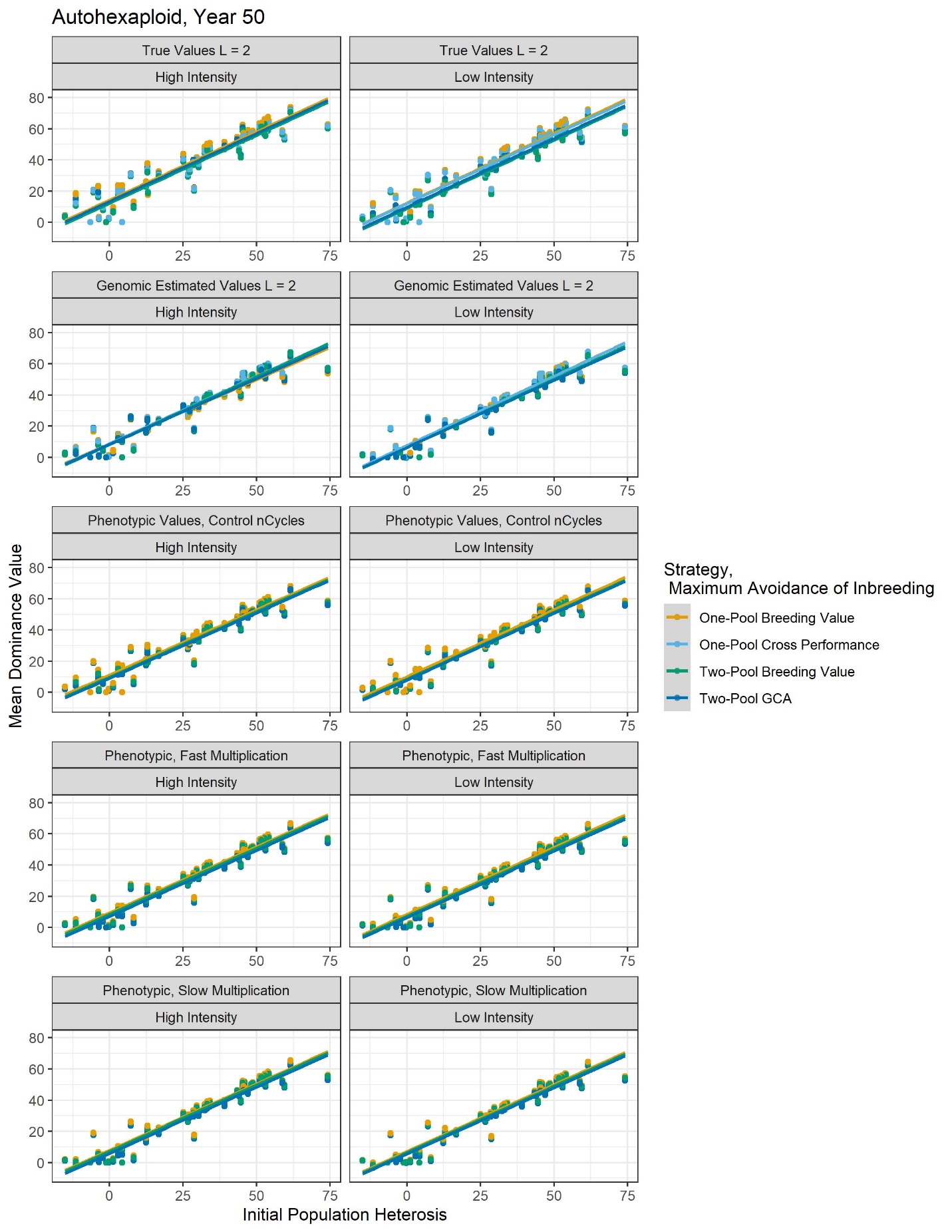

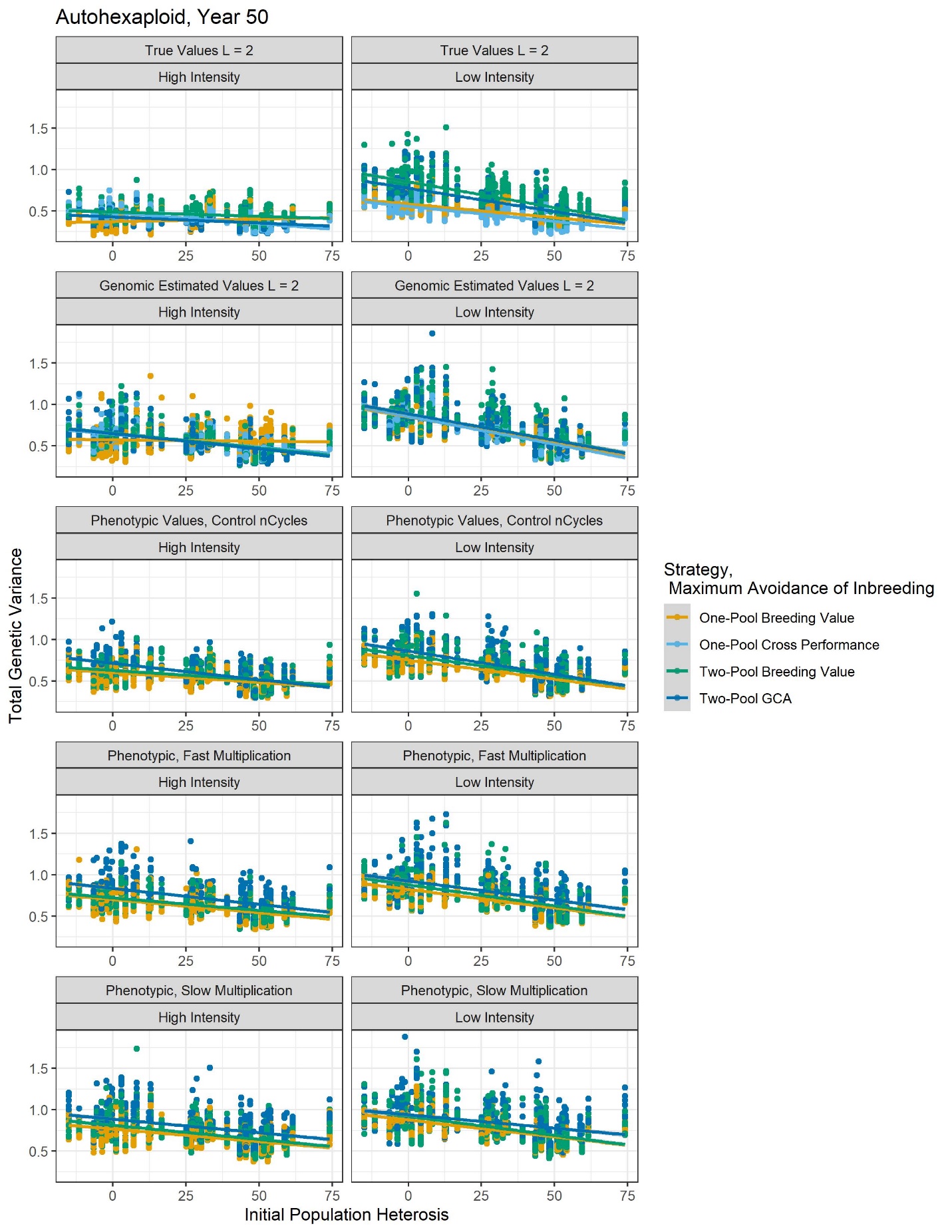

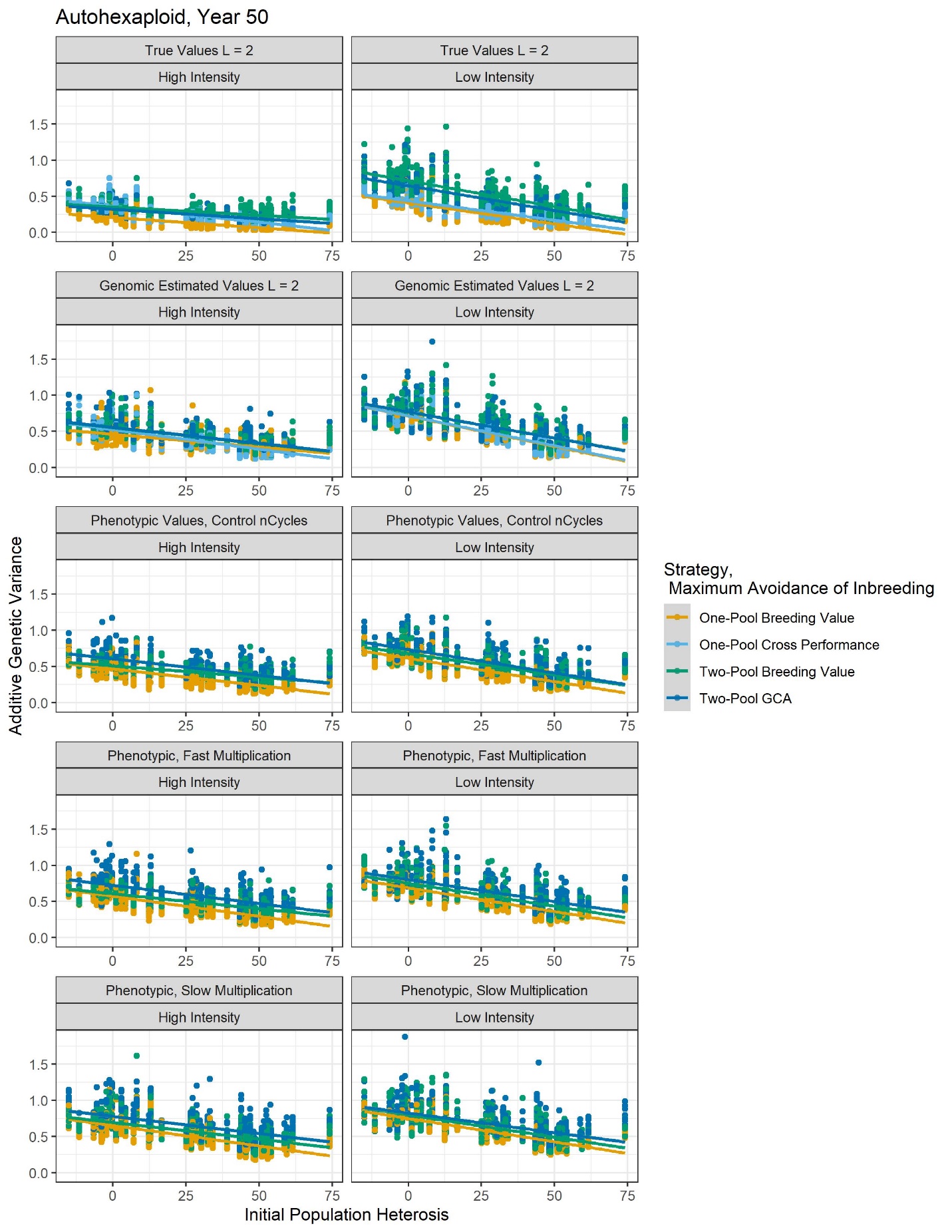

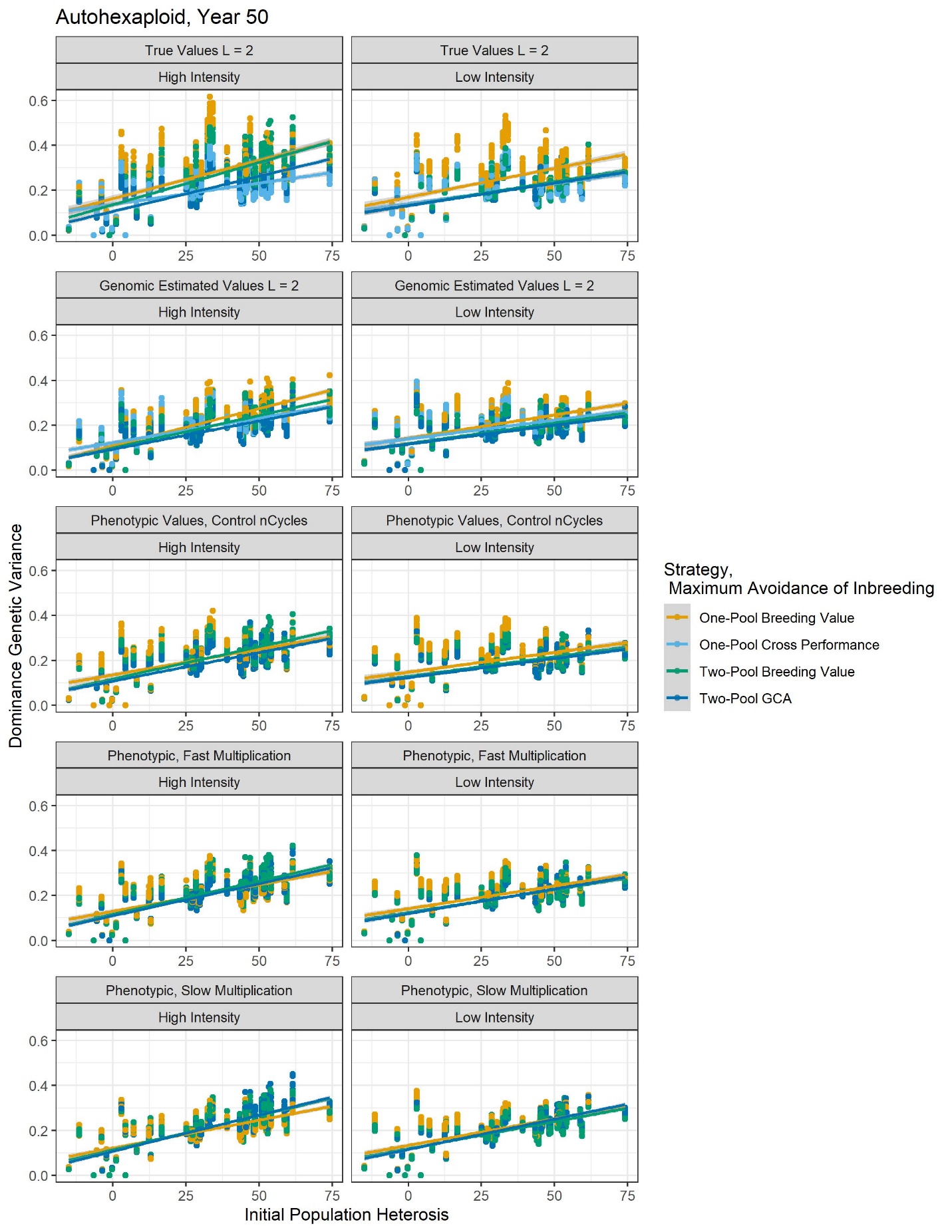

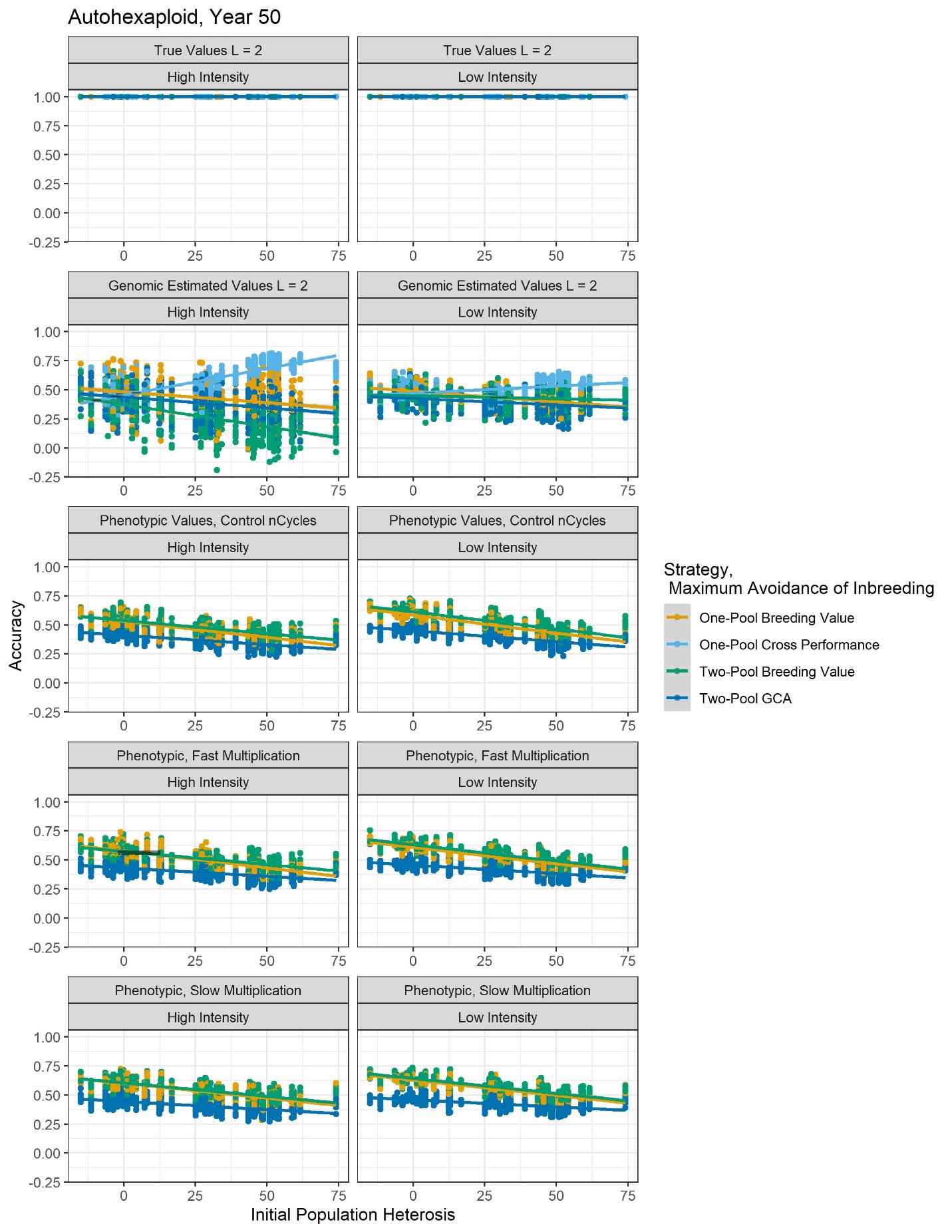

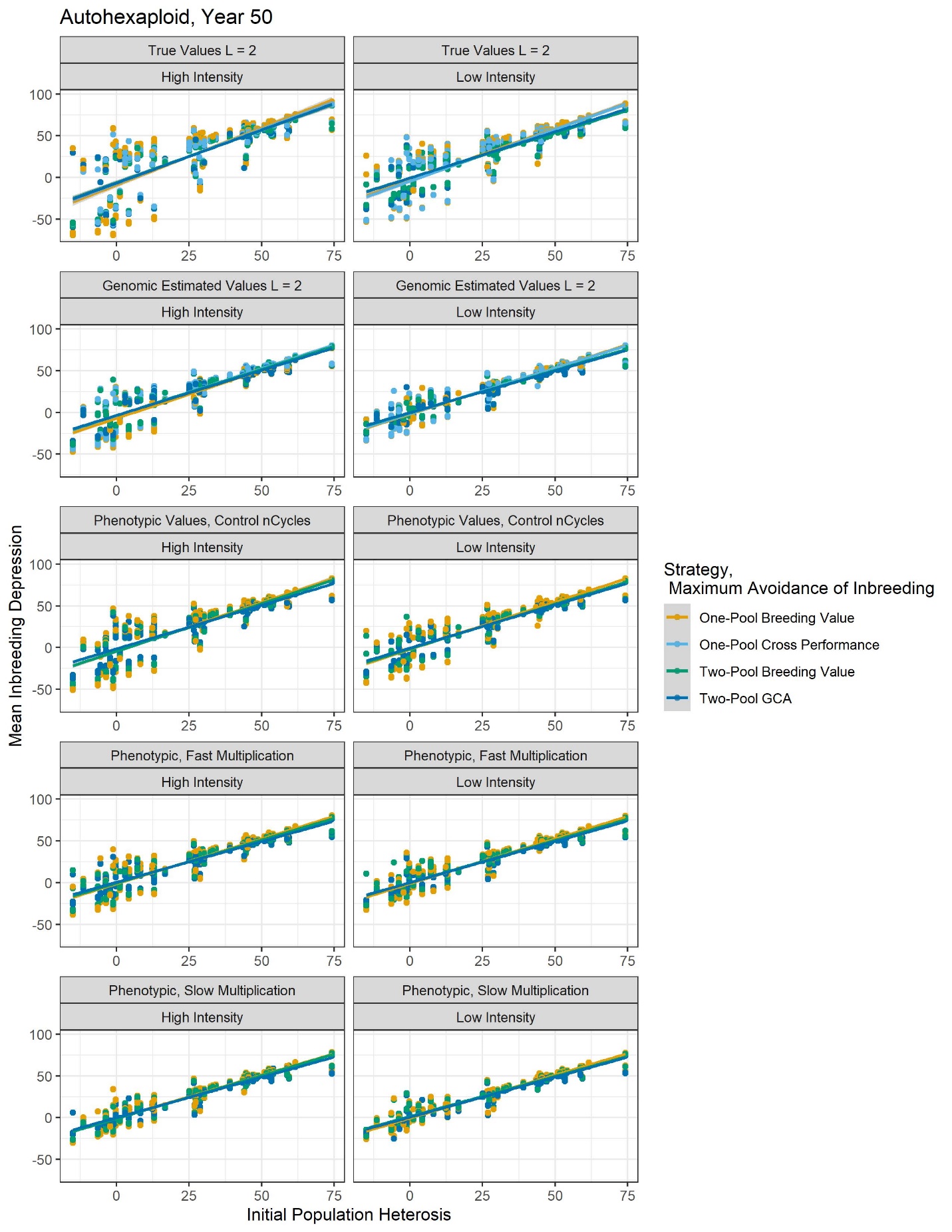

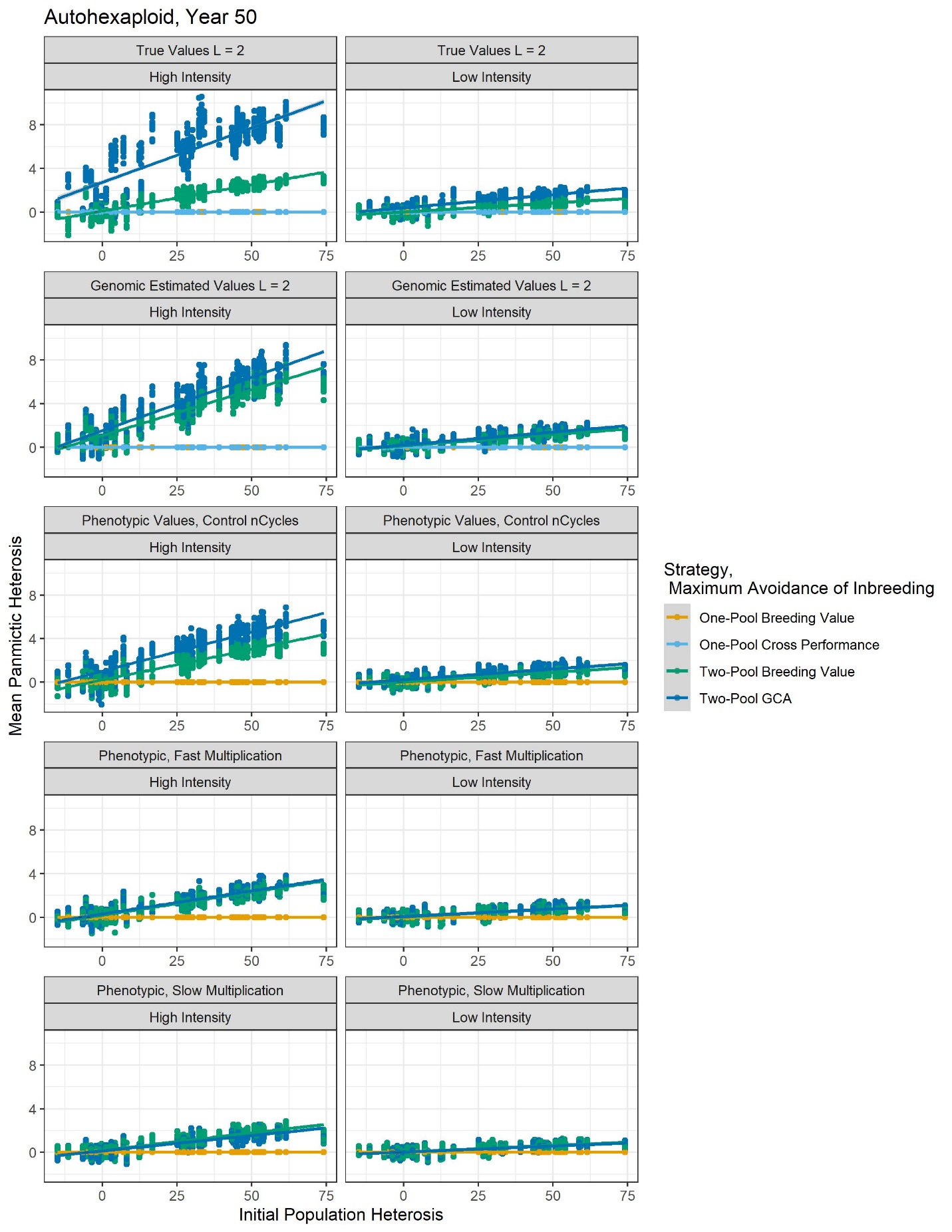

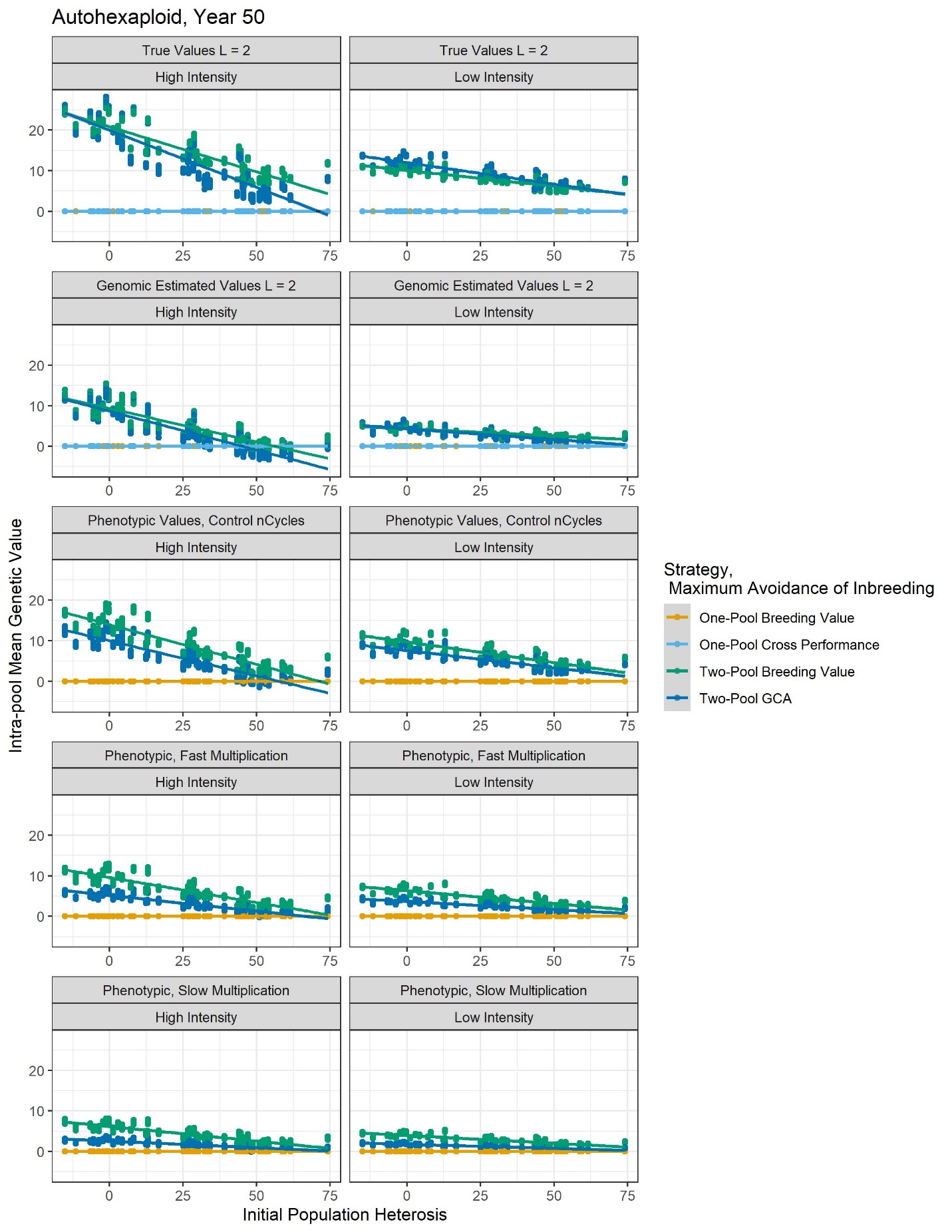

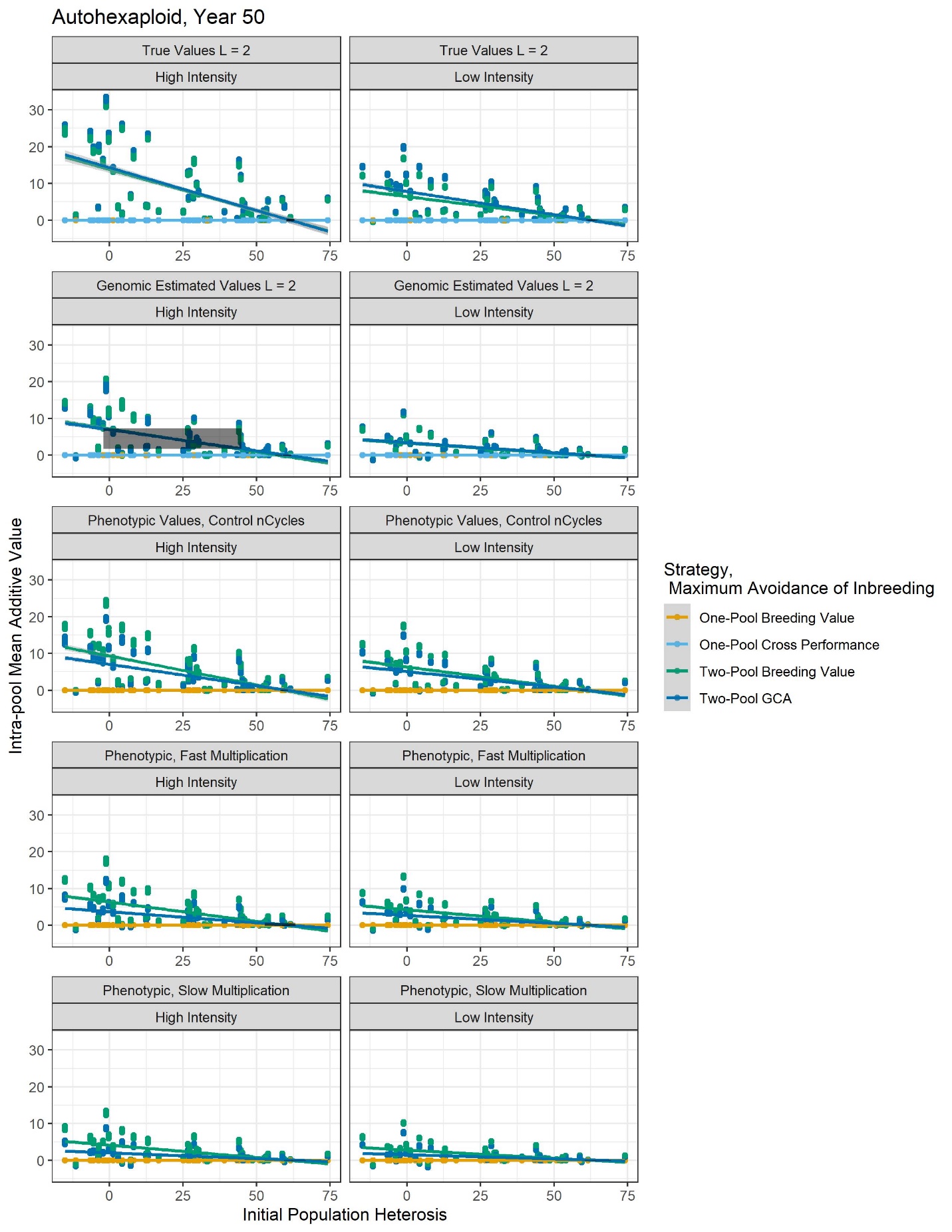

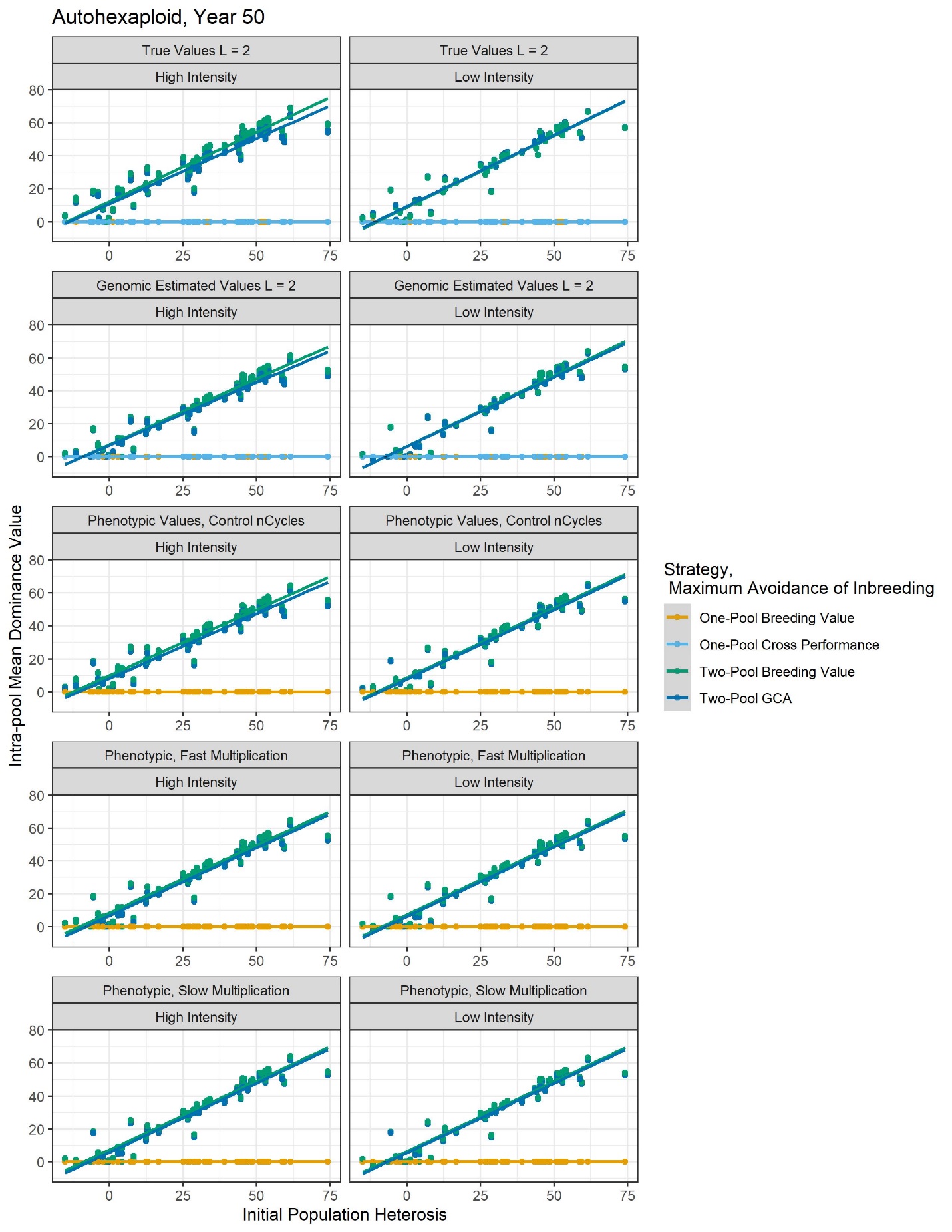

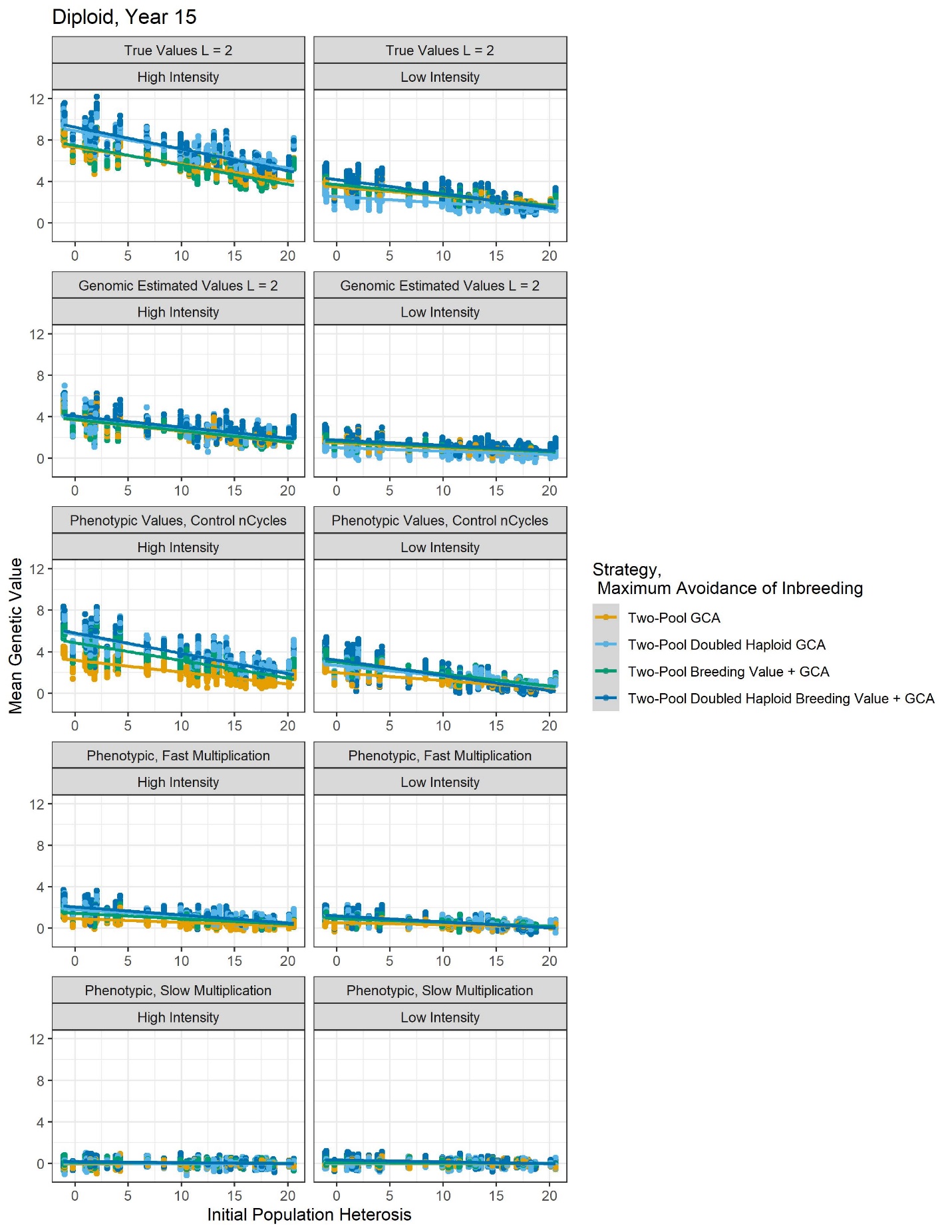

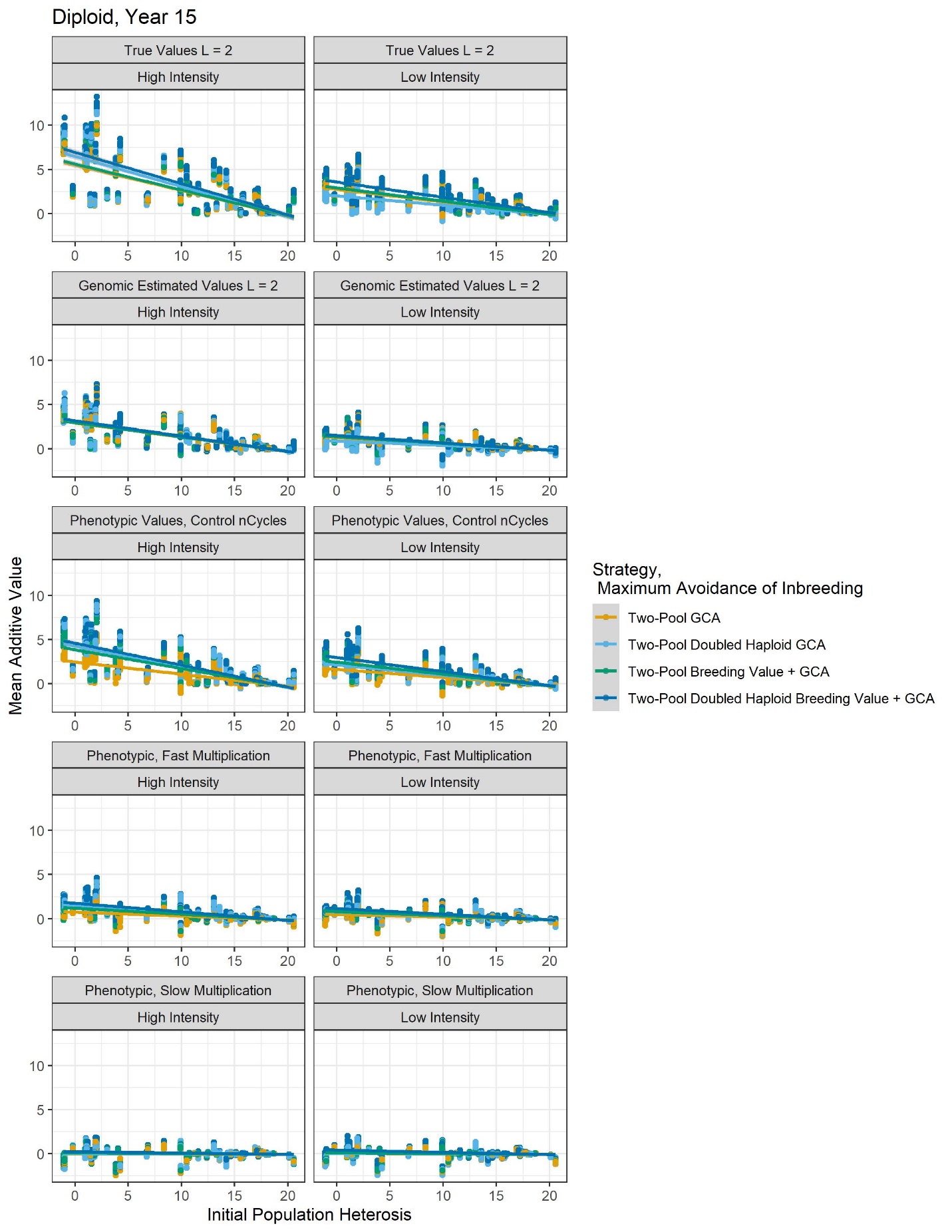

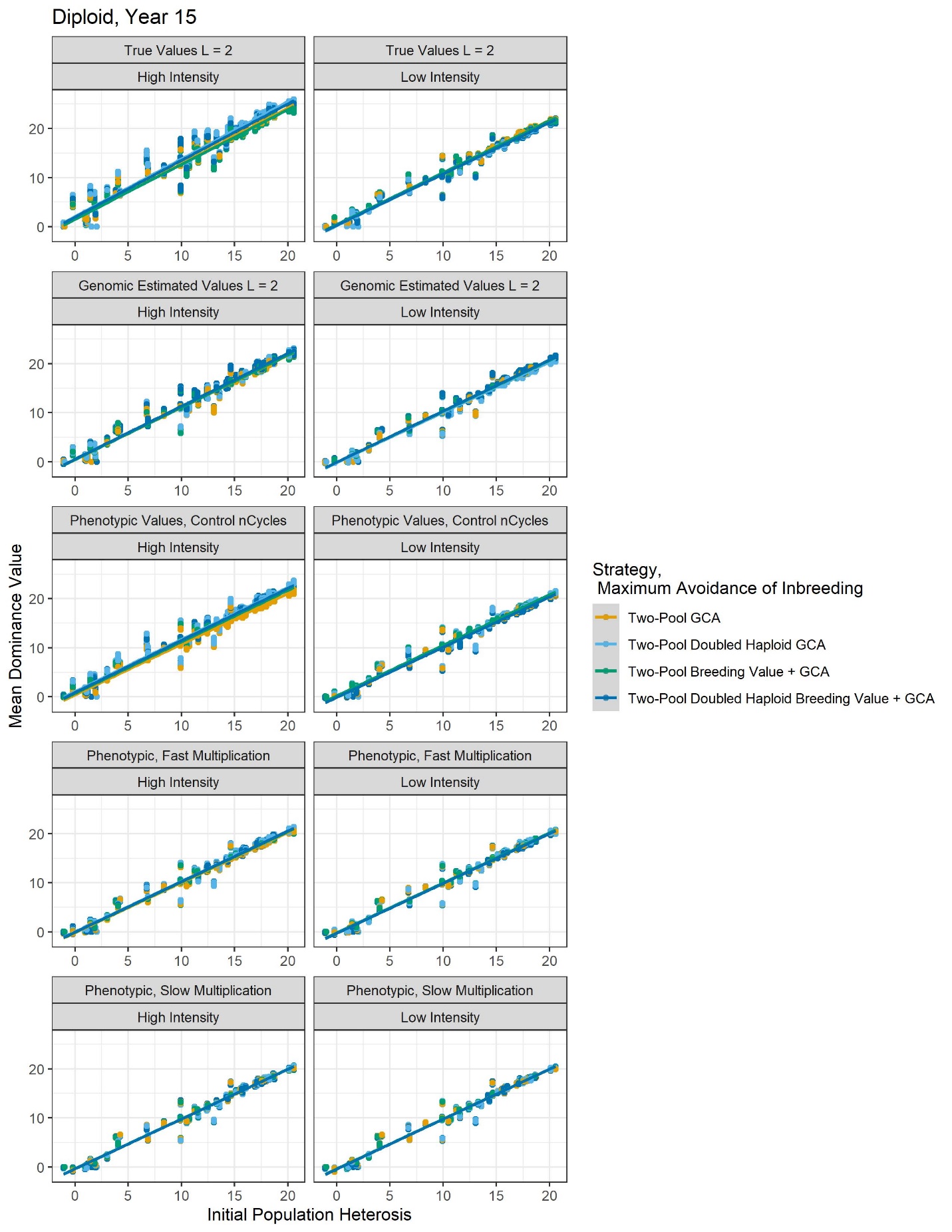

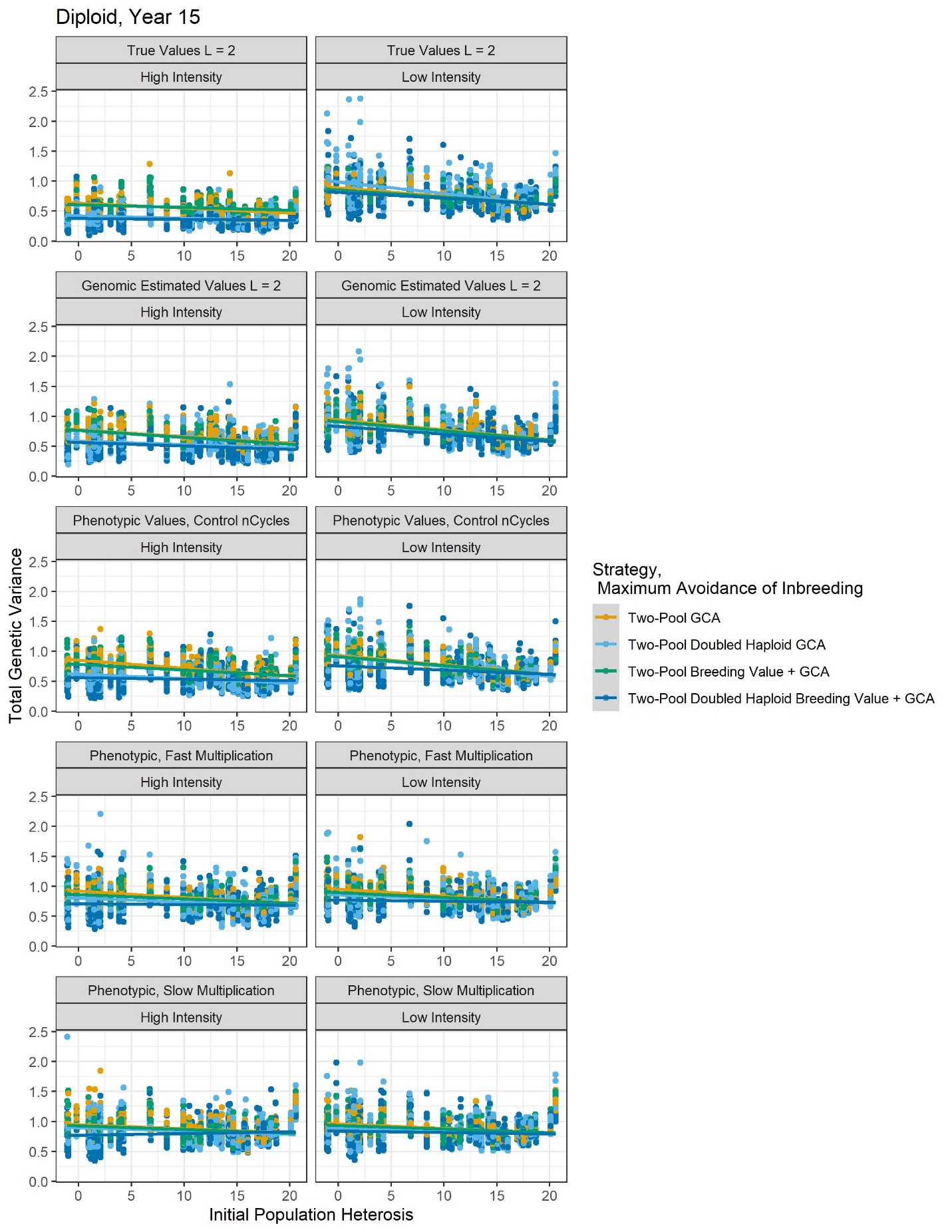

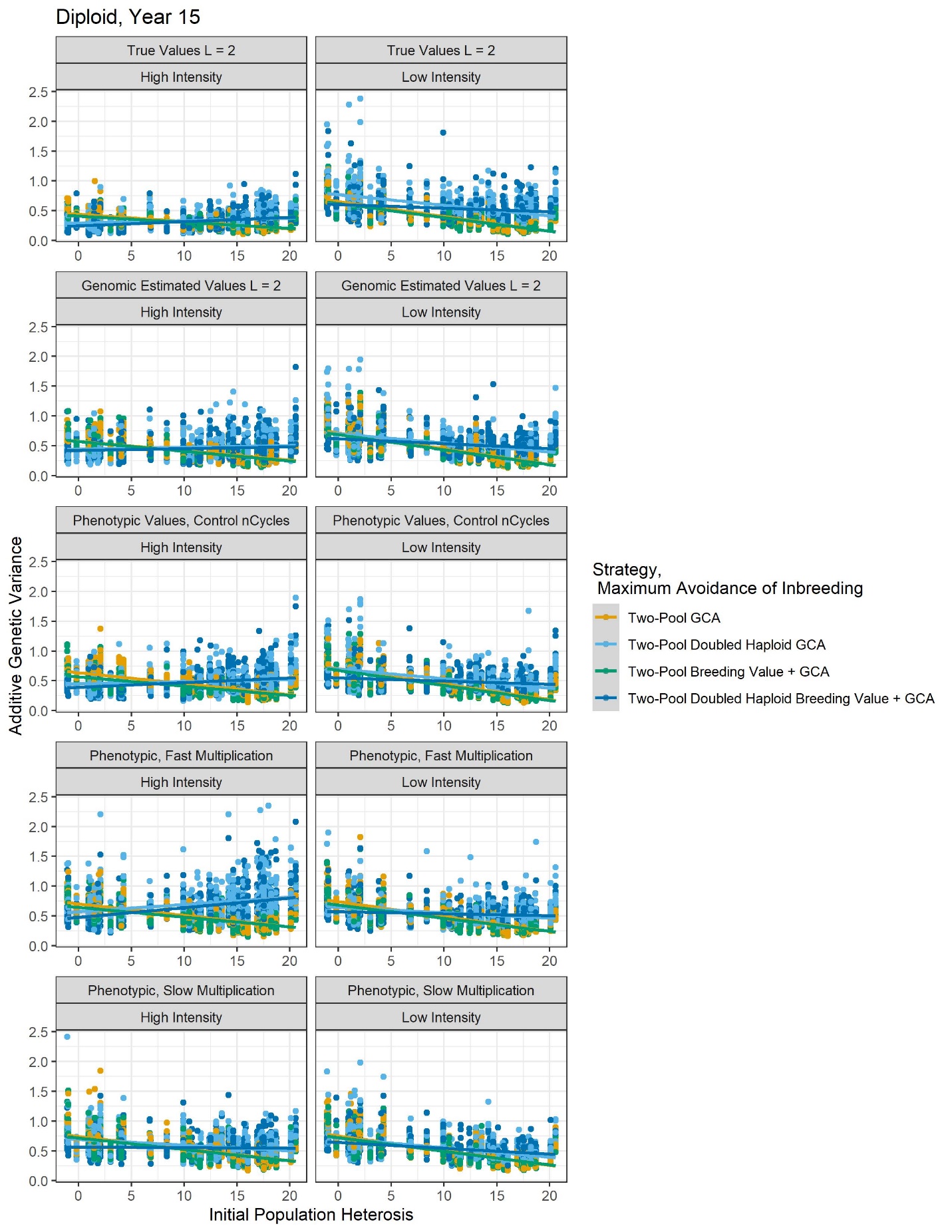

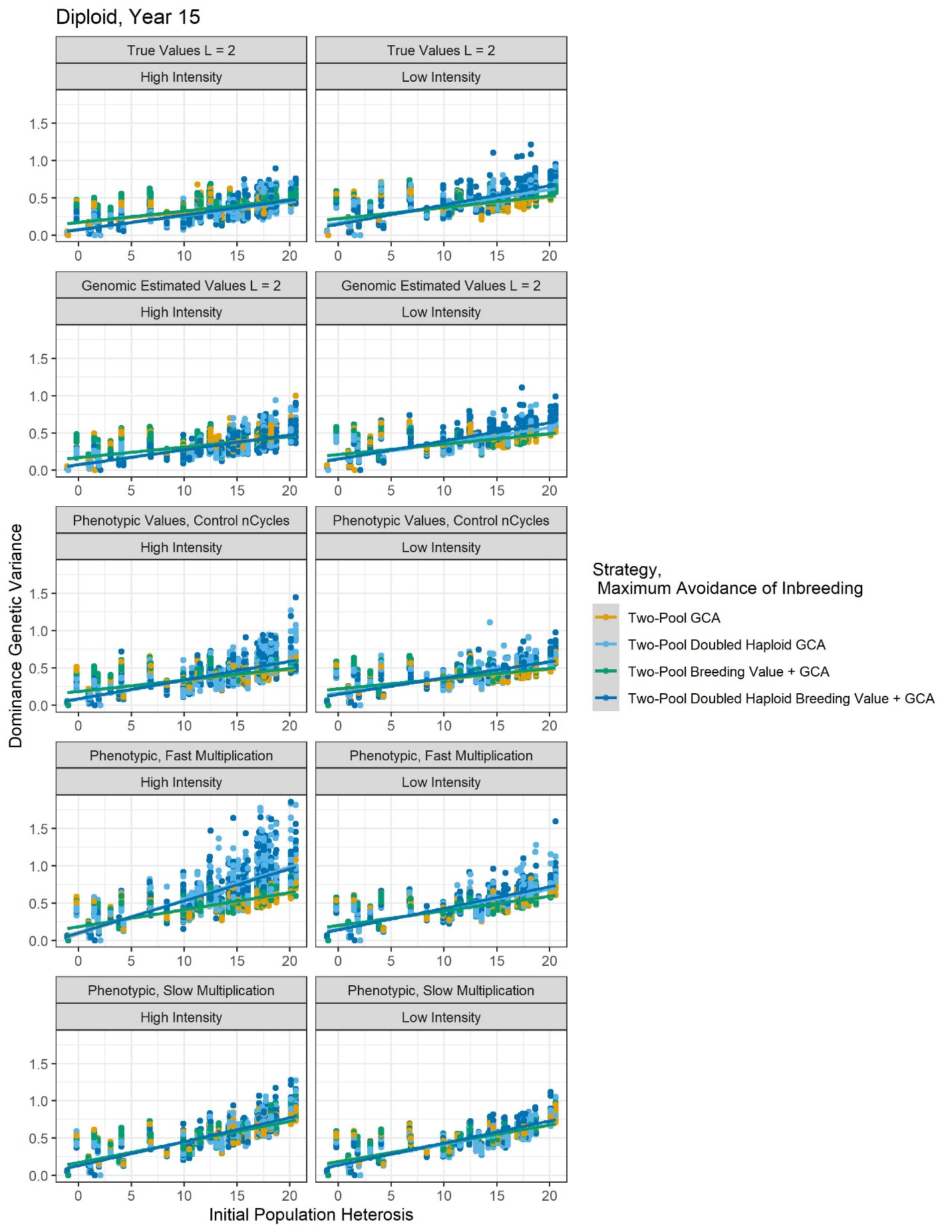

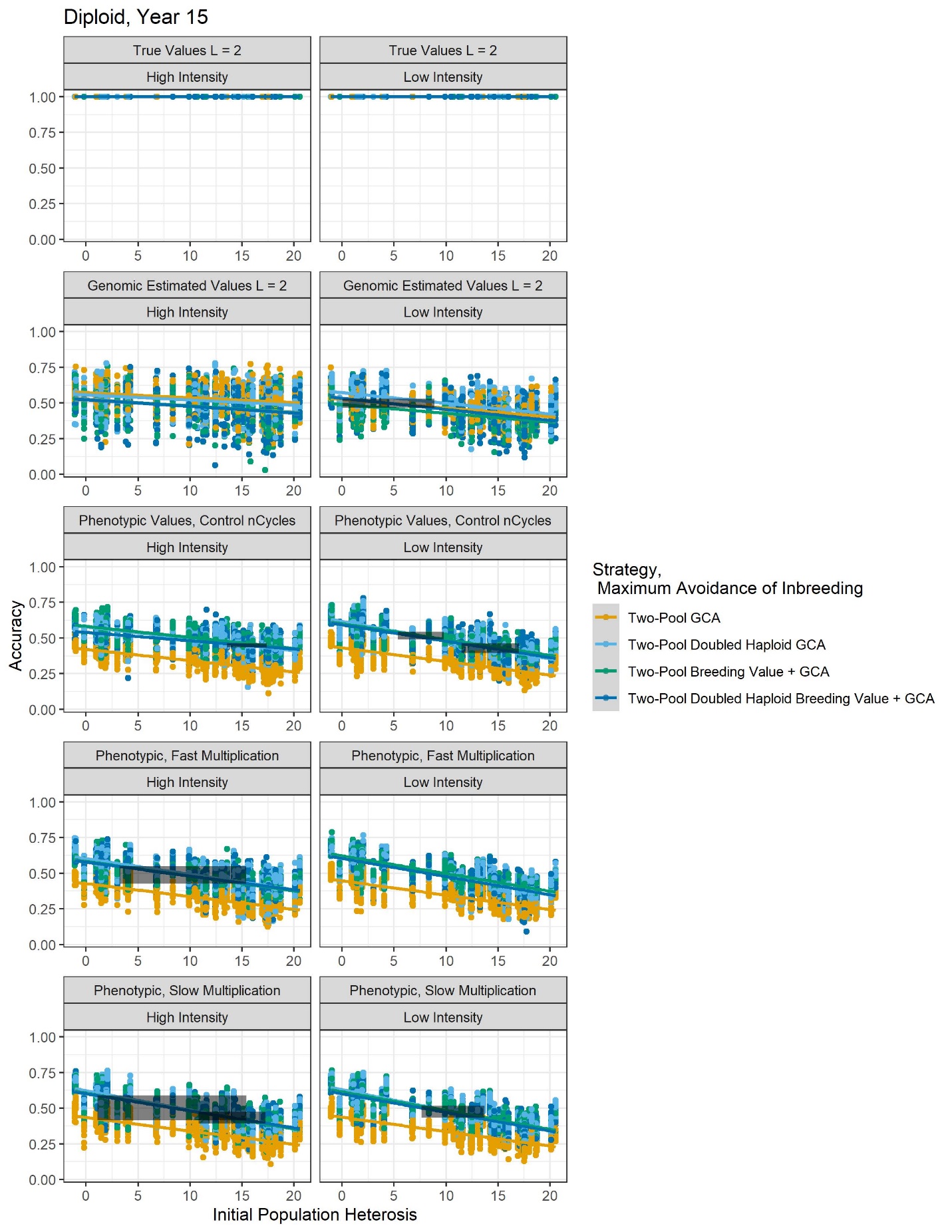

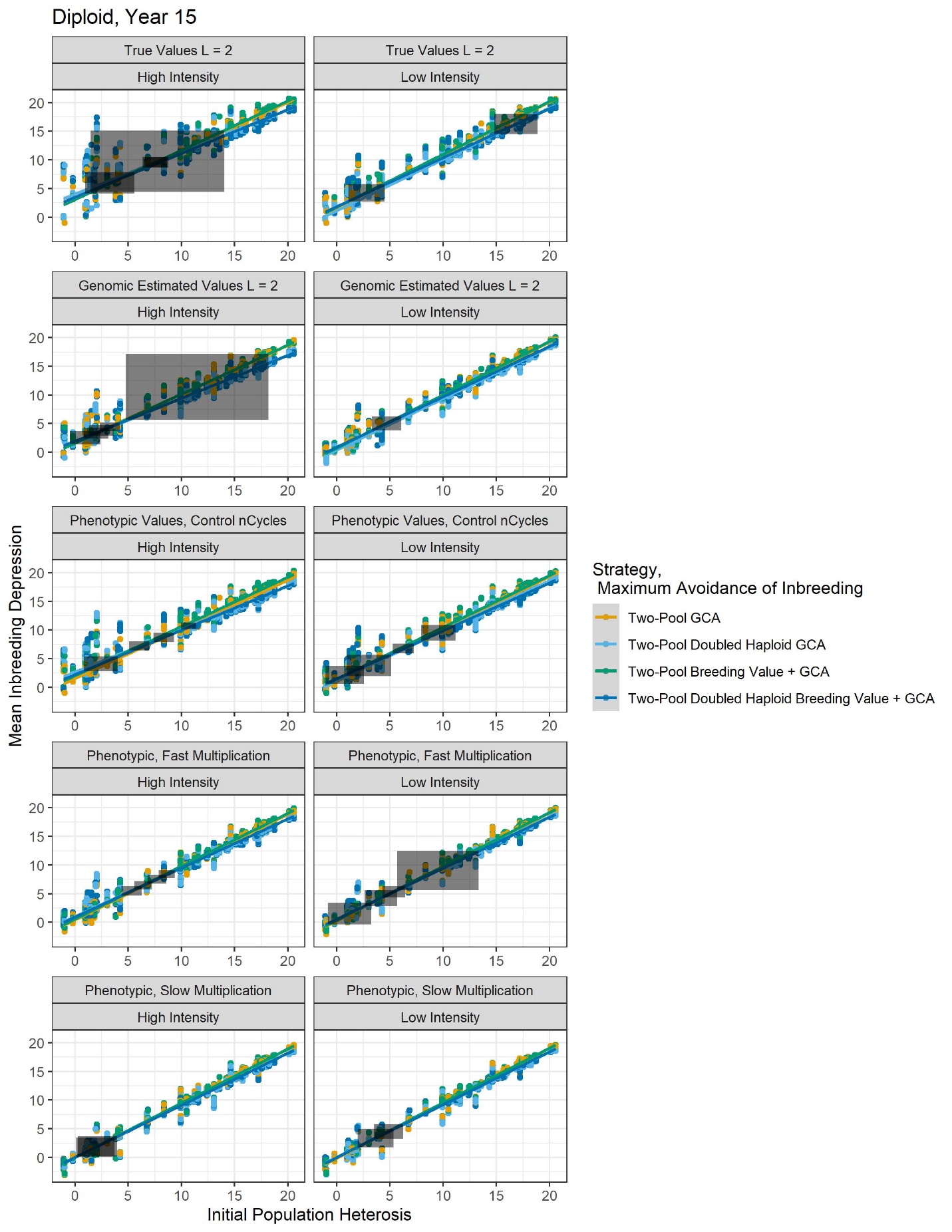

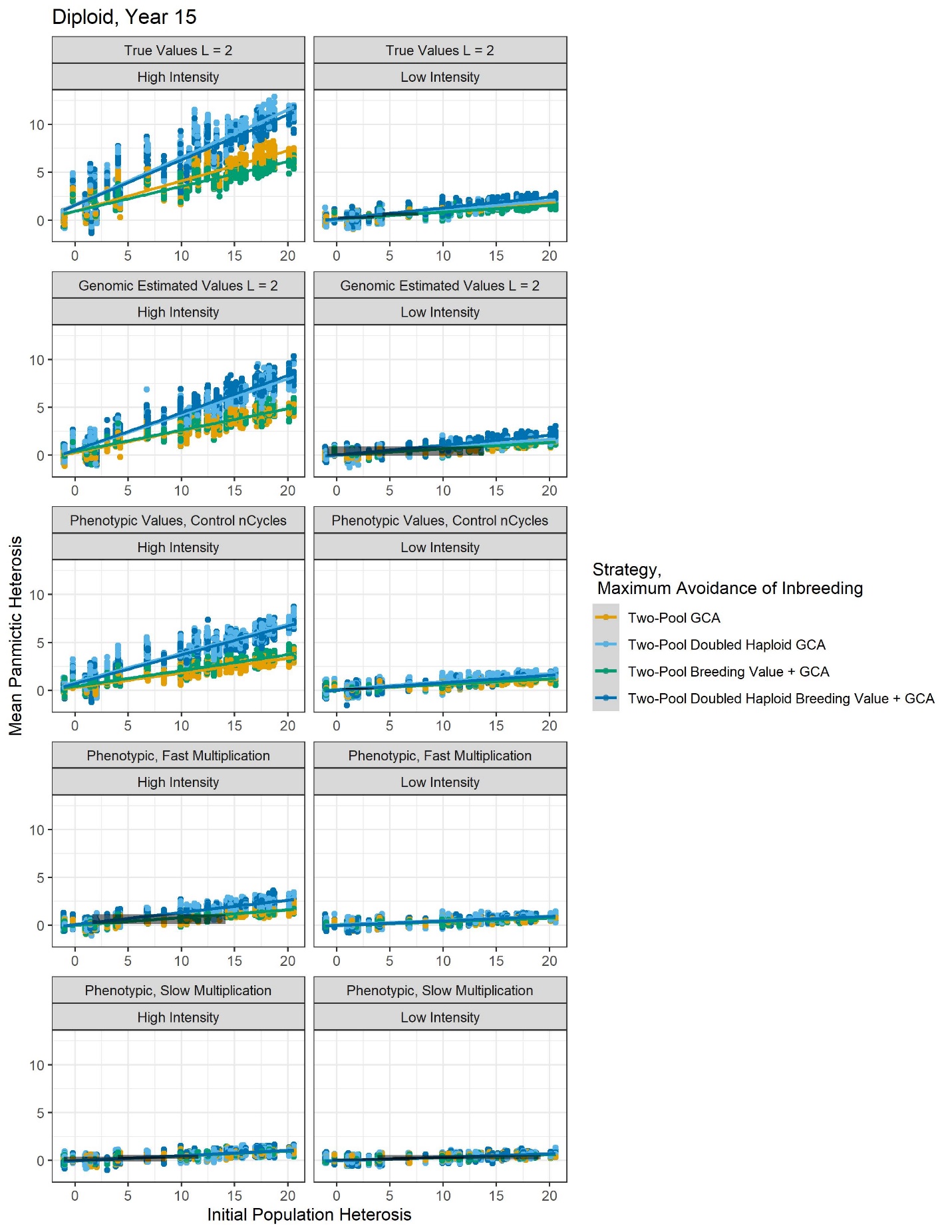

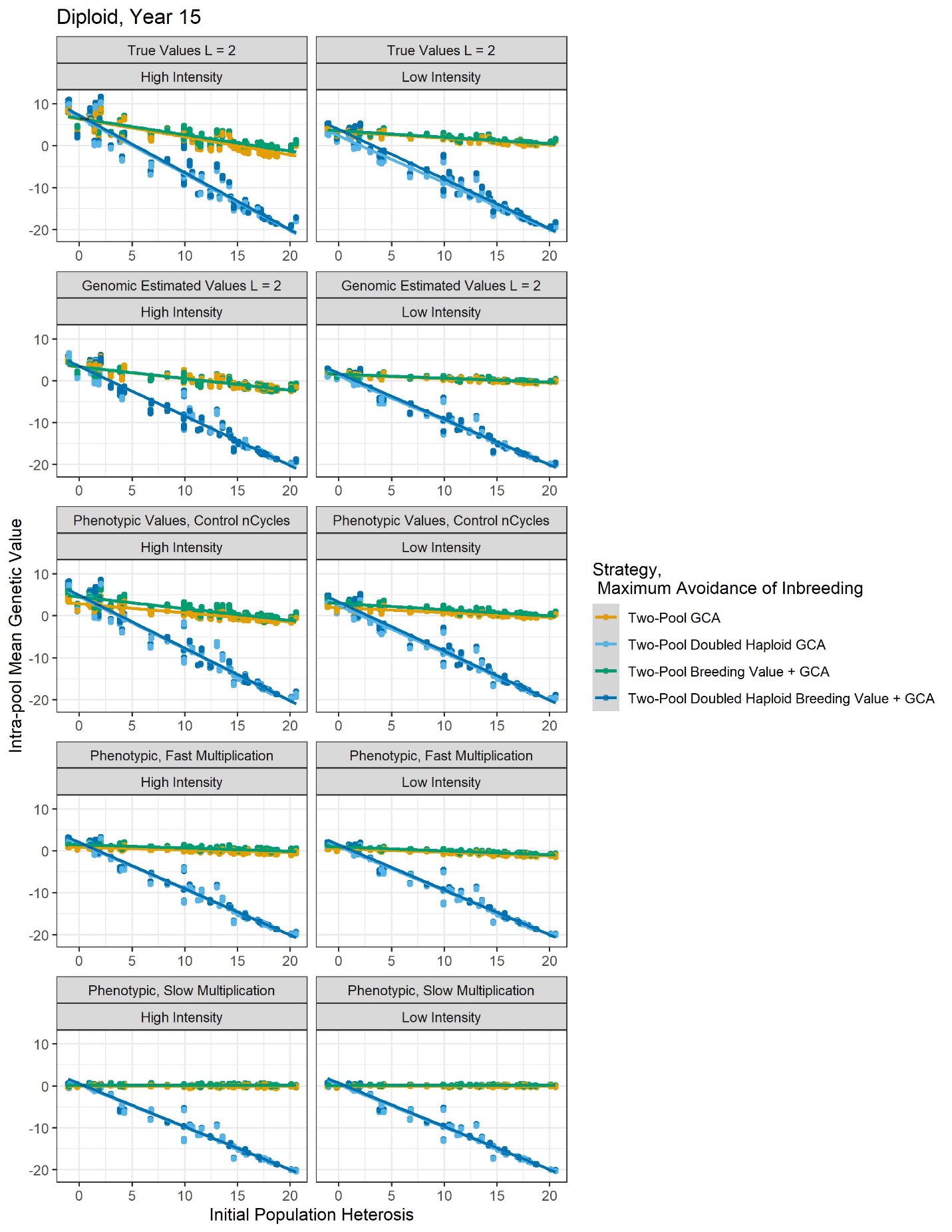

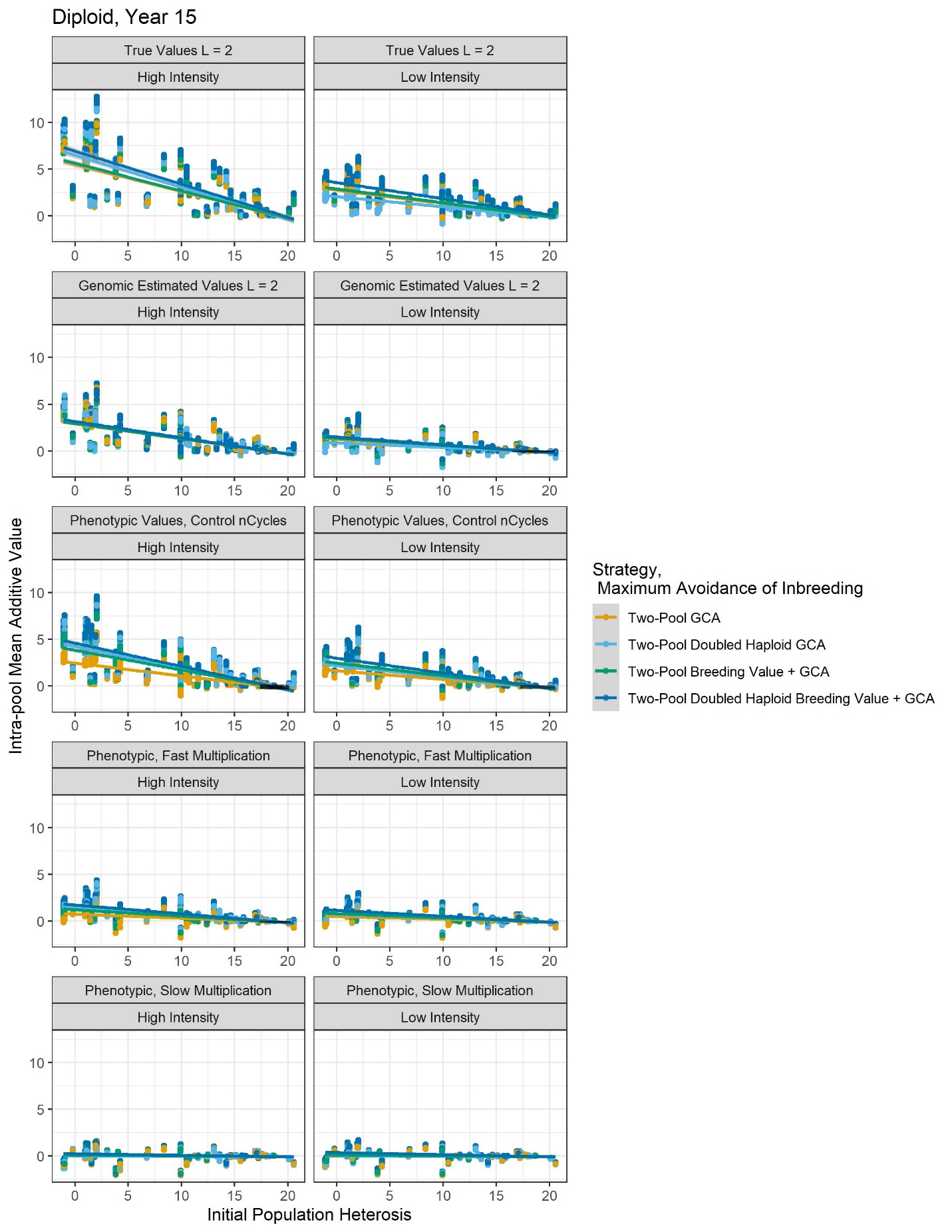

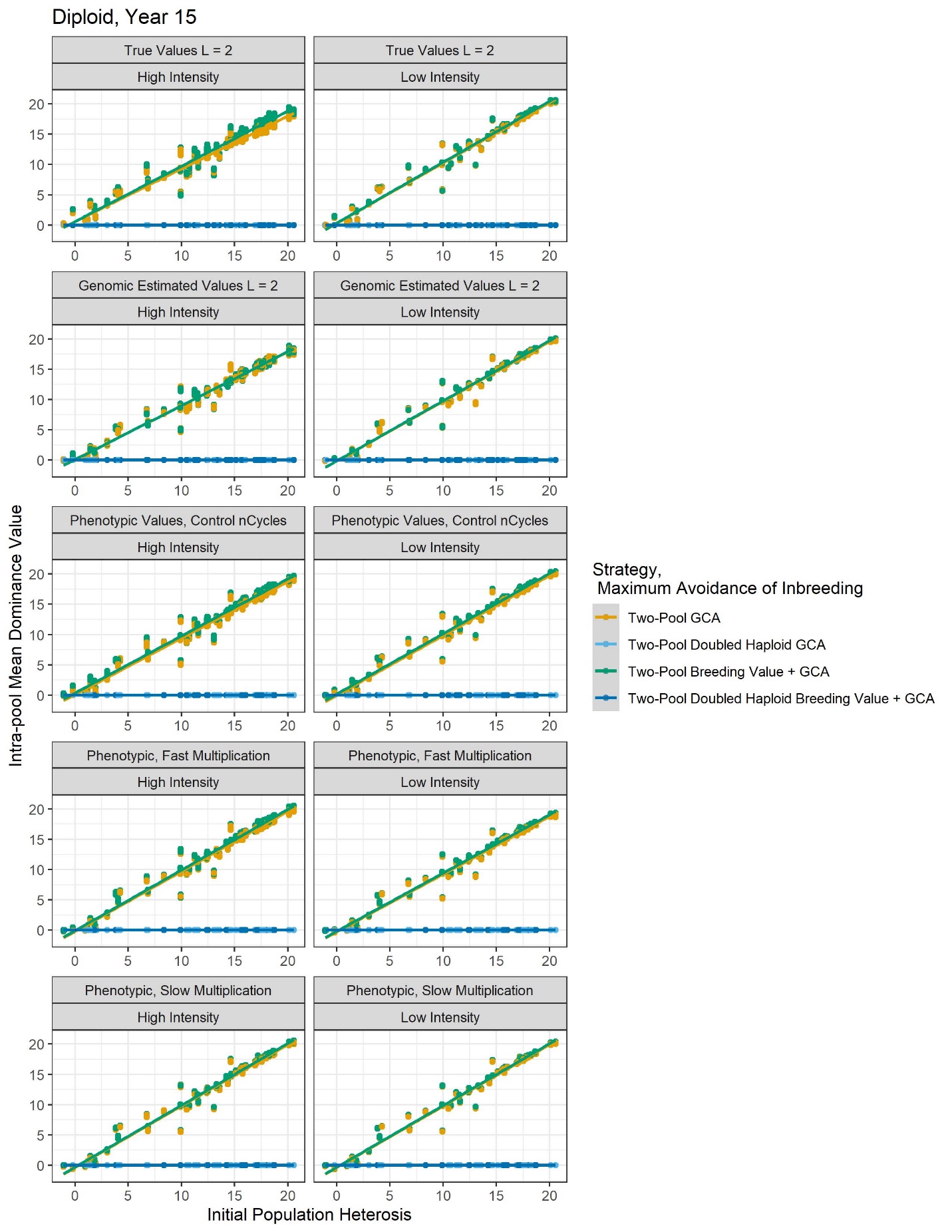

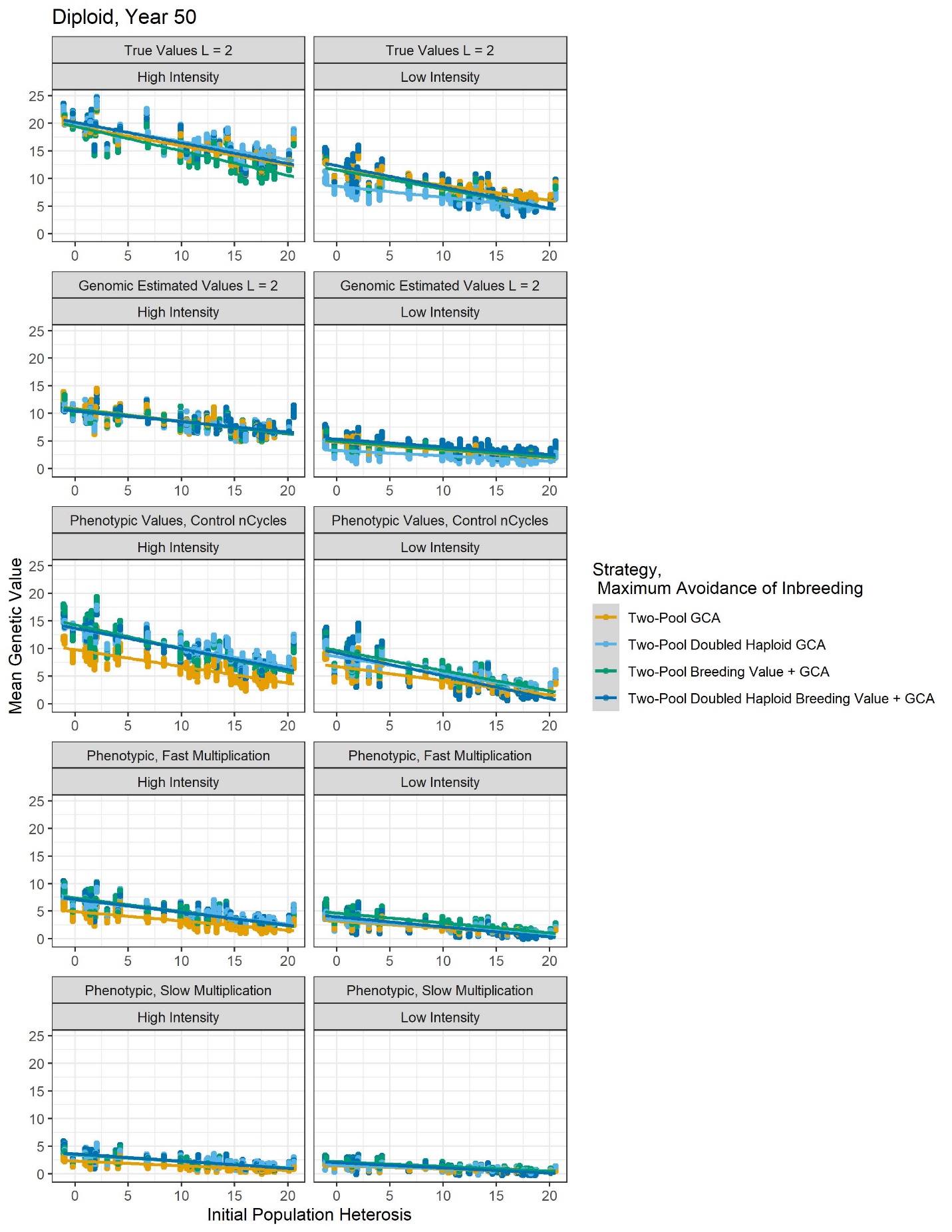

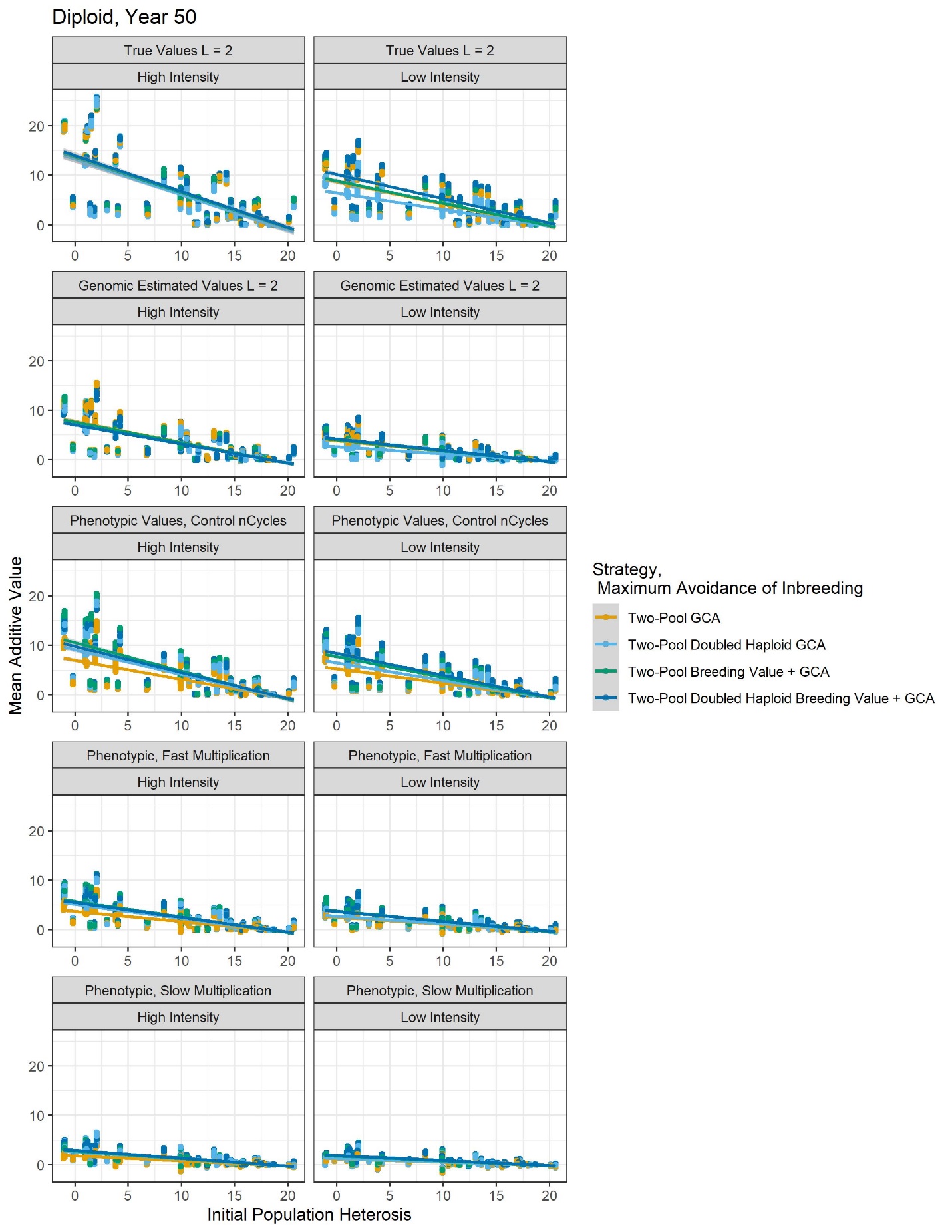

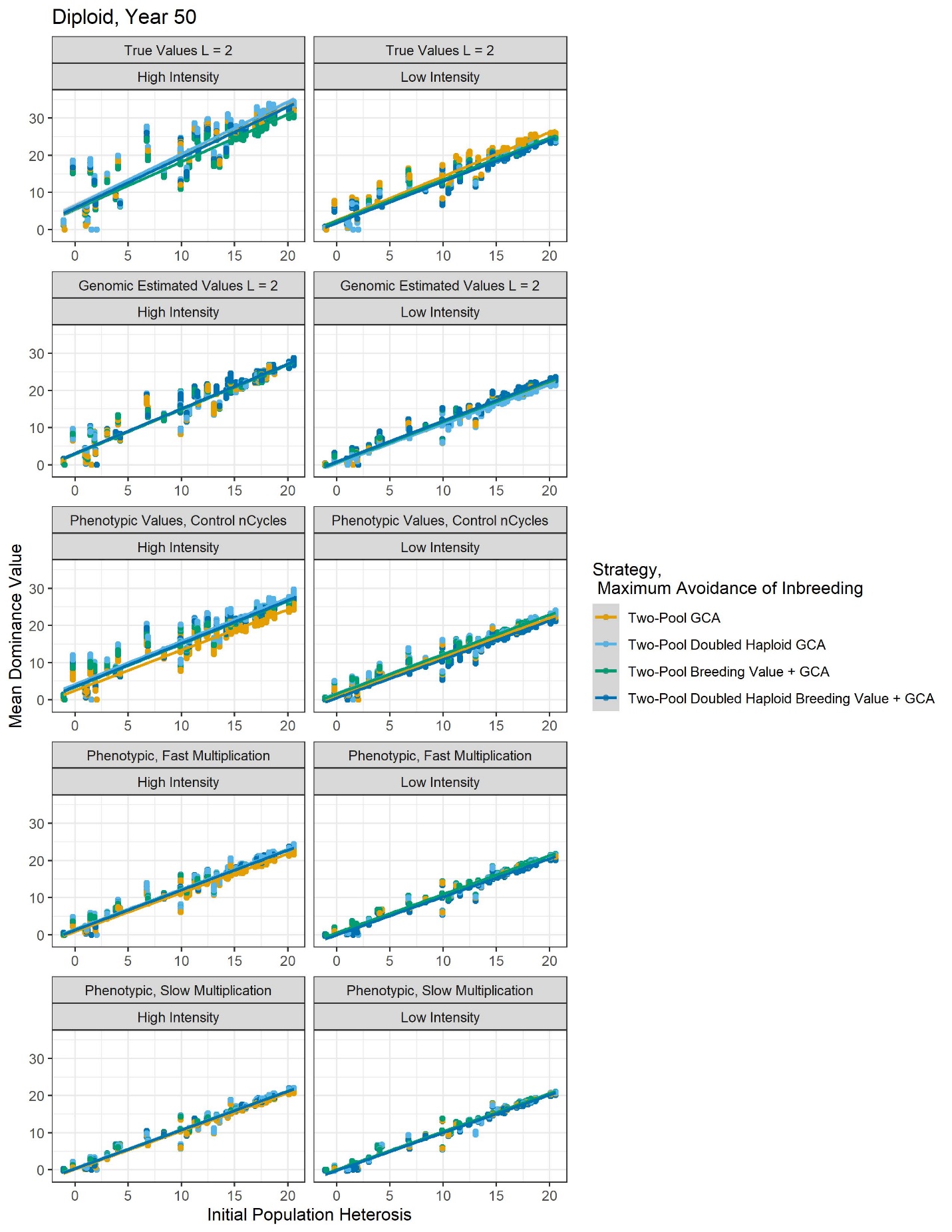

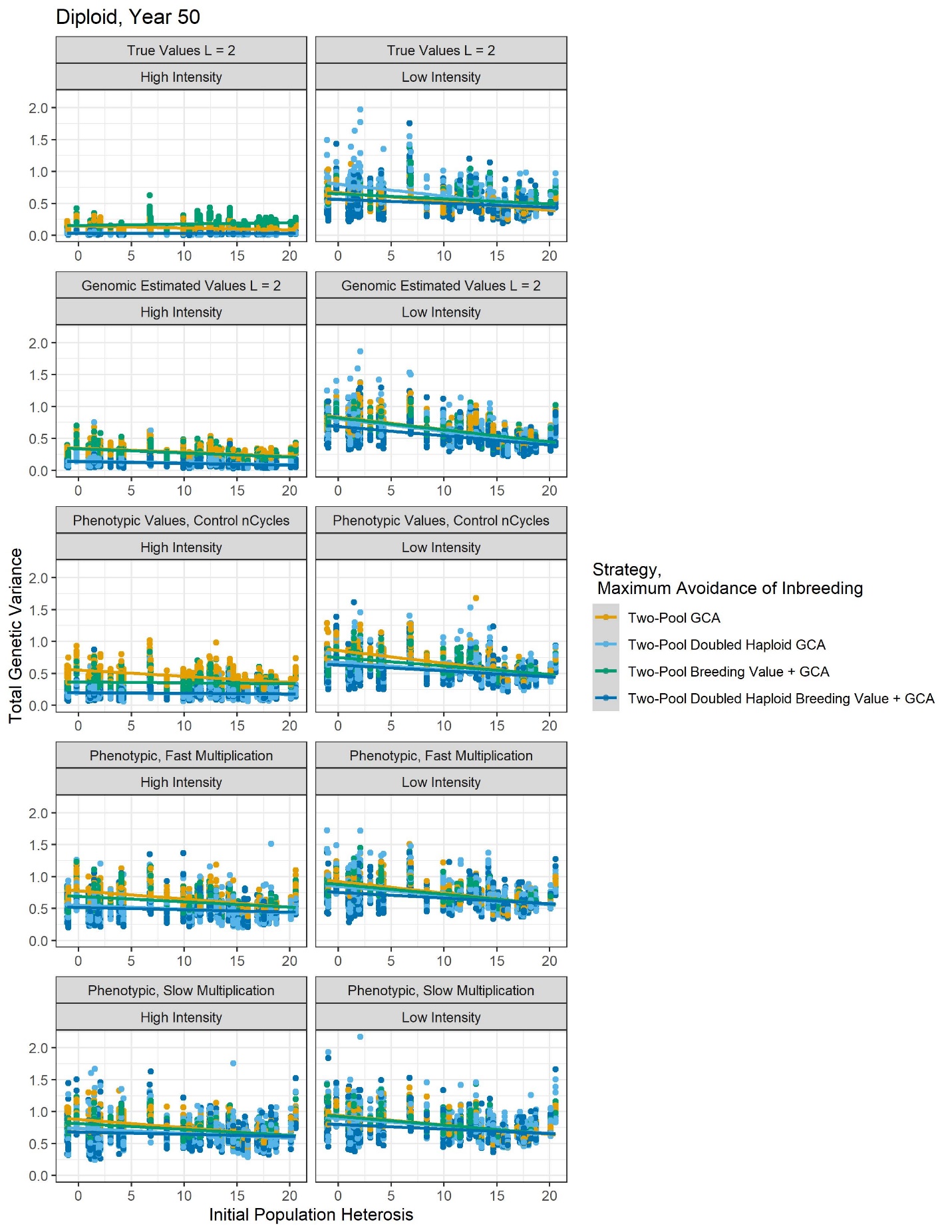

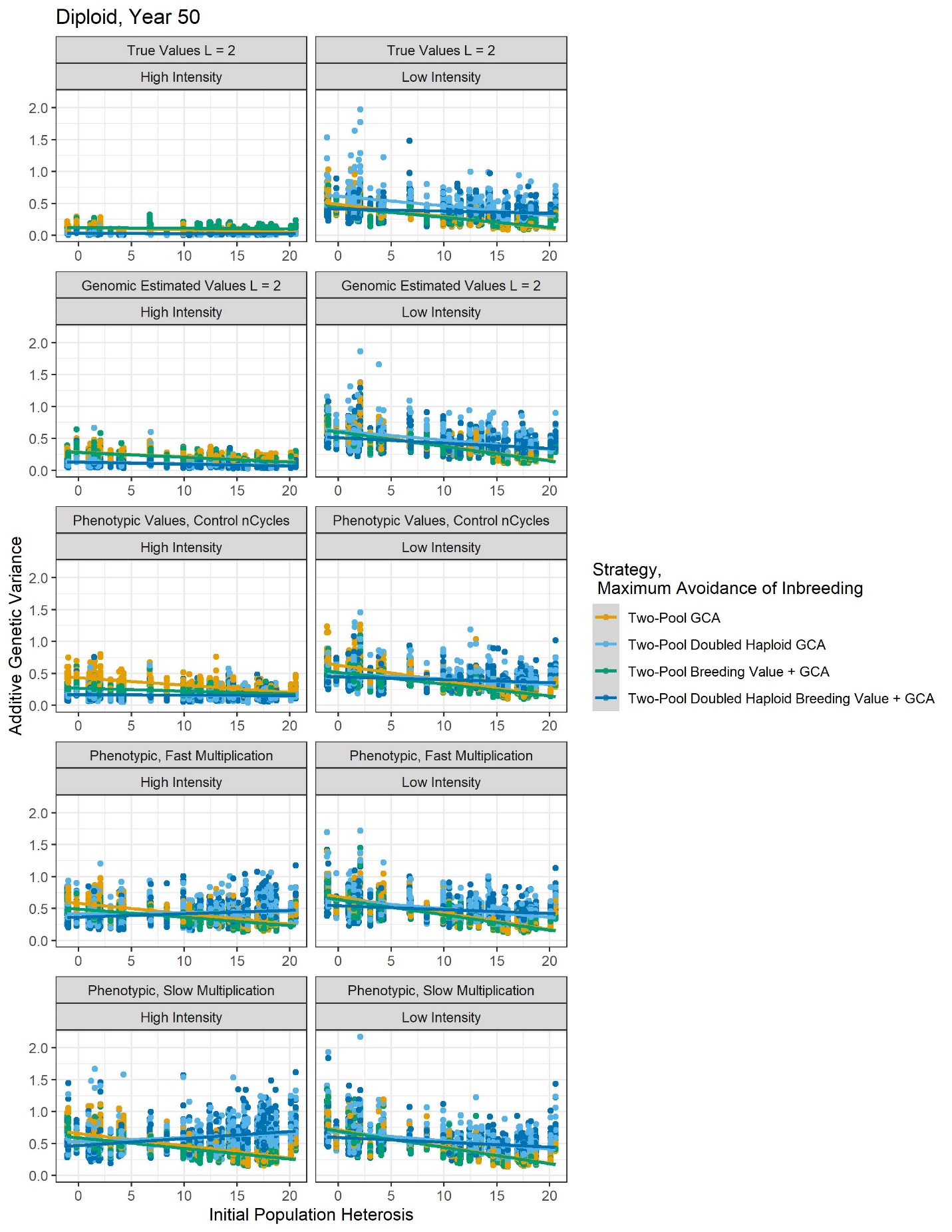

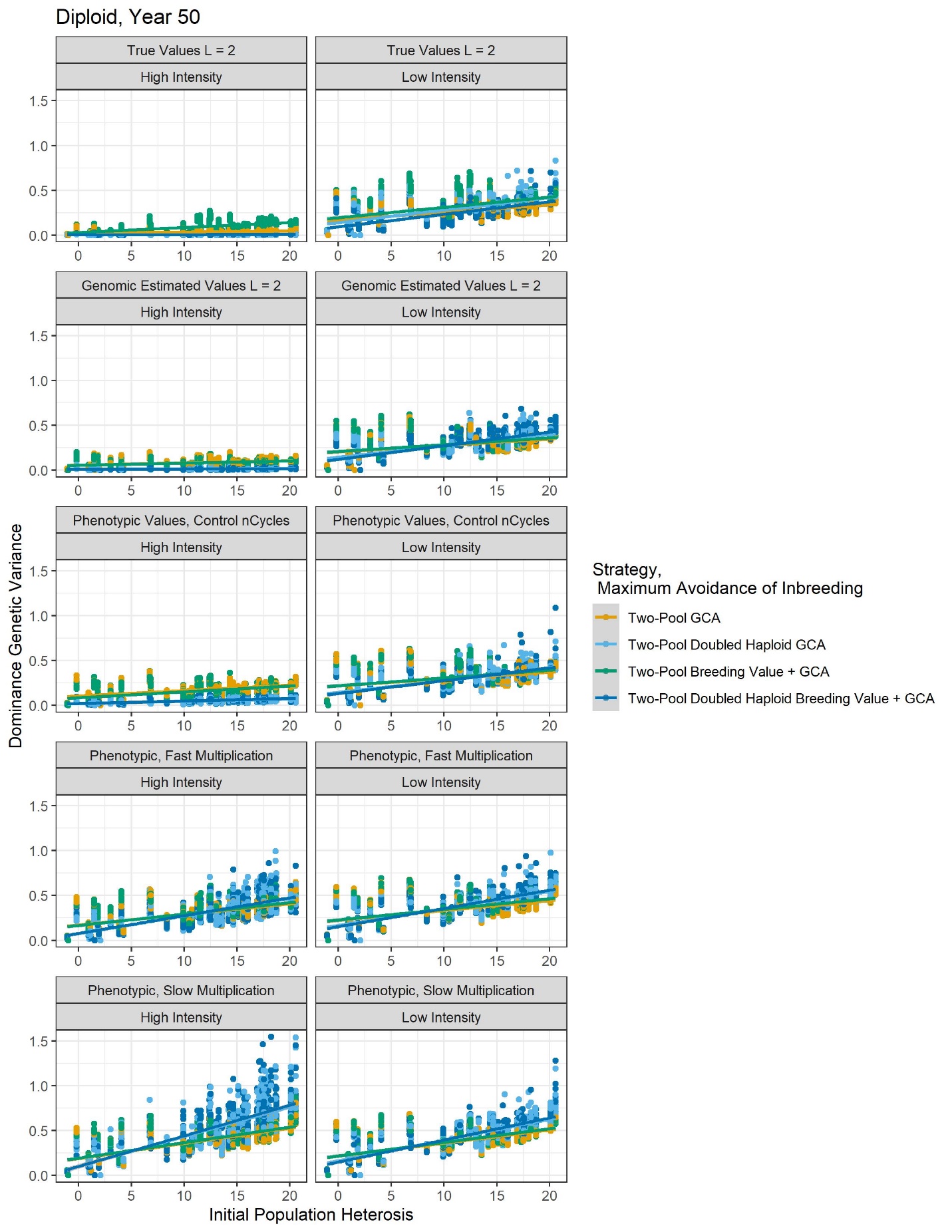

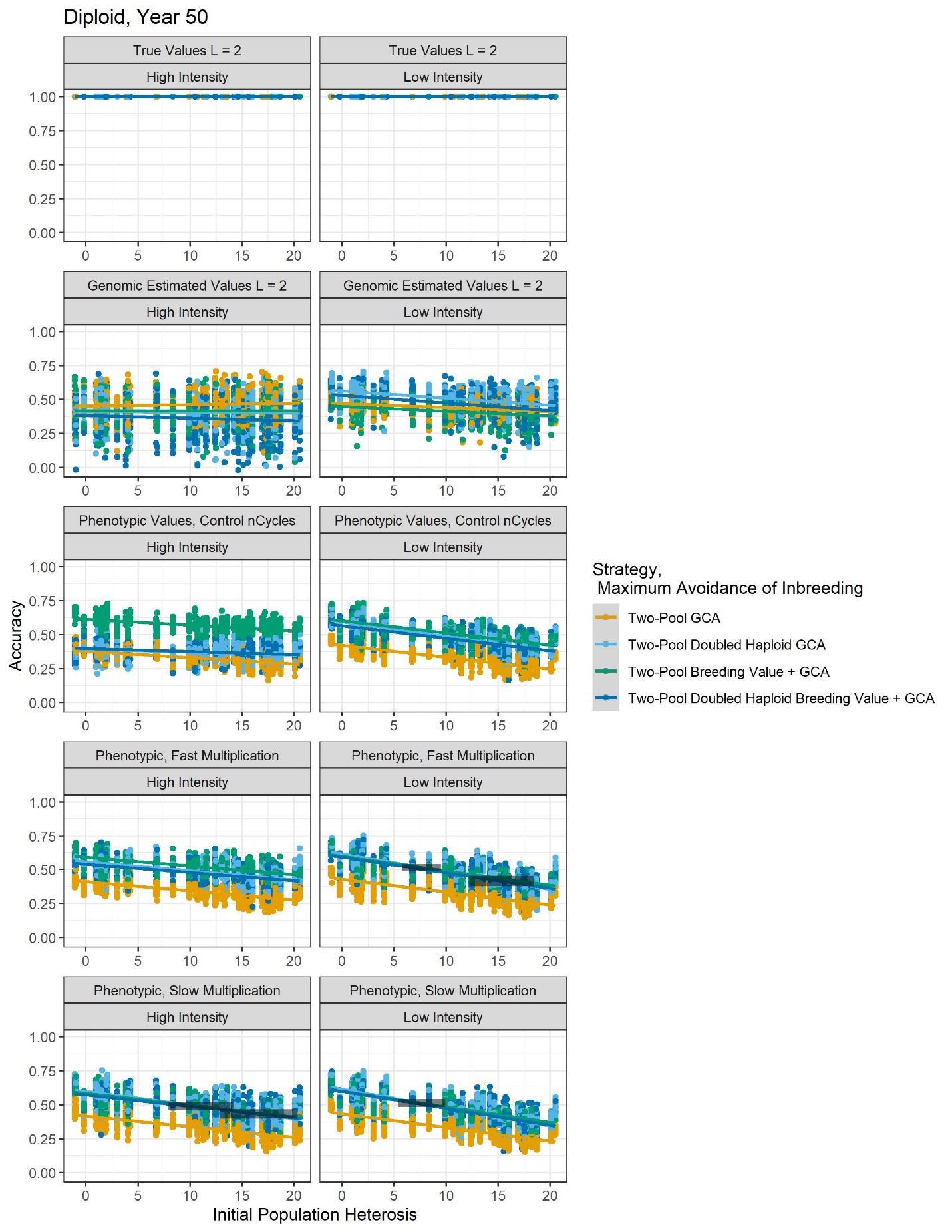

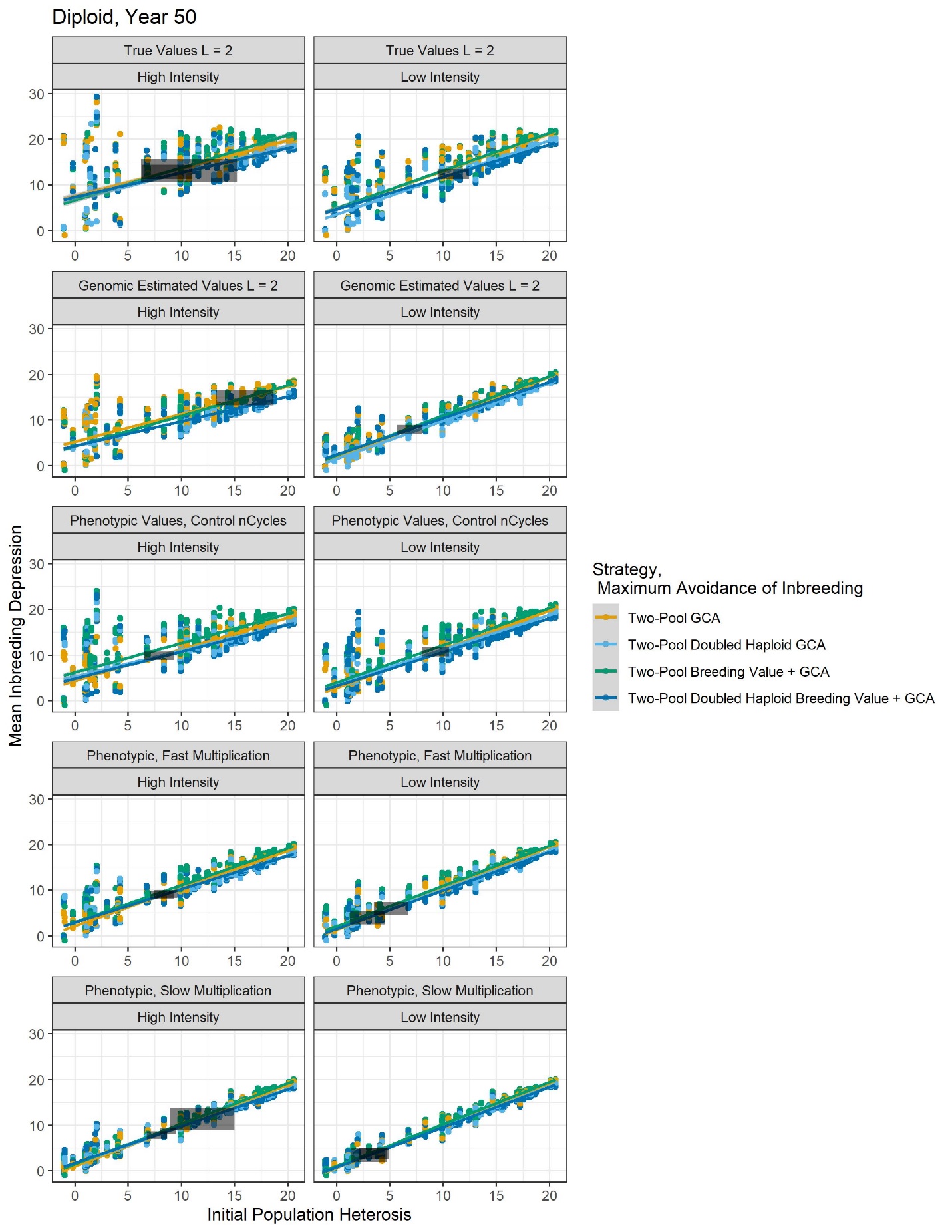

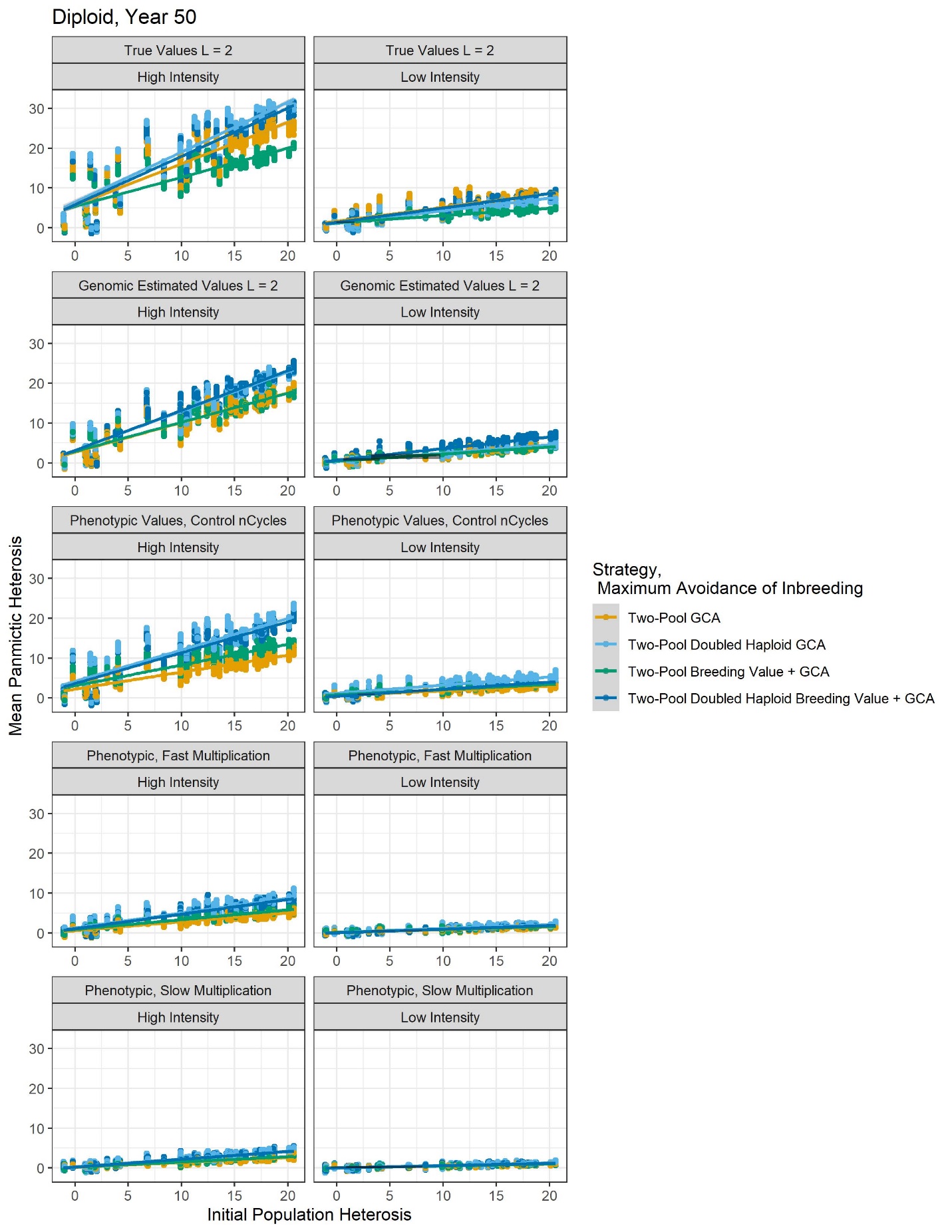

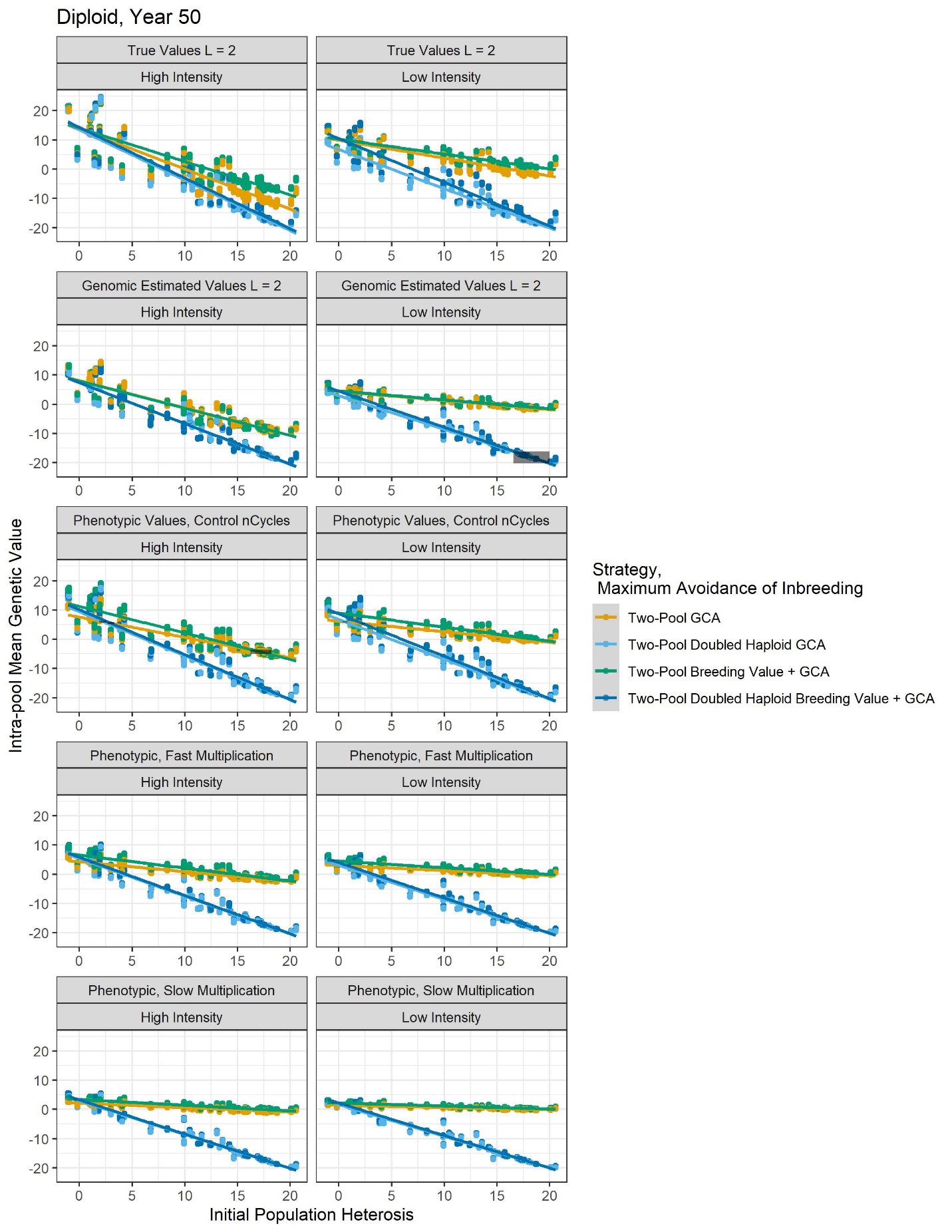

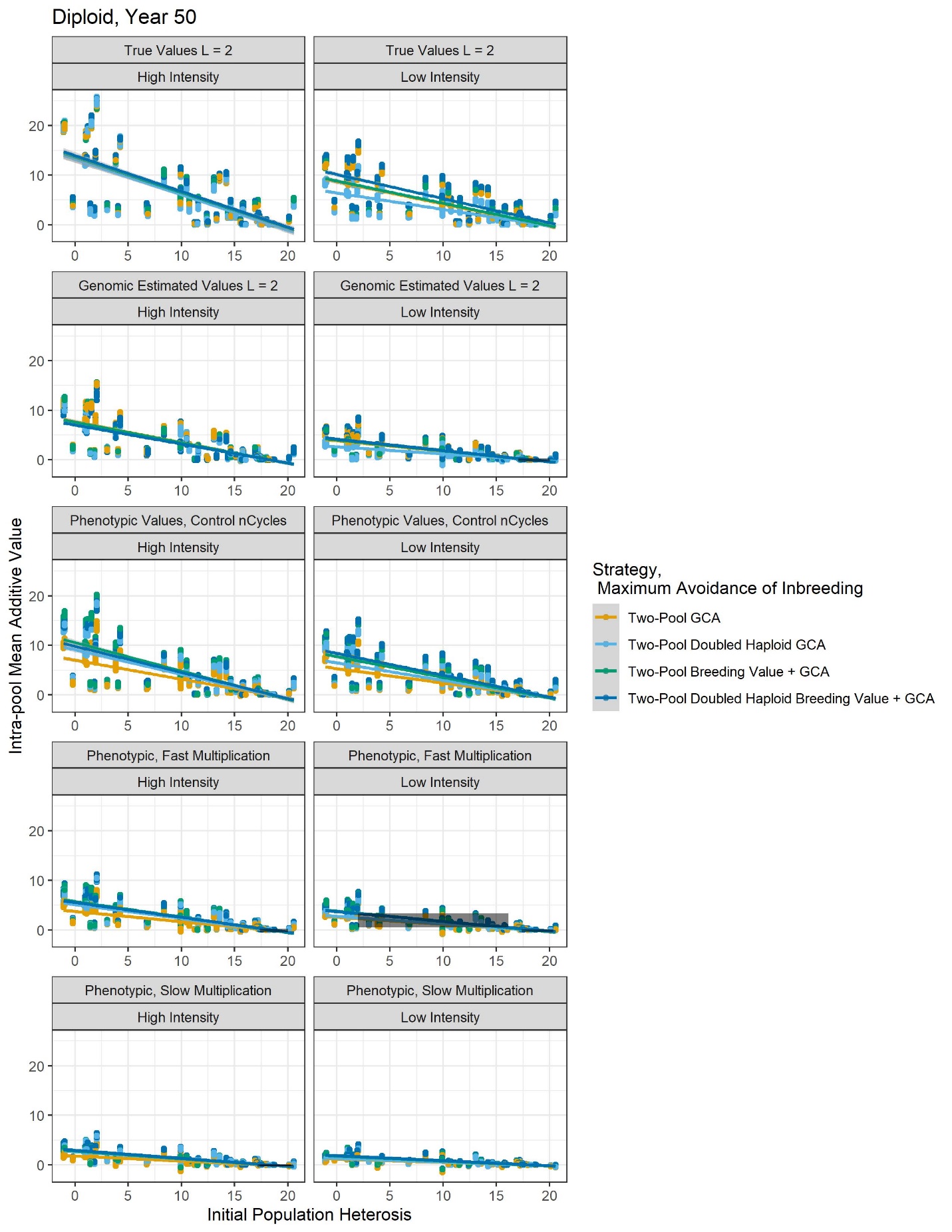

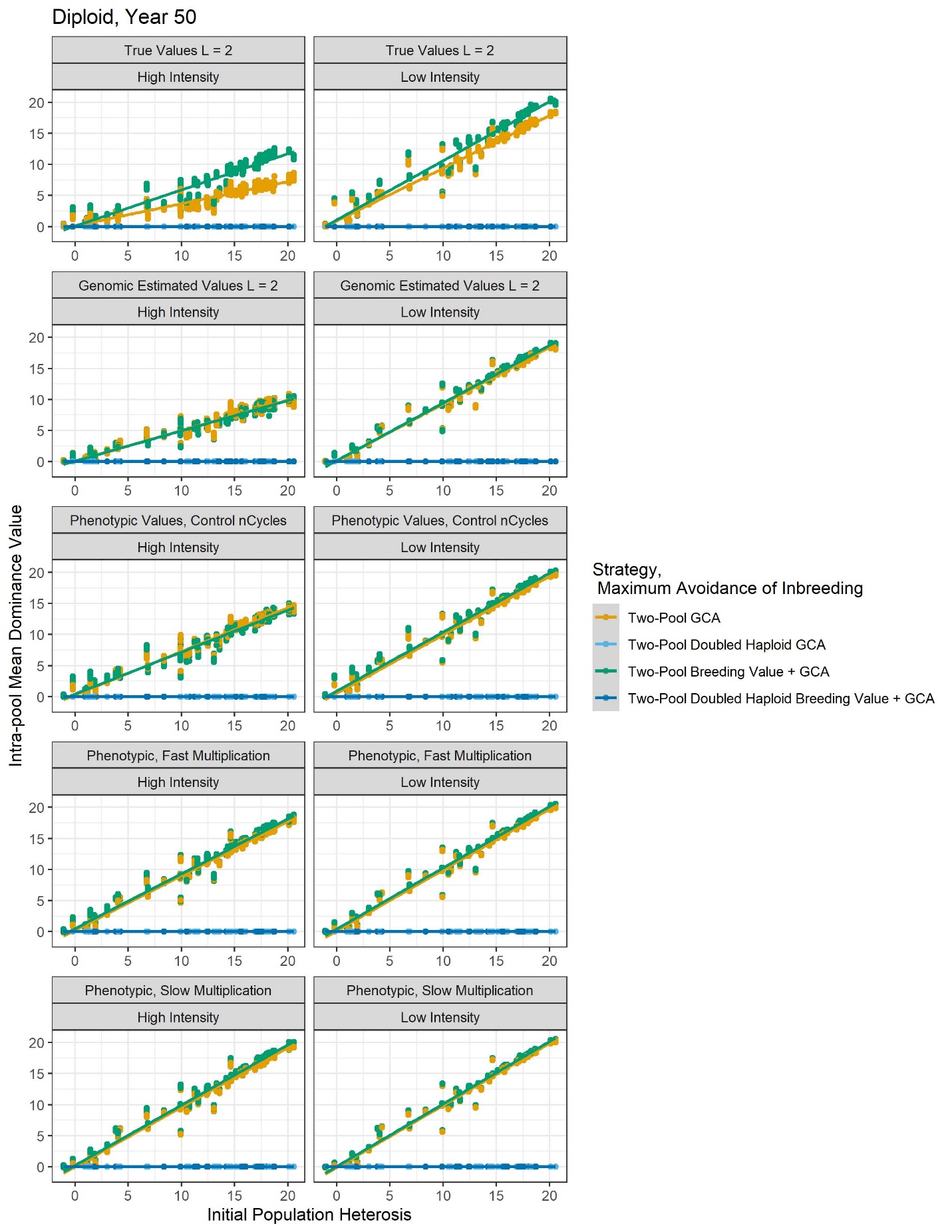

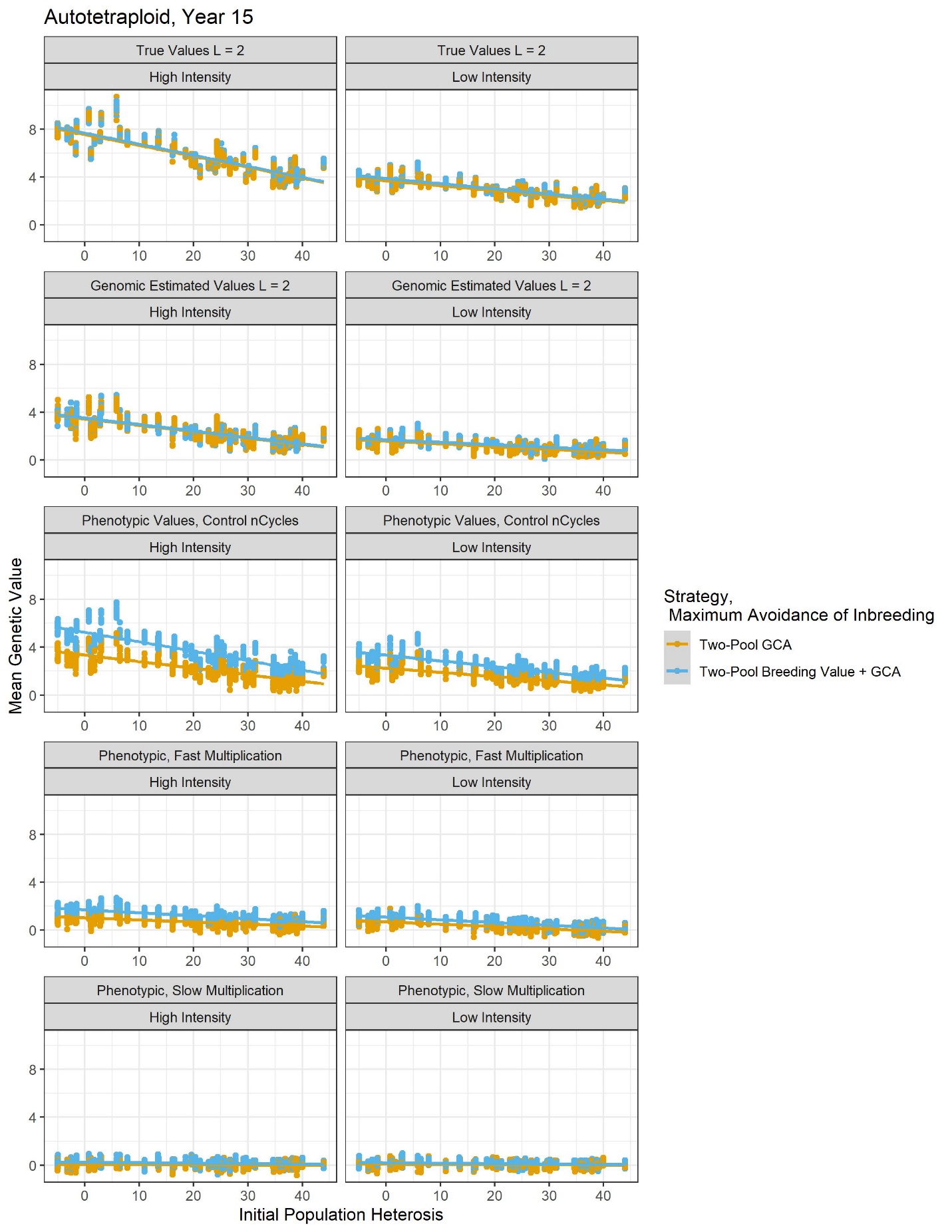

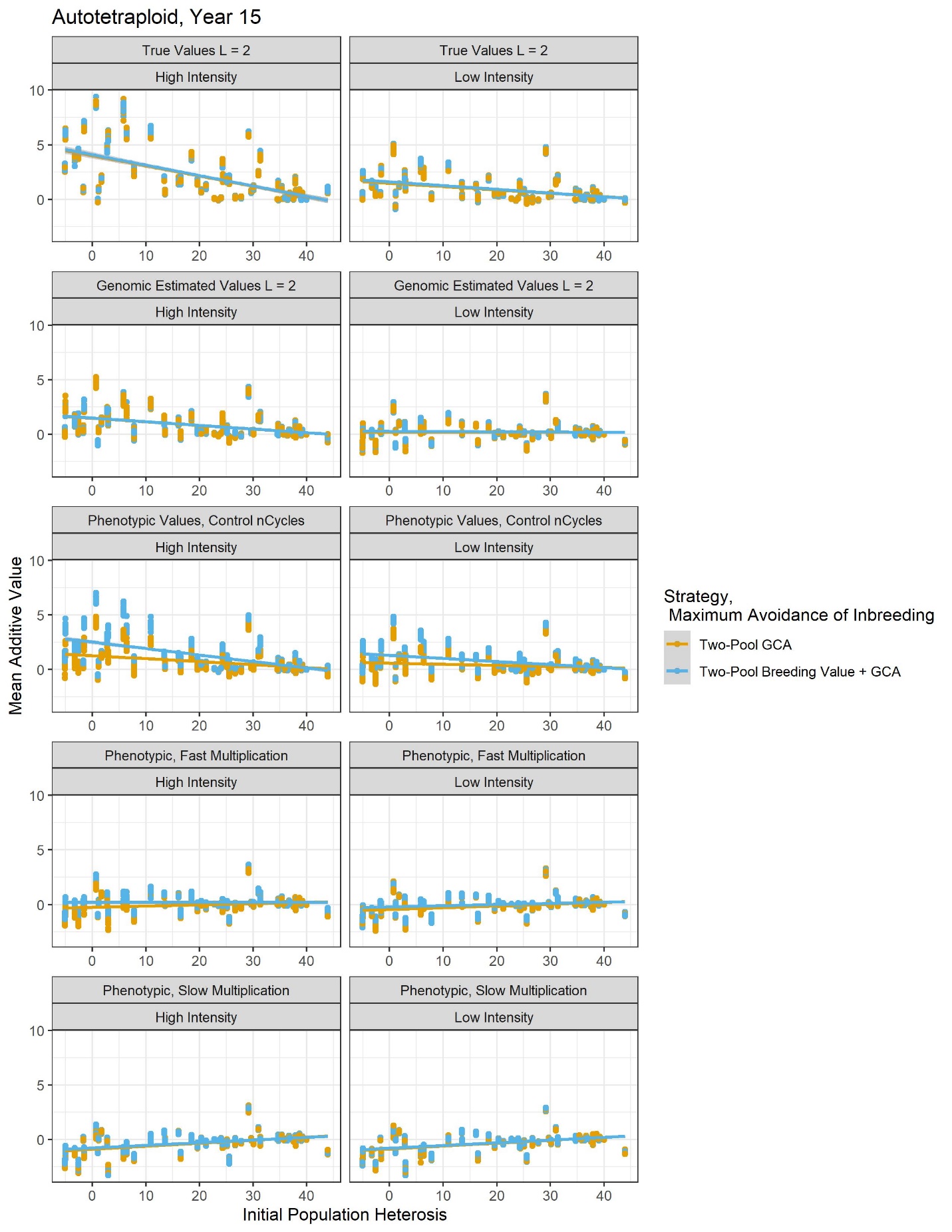

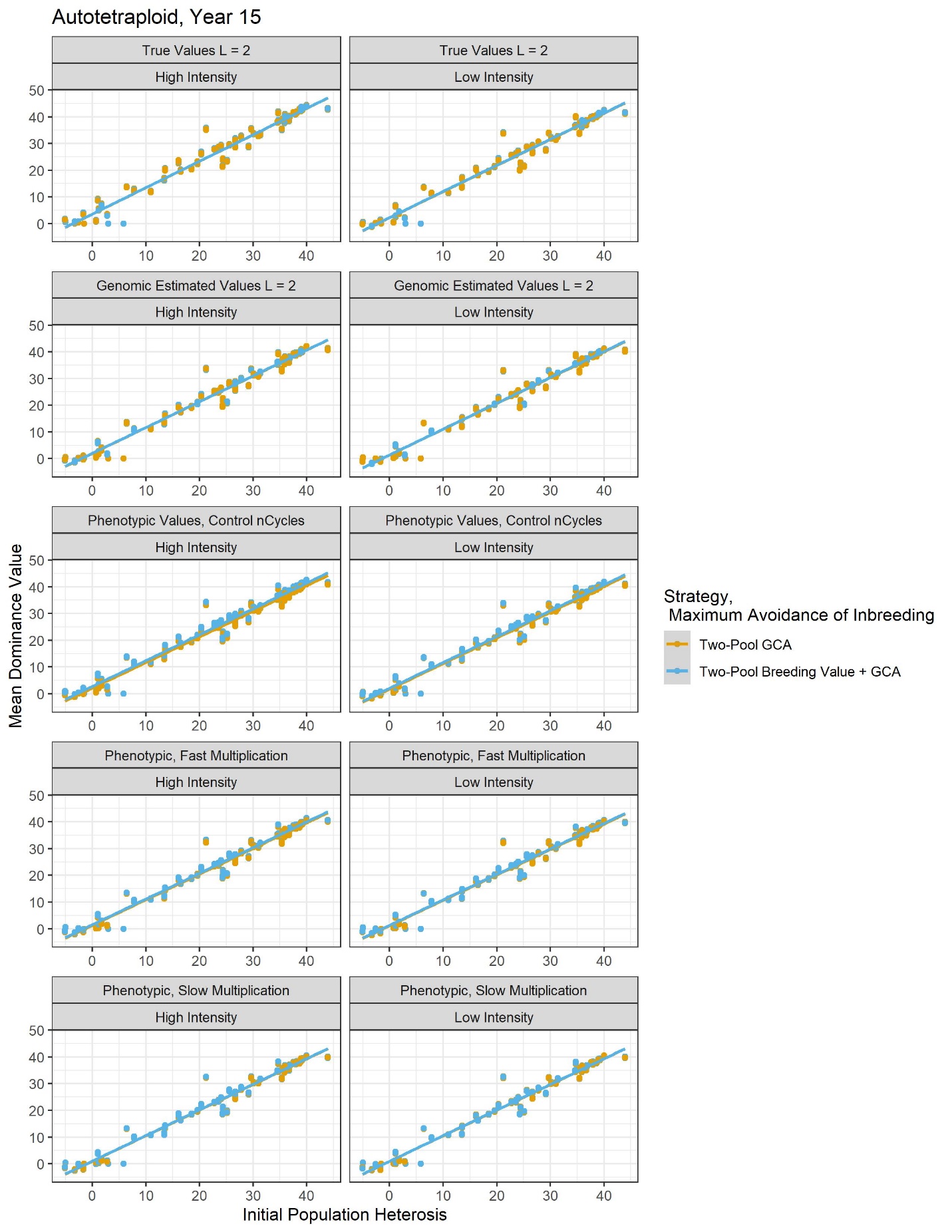

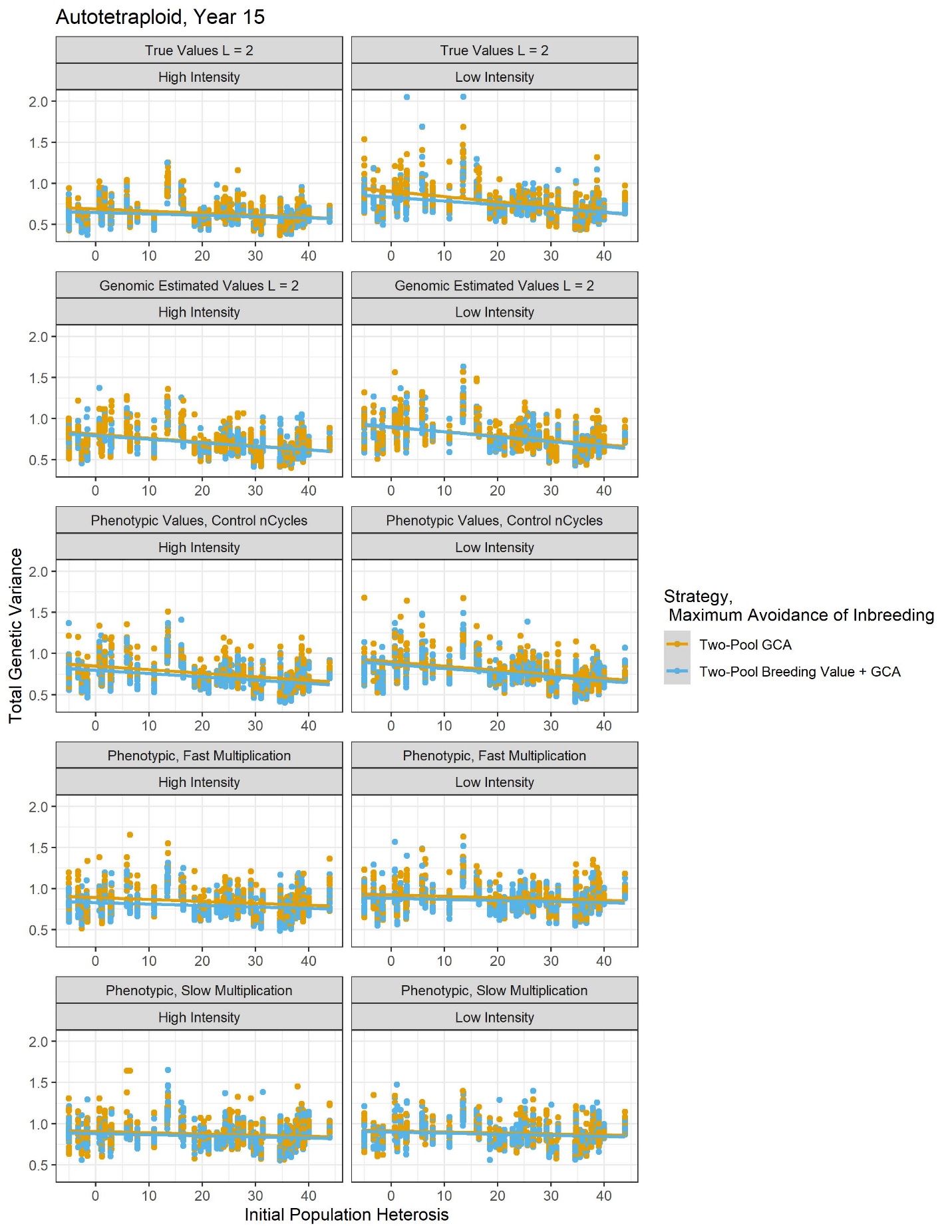
