## Supplemental File 45 for "Clonal breeding strategies to harness heterosis: insights from stochastic simulation"

**Supplemental Discussion**

*Alternatives to Two-Pool GCA for diploids*

Transitioning to Two-Pool GCA can represent a significant logistical challenge for breeding programs, and species may have radically different biological constraints or costs among strategies than those assumed in our study. We explored two other methods to exploit dominance, One-Pool Cross Performance and Two-Pool Breeding Value, which may be useful to programs for which Two-Pool GCA is not feasible. With PS at high intensity, Two-Pool Breeding Value tended to outperform Two-Pool GCA at high intensity with fast or slow multiplication, with Two-Pool GCA only providing benefits at highly positive H_0_ after 50 years. However, Two-Pool Breeding Value usually underperformed One-Pool Breeding Value. As such, Two-Pool Breeding Value may be a helpful transition for phenotypic programs using One-Pool Breeding Value which intend to move to use of both GS and Two-Pool GCA. With use of GS at high intensity, Two-Pool Breeding Value provided similar gain as Two-Pool GCA after 15 years and was the second-best strategy after 50 years, with slightly reduced gain compared to Two-Pool GCA. However, there is not an apparent logistical advantage to use of Two-Pool Breeding Value with genomic prediction, because programs can choose to predict either GCA or breeding value from identical breeding programs, and GCA is likely the better or not-worse choice. As simulated, Two-Pool Breeding Value had relatively low accuracy with GS but not PS as H_0_ increased, and we note that the low accuracy likely resulted from our choice to predict intra-pool genotypes from a training set of inter-pool genotypes (Moghaddar et al., 2014; Hidalgo et al., 2016). We chose not to explore this further for two reasons. First, this is analogous to prediction of purebred animals from crossbreds, which has been extensively studied (Wei & Van der Werf, 1994; Moghaddar et al., 2014; Hidalgo et al., 2016). Second, although we assumed that intra- and inter-pool phenotyping is achieved with equal thoroughness and accuracy, in practice two-pool plant breeding programs typically would not phenotype intra-pool material for all traits or in as many environments. It is unlikely that increasing intra-pool phenotyping is cost-effective, because the inter-pool genotypes must be thoroughly phenotyped for use as products anyway and this information doubles for use in population improvement. As such, further optimization of Two-Pool Breeding Value with GS was not pursued.

One-Pool Cross Performance is another alternative to Two-Pool GCA. As in previous studies, we observed that at high intensity genomic estimated One-Pool Cross Performance typically outperformed One-Pool Breeding Value with adequate H_0_ (Werner et al., 2020). However, One-Pool Cross Performance underperformed two-pool strategies unless H_0_ was relatively low. As such, One-Pool Cross Performance is a safer option than One-Pool Breeding Value over most H_0_ values and likely benefits programs where creating two pools is infeasible. The underperformance of One-Pool Cross Performance to two-pool strategies was rather small after 15 years, making it an attractive short-term option, but One-Pool Cross Performance was not a substitute for two-pool strategies after 50 years in populations with heterosis.

Somewhat interestingly, both One-Pool Cross Performance and Two-Pool Breeding Value lost competitiveness with use of true values. Compared to genomic estimates, One-Pool Cross Performance lost and Two-Pool Breeding Value gained comparative advantage in accuracy over other values. We hypothesize that use of genomic estimates induced more inbreeding than true values, so Two-Pool Breeding Value lost advantage even as it gained accuracy, because the overall decrease in genetic drift led the two pools to diverge less. Concordantly, we observed that panmictic heterosis decreased in Two-Pool Breeding Value with use of true values compared to genomic estimated values. One-Pool Cross Performance likely lost advantage partly due to the relatively increased accuracy of all other strategies, but perhaps also because with true values it had similar dominance gain but decreased additive gain compared to One-Pool Breeding Value. With genomic-estimated values, One-Pool Cross Performance had increased dominance value compared to One-Pool Breeding Value, likely because it more effectively prevented homozygosity due to inbreeding.
